# Designer Aromatic Cations for Photo-Induced Protein Ligation, Imaging, and Intracellular Labelling at Extended Wavelengths

**DOI:** 10.1101/2025.10.13.681063

**Authors:** Pranab C. Saha, Pooja R. Solanke, Sanwayee Biswas, Vishal Agarwal, Michael T. Taylor

**Author notes:** **Corresponding Author** Michael T. Taylor – Department of Chemistry and Biochemistry, University of Arizona, Tucson, AZ 85701, United States, **Email:**.

## Abstract

Photo-induced protein labelling strategies have become essential tools in chemical biology, but most strategies require high energy wavelengths of light as input to drive reactivity. Recently, we reported a biocompatible method for engaging photo-induced electron transfer to drive protein labelling using biaryl pyridinium salts and, here, we report the design of a series of aromatic cation salts that trigger this process using longer wavelengths of light while maintaining a sterically minimal profile. We achieved this through the systematic study of structure-reactivity relationships of various donor-acceptor pyridinium salts possessing extended conjugation, and these studies revealed the need of a constrained *trans*-stilbene relationship between the probe’s donor and acceptor substituents in order to achieve protein labelling. Probes with chromene-based donor groups in particular showed either robust protein labelling, significant fluorescence quantum yields, or state-dependent photophysical properties; in turn enabling the same probes to be used for both photo-induced protein labelling and wash-free live-cell imaging. We also demonstrate that these enhanced probes possess robust reactivity in complex biological environments through green light-triggered intracellular labelling in live HeLa cells, resulting in the identification of 659 enriched proteins. This series of experiments not only demonstrates the ability of this latest generation of probes to engage in photo-induced labelling using lower energy light in complex proteomes, but also reveals new capabilities for photophysical state-dependent reactivity and measurements.

**Table of Contents:** 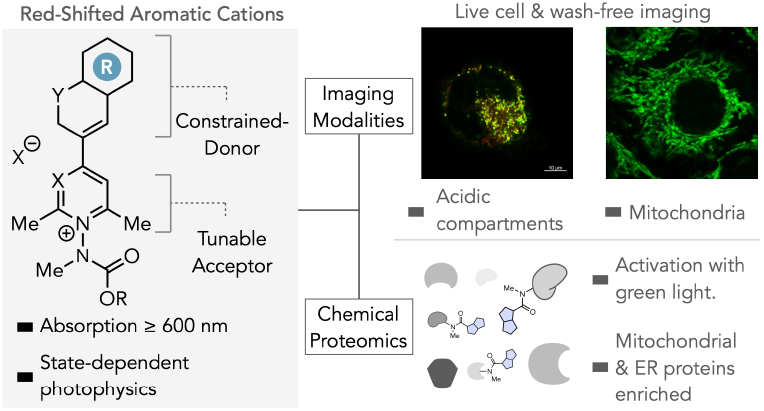

## Introduction

Light mediated chemistries have become integral tools for chemical biology as they offer spatiotemporally resolved access to reactivity that is not accessible from a ground-state. Within this area, photo-chemical processes that are activated by longer wavelengths of light are especially attractive as longer wavelengths of light decrease risks of unintended phototoxicity in living substrates and enhance the capability of light to penetrate further through turbid reaction media^1^. Extensive efforts have been made to develop red-shifted dyes for intracellular and *in vivo* imaging^2^, as well as for optochemical control methods such as photo-cleavable groups^3^, and photoswitches^4^. By contrast, red-shifted variants of photochemical protein ligation reactions have seen less development. Photochemical protein ligation reactions use light to generate high-energy reactive intermediates that drive bond forming reactions; and thus triggering these reactions with longer wavelengths can be challenging as lowering the energetic input of the light frequency may lead to a loss of thermodynamic viability of the reaction. Notable examples of red-shifted protein modification reactions include the work of Yang, who recently developed diazo-coumarin structures that enable diazo-activation using blue wavelengths of light^5a^. Lin has shown that 2,3,4,5-tetrazoles, which can be used for multiple classes of protein ligations, can be decorated with conjugated groups that shift photoactivation from UV-B to violet wavelengths using single-photon excitation^5b^. Recently, Rovis^5c^ and Macmillan^5d^ demonstrated photosensitized cell surface labelling with aryl nitrenes using red light-activated osmium- and tin-containing photocatalysts respectively. Organic dyes, such as rhodamines and silyl-rhodamines have been elegantly used for intracellular photocatalysis, both for protein ligation^5e,f^ and bioorthogonal catalysis^5g^, using green to red light.

We recently reported a photochemical protein ligation approach uses *N*-substituted pyridinium salts to enable photo-induced electron transfer (PET) as a mechanism to drive rapid protein ligation (**Figure 1A**). This approach enabled rapid and efficient protein ligation with high site-selectivity for tryptophan (Trp) residues in peptides and proteins without the need for catalysts or organic cosolvents^6^. Beyond the biocompatibility of the reaction, we demonstrated the ability to establish wavelength-mediated control over reactivity, a capability that we leveraged to develop a visible light-mediated approach that proceeded with enhanced kinetics and milder conditions and enabled for live-cell Tryptophan profiling for the first time^6c^. Building on this work we sought to develop pyridinium probes that further red-shift absorption and in turn offer the potential to further enhance biocompatibility by avoiding excitation of endogenous chromophores, enabling facile compatibility with common microscope laser lines, and lead to the potential for experiments in higher order organisms.

**Figure 1.**
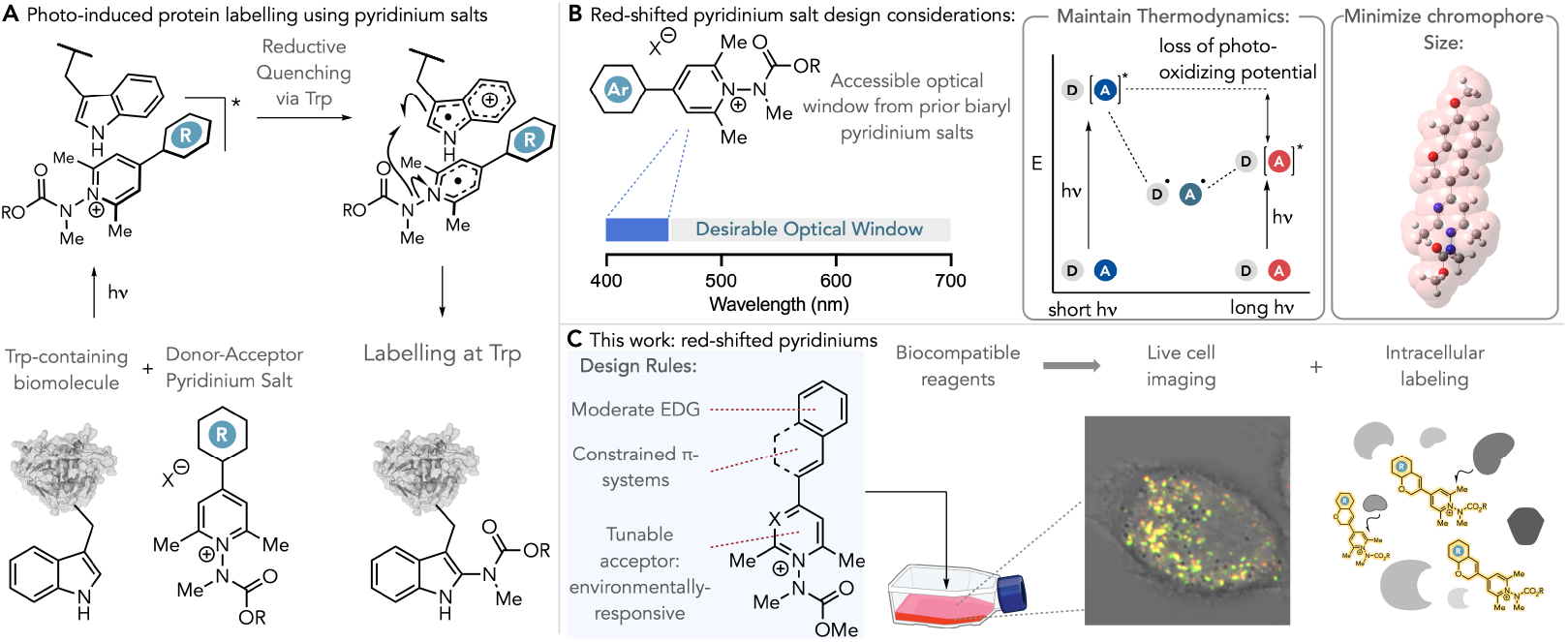
(**A**) Photo-induced electron transfer-driven protein ligation using N-substituted pyridinium salts. (**B**) Design considerations for N-substituted pyridinium salts for protein ligation triggered at longer wavelengths of light. (**C**) This work: design principles and applications of N-substituted pyridinium salts for photo-induced protein ligation at longer wavelengths of light.

However, achieving this in practice poses a number of practical challenges that must be addressed (**Figure 1B**). Traditionally, bathochromic shifts in absorbance of organic chromophores involve extending the π-conjugation to the chromophore; imparting red-shifting but at the same time making the fluorophore larger^7^. A desirable feature of our previous probe generations is their small molecule-like properties and their ease of membrane permeation, which enabled these probes to enter confined spaces within cells and label Trp residues with very low surface exposure. Thus, any future design should seek to preserve these qualities by keeping the chromophore region as small as possible^8^. Additionally, our proposed mechanism of action for these probes is a key Trp->[pyridinium]^*^ reductive quenching *via* photo-induced electron transfer event in which the pyridinium salt acts as a photo-oxidant. Photo-oxidizing potentials can be simply estimated by summation of the reduction half-potential (E_1/2_ [pyridinium]^+^/[pyridinium]^•^) and the lowest energy (E_0,0_) bandgap estimation, these two properties need to be modulated such as to maintain the thermodynamic viability of the key Trp->[pyridinium]^*^ PET event.

*N*-substituted pyridinium salts with red-shifted absorbance compared to **1** are well established in the literature. Structures such as 4-amino-naphthyl-ethenyl pyridinium salts and dialkylaminophenyl ethenyl pyridinium salts have been extensively employed as potentiometric probes for measuring cell membrane potentials^9a,b^ as well as in non-linear optical materials^9c^. These structures are categorized by possessing an aromatic donor linked to the pyridinium acceptor by a stilbene-type arrangement of one or more alkenes, as well as through the use of dialkylamino electron-donating groups.

## Results and Discussion

Using these prior reports as a starting point, we designed conjugation-extended pyridinium salts **2-5** (**Figure 2A**). Salt **2** was synthesized using our previously established route^6a,b^, whilst salts **3-5** were synthesized by adapting knoevenagel condensations between a pyridinium or pyrylium salt and the corresponding carbonyl precursor^10^. Salts **2-5** each possess similar or bathochromically shifted absorption spectra compared to previously reported **1**, ranging from ~350-463 nm in pH 7.4 phosphate buffer. Salts were screened for reactivity against model protein lysozyme (14.3 kDa, 6 Trp, 3 Tyr residues) by irradiating a 10 µM solution of lysozyme and glutathione (GSH) [GSH]= 300 µM) in pH 6.9 NH_4_OAc buffer with salts **2-5** ([probe]=200 µM) for 60 minutes using a Hepatochem PhotoredoxBox™ and a Kessil PR-160 456 nm LED source, followed by direct analysis of the crude reaction mixtures by intact protein mass spectrometry (**Figure 2B**). In each instance, little to no protein labelling was observed, with **3** giving <10% conversion and **2, 4**^10b^, and **5** giving no measurable labelling. Compared to **1, 2-5** possess functional moieties that can potentially quench photo-excited states faster than bimolecular photo-induced electron transfer (PET) with proteins; including very strong electron donating amines (Me_2_N-R, **2**) that can form twisted intramolecular charge transfer states^11^ (TICT), as well as alkenes that can under photoisomerization^12^ (**3**– **5**). These processes can lead to relaxation to ground states *via* non-radiative decay, and this is evidenced by the comparatively low fluorescence quantum yields of these compounds compared to **1**.

**Figure 2.**
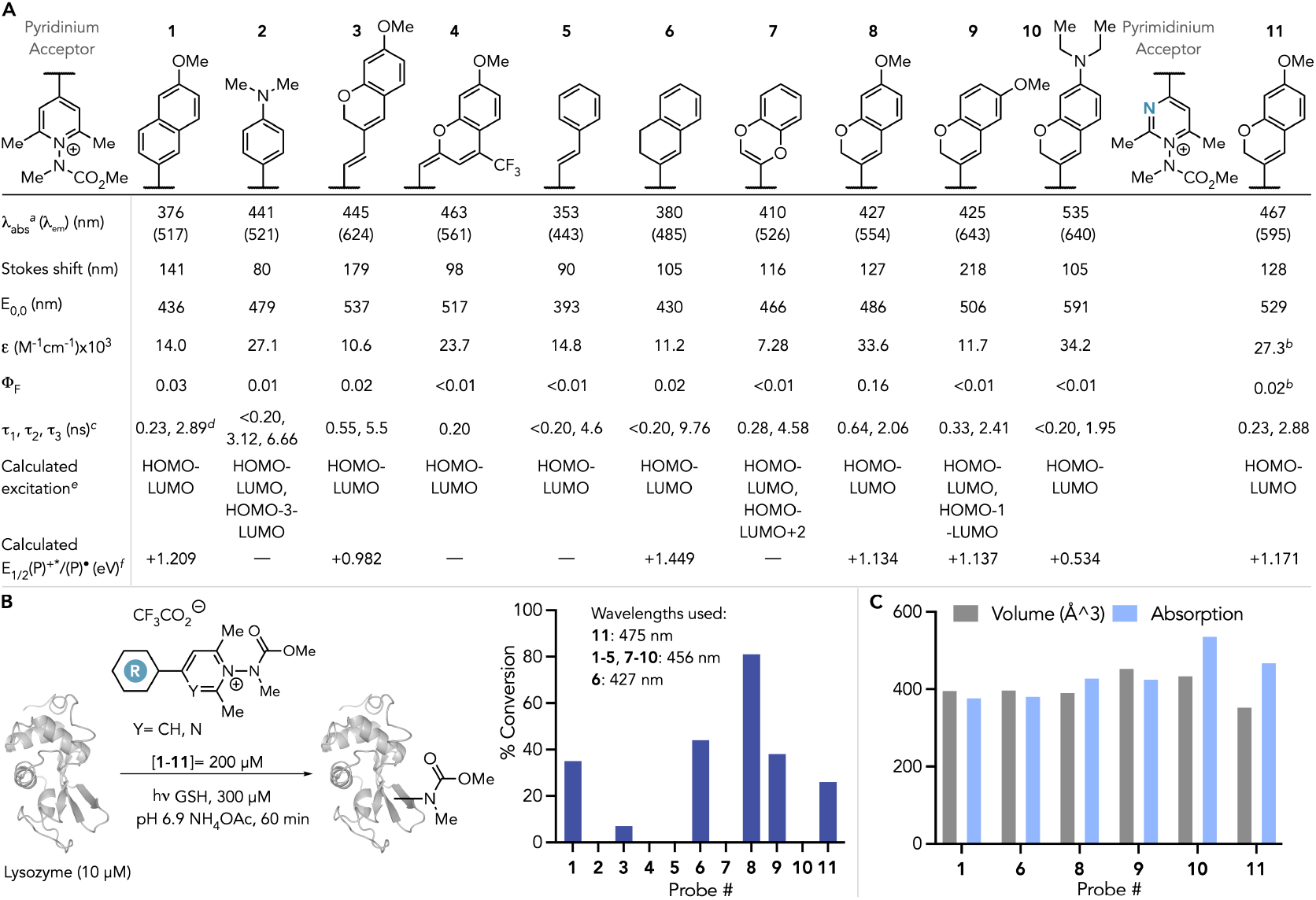
(**A**) Select photophysical and calculated properties of probes **1**-**11**. Values were measured in 20 mM Na_2_HPO_4_ (pH 7.4) except where stated otherwise. (**B**) Labelling of lysozyme with probes **1**-**11**. (**C**) Comparisons of absorption maxima and calculated molecular volume of select probes. ^*a*^Lowest energy transition. ^*b*^Measured in pH 3.5 20 mM Na_2_HPO_4_. ^*c*^Measured in water. ^*d*^from reference 6c. ^*e*^Calculated at the CAMB3LYP 6-31g+(d,p) SMD=H_2_O level of theory. ^*f*^Estimated using experimental E_0,0_ values and calculated reduction potentials^16^. ^*g*^Volume calculated from DFT-optimized structures.

We then posited that a π-system possessing extended conjugation in a conformationally constrained fashion, which has been shown to suppress undesired non-radiative decay processes in other systems^13^, could potentially reveal bathochromic shifts in absorption whilst maintaining the desired PET-driven protein ligation. Thus, we synthesized constrained pyridinium salt **6. 6** retains the stilbenyl-pyridinium chemotype type that can enable bathochromic shifts but constrains this moiety in a 1,2-dihydronaphthyl ring that should inhibit undesired photoisomerization. **6** possesses a comparable absorption window to **1** (λ_max_= 380 nm, 20 mM pH 7.4 Na_2_HPO_4_); enabling photo-activation with visible light sources. Irradiation of lysozyme with **6** under similar conditions as **2**-**5** but using a Kessil PR-160 427 nm LED source yielded a robust 52% conversion to carbamylated lysozyme. This result validated our hypothesis that ring constraint prevents the rate of competitive non-radiative decay pathways, and provided a core stilbenyl-pyridinium scaffold that is photochemically active towards Trp-ligation and could be optimized further for superior optical properties.

**6** was synthesized by an established route used previously^6^ in which the appropriate pyrylium salt is synthesized by Grignard addition of the dihydronapthalene donor group to 2,6-dimethyl-γ-pyrone, followed by condensation with the desired hydrazide (**Figure 3A**). Whilst this route works well for simple groups, such as in **1, 2**, and **6**, many of the structures envisioned for this study possess moieties that are not compatible with this chemistry. This roadblock prompted us to re-evaluate synthetic procedures for accessing these probes. In our general design of pyridinium salts (shown in **Figure 3A**), we divide the probe into three modular constituents: (1) a tunable donor group (2) the aromatic cation acceptor and (3) the carbamate group that ultimately forms covalent bonds with the biomolecule (referred to here as the “transferring group”). Synthetic routes that enable maximum interchangeability between these substituents are desirable as components could be mixed and matched to easily access probes with a particular set of properties. A more ideal synthetic approach would be to form the bond linking the donor π-system to the pyridinium acceptor *via* cross-coupling. Thus, we designed a general synthetic route in which an electrophilic amination reaction is used to form the pyridinium ring from a bespoke pyridine precursor that would first be assembled *via* cross-coupling chemistry. Briefly, either the appropriate vinyl halide or B-pin ester could be used for Suzuki-Miyaura cross-coupling with (2,6-dimethylpyridine-4-yl)boronic acid or converted to a boronic ester and coupled with 4-chloro-2,6-dimethylpyrimidine (shown with a chromene donor ring in Figure 3B for simplicity, see supplementary info section **2** for specific structures and conditions). Next, electrophilic amination of the pyridine/pyrimidine core with *O*-tosylhydroxylamine^14^ smoothly afforded the corresponding *N-*amino pyridinium tosylate salt in 52-74% yield. Building on our recent work involving selective pyridinium ylide-alkylation^15^, we developed a tandem acylation/alkylation sequence using the appropriate activated NHS-carbonate as a carbamylating agent and methyl iodide as the electrophile to install the requisite *N,N*-disubstituted carbamate and ultimately yield the target probe. Using this approach, we synthesized pyridinium salts **7**-**11**. Salt **7** features a benzodioxine donor core^16^ whilst salts **8**-**11** feature chromene donor cores with varying substitution patterns. This approach also enables modular manipulation of the carbamate transferring group beyond simple methyl groups (**7-11**, Figure 2A) to readily include relevant application-specific functional groups (see **8a, b, 9a, 11a**, Figure 6).

**Figure 3.**
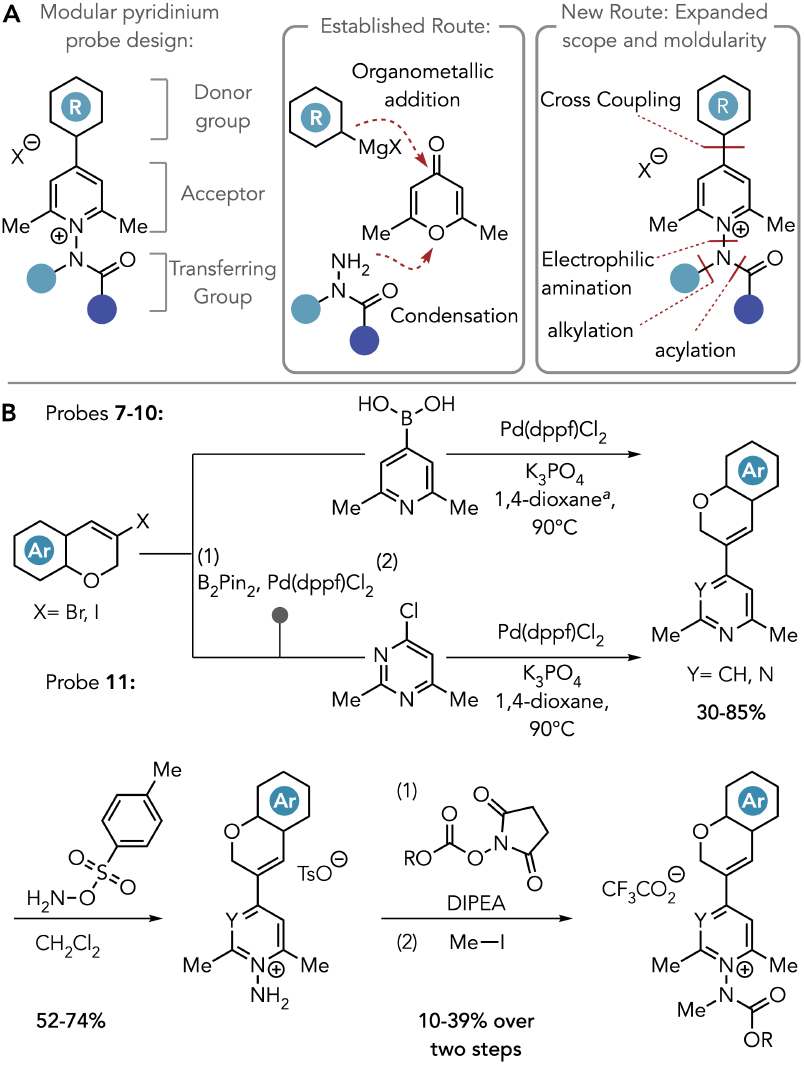
(**A**) Retrosynthetic strategies for pyridinium salts **B**) General synthetic strategy for pyridinium probes **7**-**11**. An logs **8a-11a**, and **8b** were accessed using this route (Figure 6).^*a*^DMF was used for **10**.

Having accessed this bank of pyridinium salts, we next collected photophysical data for each analog, with the results summarized in Figure 2A. Each analog synthesized, except for **5**, possesses absorption spectra that are bathochromically shifted compared to **1**. Though minimally active towards Trp-labelling, salts **2**-**4** each possess λ_max_ between 440-470 nm. While dihydronaphthalene salt **6** possesses a similar absorption spectrum to **1**, evolution of the donor unit from a dihydronaphthalene donor to analogs featuring heteroatom-based donating groups imparts significant bathochromic shifts in absorption. For example, benzodioxine-substituted salt **7** possesses an absorption maximum of 410 nm. 7-methoxychromene substituted pyridinium salt **8** is nearly 50 nm red-shifted (λ_max_ = 427 nm) compared to **1** whilst 6-methoxychromene analog **9** possesses a similar shift (λ_max_ = 425 nm). Notably, **9** has a much lower fluorescence quantum yield compared to **8**, and TDDFT calculations suggest that shifting this group changes the electronic transitions from a primarily HOMO-LUMO excitation to a HOMO-1-LUMO excitation mode. Addition of a strongly electron-donating diethyl amino group in pyridinium salt **10** further bathochromically shifts absorption to well beyond 500 nm (λ_max_ = 535 nm) and by 159 nm compared to **1**. Both in biaryl pyridinium salts such as **1** and **2**, as well as structures possessing the trans-stilbenyl structural motif such as **3**-**11**, calculations of electronic structure clearly indicate a donor-acceptor relationship between the aryl/styryl group and the aromatic cation acceptor ring, thus, perturbations at the aryl/styryl site more significantly affect HOMO orbital energies. To explore perturbations of the acceptor, we synthesized pyrimidinium-containing salt **11**. This highly electron-deficient ring significantly lowers acceptor (LUMO) energies instead of donor (HOMO); with **11** LUMO lowered by 0.233 eV and results in a bathochromic shift in relation to their pyridinium analogues (λ_max_ = 467 nm for **11** vs. 427 nm in **8**). Notably, these designs represent a gain in efficiency by measure of absorbance versus the calculated chromophore volume. For example chromophore of **8**, which is bathochromically shifted by 51 nm compared to **1**, is calculated to be approximately the same volume as **1**, whilst **11** is bathochromically shifted by 91 nm compared to **1** but is calculated to be 33 Å^3^ smaller (**Figure 2C**). In short, probes **8**-**11** shift probe absorbance to longer wavelengths without sacrificing the small molecule-like properties of the probes.

With this series of probes in hand, we next assayed them for labelling against lysozyme using the same conditions as described previously. Out of this series, probes **8, 9**, and **11** showed robust labelling; with **8** showing 81%, **9** showing 37%, and **11** showing 26% (**Figure 2B**). Peptide mapping studies of lysozyme, as well as chymostrypsinogen labelling with **8** confirmed selectivity for Trp labelling on both proteins (Lysozyme: W62, W108, chymotrypsinogen: W237, see supplementary info section 3 for experimental details). Probes **7** and **10** did not show any meaningful lysozyme labelling. As with **2, 10** possesses a strongly donating dialkylamino group, further reinforcing the importance of moderate electron-donating groups in ensuring photochemical reactivity whilst moderating calculated photooxidation potentials^16^ (calculated E_1/2_ **6** = +1.440 vs. **8**= +1.134 eV). Probes showing labelling also maintain mild calculated photo-oxidation potentials ranging between 0.9 eV to 1.4 eV, with the most active **8, 9**, and **11** (see fig 4D) ranging of ~+1.1 eV; in excess of experimental Trp oxidation potential (E^0^=+1.077 eV vs. NHE @ pH 7)^17^. By contrast, the strongly donating diethylamino group of **10** reduces the calculated photo-oxidation potential by 600 mV compared to **8**.

**Figure 4.**
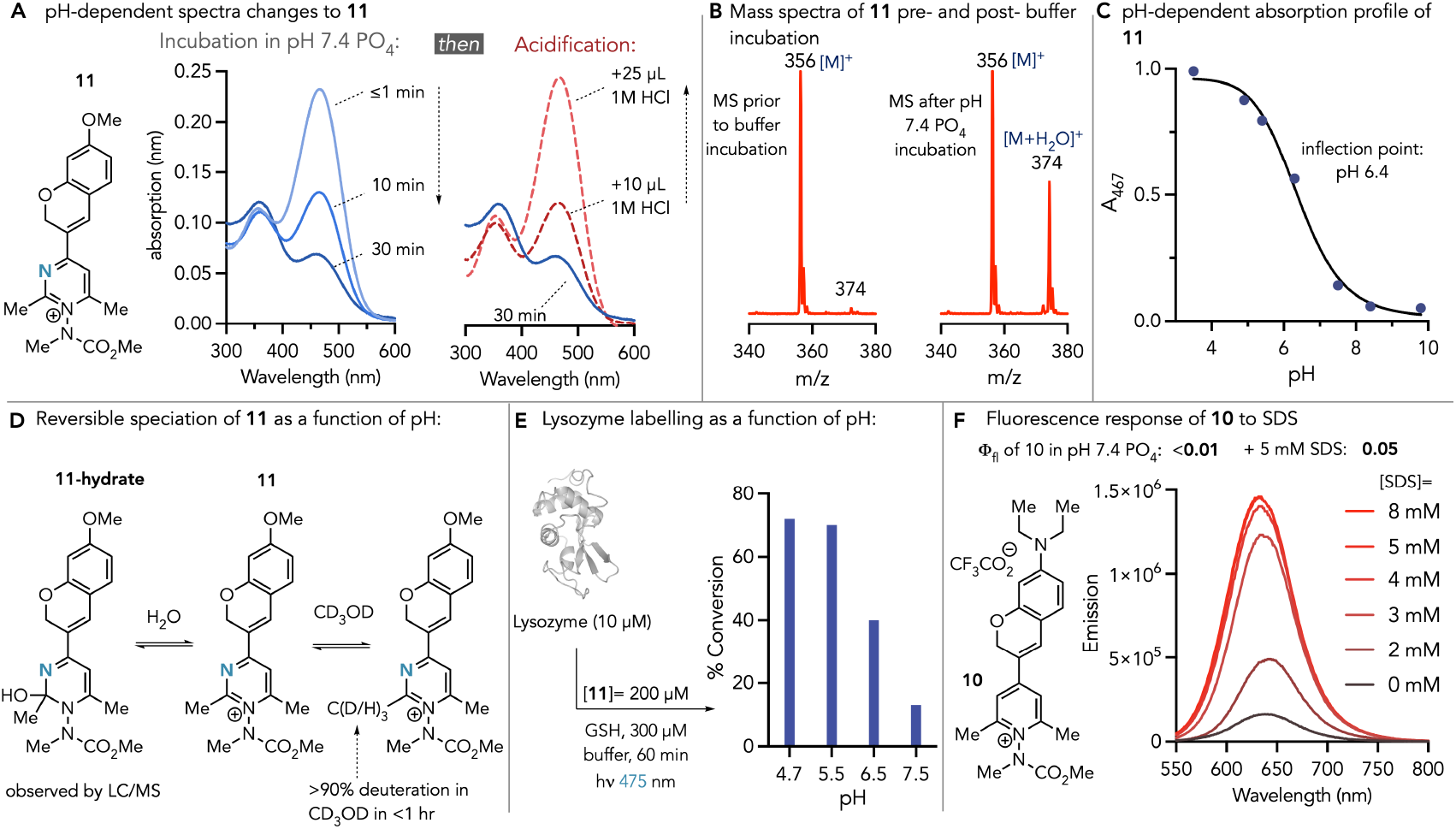
(**A**) **11** displays reversible, pH-dependent changes to electronic structure. (**B**) Observation by LC-MS of hydrate formation of **11** after incubation in pH-7.4 Na_2_HPO_4_ buffer. (**C**) pH-dependent changes to absorption at 467 nm. (**D**) Proposed mechanism of pH-sensing by **11** *via* reversible, pH-dependent, hydrate formation. (**E**) pH-dependent labelling of lysozyme using **11**. (**F**) Turn-on fluorescence effect of **10** in sodium-dodecyl sulfate (SDS).

Whilst **10** and **11** show reduced labelling activity compared to **8** and **9**, these probes each show state-dependent photophysical properties (**Figure 4**). **11** features a pyrimidinium acceptor ring that is highly electron deficient in nature. As a result, the benzylic 2-methyl group of the pyrimidinium ring undergoes rapid C-H/C-D exchange in methanol (>90% in <1 hr). Previous studies, including ours, have shown that this process can occur in the presence of mild bases, and the observation of this exchange in solely protic solvent reveals the relatively acidic nature of this position on the probe^10,14^. Beyond H/D exchange, **11** also shows pH-dependent photophysical behavior. Upon initial dissolution in pH 7.4 phosphate buffer, the absorption spectra of a 20 µM solution of **11** shows an absorption maxima at 467 nm and minor maxima at 354 nm. Monitoring absorption over time reveals a steady decrease in the intensity of the 467 nm maxima which reaches equilibrium at 30% of the initial intensity after 30 minutes (**Figure 4A**). Given our observations with H/D exchange, we posited that this effect was pH-dependent, which was validated by restoration of the original 467 nm maxima through acidification with HCl(aq). Analysis of a solution of **11** in pH 7.4 PO_4_ after 30 minutes by LC/MS using a short chromatography method showed the formation of a new peak with a mass of **11** + 18 mass units; suggesting the formation of a hydrate of **11** (**11-**hydrate) (**Figure 4B, D**). In **11**-hydrate, the conjugation of the pyrimidinium ring is broken, resulting in the loss of the 467 nm transition band. We then measured absorption spectra of **11** in phosphate buffer at various pH’s ranging from 3.5-9. and this study revealed that **11** and **11**-hydrate exist in a pH-dependent equilibrium, with the **11**/**11**-hydrate k_eq_=1 observed at pH 6.4 (**Figure 4C**). Reinvestigating protein labelling with **11** revealed pH-dependent protein labelling, with higher conversions at pH 4.7 and 5.5 (72% and 70% respectively), and a marked drop in conversion at pH 6.5 and 7.4 (40% and 13% respectively) (**Figure 4E**).

In contrast to **11**, which is highly electron-deficient, **10** possesses a strongly donating diethyl amino group that makes this compound electron rich in comparison to **11**. Fluorogenic donor-acceptor compounds, including those that possess strong donating groups such as dialkylamino aryl moieties, are highly sensitive to environmental changes arising from changes to stabilization if different excited states and charge-transfer states^11^. To understand how **10** responds to these changes, we measured absorption and emission of 20 µM solution of **10** in pH 7.4 phosphate buffer and varying amounts (0-8 mM) of sodium dodecyl sulfate (SDS) (See supplementary info figure S16, **Figure 4F**). As expected with aromatic cations, absorbance displayed a bathochromic shift of the lowest energy transition absorption maxima by 20 nm between 0 mM (533 nm) and 8 mM SDS solutions (553 nm). Analysis of emission spectra of **10** at varying concentrations of SDS reveals a turn-on fluorescence effect: **10** does not emit sufficiently in the pH 7.4 Na_2_HPO_4_ at 0 mM SDS to measure a fluorescence quantum yield with the available instrumentation; but Φ_Fl_=0.05 was measured after addition of 5 mM SDS.

Next, we sought to probe the behavior of select aromatic cation probes in complex biological environments. Since several analogs possess inherent fluorescence, as well as potentially useful state-dependent changes to photophysical and structural properties, we performed cellular localization studies *via* confocal microscopy to assess cell penetration and to assess if the state-dependent photo-physical properties of analogs **10** and **11** could enable localization and imaging of orthogonal sub-cellular compartments. We selected probes **8, 10**, and **11** for these studies owing to their significant quantum yield (**8**), their dynamic response to stimuli (**10, 11**), as well as their convenient absorption/emission profiles excitation with common confocal laser lines (**Figures 5A-C**). Thus cultured HeLa cells were incubated with 10 µM of either **8, 10**, or **11** for varying periods of time, and then either washed or imaged directly without washing depending on the probe and experiment. (**Figure 5D**). HeLa cells were incubated first with mitotracker red, followed by washing with PBS, and then with Pyridinium probe **8** for 20 minutes. Imaging was performed directly without further washing and revealed that **8** readily crosses cell membranes and displays significant colocalization with mitotracker red (Pearson Correlation = 0.82) (**Figure 5E**). The low-energy transition of dialkylamino probe **10** features a wide charge-transfer band with significant absorption beyond 600 nm, thus enabling use of the 561 nm laser line for excitation (**Figure 5B**). Incubation of HeLa cells with **10** for 15 minutes reveals that **10** also rapidly crosses membranes and remains localized in mitochondria post-washing (**Figure 5F**). However, given that **10** is essentially non-fluorescent in phosphate buffer but demonstrates polarity-dependent fluorescence turn-on (**Figure 4F**), we also demonstrated **10** allows for wash-free imaging. These experiments revealed that, under wash-free conditions, **10** displays measurable fluorescence in multiple cellular compartments.

**Figure 5.**
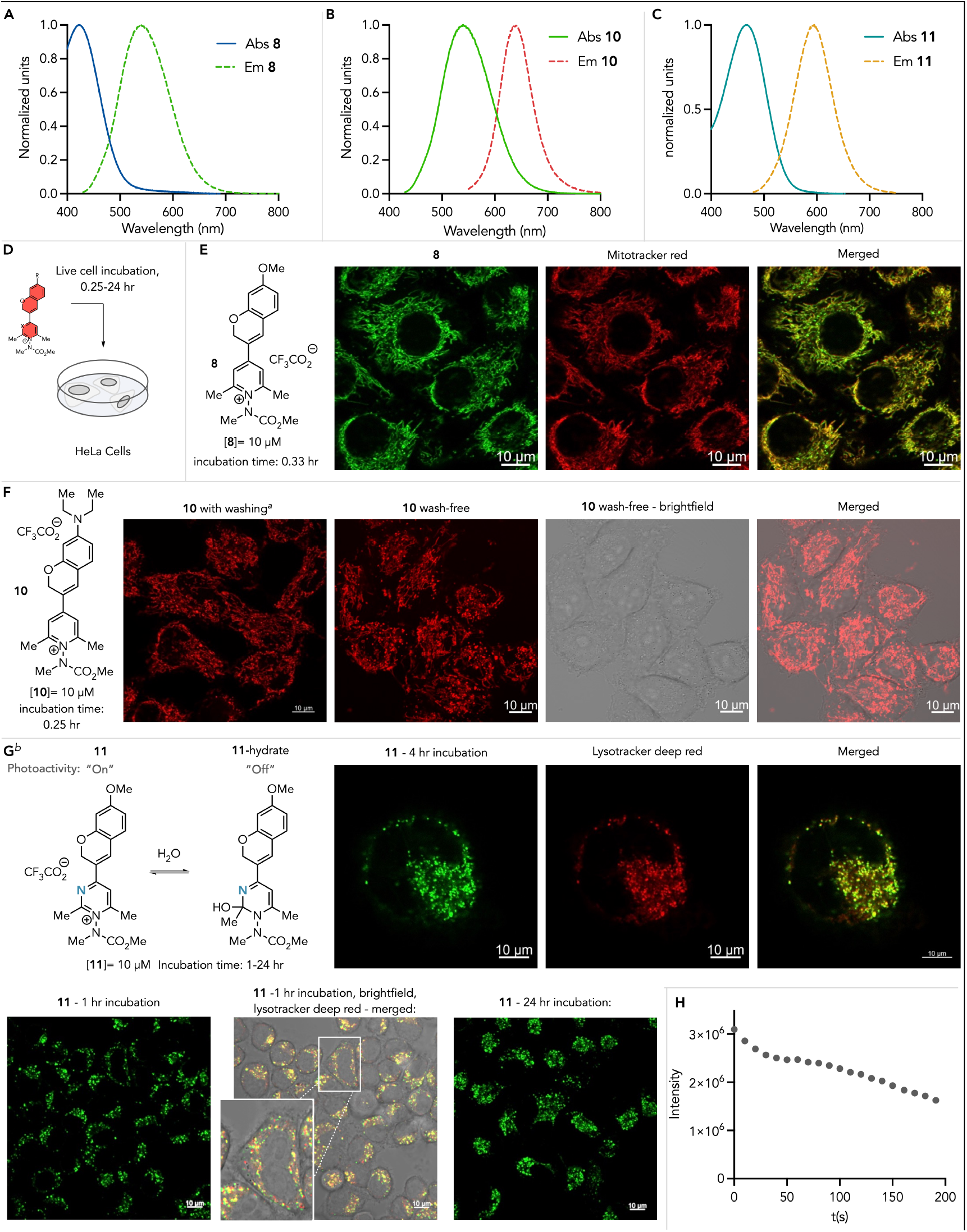
Absorption/emission spectra of pyridinium probes in 20 mM pH 7.4 NaH_2_PO_4_: (**A**) **8** (**B**) **10** (**C**) **11**. (**D**) Confocal imaging of live HeLa cells using **8**-**11**. (**E**) Live cell confocal imaging using **8** (λ_ex_=405 nm). (**F**) Live HeLa cell imaging under wash and wash-free conditions using **10** (λ_ex_=561 nm). (**G**) Wash-free live cell imaging using **11**. (λ_ex_ =488 nm) (**H**). Photobleaching profile of **11** using 488 nm excitation at 5% intensity (25 mW, Zeiss LSM 880 Argon laser). ^*a*^Cells were washed 3x with PBS buffer. ^*b*^Wash-free imaging.

Next, we explored live-cell imaging using **11**. Intracellular compartments vary by up to 3 orders of magnitude in pH^17^, whilst the HeLa cell cytosol pH is estimated to be 7.34^18^. In **11**-hydrate, chromophore conjugation is broken, resulting in **11**-hydrate not absorbing visible light. Given that the hydrate is the major species at pH>6.4, excitation of cells using a 488 nm laser should preferentially illuminate acidic subcellular compartments where it is most likely that detectable quantities of fully conjugated cation **11** will be present. Thus, HeLa cells were incubated with 10 µM **11** for 1-24 hrs followed by lysotracker deep red incubation for 15 minutes (**Figure 5G**). Dual color imaging was performed directly without washing (**11** excitation: 488 nm, lysotracker – 633 nm) and revealed significant co-localization between the two fluorophores (Pearson Correlation = 0.82) and leads us to propose that **11** can be used as a means of imaging microbodies in live cells. Additionally, given that the **11**-hydrate “off” state does not absorb 488 nm light, imaging can be conveniently conducted under wash-free conditions. There are many classes of dyes and fluorogenic compounds that show pH-responsive photophysical properties, and the primary mechanism by which these act is through direct coupling of changes in photophysical properties/electronic states to changes in protonation states as a function of pH^19^. **11** complements this mechanism of established pH sensors as **11**’s pH-dependent behaviour arises from changes to hydrate-**11** equilibrium as a function of pH as opposed to a direct Brønsted acid-base reaction. This method of lysosomal imaging also directly complements lysosome-targeted probes, exemplified by the lysotracker series, which localizes in lysosomes *via* amine protonation. Extended incubation times with structures that localize in lysosomes through amine protonation have been shown to raise lysosome pH and therefore alter morphology^20^. **11** directly complements these structures by allowing observation of lysosomes over extended incubation times, with up to 24 hrs incubation being demonstrated and could potentially enable studies of acidic cellular compartments over longer time courses. A photobleaching time course of **11** is shown in Figure 5H and shows a steady decrease in signal vs. irradiation pulses and is representative of the rational design of these probes for photo-induced reactivity. Despite this photolability, **11**, still maintains sufficient fluorescence for measurement purposes, even after continuous pulsing over three minutes (see supplementary video).

Finally, we sought to demonstrate proteome profiling using the probes developed here. Chemical proteomic profiling has emerged as a valuable new tool for discovering new druggable targets, functionally essential sites on proteins, and monitoring state-dependent changes to the proteome^21^. We demonstrated previously that *N*-substituted pyridinium probes designed around **1** enabled proteome profiling with violet light triggering of traditionally challenging sites (e.g. Trp) both at the lysate and live cell level for the first time and methods targeting this residue remain rare^6c,22^. Demonstrating proteome profiling with the pyridinium probes developed here that are activated at longer wavelengths of light should enable milder labeling conditions that can expand future applications^1^. We therefore synthesized biotinylated analogs of **8a, 9a**, and **11a** using the modular synthetic route shown in Figure 3 and a biotinylated NHS-activated carbonate (**S2**, see supplementary info for specific reagents/conditions). To understand what, if any, effect on photophysics these groups have on the pyridinium structures, we also measured select photophysical properties of **8a** and azide-derived **8b** (synthesized from activated carbonate **S1**, see supplementary info section 1) (**Figure 6A**). These measurements revealed that the biotin group of **8a** slightly increases the stokes shift of the probe and increases quantum yield by 30% (Φ_Fl_= 0.16 for **8**, Φ_Fl_= 0.21 for **8a**), whilst the azido-alkyl group of **8b** imparts more modest effects (Φ_Fl_= 0.17).

**Figure 6.**
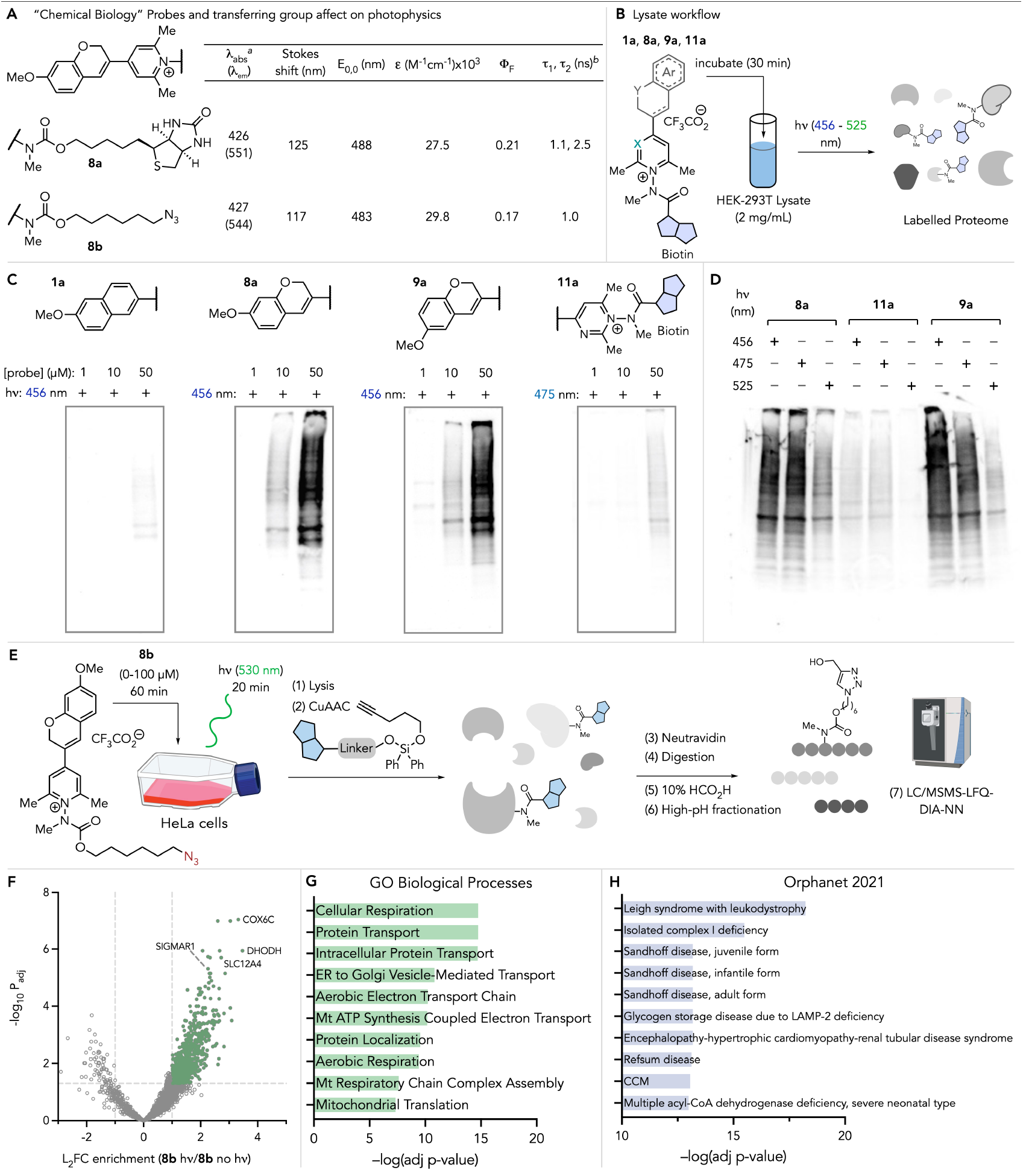
(**A**) Photophysical properties of chemical proteomic pyridinium probes **8a** and **8b** in 20 mM Na_2_HPO_4_ (pH 7.4). (**B**) Proteome profiling of HEK293T lysates using biotinylated probes. (**C**) Western analysis of proteome HEK293T lysate profiling with probes **1a, 8a, 9a**, and **11a**. (**D**) Proteome profiling with **8a, 9a**, and **11a** at 456 nm, 475 nm, and 525 nm irradiation wavelengths. (**E**) Live HeLa cell culture labelling using **8b** and 530 nm irradiation. (**F**) Volcano plot showing differential expression of 659 protein entities. (**G**) Top 10 biological processes from EnRichr GO analysis of enriched proteins. (Mt=mitochondria) (**H**) Top 10 orphan disease associations of enriched proteins. ^*a*^Lowest energy transition. ^*b*^Measured in water.

Next, HEK293T cell lysates (2 mg/mL) were treated with biotinylated pyridinium/pyrimidinium salts **1a, 8a, 9a**, or **11a** (1, 10, and 50 µM) for 30 minutes followed by irradiation for 20 minutes with either 456 nm (**1a, 8a, 9a**) or 475 nm (**11a**) LED’s (**Figure 6B**). Labelled proteomes were then analyzed by western analysis to detect biotinylated proteins (**Figure 6C**). Using these conditions, **1a** gives only low levels of labelling at the highest probe concentration (50 µM) as it absorbs little light beyond 440 nm. By contrast **8a** and **9a** give robust labelling at 10 µM probe concentrations and high levels of labelling at 50 µM concentrations. Given the propensity of **11** to form photo-inert hydrates at pH 7.4, we anticipated similar behavior from **11a**, which was confirmed by only low levels of labelling observed at 50 µM probe concentration. **8a, 9a**, and **11a** each possess significant absorption beyond 450 nm, so we next explored lysate profiling using different wavelengths of light (456, 475, and 525 nm) for photoactivation (**Figure 6D**). As expected, **11a** shows comparatively low levels of labelling for each wavelength, but we observed robust labelling for **8a** and **9a** at 456 nm and 475 nm. Gratifyingly, we also observed robust labelling when a green LED (525 nm) was used for photoactivation of **8a** and **9a**.

With these data in hand, we then developed a chemical proteomics workflow to establish labelling at the cellular level using azide-functionalized probe **8b**. HeLa cell cultures in T75 culture flasks were incubated for 60 minutes with **8b** ([**8b**] = 0, 50, 100 µM) at 37 °C, followed by photoactivation using a 96-LED array equipped with green LEDs (Amuza, Inc., 530 nm) for 20 minutes at 4 °C. After lysate generation, CuAAC was performed to derivatize labelled sites with an acid-cleavable biotin group^23^. Direct analyses of these workflows *via* western analysis indicated a clear dose-response between 50-100 µM **8b**, with controls with 100 µM but no light-activation showing that photocontrol is both maintained in complex cellular environments but also extended now to the milder green light wavelengths (see supplementary info Figure S29). After biotinylation with CuAAC, samples were also subjected to a chemical proteomics workflow for protein-level enrichment in which samples were incubated with neutravidin resin, followed by on-bead digestion, tryptic peptide cleanup, and fractionation at basic pH. Tryptic peptides were then analyzed using LC/MSMS using data-independent acquisition (DIA). Experiments were performed in biological triplicate and data was analyzed using the Fragpipe software suite^24^ using the DIA-NN workflow^25^ with closed-searching and label-free quantification with variance stabilization normalization of samples. Analysis in this way led to the identification of 659 differentially enriched proteins (log_2_-fold enrichment ≥1) between photoactivated **8b** and **8b** incubation with no photoactivation (**Figure 6F**).

Significantly enriched proteins were searched against the UniProt database for subcellular localization data, which revealed 32% of proteins are associated with the mitochondria, 24% with the ER, and 20% with the nucleus (see supplementary info, Figure S30). 62% of enriched proteins have associations with membranes. This data generally agrees with our imaging studies with **8a** in Figure 5E, but complements proteins enriched with probe **1** previously by us^6c^, which enriched primarily nuclear proteins. Gene Ontology (GO) analysis using Enrichr^26^ also reflects imaging data and shows enrichment of genes comprising mitochondrial, endoplasmic reticulum, and Golgi-related cellular components (SI Figure S31), as well as key metabolic processes that take place in the mitochondria as well as in protein transport (**Figure 6G**). Outside of dysfunction induced by ischemic events, mitochondrial diseases are primarily genetic in nature and, owing to the infrequency of these diseases, are generally considered orphan diseases^27^. Analysis of the Orphanet database with Enrichr reveals proteins enriched associated with mitochondrial orphan diseases including, Leigh Syndrome, isolated complex I deficiency, and congenital cataract – hypertrophic cardiomyopathy – mitochondrial myopathy (CCM) (**Figure 6H**). Thus, by using **8b**, we are able to perform live cell labelling at longer wavelengths of light compared to our previous designs (green for **8b** vs. violet for **1**) whilst also enriching highly complementary proteomic subsections.

## Conclusion

In summary, we have developed a suite of photoactivated aromatic cation probes for protein labelling that possess red-shifted absorption properties. Key to this development was the realization through structure-reactivity relationships of the structural prerequisites required to both preserve photoreactivity and red-shift absorption. These studies revealed several viable chromene-pyridinium and pyrimidiminium salts for photo-induced protein labelling at longer wavelengths of light. We also show that the probes developed here possess sufficient brightness for use in live cell imaging, and the state-dependent photophysics that some of the probes possess, especially **10** and **11**, can enable wash-free imaging of multiple subcellular locales. Furthermore, we demonstrated live cell labelling experiments triggered by green light. Taken together, we anticipate that the expanded capabilities provided by this series of aromatic cation probes will enable access to protein conjugates through low energy photo-conjugation as well as intracellular experiments in which multiple analytical techniques and experimental workflows can be harnessed through identical structures.

## Supporting information

Experimental Procedures

Computation Details

Proteomics Data Tables

Imaging_11_Video

## ASSOCIATED CONTENT

### Supporting Information

Supporting information includes experimental procedures for probe synthesis, protein modification experiments, photophysical studies, imaging, intracellular labelling, and small molecule NMR data. A second document contains computational analysis procedures and data. A third spreadsheet contains proteomic profiling data tables. A fourth file consists of a .AVI video of HeLa cell imaging with **11**.

The Supporting Information is available free of charge on the ACS Publications website.

## AUTHOR INFORMATION

### Authors

**Pranab C. Saha** – Department of Chemistry and Biochemistry, University of Arizona, Tucson, AZ 85701, United States

**Pooja R. Solanke** – Department of Chemistry and Biochemistry, University of Arizona, Tucson, AZ 85701, United States

**Sanwayee Biswas** – Department of Chemistry and Biochemistry, University of Arizona, Tucson, AZ 85701, United States

**Vishal Agarwal** – Department of Chemistry and Biochemistry, University of Arizona, Tucson, AZ 85701, United States

### Author Contributions

PCS synthesized, characterized, and assessed probes **1, 6, 7, 8, 8a, 8b, 11**, and **11a**. PRS synthesized, characterized, and assessed probes **5, 9**, and **9a**, and developed the synthetic route to chromene precursors. SB synthesized, characterized, and assessed probes **2** and **10**. VA synthesized, characterized, and assessed probes **3** and **4**. SB performed SDS-based fluorescence studies with **10**. PCS performed pH-dependent studies with **11**, performed all live-cell imaging experiments, western analyses, and chemical proteomics experiments. MTT conceived and managed the project, designed, and analyzed experiments, interpreted data, and performed calculations. MTT wrote the manuscript with contributions from all authors. All authors wrote the supplementary information. All authors have given approval to the final version of the manuscript.

### Notes

The authors declare no competing financial interest.

## ACKNOWLEDGMENT

We thank R35 GM143120 for supporting this work. We acknowledge the W.M. Keck Center for Nano-Scale Imaging (RRID:SCR_022884) and the Nuclear Magnetic Resonance Facility (RRID:SCR_012716) at the University of Arizona Department of Chemistry & Biochemistry for instrumentation support. We also thank NSF (grant no.’s 1920234, 840336, 9214383, 9729350) for additional instrumentation support. We thank the University of Arizona for startup funds.

