## Supplementary material for "Designer Aromatic Cations for Photo-Induced Protein Ligation, Imaging, and Intracellular Labelling at Extended Wavelengths": Experimental Procedures

**Table of Contents**


1. **General Considerations** ……………………………………………………… S3–S4
2. **Small Molecule Synthesis Procedures** ………………………..……………S5–S38
3. **Protein labeling and mapping**………………………………………………… S39–S46
4. **Photophysical Properties of pyridinium and pyrimidinium probes**……S47–S60
5. **Live cell imaging with pyridinium and pyrimidinium probes using confocal microscopy** …………………………………………………………… S61–S70
6. **Chemoproteomic Profiling Analysis, Procedures, and Validation**..……S71–S82
7. **References**………………………………………………………………………… S83
8. **NMR Spectra**……………………………………………………………………… S84–S160

### 1. General Considerations

Nuclear magnetic resonance (NMR) spectra were acquired at ambient temperature unless otherwise stated using either a Bruker Avance III 400 and a NEO 500 at the University of Arizona Nuclear Magnetic Resonance Facility. Chemical shifts (δ) are reported in ppm and coupling constants (J) are reported in Hz. Data are reported in the following format: Chemical shift (multiplicity, coupling constants, number of protons). The following convention is used to report multiplicity: s = singlet, d = doublet, t = triplet, q = quartet, qn = quintet, sext = sextet, sept = septet, m = multiplet, br = broad. $^{19}$F-NMR spectra were recorded using fluorobenzene ($C_6H_5F$; δF = – 115.3 ppm in acetonitrile-D3) as an internal standard. Many of the pyridinium salts exist as rotamers. Where possible, NMR signals are assigned integration values equal to the number of protons associated with the given functional group of the given rotamer. Signals where rotameric peaks are overlapping are assigned hydrogens based directly on their integration values. Infrared (IR) spectra were collected on a Thermo Fisher Scientific NICOLET iS50R FT-IR equipped with an ATR probe at the Keck center, University of Arizona. Absorptions are reported in wavenumbers ($cm^{-1}$). Analytical thin layer chromatography (TLC) was performed using Merck pre-coated glass backed silica gel plates (TLC Silica gel 60 F254). TLC plates were visualized using either UV-light (254 nm), potassium permanganate or ceric ammonium molybdate staining solutions. Flash column chromatography was performed under positive pressure of compressed air using SiliCycle SiliaFlash® P60 silica gel (230-400 mesh) or using an ISCO Combiflash 300. High-resolution mass spectra were acquired at the analytical & biological mass spectrometry facility at the University of Arizona. Absorption spectra were acquired on a Jasco V-760 spectrophotometer. Emission spectra, fluorescence lifetime and absolute quantum yield (using an integration sphere) were acquired on a Horiba FluoroMax Plus fluorescence spectrophotometer. Gels were visualized using a UVP ChemStudio Plus imaging instrument. Reactions were performed in heat-dried glassware using appropriate Schlenk techniques. The reagents purchased from commercial sources were used as received. 2,4,6-trimethylpyrylyium tetrafluoroborate[1] and methyl 1-methylhydrazine-1-carboxylate[2] were synthesized according to the previously reported procedure. 4-(6-methoxynaphthalen-2-yl)-2,6-dimethylpyrylium tetrafluoroborate was synthesized as reported previously.[3] 2,5-dioxopyrrolidin-1-yl methyl carbonate was synthesized adapting the reported literature.[4] Lysozyme (from chicken egg white, >40,000units/mg of protein), and α-Chymotrypsinogen A from bovine pancreas (essentially salt free, lyophilized powder), were purchased from Sigma Aldrich.

**General LC/MS methods:**

**Method A**: solvents: A: 0.1% formic acid in $H_2O$ B: 0.1% formic acid in $CH_3CN$, method: 10-50% B over 8 min, 70-90% B over 1 min, hold 90% B for 1 min, 90-10% B over 1 min, then hold 10% B 1 min, column: Kinetex® 1.3 µm C18 100 Å, LC Column 50 x 2.1 mm, m/z range: 500-2000, flow rate: 0.3 mL/min.

**Method B**: solvents: A: 0.1% formic acid in $H_2O$ B: 0.1% formic acid in $CH_3CN$, method: hold 10% B for 0.5 min, 10-40% B over 10.5 min, 40-90% B over 1 min, hold 90% B for 1.5 min, 90-10% B over 0.5 min, then hold 10% B 1 min, column: Kinetex® 1.3 µm C18 100 Å, LC Column 50 x 2.1 mm, m/z range: 500-2000, flow rate: 0.3 mL/min.

**General HPLC methods:**

**Method A**: solvents: A: 0.1% TFA in $H_2O$, B: 0.1% TFA in $CH_3CN$, method: hold 5% B 1 min, 5-30% B over 26 min, hold 30% B 8 min, 30-35% B over 5 min, 35-95% over 3min, hold 95% B 4 min, 95-10% B over 2 min, hold 10% B 3 min, UV-Vis: 390, 365, 254 nm, column: Kinetex® 5µm C18 100 Å, LC Column 150 x 21.2 mm, flow rate: 10 mL/min.

**Method B** solvents: A: 0.1% TFA in $H_2O$, B: 0.1% TFA in $CH_3CN$, method: hold 10% B 1 min, 10-45% B over 25 min, hold 45% B 5min, 45-95% B over 3 min, hold 95% B 3 min, 95-10% B over 2 min, hold 10% B 3 min, UV-Vis: 390, 365, 254 nm, column: Kinetex® 5µm C18 100 Å, LC Column 150 x 21.2 mm, flow rate: 10 mL/min.

**General reversed-phase flash chromatography method:**
**Method A:** Tedelyne ISCO 300 Instrument method: solvents: A: 0.1% TFA in $H_2O$, B: 0.1% TFA in $CH_3CN$; method: hold 5% B for 4 min, 5-10% B over 6 min, hold 10% B for 3 min, 10-40% B for 8 min, hold 40% for 2 min, 40-95% B over 7 min, hold 95% B for 2 min, 95-20% B over 1 min, hold 20% B for 2 min; UV-Vis: 280, 254 nm, column: RediSep Gold C18Aq 50g HP C18, 100 Å, Teledyne ISCO (part # 69-2203-335); flow rate: 50 mL/min, 100 psi max.

**Method B**: solvents: A: 0.1% TFA in $H_2O$, B: 0.1% TFA in $CH_3CN$; method: hold 10% B for 5.5 min, 10-12.5% B over 2.6 min, hold 12.5% B for 8.9 min, 12.5-20% B over 2 min, 20-30% B for 8 min, 30-95% B over 1 min, hold 95% for 5 min, 95-10% B over 1 min, hold 10% B for 2 min; UV-Vis: 280, 254 nm, column: RediSep Gold C18Aq 50g HP C18, 100 Å, Teledyne ISCO (part # 69-2203-335); flow rate: 40 mL/min.

**Method C:** solvents: A: 0.1% TFA in $H_2O$, B: 0.1% TFA in $CH_3CN$; method: hold 10% B for 1 min, 10-30% B over 9 min, hold 30% B for 11 min, 30-95% B over 2 min, hold 95% B for 5 min, 95-10% B over 1 min, hold 10% B for 2 min; UV-Vis: 284, 254 nm, column: RediSep Gold C18Aq 50g HP C18, 100 Å, Teledyne ISCO (part # 69-2203-335); flow rate: 30 mL/min.

### 2. Small Molecule Synthesis Procedures

#### 6-azidohexyl (2,5-dioxopyrrolidin-1-yl) carbonate (S1):

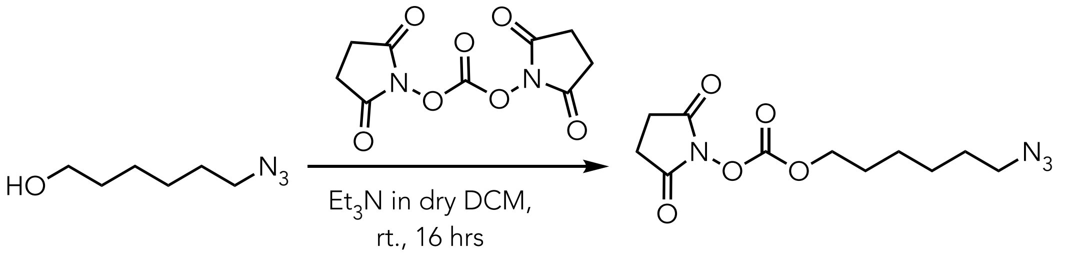


**80%** **S1**

The synthesis of **S1** was achieved by adapting the procedure described in the literature.[5] A round-bottom flask equipped with a stir bar and a side arm inlet adapter was heat dried under vacuum, allowed to cool to room temperature, and refilled with $N_2$. To the flask was added 6-azidohexan-1-ol[6] (2.001 g, 13.97 mmol, 1.0 eq.), and bis(2,5-dioxopyrrolidin-1-yl) carbonate (5.370 g, 20.91 mmol, 1.5 eq.) and the flask was then evacuated and refilled with $N_2$. To the flask 15mL dry $CH_2Cl_2$ and TEA (2.90 mL, 20.9 mmol, 1.5 eq.) was added via syringe and the reaction mixture was stirred for 16 hours at room temperature. The resultant mixture was neutralized with saturated $NH_4Cl$ (aq) and extracted with ether (10 mL x 3). The combined organic phase was washed with brine (10 mL x 1), dried over $Na_2SO_4$, filtered, and concentrated under reduced pressure to give desired product **S1** as light-yellow liquid. (Yield: 3.18 g, 80%, estimated purity by $^1$H-NMR: ≥ 95%).

**$^1$H NMR** (500 MHz, $CDCl_3$) δ: 4.33 (t, *J* = 6.5 Hz, 2H), 3.28(t, *J* = 6.9 Hz, 2H), 2.84 (s, 4H), 1.79-1.74 (m, 2H), 1.64-1.58 (m, 2H), 1.44-1.38. (m, 4H) ppm.

**$^{13}$C NMR** (126 MHz, $CDCl_3$) δ: 168.8, 151.7, 71.4, 51.5, 28.9, 28.48, 26.4, 25.9, 25.2 ppm

**FT-IR:** 2939.6, 2862.4,1810.9, 1787.6, 1736.6, 1711.1, 1463.3, 1430.1, 1362.5, 1256.2, 1198.5, 1090.2, 1046.8, 988.9, 922.7, 812.6, 765.4, 712.1 $cm^{-1}$.

**HR-MS (ESI) m/z:** Found $[M+NH_4]^+$ 302.1451Calc'd for $[C_{11}H_{20}N_5O_5]^+$ 302.1459

#### 2,5-dioxopyrrolidin-1-yl (5-((3a*S*,4*S*,6a*R*)-2-oxohexahydro-1*H*-thieno[3,4-*d*]5midazole-4-yl)pentyl) carbonate (S2):

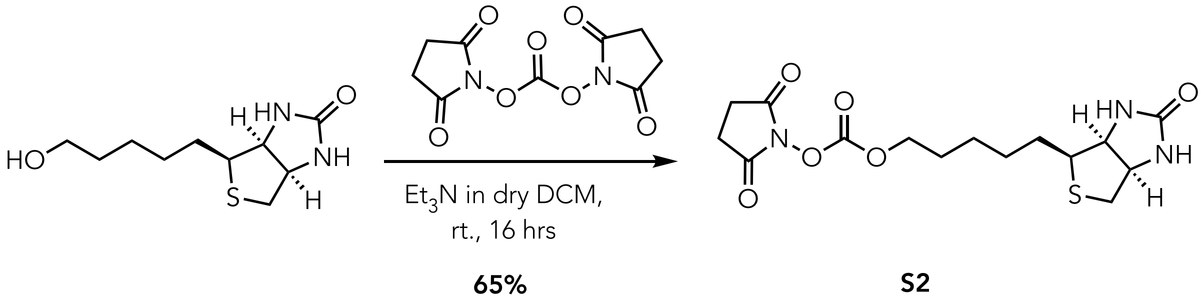


**65%** **S2**

The synthesis of **S2** was achieved by adapting the procedure described in the literature.[5] A round-bottom flask equipped with a stir bar and a side arm inlet adapter was heat dried under vacuum, allowed to cool to room temperature, and refilled with $N_2$. To the flask was added (3a*S*,4*S*,6a*R*)-4-(5-hydroxypentyl)tetrahydro-1*H*-thieno[3,4-*d*]imidazole-2(3*H*)-one[1] (1.100 g, 4.734 mmol, 1.0 eq.) and bis(2,5-dioxopyrrolidin-1-yl) carbonate (1.818 g, 7.100 mmol, 1.5 eq.) and the flask was then evacuated and refilled with $N_2$. To the flask was added 15 mL dry $CH_2Cl_2$ and triethylamine (0.990 mL, 7.10 mmol, 1.5 eq.) sequentially *via* syringe and the reaction mixture was stirred for 16 hours at room temperature. The resultant mixture was neutralized with saturated $NH_4Cl$ (aq.) and filtered to remove the residue. The filtrate was extracted with EtOAc (10 mL x 3). The combined organic phases were washed with brine (10 mL x 1), dried over $Na_2SO_4$, filtered, and concentrated using a rotary evaporator. The residue was washed with ether and dried under high vacuum to give desired product **S2** as white solid. (Yield: 1.14 g, 65%, estimated purity by $^{1}$H-NMR: 90%).

**$^{1}$H NMR** (500 MHz, $CDCl_3$) δ: 5.61 (s, 1H), 4.53-4.51 (m,1H), 4.37-4.31 (m, 3H), 3.18-3.13 (m,1H), 3.10 (m, 2H), 2.94-2.91 (m, 1H), 2.85 (s, 4H), 1.80-1.75 (m, 2H), 1.72-1.65 (m, 2H), 1.48-1.43 (m, 4H) ppm.

**$^{13}$C NMR** (126 MHz, $CDCl_3$) δ: 169.1, 164.5, 151.7, 71.6, 62.3, 60.5, 55.5, 40.6, 28.5, 28.1, 25.6, 25.4 ppm;

**FT-IR:** 3338.5, 2935.8, 1810.5, 1785.9, 1734.9, 1700.2, 1461.2,1428.1, 1257.6, 1201.6, 1088.4, 1047.3, 993.1, 935.6, 813.4, 761.2, 712.9 $cm^{-1}$.

**HR-MS (ESI) m/z:** Found $[M+H]^+$ 372.1214, Calc'd for $[C_{15}H_{21}N_3O_6S]$ 372.1224

**MP:** 88-92 °C

**1-((methoxycarbonyl)(methyl)amino)-4-(6-methoxynapthalen-2-yl)-2,6-dimethylpyridin-1-ium trifluoroacetate (1*):***

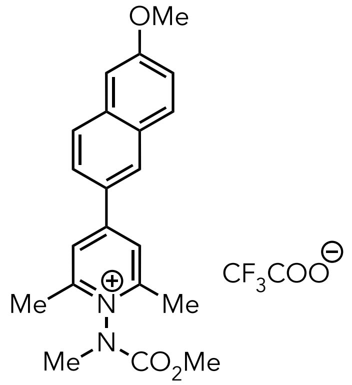


**1**

The synthesis of the compound **1** was achieved by adapting our previously reported procedure,[3] with the added step that an aliquot of the tetrafluoroborate salt of **1** was purified by HPLC Method A to yield trifluoroacetate salt **1** as a yellow powder.

**$^{1}$H NMR** (500 MHz, $CD_3OD$) δ: 8.55 – 8.53 (m, 1H), 8.40 (s, 2H), 8.03 – 7.94 (m, 3H), 7.34 (s, 1H), 7.29 – 7.23 (m, 1H), 3.99 (s, 1.7H), 3.96 (s, 3H), 3.82 (s, 1.3H), 3.61 (s, 1,7H), 3.57 (s, 1.3H), 2.81 (s, 6H) ppm.

**$^{13}$C NMR** (126 MHz, $CD_3OD$) δ: 161.8, 158.9, 155.2, 153.7, 138.6, 132.2, 130.9, 130.1, 129.7, 129.3, 125.5, 121.4, 106.9, 56.1, 55.6, 38.1, 37.1, 19.1 ppm.

**FT-IR:** 1731.8, 1688.9, 1611.2, 1562.4, 1492.6, 1457.2, 1408.2, 1375.9, 1351.2, 1326.6, 1265.9, 1193.2, 1167.6, 1050.6, 1021.5, 938.6, 853.4, 792.3, 759.2, 704.5.

**$^{19}$FNMR** (471 MHz, $CD_3OD$): δ = -77.24 ppm

**MP:** 45-49 °C

**Synthetic Route for 2:**

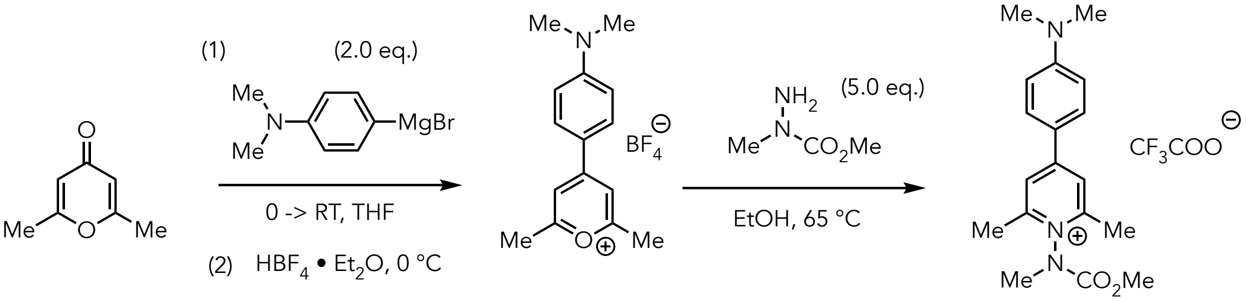


**4-(3,4-dihydronaphthalen-2-yl)-2,6-dimethylpyrylium tetrafluoroborate (S3):**

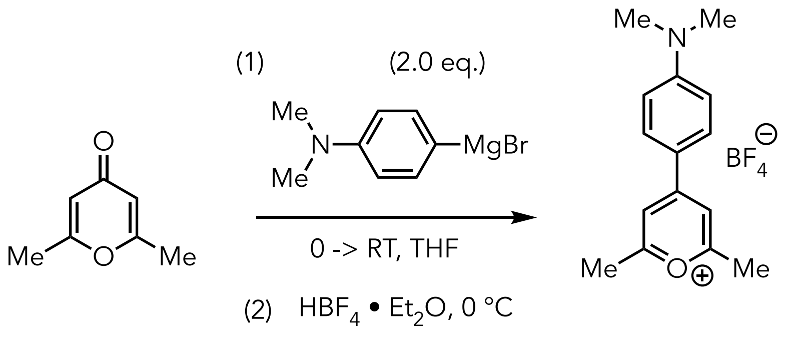


The synthesis of **S3** was achieved by adapting the previously reported procedure as follows.[3] A round-bottom flask equipped with a side arm inlet adapter and stir bar was heat dried under vacuum, allowed to cool to room temperature, and refilled with $N_2$. The flask was charged with the 2,6-dimethyl-gamma-pyrone (0.553 g, 4.45 mmol) and then was evacuated and refilled with $N_2$. To the flask was added THF (1 mL), and the resulting mixture was cooled in an ice bath and stirred. To the stirring, chilled suspension was added (4-(dimethylamino)phenyl)magnesium bromide (17.8 mL of a 0.5 M solution in THF, 8.91 mmol) dropwise over 15 minutes. The resulting solution was stirred at 0°C for 5 minutes and then allowed to warm to room temperature. The solution was stirred vigorously for 12 hours. The mixture was then chilled in an ice bath and quenched carefully by dropwise addition of tetrafluoroboric (2.30 mL of 50-55% w/w in ether, 17.8 mmol). The resulting mixture was allowed to stir for 2 hours while warming to room temperature. To the mixture was added diethyl ether with stirring, and the resulting mixture was filtered. The filter cake was then triturated in ethanol using sonication, filtered, and washed with ethanol. The resulting solid was finally washed with excess diethyl ether and dried to yield a dark maroon solid. (Yield: 0.752 g, 54%)

**$^1$H NMR** (500 MHz, $CD_3CN$) δ: 8.18 (d, *J* = 5.7 Hz, 2H), 8.04 (s, 2H), 7.54 (d, *J* = 9.0 Hz, 2H) 3.30 (s, 6H), 2.83 (s, 6H) ppm.

**$^{13}$C NMR** (125 MHz, $CD_3CN$) δ: 178.3, 177.3, 163.3, 132.4, 131.1, 129.7, 127.7, 121.0, 110.3, 47.6, 44.8, 21.1, 20.6 ppm.

**FT-IR:** 1639.9, 1583.4, 1235.8, 1494.6, 1386.8, 1338.8, 1211.4, 1059.2, 930.4, 896.6, 841.9, 765.3, 737.6 $cm^{-1}$.

**$^{19}$F NMR** (471 MHz, $CD_3CN$) δ: -152.23 - 151.11 (m) ppm.

**HR-MS (ESI) m/z:** Found $[M]^+$ 228.1382, Calcd for $[C_{15}H_{18}NO]^+$ 228.1382

**MP:** 208-210 °C

**4-(4-(dimethylamino)phenyl)-1-((methoxycarbonyl)(methyl)amino)-2,6-dimethylpyridin-1-ium trifluoroacetate (2):**

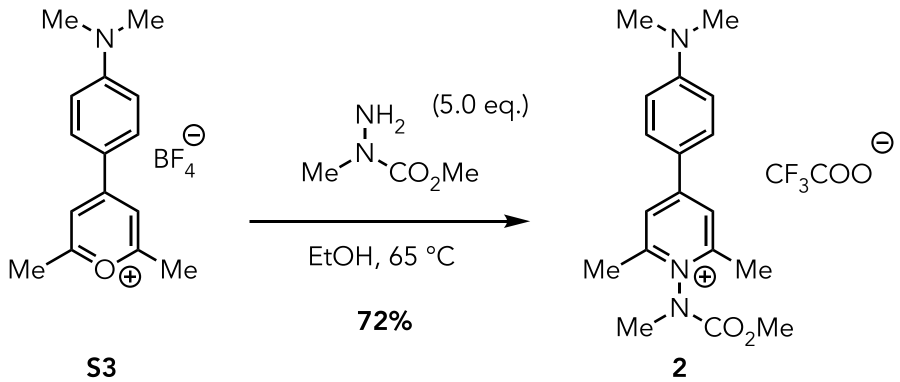


The synthesis of 4-(4-(dimethylamino)phenyl)-1-((methoxycarbonyl)(methyl)amino)-2,6-dimethylpyridin-1-ium **(2)** was achieved by adapting the previously reported procedure as follows.[3] A round-bottom flask equipped with a stir bar and a side arm inlet adapter was heat dried under vacuum, allowed to cool to room temperature, and refilled with $N_2$. The flask was charged with the 4-(3,4-dihydronaphthalen-2-yl)-2,6-dimethylpyrylium salt (0.254 g, 0.806 mmol, 1.0 eq) and was then evacuated and refilled with $N_2$. To the flask was added ethanol (2 mL), and the resulting mixture was stirred vigorously. To the stirring solution was added the 1-methylhydrazine-1-carboxylate[2] (0.252 g, 2.42 mmol, 3.0 eq., 1 M solution in ethanol) dropwise over 20 minutes, and the resulting solution was stirred vigorously at 65 °C for 3 hours. The resultant mixture was then concentrated using a rotary evaporator and purified using Method A in reverse-phase flash chromatography (0.1% TFA in water-acetonitrile) to isolate a yellow residue **(2)** (Yield: 0.248 g, 72%).

**$^1$H NMR** (500 MHz, $CD_3CN$) δ: 7.95 (s, 2H), 7.91 (dd, *J* = 9.2, 3.7 Hz, 2H), 6.90 (d, *J* = 9.2 Hz, 2H) 3.92 (s, 1.6H), 3.79 (s, 1.4H), 3.49 (s, 1.5H), 3.46 (s, 1.5H), 3.13 (s, 6H), 2.62 (s, 6H) ppm.

**$^{13}$C NMR** (125 MHz, $CD_3CN$) δ: 157.0, 157.0, 156.9, 156.7, 155.0, 154.9, 154.8, 153.6, 131.1, 121.6, 113.3, 55.6, 40.3, 38.3, 37.5, 19.1 ppm.

**$^{19}$F NMR** (471 MHz, $CD_3CN$) δ: -76.49 ppm.

**FT-IR:** 1735.6, 1687.5, 1586.5, 1447.1, 1380.4, 1337.4, 1202.6, 1171.8, 1129.9, 822.9, 718.7 $cm^{-1}$.

**HR-MS (ESI) m/z:** Found $[M]^+$ 314.1863, Calc'd for $[C_{18}H_{24}N_3O_2]^+$ 314.1863

**(*E*)-4-(2-(7-Methoxy-2*H*-chromen-3-yl)vinyl)-1-((methoxycarbonyl)(methyl) amino)-2,6-dimethylpyridin-1-ium 2,2,2-trifluoroacetate (3):**

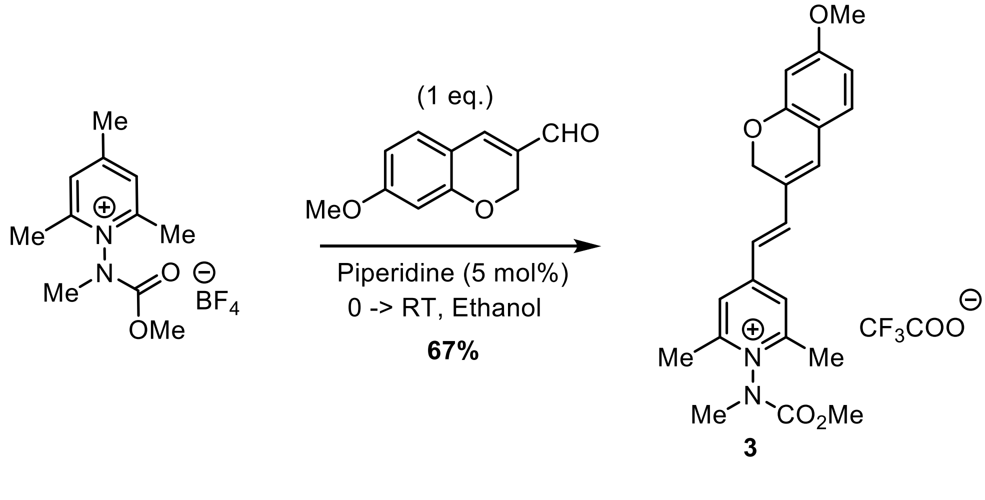

A round-bottom flask equipped with a side arm inlet adapter and stir bar was heat dried under vacuum and refilled with $N_2$. The flask was charged with the 1-((methoxycarbonyl)(methyl)amino)-2,4,6-trimethylpyridin-1-ium tetrafluoroborate[2] (0.220 g, 1.05 mmol, 1.0 eq.). To the flask was added ethanol (2.6 mL) and the mixture was chilled in an ice bath and stirred. To the stirring solution was added piperidine (5.2 µL, 52.58 µmol, 5 mol%), followed by 7-methoxy-2*H*-chromene-3-carbaldehyde (0.200 g, solution of 8 mL in ethanol, 1.05 mmol, 1.0 eq.). The resulting solution was stirred at 0°C for 30 minutes and then allowed to warm to room temperature and stirred for 30 minutes. The resulting solution was concentrated using a rotary evaporator, and the resulting residue was precipitated with dropwise addition of cold diethyl ether. The precipitate was filtered and dissolved in ethanol then again precipitated with cold diethyl ether. The resulting red solid was filtered, dissolved in acetonitrile and water (4:1) and purified using auto flash (Method A) to yield the desired pyridinium salt **3** as a dark orange/brown solid (Yield: 0.266 g, 67%). The compound is a 1.2:1 mixture of rotamers at 300K.

**$^1$H NMR** (500 MHz, $CD_3CN$) δ: 7.78 (d, *J* = 5.0 Hz, 2H), 7.60 - 7.56 (m, 1H), 7.21 - 7.19 (m, 1H), 7.00 (s, 1H), 6.60 - 6.49 (m, 3H), 5.13 (s, 2H), 3.92 (s, 1.6H), 3.83 (s, 3H), 3.75 (s, 1.6H), 3.45 (d, *J* = 15.0 Hz, 3H), 2.63 (s, 6H) ppm.

**$^{13}$C NMR** (126 MHz, $CD_3CN$) δ: 164.0, 158.0, 157.8, 157.4, 156.0, 154.7, 153.3, 141.9, 141.8, 134.1, 134.0, 130.8, 130.8, 128.1, 124.2, 124.1, 121.2, 121.2, 116.3, 109.4, 109.3, 102.3, 65.9, 56.3, 55.7, 55.7, 38.3, 37.4, 19.3, 19.3 ppm.

**$^{19}$F NMR** (471 MHz, $CD_3CN$) δ: -76.10 (s, 3F) ppm.

**FT-IR:** 1681, 1630, 1587, 1505, 1452, 1351, 1275, 1203, 1133, 1031, 838, 800, 721 $cm^{-1}$.

**HR-MS (ESI) m/z:** Found $[M – CF_3COO]^+$ 381.1808, Calc'd for $[C_{22}H_{25}N_2O_4]^+$ 381.1809.

**MP:** 68-70 °C

#### Synthetic Route for 4

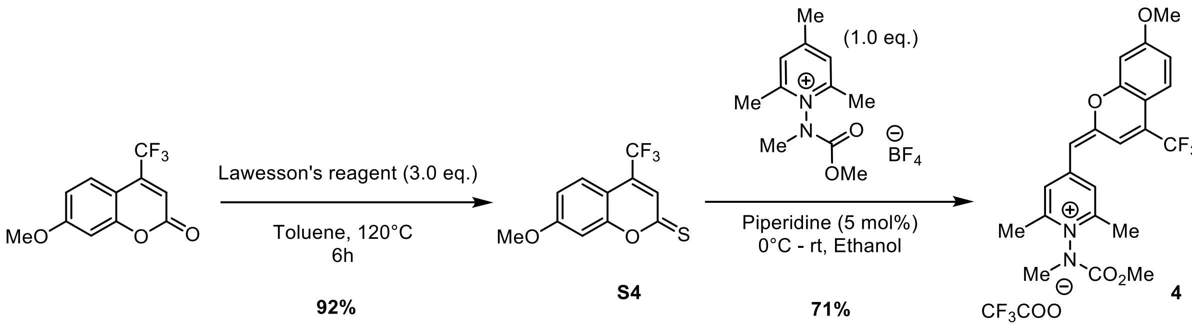


#### 7-Methoxy-4-(trifluoromethyl)-2*H*-chromene-2-thione (S4)

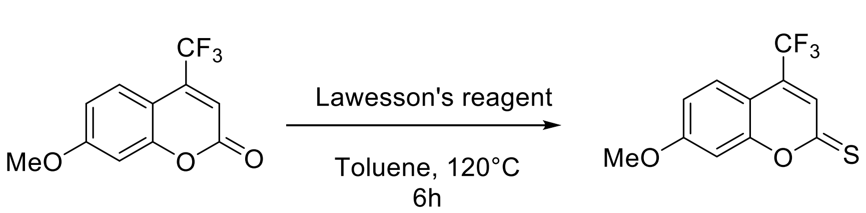


A round-bottom flask equipped with a Liebig condenser and stir bar was charged with 7-methoxy-4-(trifluoromethyl)-2*H*-chromen-2-one[7] (0.200 g, 0.819 mmol 1.0 eq.) and backfilled with $N_2$. To the flask was added sequentially dry toluene and Lawesson's reagent (0.994 g, 2.46 mmol, 3.0 eq.) and the reaction mixture was stirred at reflux until consumption of the starting material was observed by TLC (~6 hrs). After the reaction was complete, the resulting mixture was cooled to room temperature and poured into 20 ml of deionized water. The resulting biphasic mixture was extracted with dichloromethane (20 ml x 3), and the combined organic layers were dried $Na_2SO_4$, filtered, and the solvent was removed using a rotary evaporator. The resulting crude residue was purified using flash chromatography (30% EtOAc:hexane) to obtain the desired compound 7-methoxy-4-(trifluoromethyl)-2*H*-chromene-2-thione **S4** as an orange solid (Yield: 0.196 g, 92%).

**$^1$H NMR** (400 MHz, $CDCl_3$) δ: 7.64 (dd, *J* = 9.0, 1.9 Hz, 1H), 7.33 (d, *J* = 1.0 Hz, 1H), 7.03 – 6.95 (m, 2H), 3.91 (s, 3H) ppm.

**$^{13}$C NMR** (126 MHz, $CDCl_3$) δ: 195.6, 163.8, 159.1, 132.3, 132.0, 126.2, 126.2, 124.7, 124.7, 124.6, 123.4, 121.3, 114.8, 109.0, 101.0, 56.1 ppm.

**FT-IR:** 1629, 1602, 1546, 1350, 1318, 1212, 1182, 1166, 1153, 1137, 885, 853 $cm^{-1}$.

**HR-MS (ESI) (m/z):** Found $[M+H]^+$ 261.0193, calcd for $C_{11}H_8F_3O_2S$ $[M + H]^+$ 261.0197.

**MP:** 104-106 °C

#### (*E*)-4-((7-methoxy-4-(trifluoromethyl)-2*H*-chromen-2-ylidene)methyl)-1-((methoxycarbonyl)(methyl)amino)-2,6-dimethylpyridin-1-ium 2,2,2-trifluoroacetate (4)

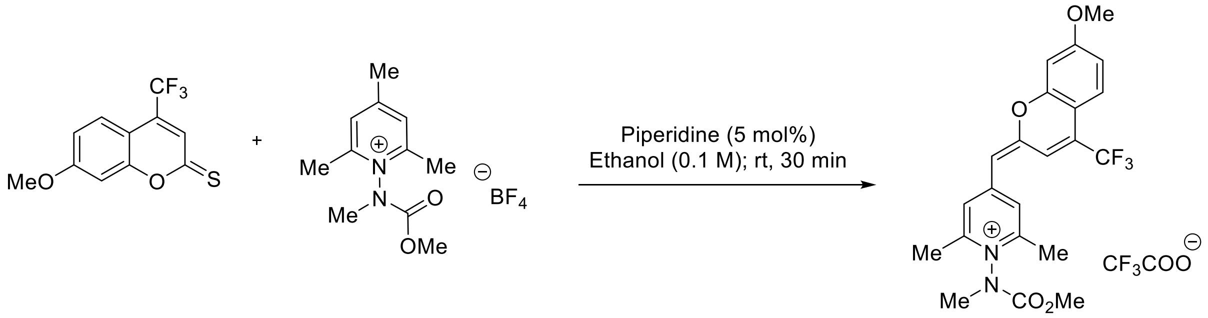


**4**

A round-bottom flask equipped with a side-arm inlet adapter and stir bar was heat-dried under vacuum and refilled with $N_2$. The flask was charged with the 1-((methoxycarbonyl)(methyl)amino)-2,4,6-trimethylpyridin-1-ium tetrafluoroborate[1] (0.113 g, 0.384 mmol, 1.0 equiv.). To the flask was added ethanol (1.3 mL) and the reaction mixture was chilled in an ice bath and stirred. To the stirring solution was added, piperidine (2.0 µL, 19.21 µmol, 5 mol%) followed by addition of 7-methoxy-4-(trifluoromethyl)-2*H*-chromene-2-thione (0.100 g solution in 2.5 mL ethanol, 0.384 mmol, 1.0 eq.). The resulting solution was stirred at 0°C for 30 minutes and then allowed to warm to room temperature and stirred for 30 minutes. The resulting solution was concentrated using rotary evaporator, and the residue precipitated with cold diethyl ether. The precipitate was filtered and dissolved in ethanol, then again precipitated with cold diethyl ether. The resulting red solid was filtered, dissolved in acetonitrile and water (4:1) and purified using auto flash (using Method A) to yield the desired pyridinium salt **4** as orange solid (Yield: 0.119 g, 71%). Compound **4** is a 1.2:1 mixture of rotamers at 300 K.

**$^1$H NMR** (500 MHz, $CD_3CN$) δ: 7.94 (d, *J* = 10.0 Hz, 2H), 7.58 (d, *J* = 10.0 Hz, 1H), 7.21 (s, 1H), 7.07 (s, 1H), 6.98 (d, *J* = 5.0 Hz, 1H), 5.97 (s, 1H), 3.96 (d, *J* = 5.0 Hz, 3H), 3.92 (s, 1.4H), 3.76 (s, 1.5H), 3.49 (s, 1.3H), 3.46 (s, 1.3H), 2.66 (s, 3H), 2.65 (s, 3H) ppm.

**$^{13}$C NMR** (126 MHz, $CD_3CN$) δ: 164.3, 164.3, 159.2, 159.1, 157.2, 157.0, 155.3, 155.3, 154.8, 153.6, 153.0, 153.0, 126.7, 124.8, 124.7, 124.3, 122.5, 122.2, 120.8, 113.4, 113.4, 109.2, 104.2, 104.2, 103.5, 57.1, 57.0, 56.9, 55.7, 55.6, 38.4, 37.5, 19.6, 19.3, 19.2 ppm.

**$^{19}$F NMR** (471 MHz, $CD_3CN$) δ: -65.15 (s, 3F), -76.49 (s, 3F) ppm.

**FT-IR:** 1684, 1628, 1579, 1557, 1514, 1463, 1399, 1356, 1325, 1298, 1278, 1203, 1171, 1129. 801, 722 $cm^{-1}$.

**HR-MS (ESI) m/z:** Found $[M – CF_3COO]^+$ 435.1526, calc'd for $[C_{22}H_{22}F_3N_2O_4]^+$ 435.1526.

**MP:** 96-98 °C

**Synthesis of (*E*)-1-((methoxycarbonyl)(methyl)amino)-2,6-dimethyl-4-styrylpyridin-1-ium trifluoroacetate (5):**

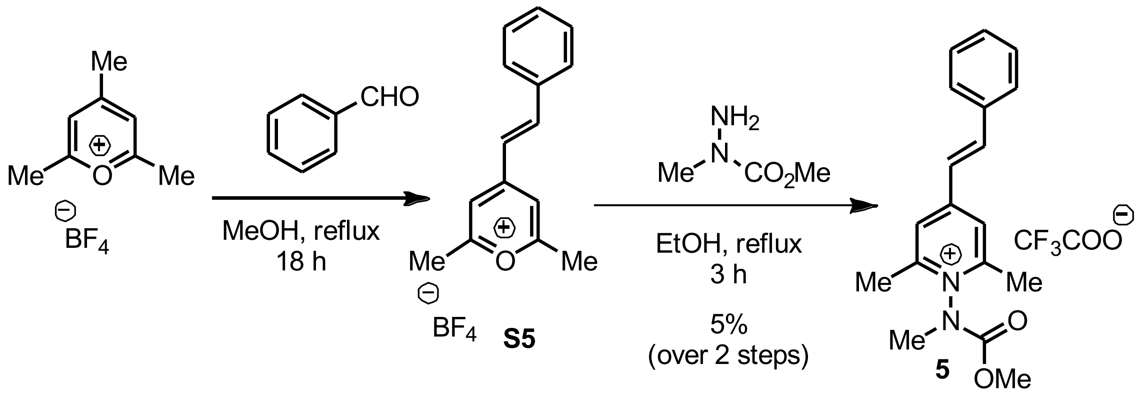


The **S5** was synthesized according to the following procedure adapted from Stark *et al.*[8] A heat-dried round-bottom flask equipped with a stir bar and reflux condenser and a sidearm inlet adapter was charged with 2,4,6-trimethylpyrylium tetrafluoroborate[1] (0.250 g, 1.19 mmol) under $N_2$ atmosphere. To the flask was added sequentially anhydrous MeOH (10 mL) and benzaldehyde (0.132 mL, 1.31 mmol). The reaction mixture was heated to reflux for 18 h with vigorous stirring. The reaction was cooled to room temperature, and the volatiles were removed using a rotary evaporator. The compound **S5** displayed poor stability and was therefore used immediately for the next step without purification. The presence of **S5** was confirmed by LC/MS (method A).
**LR-MS *m/z***: Found $[M]^+$ 211.2, calc'd for $[C_{15}H_{15}O]^+$ 211.1

A round-bottom flask equipped with a stir bar, reflux condenser and a sidearm inlet adapter was charged with the crude solid containing **S5** under $N_2$ atmosphere. To the flask was added EtOH (1 mL) and methyl 1-methylhydrazine-1-carboxylate[2] as a solution in ethanol (0.316 g, 2.97 mmol, 2 mL EtOH) and the resulting mixture was stirred. The mixture was slowly heated to reflux and stirred for another 3 hours. The resulting mixture was cooled to room temperature, and the solvent was removed using a rotary evaporator. The resultant crude material was washed with diethyl ether (3 mL X 2) to remove nonpolar impurities and dried under reduced pressure. The obtained residue was further purified by using reverse phase flash chromatography method B to get the desired product **5** as a pale-yellow residue. (Yield: 0.021 g, 5 %, 90% purity by $^1$H NMR).

**$^1$H NMR** (500 MHz, $CD_3CN$) δ: 7.91 (s, 2H), 7.87 (dd, *J* = 16.3, 5.1 Hz, 1H), 7.76 – 7.71 (m, 2H), 7.53 – 7.46 (m, 3H), 7.30 (d, *J* = 16.3 Hz, 1H), 3.90 (s, 1.6H, rotamer), 3.73 (s, 1.4H, rotamer), 3.47 (3.44) (2 s, 3H, rotamer), 2.65 (d, *J* = 1.7 Hz, 6H) ppm.

**$^{13}$C NMR** (126 MHz, $CD_3CN$) δ: 158.5, 158.3, 156.1, 156.1, 154.5, 153.2, 144.2, 144.1, 135.8, 132.0, 132.0, 129.4, 129.4, 124.9, 124.8, 123.4, 123.3, 55.7, 55.6, 38.2, 37.3, 19.3, 19.3 ppm.

**$^{19}$F NMR** (471 MHz, $CD_3CN$) δ: -75.99 ppm.

**FT-IR:** 725.1, 802.8, 843.4, 1135.4, 1209.3, 1443.2, 1616.3, 1680.9 $cm^{-1}$.

**HR-MS (ESI) *m/z:*** Found $[M]^+$ 297.1594, Calc'd for $[C_{18}H_{21}N_2O_2]^+$ 297.1603

**Synthetic Route for 6**

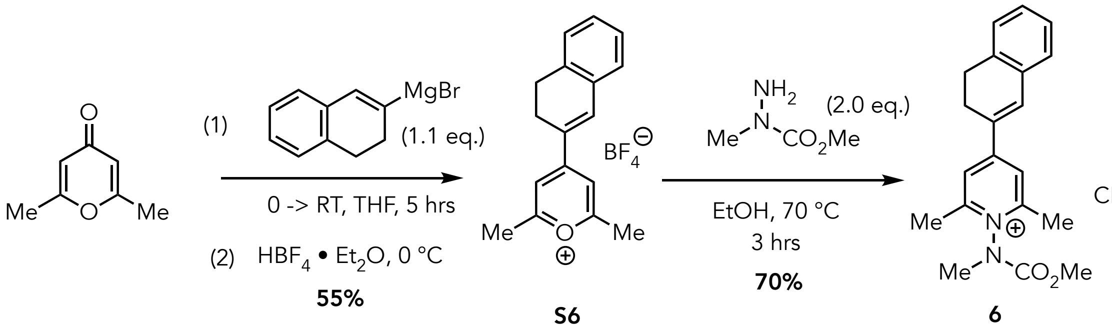

**4-(3,4-dihydronaphthalen-2-yl)-2,6-dimethyl- pyrylium tetrafluoroborate (S6):**

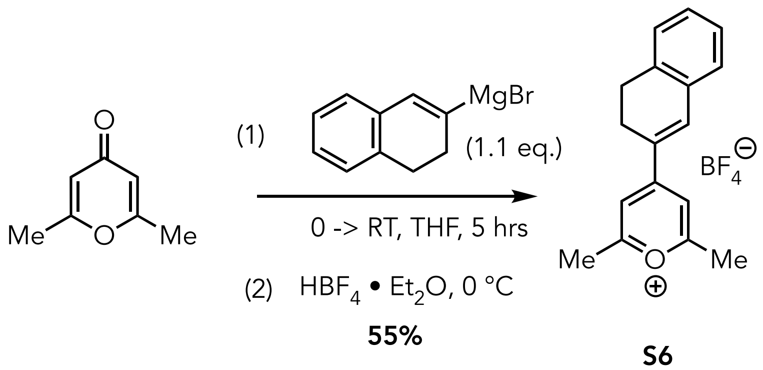

The synthesis of **S6** was achieved by adapting our previously reported procedure.[2] A round bottom flask equipped with a side arm inlet adapter and stir bar was heat dried under vacuum, allowed to cool to room temperature, and refilled with $N_2$. The flask was charged with 2,6-dimethyl-γ-pyrone (0.393 g, 3.17 mmol, 1.0 eq.) and then was evacuated and refilled with $N_2$. To the flask was added tetrahydrofuran *via* syringe (1.0 mL), and the resulting mixture was cooled in an ice batch and stirred. To the stirring, chilled suspension was added freshly prepared (3,4-dihydronaphthalen-2-yl)magnesium bromide[9] (7.0 mL of a 0.5 M solution in THF, 3.5 mmol, 1.1 eq.) dropwise over 15 minutes. The resulting solution was stirred for 5 minutes and then the ice batch was removed and the reaction allowed to warm to room temperature. The solution was stirred vigorously for 5 hours. The mixture was then cooled in an ice bath before being quenched carefully by dropwise addition of tetrafluoroboric acid (1.1 mL of 50-55% w/w in ether, 9.0 mmol). The resulting mixture was allowed to stir for 20 minutes while steadily warming to room temperature. To the mixture was added 10 mL diethyl ether with stirring, and the resulting mixture was filtered. The filter cake was then triturated in ethanol using sonication, filtered, and washed with ethanol. The resulting solid was washed with excess diethyl ether and dried to yield pyrylium salt **S6** as a yellow solid (Yield: 0.565 g, 55%).

**$^1$H NMR** (500 MHz, $CD_3OD$) δ: 8.17 (s, 1H), 8.11 (s, 1H), 7.51-7.49 (m, 1H), 7.45-7.43 (m, ,1H), 7.36-7.33 (m, 1H), 3.09-3.06 (m, 2H), 3.90 (s, 1.6H), 2.88 (s, 6H) ppm.

**$^{13}$C NMR** (126 MHz, $CD_3OD$) δ: 176.5, 166.1, 142.9, 138.1, 132.1, 130.4, 127.6, 127.1, 115.5, 65.5, 26.7, 23.6, 19.8 ppm.

**$^{19}$FNMR** (471 MHz, $CD_3OD$): δ = -154.54 ppm.

**FT-IR:** 1640.4, 1598.0, 1552.1, 1487.7,1337.8, 1183.2, 1036.0, 975.5, 911.4, 874.2 $cm^{-1}$.

**HR-MS (ESI) m/z:** Found $[M]^+$ 237.1274, Calc'd for $[C_{17}H_{17}O]^+$ 237.1274

**MP:** 170-175 °C

**4-(3,4-dihydronaphthalen-2-yl)-1-((methoxycarbonyl)(methyl)amino)-2,6-dimethylpyridin-1-ium trifluoroacetate (6):**

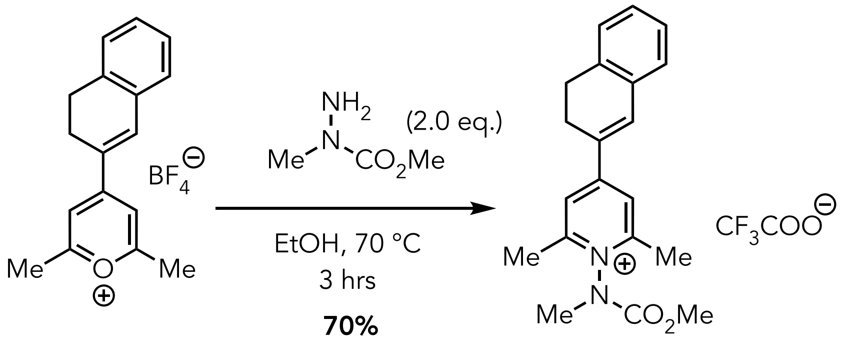


**S6** **6**

Compound **6** was synthesized by adapting the previously reported procedure as follows.[3] To a heat dried 20 mL microwave vial equipped with a stir bar, was added sequentially **S6** (0.300 g, 0.925 mmol, 1.0 eq.), methyl 1-methylhydrazine-1-carboxylate[2] (0.193 g in a solution in 1.0 mL EtOH, 1.85 mmol, 2.0 eq.) and ethanol (4 mL). The reaction was allowed to stir vigorously for 3 hours at 70 °C. The reaction mixture was cooled to room temperature and cold ether was added. The resulting precipitant was allowed to settle, and the supernatant was carefully decanted, and resulting precipitate was purified by HPLC Method A to get desired product as yellow powder (Yield: 0.282 g, 70%).

**$^1$H NMR** (500 MHz, $CD_3OD$) δ: 8.13 (s, 2H), 7.74 (s, 1H), 7.38-7.28 (m,2H), 3.97 (s, 1.6H), 3.80 (s, 1.2H), 3.59 (s, 1.6H), 3.55 (s, 1.2H), 3.04 (t, *J*= 8.2 Hz, 2H), 2.83 (d, *J*= 7.6 Hz, 2H), 2.74 (s, 6H) ppm.

**$^{13}$C NMR** (126 MHz, $CD_3OD$) δ: 160.4, 157.2, 157.0, 153.8, 136.8, 132.6, 130.3, 129.0, 127.3, 126.8, 122.2, 54.3, 36.6, 35.7, 27.0, 24.2, 17.5 ppm.

**$^{19}$FNMR** (471 MHz, $CD_3CN$) δ: -77.25 ppm.

**FT-IR:** 1735.4, 1687.5, 1631.4, 1603.9, 1562.5, 1454.6, 1338.1, 1272.4, 1181.6, 1135.8, 1030.4, 987.2, 923.2, 864.3, 794.4, 762.4, 706.5 $cm^{-1}$.

**HR-MS (ESI) m/z:** Found $[M]^+$ 323.1753, Calc'd $[C_{20}H_{23}N_2O_2]^+$ 323.1754

**MP:** 45-49 °C

### Synthetic Route for 7

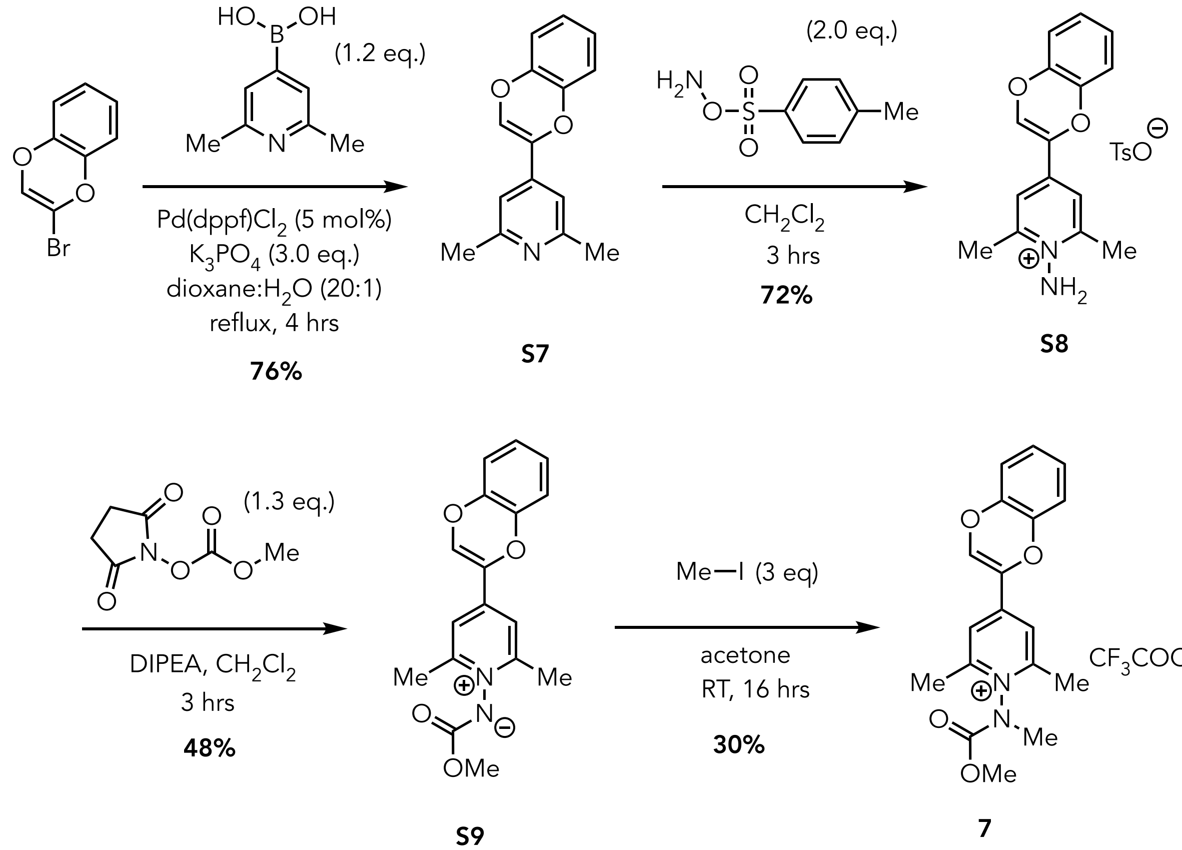


**4-(benzo[*b*][1,4]dioxin-2-yl)-2,6-dimethylpyridine (S7)**

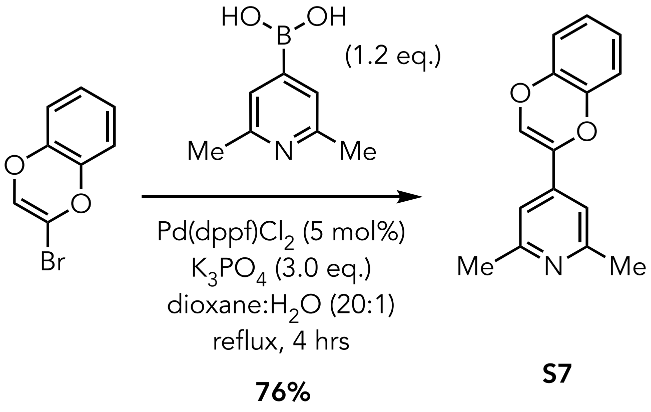


A heat dried round bottom flask equipped with a stir bar, reflux condenser, and side arm inlet adapter was charged with 2-bromobenzo[*b*][1,4]dioxine[10] (1.043 g, 4.896 mmol, 1.0 eq.), 6-dimethylpyridin-4-yl)boronic acid (0.887 g, 5.87 mmol, 1.2 eq.), Pd(dppf)Cl$_2$ (0.179 g, 0.245 mmol, 5.0 mol%), and K$_3$PO$_4$ (3.117 g, 14.69 mmol, 3.0 eq.) and the flask was evacuated and refilled with N$_2$. To flask was added 12 mL of 1,4-dioxane: H$_2$O (20:1, v/v) and resulting mixture was heated to reflux for 18 hours under N$_2$ with continuous stirring. The reaction mixture was cooled down to room temperature, and the solid was removed by filtration through Celite. The filter cake was washed with EtOAc and the filtrate was then washed with H$_2$O and brine. The organic layer was dried over Na$_2$SO$_4$, filtered, and the solvent removed using a rotary evaporator. The crude residue obtained was purified by column chromatography using 40% EtOAC:Hexane to yield the desired product **S7** as a white solid. (0.891 g, 76%)

**$^1$H NMR** (500 MHz, $CD_3OD$) δ: 7.17 (s, 2H), 7.02 (s, 1H), 6.94-6.85 (m, 3H), 6.75-6.73 (m, 1H), 2.48 (s, 6H) ppm.

**$^{13}$C NMR** (126 MHz, $CD_3OD$) δ: 157.5, 142.2, 141.5, 140.47, 134.2, 127.0, 124.5, 124.3, 116.0, 115.8, 113.5, 22.6.ppm.

**FT-IR:** 1676.7, 1596.6, 1556.4, 1420.0, 1383.9, 1347.8, 1329.8, 1280.4, 1254.9, 1208.1, 1117.8, 1078.1, 1026.1, 996.4, 931.2, 867.9, 857.3, 805.7, 792.8, 769.0, 744.8, 723.2 $cm^{-1}$.

**HR-MS (ESI) m/z:** Found $[M + H]^+$ 240.1012, Calc'd for $[C_{15}H_{14}NO_2]^+$ 240.1019

**MP:** 82-85 °C

**1-amino-4-(benzo[*b*][1,4]dioxin-2-yl)-2,6-dimethylpyridinium tosylate (S8):**

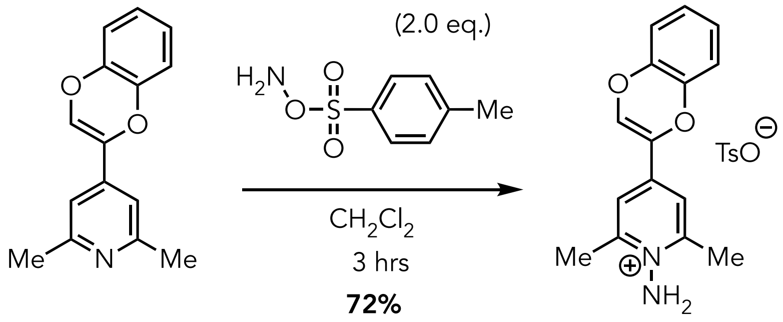


**S7** **S8**

A vacuum dried round bottomed flask equipped with a stir bar and sidearm inlet adapter was charged with 4-(benzo[*b*][1,4]dioxin-2-yl)-2,6-dimethylpyridine **S7** (0.849 g, 3.55 mmol, 1.0 eq.), and evacuated and refilled with $N_2$. To the flask was added freshly prepared O-tosylhydroxylamine[11] (10 mL, 0.71 M in anhydrous $CH_2Cl_2$, 7.1 mmol, 2.0 eq., in), and the resulting mixture was stirred for 3 hours at rt. The reaction mixture was concentrated using a rotary evaporator and the resulting residue was washed with a 1:1 mixture of EtOAc:hexane (25 mL x 3). The resulting residue was then sonicated with 5% MeOH in 4:1 mixture of $CH_2Cl_2$:Ether. The supernatant was decanted, and the sonication process was repeated twice. Subsequently, the residue was dried under vacuum to yield 1-amino-4-(6-methoxy-2*H*-chromen-3-yl)-2,6-dimethylpyridin-1-ium tosylate **S8** as a pale-yellow solid (Yield: 1.088 g, 72%).

**$^1$H NMR** (500 MHz, $CD_3OD$) δ: 7.77 (s, 2H), 7.65 (d, *J* = 8.5 Hz, 2H), 7.43 (s, 1H), 7.19 (d, *J* = 8.3 Hz, 2H), 7.00-6.92 (m, 3H), 6.82 (d, *J* = 5.9 Hz, 1H), 2.77 (s, 6H), 2.34 (s, 3H) ppm.

**$^{13}$C NMR** (126 MHz, $CD_3OD$) δ: 155.9, 144.9, 143.4, 143.0, 141.7, 133.9, 129.8, 126.9, 126.2, 119.7, 117.5, 21.3, 20.2.ppm.

**FT-IR:** 1630.8, 1595.0, 1494.6, 1360.9, 1255.5, 1210.5, 1186.2, 1167.7, 1122.4, 1103.1, 1083.0, 1033.5, 1010.4, 942.7, 817.5, 763.0, 746.6, 683.5 $cm^{-1}$.

**HR-MS (ESI) m/z:** Found $[M+H]^+$ 255.1120, Calc'd for $[C_{15}H_{15}N_2O_2]^+$ 255.1128

**MP:** 145-148 °C

**(4-(benzo[*b*][1,4]dioxin-2-yl)-2,6-dimethylpyridin-1-ium-1-yl)(methoxycarbonyl)amide (S9):**

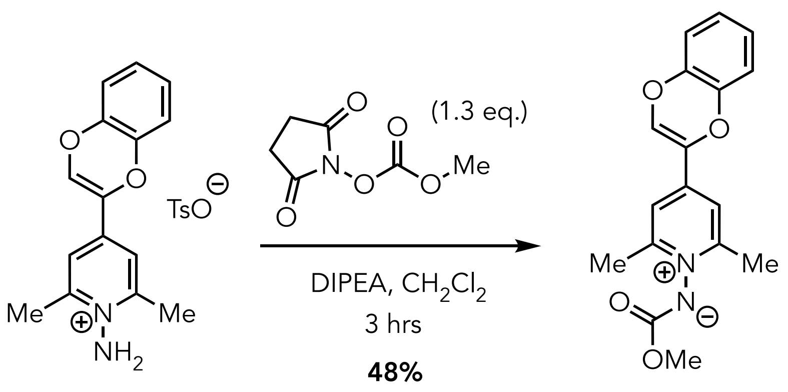


**S8** **S9**

An oven-dried round bottomed flask equipped with a stir bar and sidearm inlet adapter was charged with 1-amino-4-(benzo[b][1,4]dioxin-2-yl)-2,6-dimethylpyridinium tosylate **S8** (0.350 g, 0.820 mmol, 1.0 eq) and 2,5-dioxopyrrolidin-1-yl methyl carbonate[4] (0.185 g, 1.07 mmol, 1.3 eq.) and the flask was evacuated and refilled with $N_2$. To the flask was added dry $CH_2Cl_2$ (4.0 mL) via syringe, and the resulting mixture was stirred. To the stirring solution was added *N*,*N*-diisopropylethylamine (0.430 mL, 2.46 mmol, 3.0 eq.) dropwise using a syringe, and the resulting solution was stirred vigorously at room temperature for 3 hours. The resultant mixture was then concentrated to dryness using a rotary evaporator. The resulting crude residue was purified by normal phase column chromatography using 5% MeOH:$CH_2Cl_2$ as an eluent to yield the desired ylide **S9** as yellow solid. (Yield: 0.123 g, 48%).

**$^1$H NMR** (500 MHz, $CD_3OD$) δ: 7.98 (s, 2H), 7.65 (s, 1H), 7.06-6.96 (m, 3H), 6.89-6.87 (m, 1H), 3.92 (s, 3H), 2.71 (s, 3H) ppm.

**$^{13}$C NMR** (126 MHz, $CD_3OD$) δ: 157.8, 148.0, 141.5, 140.3, 135.4, 132.3, 125.8, 125.0, 118.3, 116.3, 53.5, 18.1.ppm.

**FT-IR:** 1685.0, 1629.3, 1594.9, 1494.2, 1413.1, 1348.5, 1255.9, 1199.5, 1131.5, 1103.3, 1084.1, 798.7, 756.4, 718.6 $cm^{-1}$.

**HR-MS (ESI) m/z:** Found $[M+H]^+$ 313.1171, Calc'd for $[C_{17}H_{17}N_2O_4]^+$ 313.1183

**MP:** 65-70 °C

**4-(benzo[b][1,4]dioxin-2-yl)-1-((methoxycarbonyl)(methyl)amino)-2,6-dimethylpyridin-1-ium trifluoroacetate (7):**

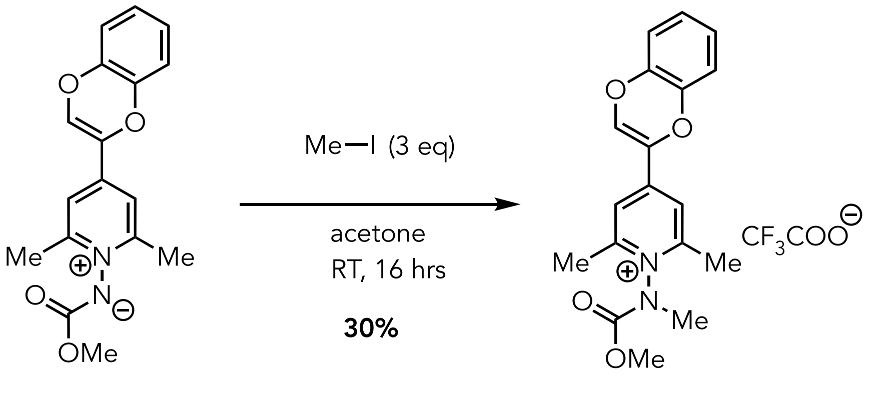


**S9** **7**

A scintillation vial (10 mL) equipped with a stir bar was charged with ylide **S9** (0.120 g, 0.384 mmol,1.0 eq) and the vial was evacuated and filled with $N_2$. To the vial was added acetone (1 mL) and the resulting mixture stirred under an inert atmosphere. To the stirring solution was added methyl iodide (71.0 µL, 1.15 mmol, 3.0 eq.) the resulting mixture was allowed to stir for 16 hours at room temperature. The resulting mixture was then concentrated using a rotary evaporator and the residue was washed with cold diethyl ether. The crude product was purified by HPLC Method A to yield product **6** as light-yellow solid. (Yield : 0.051 g ,30%).

**$^1$H NMR** (500 MHz, $CD_3OD$) δ: 8.00 (s, 2H), 7.68 (s, 1H), 7.04-6.96 (m, 3H), 6.90-6.88 (m, 1H), 3.95 (s, 1.7H), 3.79 (s, 1.3H), 3.56 (s, 1.7H), 3.52 (s, 1.3H), 2.71 (s, 3H) ppm.

**$^{13}$C NMR** (126 MHz, $CD_3OD$) δ: 158.3, 154.3, 145.9, 130.2, 128.8, 126.5, 124.6, 121.4, 120.8, 55.0, 37.2, 36.3, 18.3 ppm.

ppm.

**$^{19}$FNMR** (471 MHz, $CD_3CN$) δ: -76.85 ppm.

**FT-IR:** 1685.0, 1629.3, 1594.9, 1494.2, 1413.1, 1348.5, 1255.9, 1199.5, 1131.5, 1103.3, 1084.1, 798.7, 756.4, 718.6 cm$^{-1}$.

**HR-MS (ESI) m/z:** [Found [M]$^+$ 327.1335, Calc'd for $[C_{18}H_{19}N_2O_4]^+$ 327.1334

**MP:** 48-51 °C

**Synthetic Route for 8**

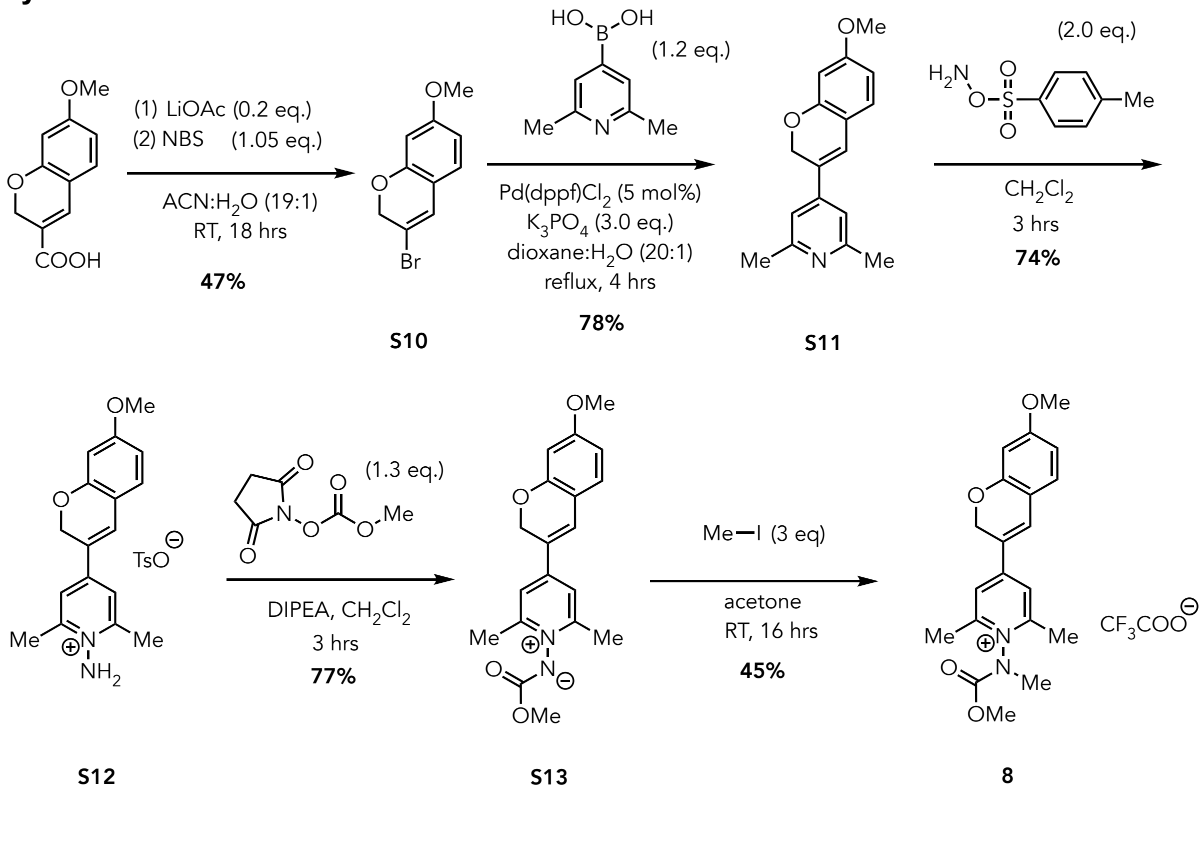

**4-(7-methoxy-2*H*-chromen-3-yl)-2,6-dimethylpyridine (S11):**

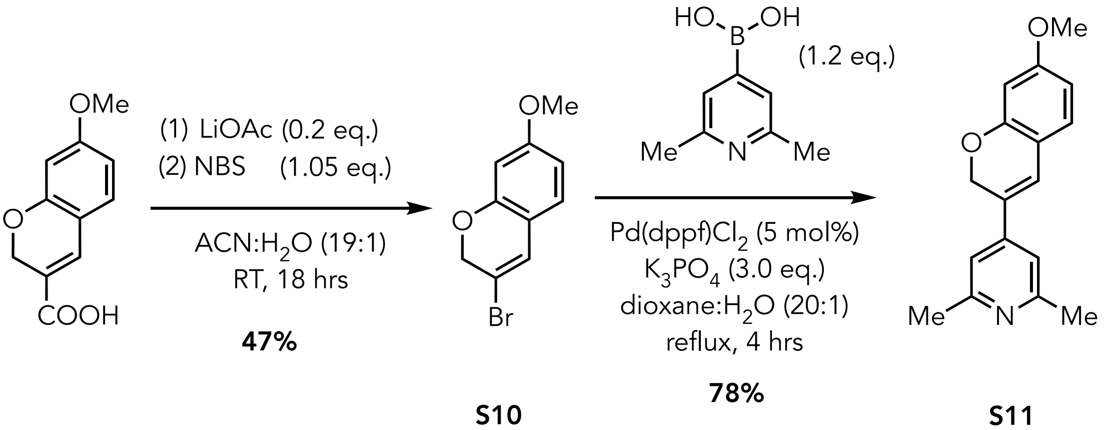


**S10** **S11**

A 50 mL, round bottom flask equipped with a stir bar and side arm inlet adapter was charged with lithium acetate (0.135 g, 2.04 mmol, 0.2 eq.). To the flask was added 0.750 mL of distilled water and the flask was evacuated and refilled with $N_2$ and the resulting solution was stirred. To the stirring solution was added a solution of 7-methoxy-2*H*-chromene-3-carboxylic acid[12] in $CH_3CN$ (2.112 g, 10.24 mmol, 1.0 eq., 10 mL $CH_3CN$) and the resulting reaction mixture stirred vigorously. To the stirring mixture was added dropwise a solution of *N*-bromosuccinimide in $CH_3CN$ (1.913 g, 10.75 mmol, 1.05 eq., 4.5 mL $CH_3CN$) and the resulting mixture was stirred for 18 hours and finally quenched by adding 5% (w/v) $Na_2S_2O_3$ (aq.). The resulting mixture was extracted with EtOAc (3 x 15mL) and the combined organic layers were washed with brine, dried over anhydrous $Na_2SO_4$, filtered, and concentrated using rotary evaporator. The oily crude residue obtained was purified by column chromatography using 5% EtOAc:hexane to yield the desired product as colorless oil. (yield: 1.171g, 47%).

**$^1$H NMR** (500 MHz, $CDCl_3$) δ: 6.84-6.83 (m, 1H), 6.69 (s,1H), 6.45-6.43 (m, 1H), 6.38 (s, 1H), 4.84 (s, 2H), and 3.77 (s, 3H) ppm.

**$^{13}$C NMR** (126 MHz, $CDCl_3$) δ: 160.9, 153.4, 126.9, 125.7, 115.6, 111.5, 107.6, 102.0, 70.4, 55.6.ppm.

**LR-MS (ESI) m/z:** Found $[M+H]^+$ 239.1, Calc'd for $[C_{10}H_9BrO_2]^+$ 239.9

An oven-dried round bottom flask equipped with a stir bar, reflux condenser and side arm inlet adapter was charged with freshly prepared 3-bromo-7-methoxy-2*H*-chromene (1.120 g, 4.645 mmol, 1.0 eq.), 2,6-dimethylpyridin-4-yl)boronic acid (0.842 g, 5.57 mmol, 1.2 eq.), Pd(dppf)$Cl_2$ (0.170 g, 0.465 mmol, 5.0 mol%), and $K_3PO_4$ (2.958 g , 13.93 mmol, 3.0 eq.) and the flask was evacuated and refilled with $N_2$. To flask was added 15 mL of 1,4-dioxane:$H_2O$ (20:1, v/v) and the reaction mixture was refluxed for 4 hours under $N_2$ with continuous stirring. The reaction mixture was cooled down to room temperature, and the solid was removed by filtration through Celite. The filter cake was washed with EtOAc and the filtrate was then washed with $H_2O$ and brine. The organic layer was dried over $Na_2SO_4$, filtered, and the solvent removed using a rotary evaporator. The resultant residue was purified by column chromatography using 10-50% EtOAc:Hexane to yield **S11** as yellow solid. (Yield: 0.969 g, 78%)

**$^1$H NMR** (500 MHz, $CD_3OD$) δ: 7.14 (s, 2H), 7.13 (s, 1H), 7.08 (d, *J* = 8.4 Hz, 1H), 6.51 (dd, *J* = 8.4, 2.5 Hz, 1H), 6.41(s, 1H), 5.08 (s, 3H), 3.77 (s, 3H), 2.49 (s, 6H) ppm.

**$^{13}$C NMR** (126 MHz, $CD_3OD$) δ: 161.5, 158.3, 155.3, 144.7, 128.5, 126.1, 123.0, 115.5, 107.9, 101.6, 66.7, 55.6, 25.0 ppm.

**FT-IR:** 1596.1, 1570.0, 1552.9, 1503.2, 1442.5, 1380.6, 1303.2, 1274.4, 1193.9, 1157.3, 1131.4, 1115.6, 1030.2, 844.6 $cm^{-1}$.

**HR-MS (ESI) m/z:** Found $[M+H]^+$ 268.1333, Calc'd for $[C_{17}H_{18}NO_2]^+$ 268.1332

**MP:** 72-75 °C

**1-amino-4-(7-methoxy-2*H*-chromen-3-yl)-2,6-dimethylpyridinium tosylate (S12):**

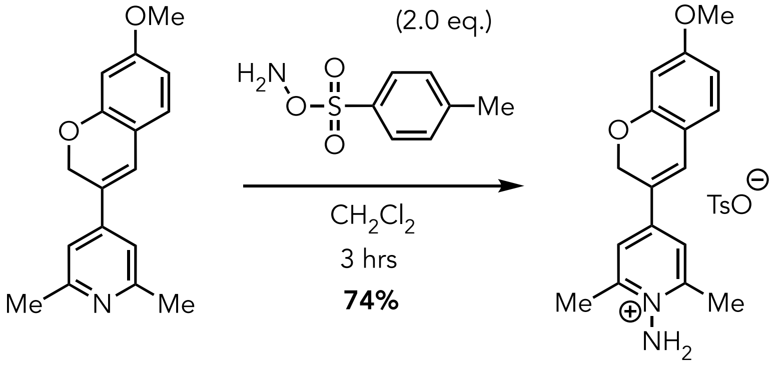


**S11** **S12**

A vacuum dried round bottomed flask equipped with a sidearm inlet adapter was charged with 4-(7-methoxy-2*H*-chromen-3-yl)-2,6-dimethylpyridine **S11** (0.950 g, 3.55 mmol, 1.0 eq.), and evacuated and refilled with $N_2$. To the flask was added freshly prepared O-tosylhydroxylamine[11] (10 mL, 0.71 M in anhydrous $CH_2Cl_2$, 7.1 mmol, 2.0 eq.), and the resulting mixture was stirred for 3 hours at rt. The reaction mixture was washed with 1:1 mixture of EtOAc in hexane (25 mL x 3). The resulting residue was then sonicated with 5% MeOH in 4:1 mixture of $CH_2Cl_2$:Ether. The supernatant was decanted, and the sonication process was repeated twice. Subsequently, the residue was dried under vacuum to yield 1-amino-4-(6-methoxy-2H-chromen-3-yl)-2,6-dimethylpyridin-1-ium tosylate **S12** (1.195g, 2.629 mmol) as a bright yellow solid in 74% yield.

**$^1$H NMR** (500 MHz, $CD_3OD$) δ: 7.78 (s, 2H), 7.67 (d, *J* = 8.3 Hz, 2H), 7.57 (s, 1H), 7.21 (d, *J* = 8.4 Hz, 3H), 6.58 (dd, *J* = 8.4, 2.4 Hz, 1H), 6.47 (s, 1H), 5.15 (s, 2H), 3.81 (s, 3H), 2.79 (s, 6H), 2.35 (s, 3H) ppm.

**$^{13}$C NMR** (126 MHz, $CD_3OD$) δ: 163.4, 156.3, 154.3, 148.6, 142.1, 140.3, 129.9, 128.4, 125.5, 122.1, 120.4, 114.9, 108.2, 101.0, 65.1, 54.7, 19.9, 18.8 ppm;

**FT-IR:** 2925.6, 1733.9, 1600.3, 1561.1, 1443.7, 1373.6, 1243.4, 1219.1, 1197.8, 1166.1, 1120,3, 1031.11, 1009.9, 945.7, 856.1, 810.7, 710.2, 682.3 $cm^{-1}$.

**HR-MS (ESI) m/z:** Found $[M]^+$ 283.1441, Calc'd for $[C_{17}H_{19}N_2O_2]^+$ 283.1441

**MP:** 180-185 °C

**(4-(7-methoxy-2*H*-chromen-3-yl)-2,6-dimethylpyridin-1-ium-1-yl)(methoxycarbonyl)-amide (S13):**

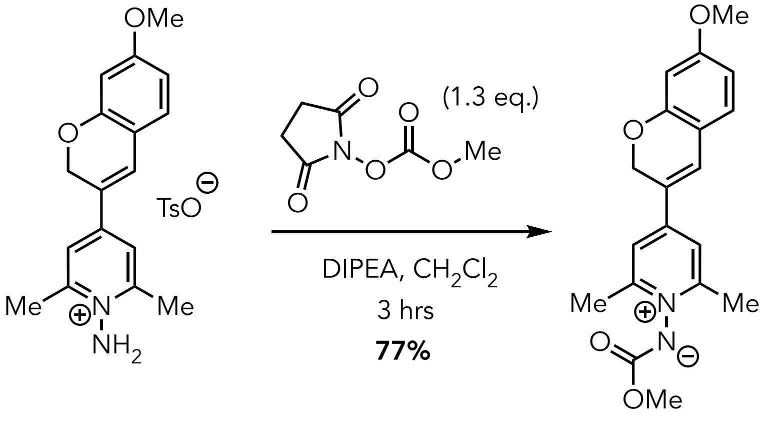


**S12** **S13**

An oven-dried round bottomed flask equipped with a stir bar and sidearm inlet adapter was charged with 1-amino-4-(7-methoxy-2*H*-chromen-3-yl)-2,6-dimethylpyridinium tosylate **S12** (0.327g, 0.697mmol, 1.0 eq.) and 2,5-dioxopyrrolidin-1-yl methyl carbonate[4] (0.157 g, 0.906 mmol,1.3 eq.) and the flask was evacuated and refilled with $N_2$. To the flask was added dry $CH_2Cl_2$ (3.5 mL) *via* syringe, and the resulting mixture was stirred. To the stirring solution was added dropwise *N*,*N*-diisopropylethylamine (365.0 μL, 2.091 mmol, 3.0 eq.) and the resulting solution was stirred vigorously at room temperature for 3 hours. The resultant mixture was then concentrated to dryness using a rotary evaporator. The residue was purified by flash chromatography using 5% MeOH:$CH_2Cl_2$ as an eluent to yield the desired ylide product **S13** as greenish-yellow residue. (Yield: 0.183 g, 77%).

**$^1$H NMR** (500 MHz, $CD_3CN$) δ: 7.60 (s, 2H), 7.38 (s, 1H), 7.19 (d, *J* = 8.4 Hz, 2H), 6.61-6.58 (m, 1H), 6.50 (s, 1H), 5.16 (s, 2H), 3.83 (s, 3H), 3.64 (s, 3H), 2.55 (s, 3H) ppm.

**$^{13}$C NMR** (126 MHz, $CD_3CN$) δ: 163.7, 162.3, 156.7, 155.8, 146.1, 130.6, 128.3, 124.3, 121.1, 116.1, 109.2, 102.3, 66.6, 56.3, 52.6, 19.9 ppm.

**FT-IR:** 1605.8, 1561.9, 1442.5, 1504.9, 1279.4, 1197.3, 1161.3, 1133.6, 1117.9, 1080.9, 1030.9 $cm^{-1}$.

**HR-MS (ESI) m/z:** Found $[M+H]^+$ 341.1488, Calc'd for $[C_{19}H_{21}N_2O_4]^+$ 341.1496

**4-(7-methoxy-2*H*-chromen-3-yl)-1-((methoxycarbonyl)(methyl)amino)-2,6-dimethylpyridinium trifluoroacetate (8):**

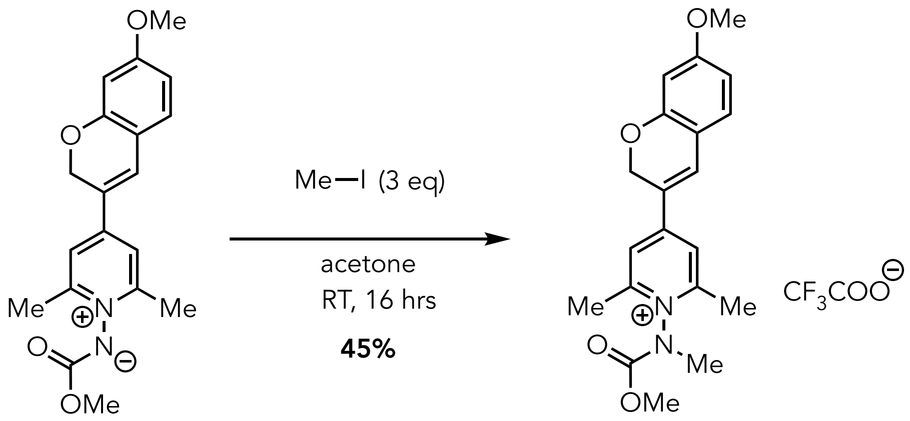


**S13** **8**

A 10 mL scintillation vial, equipped with a stir bar, was charged with ylide **S13** (0.175 g, 0.514 mmol, 1.0 eq.), and the vial was evacuated and filled with $N_2$. To the vial was added acetone (1.5 mL) and the resulting mixture stirred under an inert atmosphere. To the stirring solution was added methyl iodide (96.0 μL, 1.54 mmol, 3 eq.) and the resulting mixture was allowed to stir for 16 hours at room temperature. The resulting mixture was then concentrated using a rotary evaporator and the residue was washed with cold diethyl ether. The crude product was purified by HPLC Method A to yield product **8** as a pumpkin-orange solid. (Yield: 0.108 g, 45%).

**$^1$H NMR** (500 MHz, $CD_3CN$) δ: 7.78 (s, 2H), 7.72 (d, *J* = 7.8, 1H), 7.26 (dd, *J* = 8.4, 4.3 Hz, 1H), 6.63-6.61 (m, 1H), 6.51 (s, 1H), 5.18 (s, 2H), 3.90 (s, 1.7H), 3.83 (s, 3H), 3.73 (s, 1.4H), 3.47 (s, 1.7H), 3.44 (s, 1.5H), 2.63 (s, 6H) ppm.

**$^{13}$C NMR** (126 MHz, $CD_3CN$) δ: 164.8, 159.4, 157.7, 157.5, 154.3, 153.5, 152.9, 134.2, 131.6, 122.2, 115.4, 109.6, 101.8, 65.8, 56.2, 55.3, 37.9, 37.0, 18.9 ppm

**$^{19}$FNMR** (471 MHz, $CD_3CN$) δ: -75.33 ppm.

**FT-IR:** 1729.3, 1685.8, 1635.2, 1594.4, 1550.8, 1504.5, 1458.1, 1340.0, 1274.7, 1196.4, 1161.1, 1112.5, 1019.0, 922.7, 818.9, 798.2, 761.2, 716.0, 677.9 $cm^{-1}$.

**HR-MS (ESI) m/z:** Found $[M]^+$ 355.1653, Calc'd for $[C_{20}H_{23}N_2O_4]^+$ 355.1652

**MP:** 40-44 °C

**4-(7-methoxy-2*H*-chromen-3-yl)-2,6-dimethyl-1-(methyl(((5-((3a*R*,4*S*,6a*S*)-2-oxohexahydro-1*H*-thieno[3,4-*d*]imidazol-4-yl)pentyl)oxy)-carbonyl)amino)pyridin-1-ium trifluoroacetate (8a):**

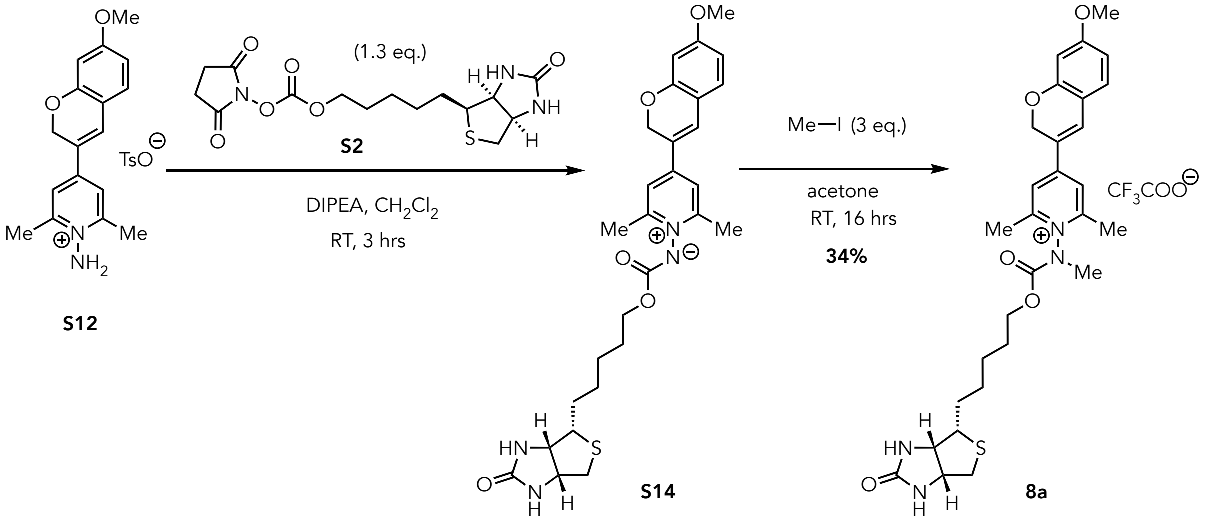


An oven-dried round bottomed flask equipped with a stir bar and sidearm inlet adapter was charged with 1-amino-4-(7-methoxy-2*H*-chromen-3-yl)-2,6-dimethylpyridinium tosylate **S12** (0.220 g, 0.330 mmol, 1.0 eq.) and **S2** (0.159 g, 0.429 mmol,1.3 eq.) and then the flask was evacuated and refilled with $N_2$. To the flask was added anhydrous $CH_2Cl_2$ (1.5 mL) *via* syringe, and the resulting mixture was stirred. To the stirring solution was added dropwise *N*,*N*-diisopropylethylamine (0.170 mL, 0.990 mmol, 3.0 eq.) and the resulting solution was stirred vigorously at room temperature for 3 hours. The resulting mixture was concentrated using a rotary evaporator, and the resulting residue was washed with ether to yield a dark brown crude residue containing **S14** that used for next step without further purification. The presence of **S14** was verified by LRMS.

**LR-MS (ESI) m/z:** Found $[M+H]^+$ 539.4, Calc'd for $[C_{28}H_{35}N_4O_5S]^+$ 539.2.

A 10 mL scintillation vial, equipped with a stir bar, was charged with crude ylide **S14,** and the vial was evacuated and filled with $N_2$. To the vial was added anhydrous acetone (1.0 mL) and the resulting mixture stirred under an inert atmosphere. To the stirring solution was added methyl iodide (0.140 g, 0.990 mmol, 3 eq.) and stirring continued for 16 hours at room temperature. The resulting mixture was then concentrated under reduced pressure using a rotary evaporator, and the residue was washed with cold diethyl ether. The crude product was purified by HPLC Method B to yield the desired product **8a** as pumpkin-orange solid. (Yield: 0.075g, 34%)

**$^1$H NMR** (500 MHz, $CD_3OD$) δ: 8.00-7.97 (m, 2H), 7.86-7.83 (m, 1H), 7.31-7.27 (m, 1H), 6.64-6.61 (m, 1H), 6.50 (s, 1H), 5.22 (s, 2H), 4.37 (m, 0.4H), 4.33 (m, 0.5H), 4.24 (m, 1.6H), 4.20 (m, 1.5H), 3.84 (s, 3H), 3.57 (s, 1.5H), 3.52 (s, 1.5H), 3.24 (m, 0.6H), 3.23 (m, 0.4H), 2.92 (m, 0.5H), 2.85 (m, 0.5H), 2.70 (s, 6H), 1.81-1.77 (m, 1.7H), 1.53-1.45 (m, 5H), 1.28-1.10 (m, 1.8H) ppm.

**$^{13}$C NMR** (126 MHz, $CD_3CN$) δ: 164.8, 157.7, 157.5, 157.4, 153.8, 153.4, 152.4, 134.1, 131.6, 122.0, 117.9, 115.4, 109.6, 101.8, 68.9, 68.7, 65.8, 56.1, 37.8, 36.8, 29.0, 28.72, 28.38, 25.98, 19.0 ppm.

**$^{19}$FNMR** (471 MHz, $CD_3CN$) δ: -76.86 ppm.

**FT-IR:** 1683.8, 1635.7, 1599.5, 1555.4, 1433.1, 1335.6, 1276.9, 1243.6, 1200.9, 1133.2, 1021.8, 838.2, 800.5, 722.3 $cm^{-1}$.

**HR-MS (ESI) m/z:** Found $[M+H]^+$ 553.2479, Calc'd for $[C_{29}H_{37}N_4O_5S]^+$ 553.2479.

**MP:** 61-65 °C

**1-((((6-azidohexyl)oxy)carbonyl)(methyl)amino)-4-(7-methoxy-2*H*-chromen-3-yl)-2,6-dimethylpyridin-1-ium trifluoroacetate (8b):**

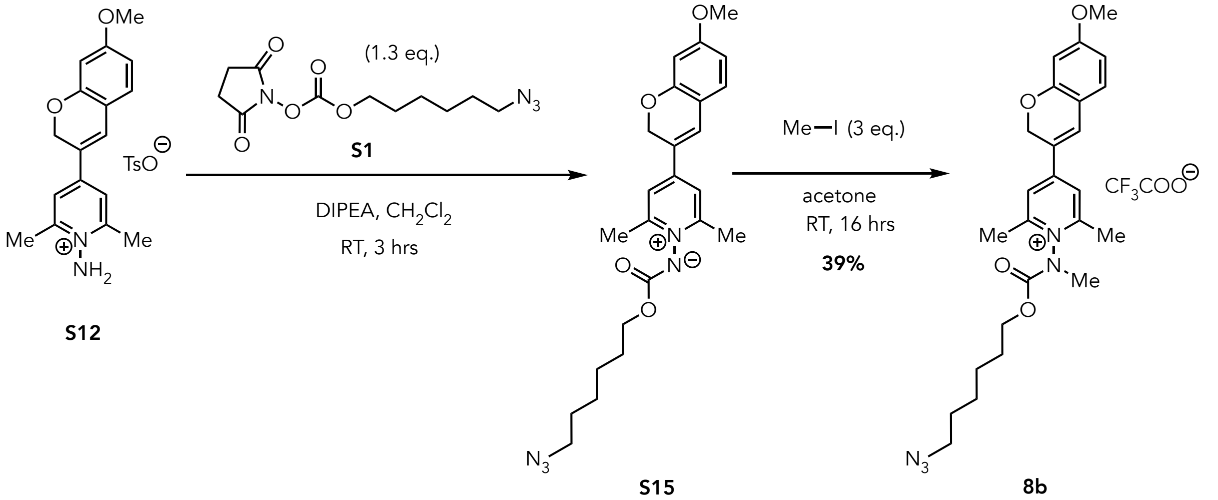


An oven-dried round bottomed flask equipped with a stir bar and sidearm inlet adapter was charged with 1-amino-4-(7-methoxy-2*H*-chromen-3-yl)-2,6-dimethylpyridinium tosylate salt **S12** (0.162 g, 0.356 mmol, 1.0 eq.) and **S1** (0.132 g, 0.463 mmol,1.3 eq.) and then the flask was evacuated and refilled with $N_2$. To the flask was added dry $CH_2Cl_2$ (1.8mL) via syringe, and the resulting mixture was stirred. To the stirring solution was added dropwise *N*,*N*-diisopropylethylamine (0.190 mL, 1.07 mmol, 3.0 eq.) via syringe, and the resulting solution was stirred vigorously at room temperature for 3 hours. The resulting mixture was concentrated using a rotary evaporator and the resulting residue was washed with ether and to yield crude residue containing **S15** that used for next step without further purification. The presence of **S15** was verified by LRMS.

**LR-MS (ESI) m/z:** Found $[M+H]^+$ 452.4 Calc'd for $[C_{24}H_{30}N_5O_4]^+$ 452.2

A 10 mL scintillation vial, equipped with a stir bar, was charged with crude ylide **S15,** and the vial was evacuated and filled with $N_2$. To the vial was added anhydrous acetone (1.2 mL) and the resulting mixture stirred under an inert atmosphere. To the stirring solution was added methyl iodide (0.151 g, 0.107 mmol, 3 eq.) and the resulting mixture was allowed to stir for 16 hours at room temperature. The resulting mixture was then concentrated using a rotary evaporator and the residue was washed with cold diethyl ether. The crude product was purified by HPLC Method B to yield the desired product **8b** as orange residue. (Yield: 0.080 g, 39%)

**$^1$H NMR** (500 MHz, $CD_3CN$) δ: 7.84 (d, $J$ = 8.7 Hz, 2H), 7.82-7.79 (m, 1H), 7.31-7.29 (m, 1H), 6.65-6.63 (m, 1H), 6.53 (s, 1H), 5.21 (s, 2H), 4.28 (t, $J$ = 6.5 Hz, 1H), 4.14 (t, $J$ = 6.4 Hz, 1H), 3.82 (s, 3H), 3.47 (s, 1.5H), 3.43 (s, 1.5H), 3.31 (t, $J$ = 6.8, 1H), 3.19 (t, $J$ = 6.8, 1H), 2.64 (s, 6H), 1.76-1.72 (m 1H), 1.64-1.58 (m, 1H), 1.51-1.42 (m, 4H), 1.24-1.19 (m, 1H), 1.15- 1.11 (m, 1H) ppm.

**$^{13}$C NMR** (126 MHz, $CD_3CN$) δ: 164.8, 157.6, 157.5, 153.8, 153.4, 152.3, 134.1, 131.7, 122.0, 115.4, 109.6, 101.8, 68.8, 68.6, 65.8, 56.1, 51.6, 37.8, 36.8, 28.9, 28.5, 26.5, 26.2, 25.4, 19.1.ppm.

**$^{19}$FNMR** (471 MHz, $CD_3CN$) δ: -75.97 ppm.

**FT-IR:** 2097.3, 1731.9, 1686.5, 1635.8, 1599.2, 1554.9, 1465.8, 1336.9, 1277.7, 1200.4, 1129.4, 800.1, 717.9 cm$^{-1}$.

**HR-MS (ESI) m/z:** Found $[M]^+$ 466.2446, Calc'd for $[C_{25}H_{32}N_5O_4]^+$ 466.2449.

#### Synthetic Route for 9

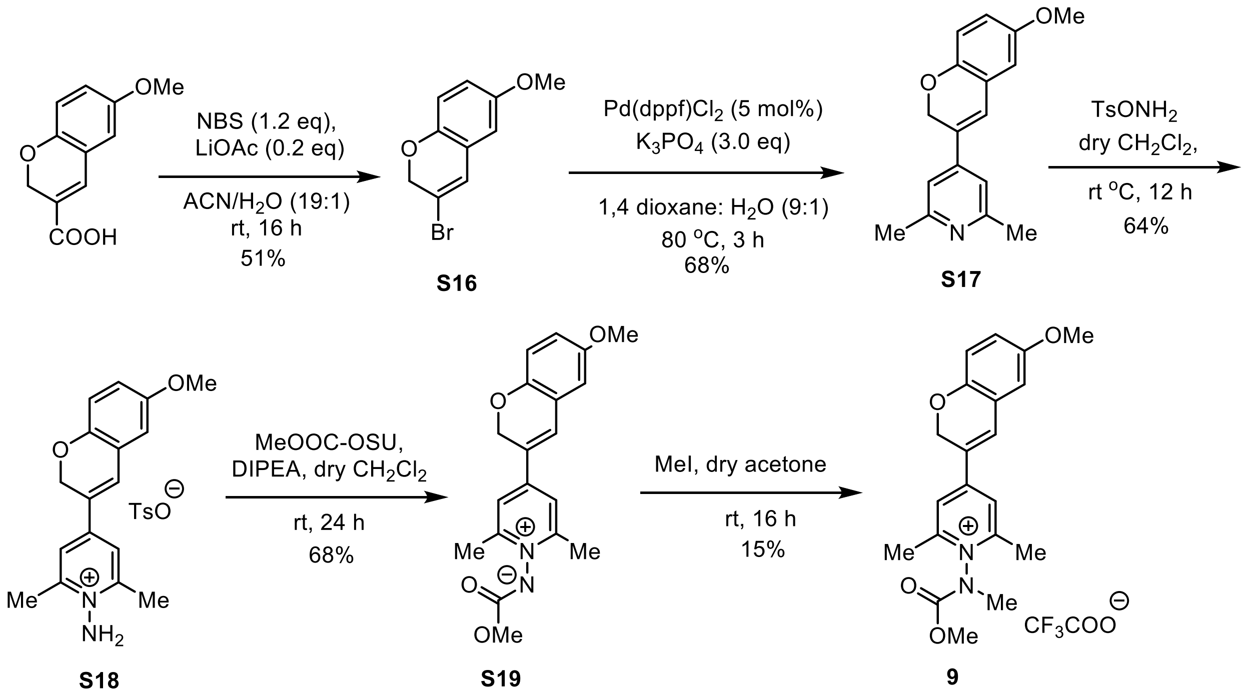


#### 3-bromo-6-methoxy-2*H*-chromene (S16):

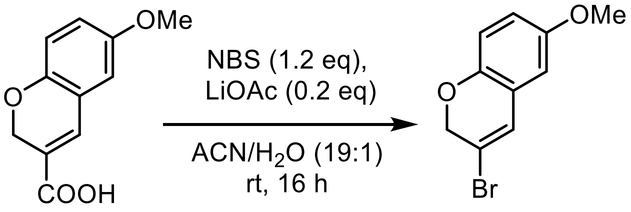


**S16**

The synthesis of 3-bromo-6-methoxy-2*H*-chromene **S16** was synthesized according to the following procedure adapted from Chen *et al.*[13] To a solution of LiOAc (0.032 g, 0.48 mmol) in a mixture of $CH_3CN/H_2O$ (19:1) was added 6-methoxy-2*H*-chromene-3-carboxylic acid[2] (0.500 g, 2.43 mmol), followed by portion-wise addition of *N*-bromosuccinimide (0.474 g, 2.66 mmol) over 10 min. The resultant suspension was stirred at 30 °C for 16 h, and then the reaction mixture was concentrated to dryness using a rotary evaporator. Purification of the crude product by silica gel column chromatography (5-10% EtOAc in hexanes) gave 3-bromo-6-methoxy-2*H*-chromene **S16** as a white solid (Yield 0.295 g, 51%).

**$^1$H NMR** (500 MHz, $CD_3CN$) δ: 6.83 (d, $J$ = 1.6 Hz, 1H), 6.74 – 6.70 (m, 2H), 6.65 – 6.61 (m, 1H), 4.81 (m, 2H), 3.72 (s, 3H) ppm.

**$^{13}$C NMR** (126 MHz, $CD_3CN$) δ: 155.5, 146.6, 126.8, 124.0, 117.1, 117.0, 115.5, 112.2, 70.9, 56.2 ppm.

**FT-IR**: 3197.7, 3103.9, 3022.9, 2993.7, 2223.6, 1598.4, 1486.1 $cm^{-1}$.

**HR-MS (ESI) m/z:** Found $[M+H]^+$ 240.9858, Calc'd for $[C_{10}H_{10}BrO_2]^+$ 240.9864

**MP**: 218 – 220 °C

**4-(6-methoxy-2*H*-chromen-3-yl)-2,6-dimethylpyridine (S17):**

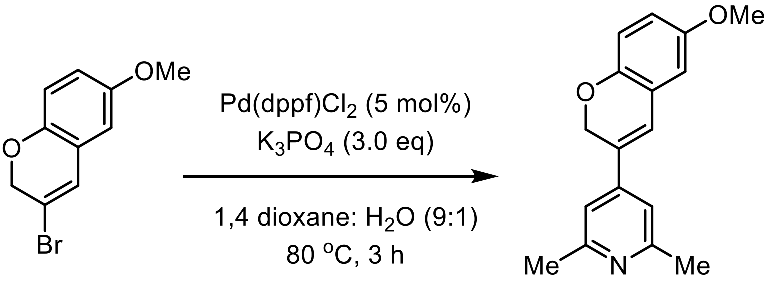


**S16** **S17**

A round-bottom flask equipped with a stir bar and reflux condenser, and a sidearm inlet adapter was dried under vacuum and refilled with $N_2$. To the flask 3-bromo-6-methoxy-2*H*-chromene **S16** (2.810 g, 11.66 mmol), 6-dimethylpyridin-4-yl)boronic acid (1.757 g, 11.66 mmol), $Pd(dppf)Cl_2$ (0.426 g, 0.583 mmol), and $K_3PO_4$ (7.425 g, 34.98 mmol) were added and the flask was evacuated and refilled with $N_2$. To the flask was added 1,4-dioxane/water (9:1, 35 mL) and the resulting mixture was heated to 80 °C with stirring for 4 hr. The reaction mixture was then cooled to rt, diluted with EtOAc (100 mL) and filtered through a celite pad. The filtrate was washed with water (20 mL X 2) and brine (20 mL), dried over $Na_2SO_4$, filtered, and concentrated using a rotary evaporator. The crude residue obtained was purified using silica gel column chromatography (30-40% of EtOAc in hexanes) to give 4-(6-methoxy-2*H*-chromen-3-yl)-2,6-dimethylpyridine **S17** as an off-white solid (Yield 2.132 g, 68%).

**$^1$H NMR** (500 MHz, $CDCl_3$) δ: 6.96 (d, *J* = 2.7 Hz, 2H), 6.92 (s, 1H), 6.80 (dd, *J* = 8.7, 2.02 Hz, 1H), 6.75 – 6.71 (m, 1H), 6.68 – 6.66 (m, 1H), 5.06 – 5.03 (m, 2H), 3.78 (d, *J* = 2.1 Hz, 3H), 2.55 (d, *J* = 2.0 Hz, 6H) ppm.

**$^{13}$C NMR** (126 MHz, $CDCl_3$) δ: 158.4, 154.6, 147.8, 144.6, 130.6, 123.3, 123.1, 116.4, 115.8, 115.4, 112.5, 66.6, 55.9, 24.7 ppm.

**FT-IR**: 3158.8, 2919.7, 2856.1, 1610.3, 1571.7, 1492,6, 1436.7, 1274.7, 1261.2, 1218.8, 1033.6, 991.2, 865.8, 817.7, 761.8 $cm^{-1}$.

**HR-MS (ESI) m/z:** Found $[M+H]^+$ 268.1329, Calc'd for $[C_{17}H_{18}NO_2]^+$ 268.1337

**MP**: 103–105 °C

**1-amino-4-(6-methoxy-2*H*-chromen-3-yl)-2,6-dimethylpyridin-1-ium tosylate (S18)**:

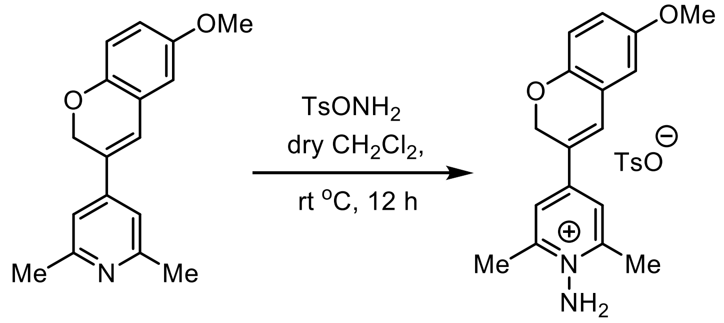


**S17** **S18**

A round bottomed flask equipped with a sidearm inlet adapter was charged with **S17** (0.740 g, 2.77 mmol) and evacuated and refilled with $N_2$. To the flask was added freshly prepared O-tosylhydroxylamine[11] (13 mL, 0.64 M solution in anhydrous $CH_2Cl_2$), and the resulting reaction mixture was stirred for 12 h at rt. The reaction mixture was concentrated using a rotary evaporator and the resulting residue was purified using silica gel column chromatography (45-60% of EtOAc in hexanes) to give 1-amino-4-(6-methoxy-2*H*-chromen-3-yl)-2,6-dimethylpyridin-1-ium tosylate **S18** as an orange solid (Yield 0.805 g, 64%).

**$^1$H NMR** (500 MHz, $CD_3OD$) δ: 7.88 (s, 2H), 7.69 (d, *J* = 7.9 Hz, 2H), 7.56 (s, 1H), 7.22 (d, *J* = 7.9 Hz, 2H), 6.90 – 6.81 (m, 3H), 5.13 (s, 2H), 3.78 (s, 3H), 2.82 (s, 6H), 2.36 (s, 3H) ppm.

**$^{13}$C NMR** (126 MHz, $CD_3OD$) δ: 156.2, 155.7, 149.9, 149.8, 143.6, 141.7, 131.0, 129.8, 128.0, 127.0, 123.7, 122.6, 118.9, 117.6, 114.1, 66.4, 56.2, 30.7, 21.3, 20.1 ppm.

**FT-IR**: 3505.9, 3197.4, 3137.6, 3041.2, 2971.7, 2911.9, 1623.7, 1616.0, 1579.4, 1444.4, 1191.7, 1035.6, 1006.6, 929.5, 713.5 $cm^{-1}$.

**HR-MS (ESI) m/z:** Found $[M]^+$ 283.1440, Calc'd for $[C_{17}H_{19}N_2O_2]^+$ 283.1446

**MP**: 88-90 °C

**(4-(6-Methoxy-2*H*-chromen-3-yl)-2,6-dimethylpyridin-1-ium-1-yl)(methoxycarbonyl)amide (S19):**

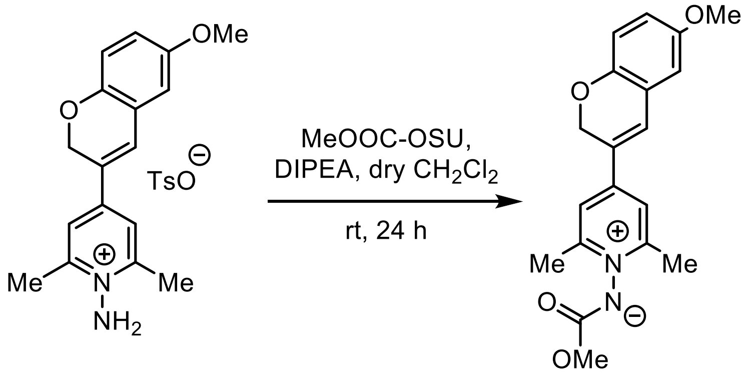


**S18** **S19**

To a round bottom flask charged with 1-amino-4-(6-methoxy-2*H*-chromen-3-yl)-2,6-dimethylpyridin-1-ium tosylate **S18** (0.750 g, 1.65 mmol) and 2,5-dioxopyrrolidin-1-yl methyl carbonate[4] (0.432 g, 2.48 mmol) were dissolved in dry $CH_2Cl_2$ (6.6 mL) at rt under $N_2$. To the stirring solution DIPEA (0.861 mL, 4.96 mmol) was added dropwise and the resulting solution was stirred vigorously at room temperature for 24 hours. The resultant mixture was then concentrated to dryness using a rotary evaporator. The residue was purified by normal phase column chromatography using DCM: MeOH as eluent to yield the desired ylide **S19** as an orange (Yield 0.385 g, 68%).

**$^1$H NMR** (500 MHz, $CD_3CN$) δ: 7.60 (s, 2H), 7.30 (m, 1H), 6.81 – 6.79 (m, 3H), 5.07 (s, 2H), 3.75 (s, 3H), 3.62 (s, 3H), 2.53 (s, 6H) ppm.

**$^{13}$C NMR** (126 MHz, $CD_3CN$) δ: 161.1, 156.4, 155.5, 149.0, 147.0, 129.1, 128.2, 123.3, 121.7, 117.3, 113.6, 66.3, 56.2, 53.0, 19.8 ppm.

**FT-IR**: 1742.0, 1617.7, 1572.0, 1493.5, 1469.1, 1437.5, 1262.8, 1225.9, 1037.3 $cm^{-1}$.

**HR-MS (ESI) m/z:** Found $[M+H]^+$ 341.1496, Calc'd for $[C_{19}H_{21}N_2O_4]^+$ 341.1501

**MP**: 69-71 °C

**4-(6-Methoxy-2H-chromen-3-yl)-1-((methoxycarbonyl)(methyl)amino)-2,6-dimethylpyridin-1-ium trifluoroacetate salt (9):**

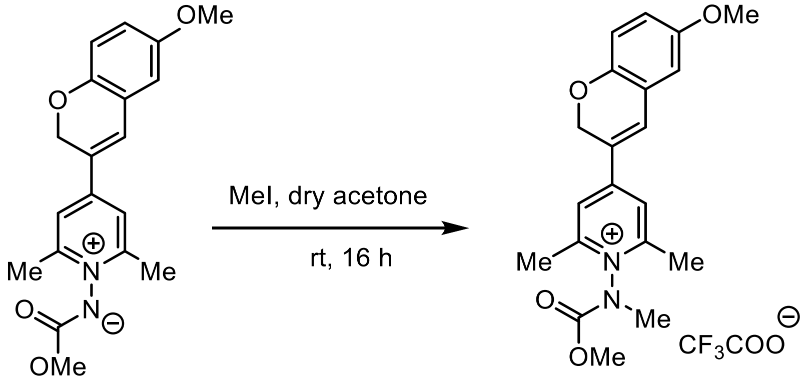


A scinitillation vial (10 mL) was charged with ylide **S13** (0.150 g, 0.441 mmol) and the vial was evacuated and filled with $N_2$. To the vial was added anhydrous acetone (1.5 mL) to dissolve the ylide and stirred vigorously under $N_2$. To the stirring solution was added methyl iodide (0.085 mL, 1.3 mmol) and the resulting mixture was stirred vigorously for 16h at room temperature. The reaction mixture was then concentrated using a rotary evaporator and the residue was washed with cold diethyl ether. The crude product was purified by reverse phase autoflash method C to get desired *N*-alkylated pyridinium **9** as an orange solid (Yield 0.031 g, 15%).

**$^1$H NMR** (500 MHz, $CD_3CN$) δ: 7.93 – 7.86 (m, 2H), 7.73 – 7.65 (m, 1H), 6.90 – 6.86 (m, 3H), 5.14 (s, 2H), 3.90 (s, 1.63H, rotamer), 3.78 (s, 3H), 3.73 (s, 1.35H, rotamer), 3.47 (2 s, 3H, rotamer), 2.66 (d, *J* = 1.7 Hz, 6H) ppm.

**$^{13}$C NMR** (126 MHz, $CD_3CN$) δ: 158.6, 158.4, 155.7, 154.5, 153.8 (153.7), 153.1, 149.8 (149.8), 134.0 (133.9), 127.2 (127.1), 123.4 (123.4), 123.04, 119.9 (119.9), 117.7, 114.1 (114.1), 66.0, 56.4, 55.7 (55.6), 38.2 (37.3), 19.3 (19.3) ppm.

**$^{19}$F NMR** (471 MHz, $CD_3CN$) δ: -75.53 ppm.

**FT-IR**: 3091.3, 2952.4, 2850.2, 1739.5, 1685.5, 1635.3, 1616.1, 1571.7, 1494.5, 1336.4, 1189.9, 1141.7, 1035.6, 802.3, 707.7 cm$^{-1}$.

**HR-MS (ESI) m/z:** Found $[M]^+$ 355.1652, Calc'd for $[C_{20}H_{23}N_2O_4]^+$ 355.1657

**MP**: 77-79 °C

**4-(6-methoxy-2H-chromen-3-yl)-2,6-dimethyl-1-(methyl(((5-((3aS,4S,6aR)-2-oxohexahydro-1H-thieno[3,4-d]imidazol-4-yl)pentyl)oxy)carbonyl)amino)pyridin-1-ium trifluoroacetate salt (9a):**

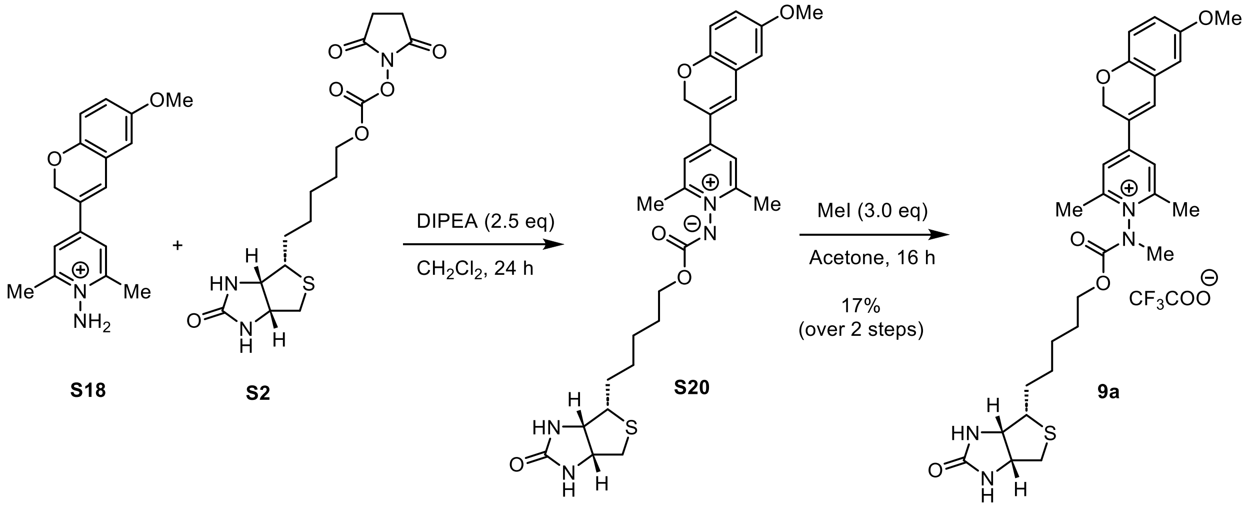


To a round bottom flask charged with **S18** (0.230 g, 0.507 mmol) and **S2** (0.226 g, 0.609 mmol) was added $CH_2Cl_2$ (1.7 mL) at rt under $N_2$ and resulting solution was stirred. To the stirring solution was added DIPEA (0.210 mL, 1.27 mmol) dropwise and the resulting mixture was stirred vigorously at rt for 24 hours. The reaction mixture was concentrated to remove volatiles using a rotary evaporator to yield crude **S20** which was used directly for the next step without any additional purification. The presence of **S20** in the crude material was confirmed by LC/MS (method A).

**LR-MS (ESI) m/z:** Found $[M+H]^+$ 539.5, Calc'd for $[C_{28}H_{34}N_4O_5S]^+$ 539.2

Crude ylide **S20** was transferred into a scinitillation vial (10 mL) equipped with a stir bar and the vial was evacuated and filled with $N_2$. To the vial was added anhydrous acetone (1.5 mL) and the resulting solution was stirred vigorously under $N_2$. To the stirring solution was added methyl iodide (0.098 mL, 1.5 mmol) and the resulting mixture was stirred vigorously for 16h at room temperature. The reaction mixture was then concentrated using a rotary evaporator and the crude residue was washed with cold diethyl ether to remove impurities. The crude product was purified by reverse phase auto flash method C to get desired product **9a** as an orange solid (Yield (0.058 g, 17%).

**$^1$H NMR** (500 MHz, $CD_3CN$) δ: 7.86 (d, $J$ = 9.9 Hz, 2H), 7.67– 7.64 (m, 1H), 6.94 – 6.88 (m, 4H), 5.23 – 5.14 (m, 3H), 4.43 – 4.37 (m, 1H), 4.30– 4.37 (m, 1H), 4.16 – 4.15 (m, 1H), 3.78 (s, 3H), 3.63– 3.58 (m, 1H), 3.48 – 3.44 (m, 3H), 3.19 – 3.06 (m, 2H), 2.91 – 2.81 (m, 2H), 2.66 – 2.65 (m, 6H), 2.60– 2.58 (m 3H), 1.76 – 1.14 (m, 8H) ppm.

**$^{13}$C NMR** (126 MHz, $CD_3CN$) δ: 158.5, 158.3, 155.5, 153.6, 152.5, 149.7, 133.9, 133.6, 126.9, 123.2, 123.1, 122.9, 119.8, 119.7, 117.6, 114.0, 114.0, 69.2, 69.0, 65.9, 62.4, 60.7, 56.3, 56.3, 56.2, 56.2, 40.8, 38.0, 37.0, 29.2, 29.0, 28.6, 26.2, 19.3, 19.2, 14.3 ppm.

**$^{19}$F NMR** (471 MHz, $CD_3CN$) δ: -75.92 ppm.

**HR-MS (ESI) m/z:** Found $[M]^+$ 553.2473, Calc'd for $[C_{29}H_{37}N_4O_5S]^+$ 553.2484

**FT-IR:** 3307.3, 3259.1, 3079.7, 2942.8, 1685.4, 1637.2, 1616.1, 1571.7, 1201.4, 1182.1, 1130.0, 829.2, 800.3, 723.1.

**MP**: 82–85 °C

**Synthesis Scheme for Probe 10:**

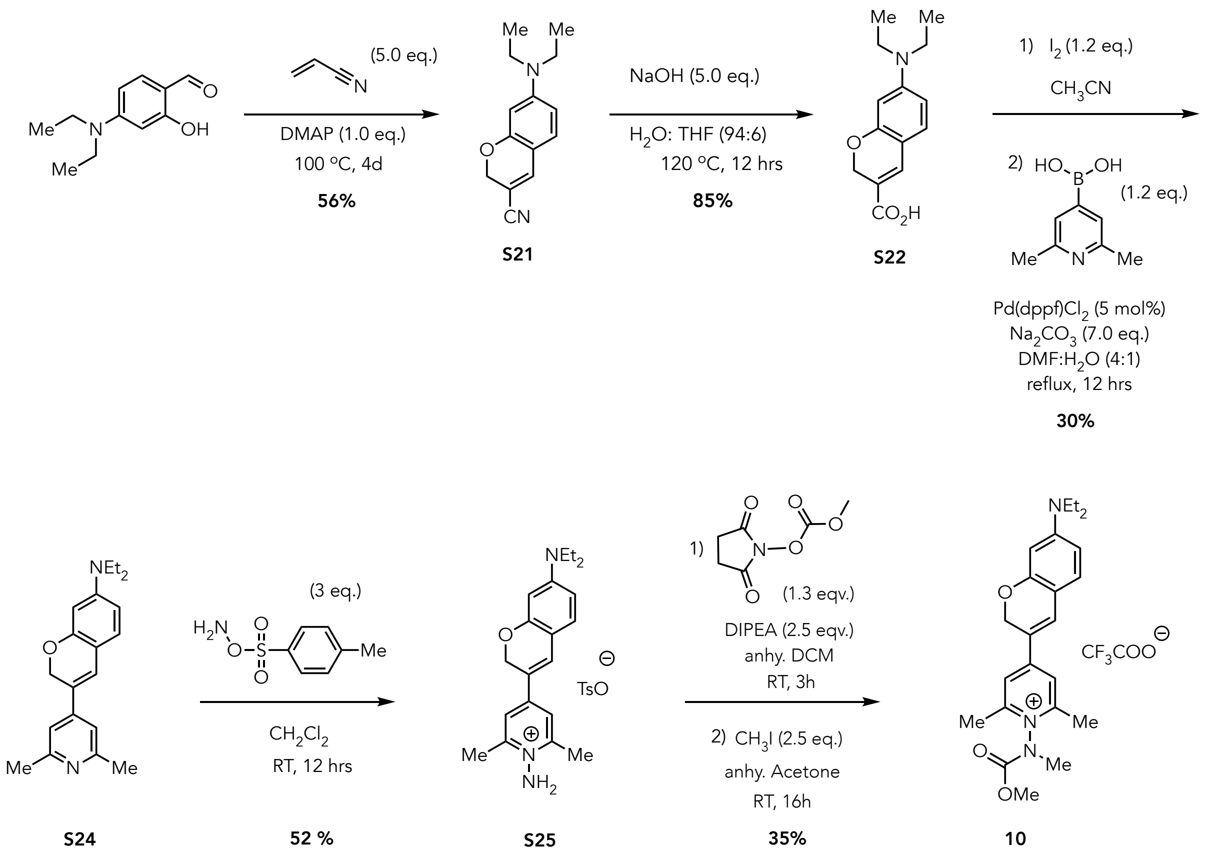

**7-(diethylamino)-2*H*-chromene-3-carbonitrile (S21):**

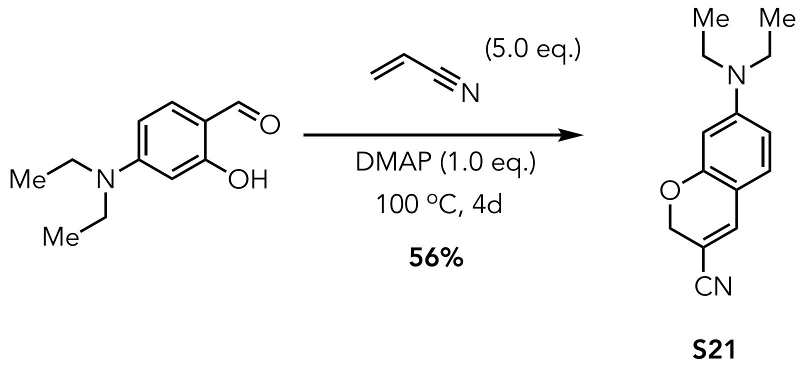

**S21** was synthesized according to the following procedure adapted from Ikeda *et al.* as follows.[14] To a stirring solution of 4-(Diethylamino)-2-hydroxybenzaldehyde (10.098 g, 52.253 mmol, 1.0 eq.) in acrylonitrile (20mL) in a 75 ml screw cap vial was added DMAP (6.421g, 52.56 mmol, 1.0 eq.). The resulting mixture was stirred for 4 days at 100°C. The resulting mixture was cooled to room temperature and was quenched with 1M HCl (aq), extracted with EtOAc. The organic layer was washed with water and brine, dried over $Na_2SO_4$, filtered, and concentrated using a rotary evaporator. The crude was purified by silica gel column chromatography (10% EtOAc: hexane to obtain **S21** as a yellow solid (Yield 6.714 g, 56%).

**$^1$H NMR** (500 MHz, $CD_3CN$) δ: 7.19 (s, 1H), 6.98 (d, *J* = 8.7 Hz, 1H), 6.32 (dd, *J* = 8.7, 2.6 Hz, 1H), 6.13 (d, *J* = 2.6 Hz, 1H), 4.71 (s, 2H), 3.36 (q, *J* = 7.0, 7.0, 7.0 Hz, 4H), 1.12 (t, *J* = 7.0, 7.0 Hz, 6H) ppm.

$^{13}$**C NMR** (125 MHz, $CD_3CN$) δ: 157.3, 152.4, 140.1, 130.9, 109.6, 106.7, 98.6, 95.6, 65.2, 45.1, 12.8 ppm.

**FT-IR:** 2971.0, 2929.2, 2198.2, 1600.1, 1536.4, 1518.1, 1469.6, 1404.7, 1373.8, 1353.0, 1271.8, 1200.5, 1158.3, 1142.8, 1111.4, 1076.6, 1012.7, 939.4, 899.8, 827.6, 786.9, 700.4 $cm^{-1}$.

**HR-MS (ESI) m/z:** Found $[M+H]^+$ 229.1333, Calcd for $[C_{14}H_{16}N_2O]^+$ 229.1335

**MP:** 68-70 °C

**7-(diethylamino)-2*H*-chromene-3-carboxylic acid (S22):**

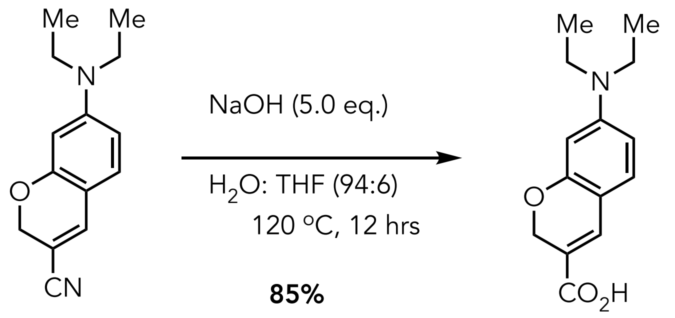


**S21** **S22**

**S22** was synthesized according to the following procedure adapted from Ikeda *et al.* as follows.[14] A mixture of 7-(diethylamino)-2*H*-chromene-3-carbonitrile (2.361 g, 10.34 mmol, 1.0 eq.), NaOH (2.068 g, 51.70 mmol, 5.0 eq.), and 94:6 $H_2O$: THF (9.0 mL) was stirred at 120°C for 12h. The reaction mixture was cooled to room temperature and then diluted with $H_2O$. The resulting mixture was then chilled in an ice bath and acidified with HCl (1M) to pH~1 (measured by pH paper). The desired compound was precipitated in the aqueous phase and was isolated by filtration, washed with water, and then dried under vacuum to yield the desired product **S22** as a light brown solid (Yield 2.164 g, 85%.8.750 mmol)

$^{1}$**H NMR** (500 MHz, MeOD) δ: 7.38 (s, 1H), 7.00 (d, $J$ = 8.5 Hz, 1H), 6.30 (dd, $J$ = 8.6, 2.5 Hz, 1H), 6.12 (d, $J$ = 2.6 Hz, 1H), 4.8 (d, $J$ = 1.2 Hz, 2H), 3.39 (q, $J$ = 7.1, 7.1, 7.0 Hz, 4H), 1.16 (t, $J$ = 7.1, 7.1 Hz, 6H) ppm.

$^{13}$**C NMR** (125 MHz, MeOD) δ: 167.6, 157.1, 151.1, 134.4, 130.0, 115.7, 109.4, 105.1, 97.5, 64.3, 44.0, 11.6 ppm.

**FT-IR:** 2972.6, 2585.9, 1664.0, 1599.0, 1539.2, 1517.6, 1428.9, 1404.9, 1377.3, 1309.5, 1277.4, 1244.9, 1210.1, 1188.0, 1156.2, 1136.0, 1080.8, 822.7 $cm^{-1}$.

**HR-MS (ESI) m/z:** Found $[M+H]^+$ 248.1279, Calc'd for $[C_{14}H_{18}NO_3]^+$ 248.1281

**MP:** 215-220°C

**3-(2,6-dimethylpyridin-4-yl)-*N*,*N*-diethyl-2*H*-chromen-7-amine (S24):**

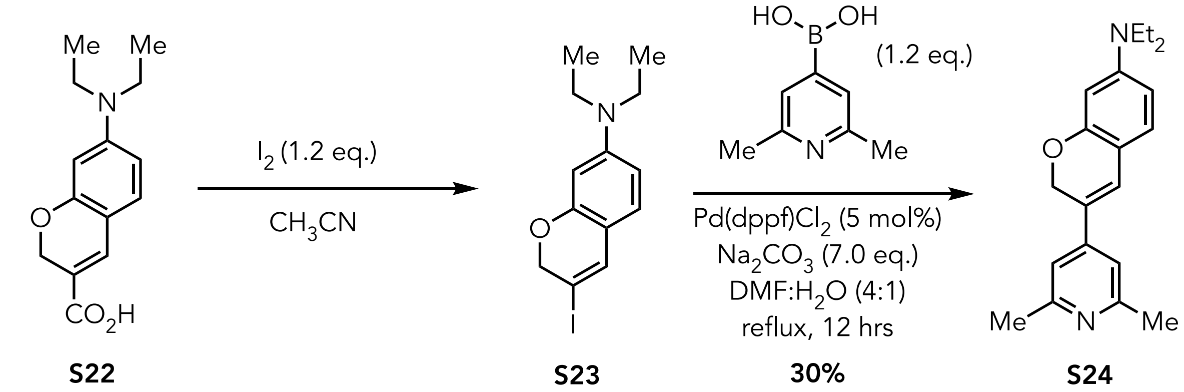

**S24** was synthesized using the two-step sequence as follows:
*N*,*N*-diethyl-3-iodo-2*H*-chromen-7-amine (**S23**) was synthesized by adapting the procedure of Larossa as follows.[15] A stirring solution of 7-(diethylamino)-2*H*-chromene-3-carboxylic acid (1.982 g, 8.014 mmol, 1.0 eq.) in acetonitrile (40 mL) was chilled in an ice bath. To the chilled, stirring solution was added iodine dropwise using a dropping addition funnel ([$I_2$]= 0.240 M in $CH_3CN$, 40 mL, 9.6 mmol, 1.2 eq) over 10 hrs. Then the reaction mixture was warmed up to room temperature and allowed to stir for 12 hrs. The reaction mixture was quenched by adding 15% aq. $Na_2S_2O_3$ solution and extracted with EtOAc (x1), and the combined organic layers were washed with $NaHCO_3$ (aq.), dried over $Na_2SO_4$, filtered, and concentrated using a rotary evaporator. Owing to the poor stability of **S23**, the crude residue containing **S23** was immediately used in the next step without further purification. The presence of **S23** was verified by LRMS.

**LR-MS (ESI) m/z:** Found $[M+H]^+$ 330.3, Calc'd for $[C_{13}H_{16}INO]^+$ 330.0

A round-bottom flask equipped with a stir bar was charged with the crude residue containing *N*,*N*-diethyl-3-iodo-2*H*-chromen-7-amine (**S23**) (0.870 g, 2.664 mmol, 1.0 eq.) and purged with nitrogen for 5 minutes. To the purged flask was added sequentially 2,6-dimethylpyridin-4-yl)boronic acid (0.482 g, 3.20 mmol, 1.2 eq.), Pd(dppf)$Cl_2$ (0.087 g, 5 mol%), and DMF (8 mL), and the solution was stirred. To this mixture was added $Na_2CO_3$ (aq) ([$Na_2CO_3$]= 9.34 M, 2 mL, 18.680 mmol, 7.0 eq.) and the reaction mixture and stirred for 12h at 85 °C using a reflux condenser. The reaction mixture was then cooled to room temperature and extracted with EtOAc (x3), and the combined organic layers were washed with water and brine, dried over $Na_2SO_4$, filtered, and the solvent was removed using a rotary evaporator. The resulting crude material was purified by silica gel column chromatography (30% EtOAc: hexane) to obtain **S24** as an orange residue (Yield 0.724 g, 30%).

**$^1$H NMR** (500 MHz, MeOD) δ: 7.11 (s, 1H), 7.10 (s, 1H), 6.29 (dd, *J* = 8.6, 1.4 Hz, 1H), 6.15 (d, *J* = 2.5 Hz, 1H), 5.03(s, 2H), 3.37 (q, *J* = 7.0, 7.0, 7.0 Hz, 4H), 1.16 (t, *J* = 7.1, 7.1 Hz, 6H) ppm.

**$^{13}$C NMR** (125 MHz, MeOD) δ: 157.3, 155.7, 149.9, 146.1, 128.7, 124.3, 122.0, 115.1, 110.7, 105.1, 97.7, 65.8, 44.0, 22.3, 11.6 ppm.

**IR:** 2969.9, 2926.0, 1616.4, 1594.8, 1545.1, 1517.0, 1467.7, 1402.4, 1374.7, 1355.8, 1279.2,1204.4, 1181.4, 1129.5, 1077.6 ppm.

**HR-MS (ESI) m/z:** Found $[M+H]^+$ 309.1957, Calc'd for $[C_{20}H_{24}N_2O]^+$ 309.1961

**1-amino-4-(7-(diethylamino)-2*H*-chromen-3-yl)-2,6-dimethylpyridin-1-ium tosylate (S25)**

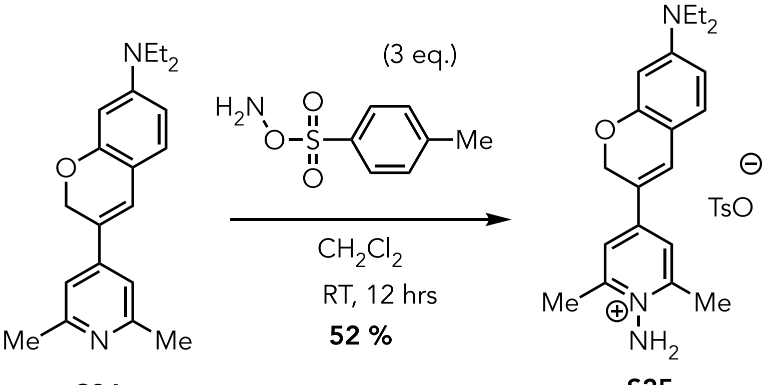


**S24** **S25**

A round-bottomed flask equipped with a stir bar was charged with 3-(2,6-dimethylpyridin-4-yl)-*N*,*N*-diethyl-2*H*-chromen-7-amine (0.452 g, 1.46 mmol, 1.0 eq.) and purged with $N_2$.. To the flask was added freshly prepared O-tosylhydroxylamine[11] (2M solution in anhydrous $CH_2Cl_2$), and the resulting mixture was stirred for 12 h at rt. The reaction mixture was concentrated using a rotary evaporator, and the resulting residue was purified using silica gel column chromatography (10% MeOH/$CH_2Cl_2$) to yield **S25** as a bright orange semi-solid **S25** (Yield 0.373 g, 52%).

**$^1$H NMR** (500 MHz, $CD_3CN$) δ: 7.57 (d, *J* = 8.2 Hz, 2H), 7.51 (d, *J* = 3.2 Hz, 3H), 7.13 (d, *J* = 8.0 Hz, 2H), 7.08 (d, *J* = 8.7 Hz, 1H), 6.37 (dd, *J* = 8.7, 2.6 Hz, 1H), 6.18 (d, *J* = 2.6 Hz, 1H), 5.89 (s, 2H), 5.06 (s, 2H), 3.40 (q, *J* = 7.1, 7.1, 7.0 Hz, 4H), 2.66 (s, 6H), 2.32 (s, 6H) 1.15 (t, *J* = 7.1, 7.1 Hz, 6H) ppm.

**$^{13}$C NMR** (125 MHz, $CD_3CN$) δ: 156.9, 154.4, 151.8, 148.9, 145.5, 138.7, 131.5, 130.8, 128.3, 125.6, 119.4, 110.1, 106.0, 97.3, 65.3, 44.3, 20.3, 19.5, 11.9 ppm.

**FT-IR:** 2970.0, 2927.8, 1622.0, 1586.6, 1522.5, 1457.3, 1413.1, 1376.6, 1348.9, 1174.9, 1122.8, 1077.8, 1033.9, 1011.2, 682.0 cm$^{-1}$.

**HR-MS (ESI) m/z:** Found $[M]^+$ 324.2067, Calc'd for $[C_{20}H_{26}N_3O]^+$ 324.2070

**4-(7-(diethylamino)-2*H*-chromen-3-yl)-1-((methoxycarbonyl)(methyl)amino)-2,6-dimethylpyridin-1-ium (10)**

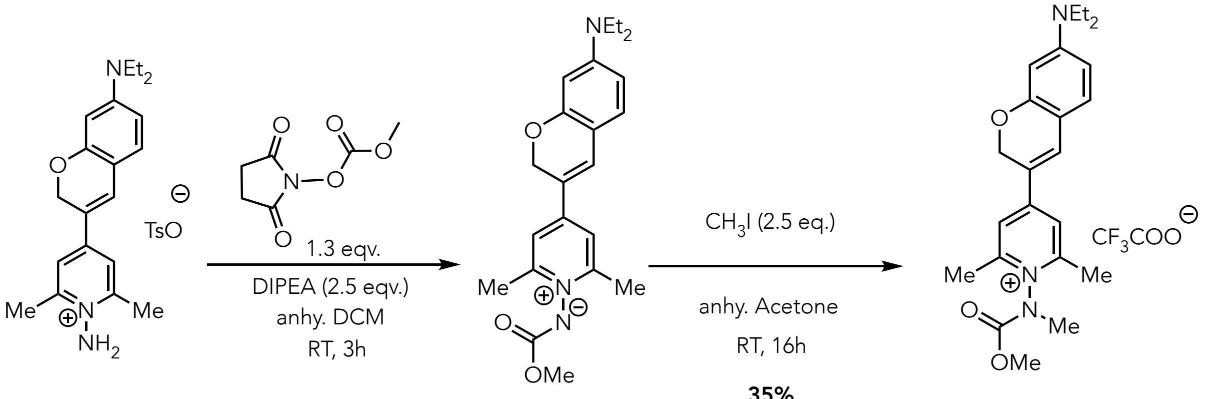


**S25** **S26** **10**

1-amino-4-(7-(diethylamino)-2*H*-chromen-3-yl)-2,6-dimethylpyridin-1-ium tosylate (S25) (0.356g, 0.716 mmol, 1.0 eq.) and 2,5-dioxopyrrolidin-1-yl methyl carbonate (0.167g, 0.930 mmol, 1.3.) were taken in a round-bottom flask purged with $N_2$. To the flask, dry DCM (2 mL) was added via syringe, and the resulting mixture was stirred. To the stirring solution, DIPEA (0.365 mL, 2.15 mmol, 3.0 eq.) was added dropwise, and the resulting solution was stirred vigorously at room temperature for 3 hours. The resultant mixture was then concentrated to dry under reduced pressure. The crude residue containing **S26** was immediately used in the next step without further purification, to utilize the ylide form for the next alkylation step. The presence of **S26** was verified by LRMS.

**LR-MS (ESI) m/z:** Found $[M+H]^+$ 382.4, Calc'd for $[C_{22}H_{27}N_3O_3]^+$ 382.0

A round-bottom flask equipped with a stir-bar purged with $N_2$ and was charged with crude (4-(7-(diethylamino)-2*H*-chromen-3-yl)-2,6-dimethylpyridin-1-ium-1-yl)(methoxycarbonyl)amide (**S26**) (0.108 g, 0.283 mmol, 1.0 eq.). To the flask was added anhydrous acetone (1.0 mL), and methyl iodide (0.04 mL, 0.708 mmol, 2.5 eq.) sequentially. The resulting mixture was stirred for 16 h at room temperature. The resultant mixture was then concentrated to dry under reduced pressure. The crude product was purified using Method A in reverse-phase flash chromatography (0.1% TFA in water-acetonitrile) to isolate a purple, semi-solid product **10** (Yield 0.078 g, 35%).

**$^1$H NMR** (500 MHz, $CD_3CN$) δ: 7.75 (d, *J* = 5.3 Hz, 1H), 7.60 (d, *J* = 3.8 Hz, 1H), 7.17 (dd, *J* = 8.7, 1.7 Hz, 1H), 6.46-6.42 (m, 1H), 6.22 (d, *J* = 2.6 Hz, 1H), 5.16 (s, 2H), 3.92 (s, 1.5H), 3.75 (s, 1.5H), 3.51-3.43 (m, 7H), 2.58 (s, 6H) 1.19 (t, *J* = 7.1, 7.1 Hz, 6H) ppm.

**$^{13}$C NMR** (125 MHz, $CD_3CN$) δ: 158.5, 156.7, 156.5, 154.8, 153.2, 153.2, 136.1, 135.9, 132.8, 132.7, 120.6, 120.5, 108.3, 66.1, 55.5, 55.5, 49.8, 46.0, 46.0, 38.3, 37.4, 30.8, 19.1, 19.0, 12.6 ppm.

**$^{19}$F NMR** (471MHz, $CD_3CN$) δ: -76.49 ppm.

**FT-IR:** 1813.3, 1789.0, 1739.2, 1718.6, 1575.70, 1519.5, 1439.9, 1325.8, 1261.0, 1207.9, 1122.0, 1078.0, 1033.2, 1010.5, 923.0, 816.9, 712.6, 681.9 $cm^{-1}$.

**HR-MS (ESI) m/z:** $[M]^+$ Calcd for $C_{23}H_{30}N_3O_3$ 396.2281; Found 396.2281

The **S26** residue can be purified by Method A in reverse-phase flash column chromatography (0.1% TFA in water-acetonitrile) to yield the desired product (**S26**) as a red semi-solid.

**$^1$H NMR** (500 MHz, $CD_3CN$) δ: 7.93- 7.80 (m, 2H), 7.66 (s, 1H), 7.50-7.42 (m, 1H), 7.12-67.03 (m, 1H), 5.30 (s, 2H), 3.92 (s, 1.5H), 3.90 (s, 3H), 3.65-3.45 (m, 4H), 2.71 (s, 6H) 1.15 (t, *J* = 7.2, 7.2 Hz, 6H) ppm.
**$^{13}$C NMR** (125 MHz, $CD_3CN$) δ: 159.2, 158.9, 156.6, 154.4, 152.5, 131.4, 130.9, 119.3, 114.8, 112.5, 109.9, 66.2, 54.8, 53.8, 19.5, 10.4 ppm.
**FT-IR:** 1813.3, 1789.1, 1736.9, 1440.2, 1368.7, 1261.9, 1238.8, 1203.2, 1096.0, 923.0, 813.3, 769.8 $cm^{-1}$.

**HR-MS (ESI) m/z:** Found $[M]^+$ 382.2120, Calc'd for $[C_{22}H_{28}N_3O_3]^+$ 382.2125

### Synthetic Route for 11

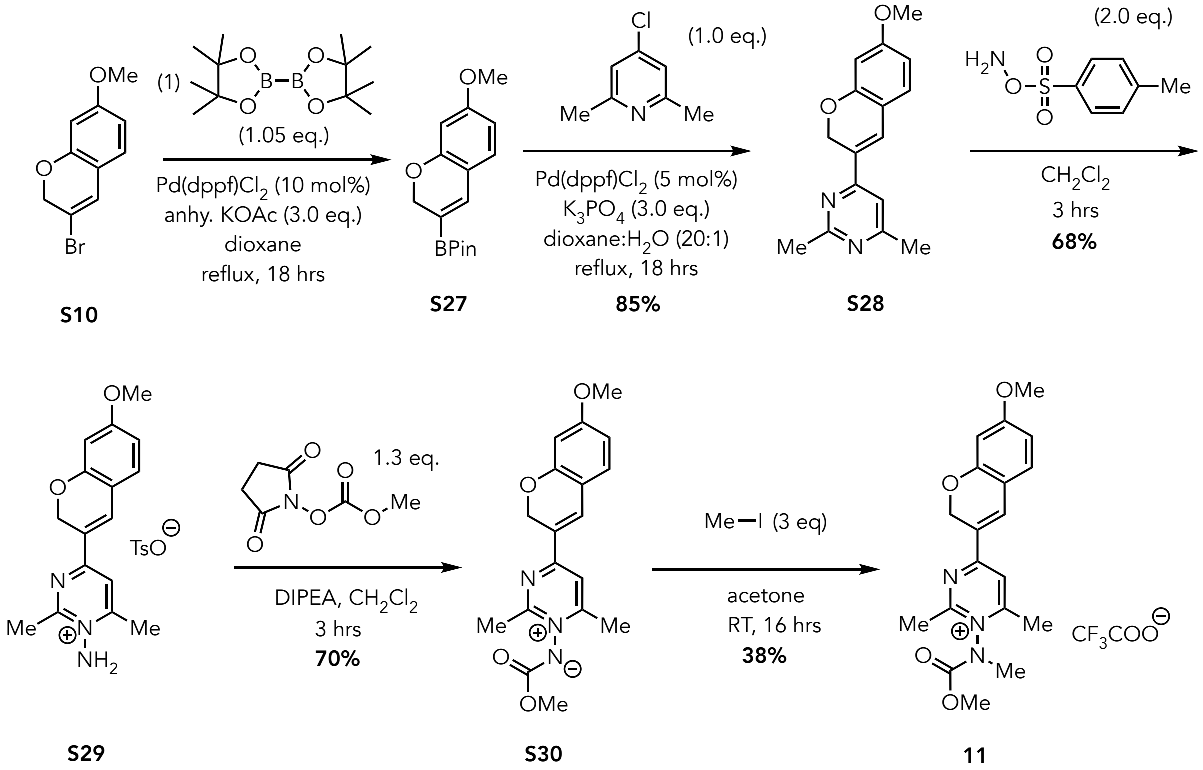


#### 4-(7-methoxy-2*H*-chromen-3-yl)-2,6-dimethylpyrimidine (S28)

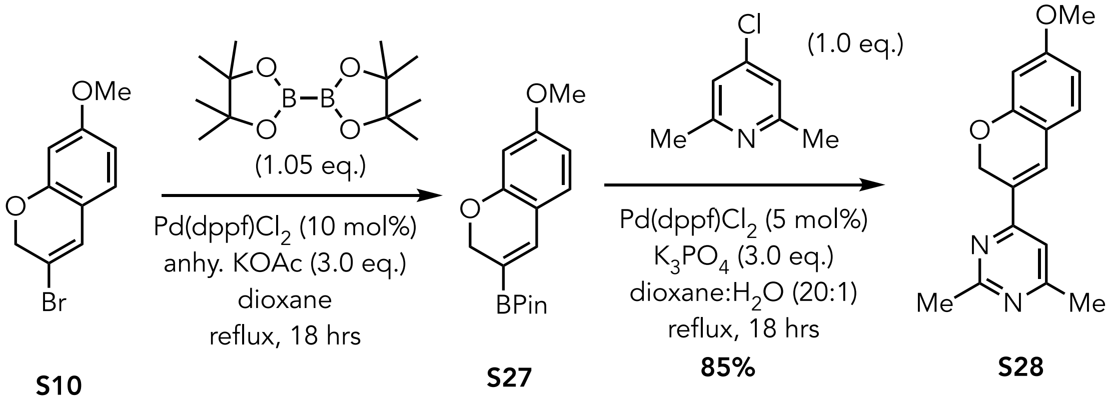


A vacuum dried round bottom flask equipped with stir bar, reflux condenser, and side arm inlet adapter was charged with 3-bromo-7-methoxy-2*H*-chromene (1.160 g, 4.811 mmol, 1.0 eq.), 4,4,4',4',5,5,5',5'-octamethyl-2,2'-bi(1,3,2-dioxaborolane) (1.466 g, 5.773 mmol, 1.2 eq.), Pd(dppf)$Cl_2$ (0.352 g, 0.481 mmol, 10.0 mol%), and anhydrous KOAc (1.416 g, 14.43 mmol, 3.0 eq.) and the flask was evacuated and refilled with $N_2$. To flask was added 1,4-dioxane (10 mL) and resulting mixture was heated to reflux for 18 hours under $N_2$ with continuous stirring. The reaction mixture was cooled down to room temperature, and the solid was removed by filtration through Celite. The filter cake was washed with EtOAc and the filtrate was then washed with $H_2O$ and brine. The organic layer was dried over $Na_2SO_4$, filtered, and the solvent removed using a rotary evaporator to yield a crude residue containing **S27** that was used directly without purification for the next step. The presence of **S27** was verified by $^1$H-NMR and LRMS.

**$^1$H NMR** (500 MHz, $CDCl_3$) δ: 7.07 (s, 1H), 6.93 (d, *J* = 8.4 Hz,1H), 6.42 (dd, *J* = 8.4, 2.5 Hz,1H), 6.36 (s,1H), 4.84 (s, 2H), 3.77 (s,3H), 1.29 (s,12H) ppm.

**LR-MS (ESI) m/z:** Found $[M+H]^+$ 289.3 calcd for $[C_{16}H_{22}BO_4]^+$ 289.1

An oven dried 50 mL round bottom flask equipped with a stir bar, reflux condenser, and side arm inlet adapter was charged with 2-(7-methoxy-2*H*-chromen-3-yl)-4,4,5,5-tetramethyl-1,3,2-dioxaborolane **S27** (1.102 g, 3.824 mmol,1.2 eq.), 4-chloro-2,6-dimethylpyrimidine (0.454 g, 3.19 mmol, 1.0 eq.), Pd(dppf)$Cl_2$ (0.140 g, 0.191 mmol, 5.0 mol%), and $K_3PO_4$ (2.029 g, 9.561 mmol, 3.0 eq.) and the flask was evacuated and refilled with $N_2$. To flask was added 10 mL of 1,4-dioxane:$H_2O$ (20:1, v/v) and the resulting mixture was heated to reflux for 18 hours under $N_2$ with continuous stirring. The reaction mixture was cooled down to room temperature and the solid was removed by filtration through Celite. The filter cake was washed with EtOAc and the filtrate was then washed with $H_2O$ and brine. The organic layer was dried over $Na_2SO_4$, filtered, and the solvent removed using a rotary evaporator. The crude residue obtained was purified by column chromatography using 40% EtOAc:Hexane to yield the desired product as a yellow solid. (Yield: 0.728 g, 85%)

**$^1$H NMR** (500 MHz, $CDCl_3$) δ: 7.39 (s, 1H), 7.08-7.06 (m, 2H), 6.50 (dd, *J* = 8.3, 2.4 Hz, 1H), 6.45 (s, 1H), 5.25 (s, 2H), 3.80 (s, 3H), 2.68 (s, 3H), 2.50 (s, 3H) ppm.

**$^{13}$C NMR** (126 MHz, $CDCl_3$): δ167.7, 166.6, 162.2, 156.4,129.2, 126.4, 115.3, 111.4, 108.1, 101.6, 65.8, 55.6, 26.2, 24.4 ppm.

**FT-IR:** 1595.8, 1569.2, 1553.0, 1503.2, 1442.3, 1380.7, 1302.9, 1274.3, 1193.7, 1156.9, 1131.3, 1115.7, 1030.0, 844.3 $cm^{-1}$.

**HR-MS (ESI) m/z:** Found $[M+H]^+$ 269.1281, Calc'd for $[C_{16}H_{17}N_2O_2]^+$ 269.1285

**MP:** 78-80 °C

**1-amino-4-(7-methoxy-2*H*-chromen-3-yl)-2,6-dimethylpyrimidinium tosylate (S29):**

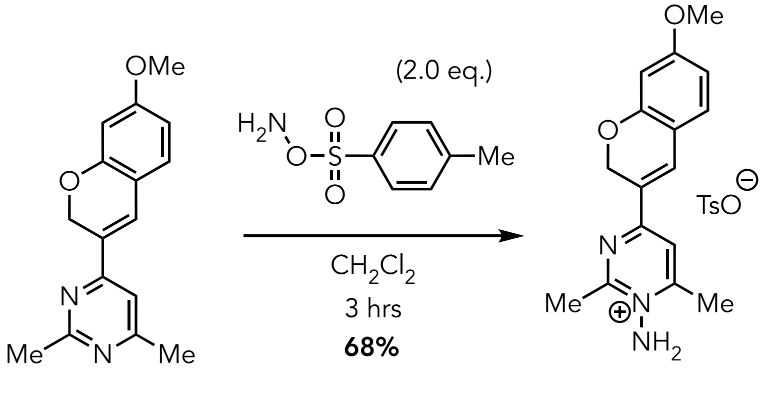


**S28** **S29**

A vacuum dried round bottomed flask equipped with a sidearm inlet adapter was charged with 4-(7-methoxy-2*H*-chromen-3-yl)-2,6-dimethylpyrimidine **S28** (0.719 g, 2.68 mmol, 1.0 eq.), and evacuated and refilled with $N_2$. To the flask was added freshly prepared O-tosylhydroxylamine[11] (in 8 mL in anhydrous $CH_2Cl_2$, 0.68 M, 5.4 mmol, 2.0 eq.,), and the resulting mixture was stirred for 3 hours at rt. The reaction mixture was concentrated using a rotary evaporator, and the resulting residue was 1:1 mixture of EtOAc in hexane (25 x 3 mL). The resulting residue was then sonicated with 5% MeOH in 4:1 mixture of $CH_2Cl_2$:Ether. The supernatant was decanted, and the sonication process was repeated twice. Subsequently, the residue was dried under vacuum to yield1-amino-4-(6-methoxy-2*H*-chromen-3-yl)-2,6-dimethylpyrimidin-1-ium tosylate **S29** (0.8290g, 68% yield) as an orange solid.

**$^1$H NMR** (500 MHz, $CD_3OD$) δ: 8.00 (d, *J* = 9.5 Hz, 2H), 7.69 (d, *J* = 7.9 Hz, 2H), 7.30 (d, *J* = 8.6 Hz, 1H), 7.28-7.23 (m, 2H), 6.62 (s, 1H), 6.51 (s, 1H), 5.26 (s, 2H), 3.85 (s, 3H), 2.96 (s, 3H), 2.80 (s, 3H), 2.37 (s, 3H) ppm.

**$^{13}$C NMR** (126 MHz, $CD_3OD$) δ: 166.0, 165.1, 163.9, 163.4, 159.0, 143.5, 141.7, 136.5, 132.3, 129.8, 126.9, 124.2, 116.6, 116.1, 110.1, 102.4, 65.7, 56.2, 22.6, 21.3, 19.9 ppm.

**FT-IR:** 3151.2, 1593.1, 1552.1, 1503.8, 1446.0, 1311.1, 1277.3, 1199.9, 1176.2, 1121.8, 1033.1, 1033.1, 1010.1, 817.2, 683.3 $cm^{-1}$.

**HR-MS (ESI) m/z:** Found $[M]^+$ 284.1392, Calc'd for $[C_{16}H_{18}N_3O_2]^+$ 284.1393

**MP:** 140-145 °C

**(4-(7-methoxy-2*H*-chromen-3-yl)-2,6-dimethylpyrimidin-1-ium-1-yl)(methoxycarbonyl)-amide (S30):**

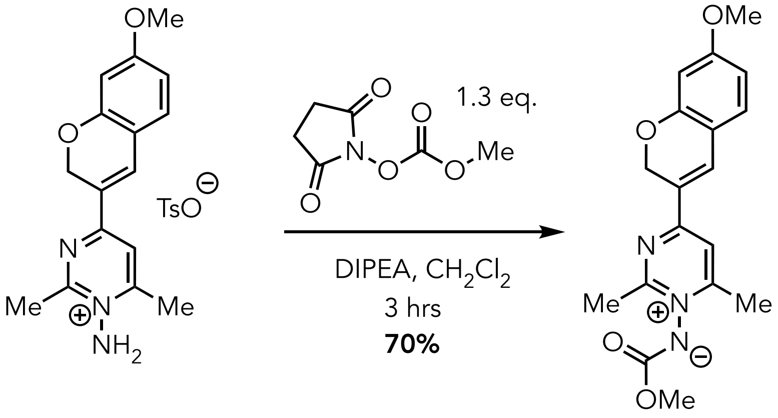


**S29** **S30**

An oven-dried round bottomed flask equipped with a stir bar and sidearm inlet adapter was charged with 1-amino-4-(7-methoxy-2*H*-chromen-3-yl)-2,6-dimethylpyrimidinium tosylate salt **S29** (0.345 g, 0.757 mmol, 1.0 eq.) and 2,5-dioxopyrrolidin-1-yl methyl carbonate[3] (0.170 g, 0.984mmol, 1.3eq.) and the flask was evacuated and refilled with $N_2$. To the flask was added dry $CH_2Cl_2$ (3.8 mL) via syringe, and the resulting mixture was stirred. To the stirring solution was added dropwise *N*,*N*-diisopropylethylamine (396.0 μL, 2.267 mmol, 3.0 eq.) and the resulting solution was stirred vigorously at room temperature for 3 hours. The resultant mixture was then concentrated to dryness using a rotary evaporator. The residue was purified by normal phase column chromatography using 5% MeOH:$CH_2Cl_2$ as an eluent to yield the desired ylide product **S30** as reddish-brown residue (Yield: 0.181 g, 70%).

**$^1$H NMR** (500 MHz, $CD_3CN$) δ: 7.78-7.74 (s, 2H), 7.24 (d, $J$ = 8.4 Hz, 1H), 6.59 (dd, $J$ = 8.3, 2.5 Hz 1H), 6.49 (s, 1H), 5.24 (s, 2H), 3.81 (s, 3H), 3.62 (s, 3H), 2.69 (s, 3H), 2.55 (s, 3H) ppm.

**$^{13}$C NMR** (126 MHz, $CD_3CN$) δ: 166.8, 166.3, 165.7, 165.1, 159.2, 139.2, 133.0, 123.7, 116.6, 115.8, 110.5, 102.3, 65.6, 56.7, 54.8, 22.3, 19.7 ppm.

**FT-IR:** 1680.9, 1591.5, 1541.2, 1436.9, 1377.9, 1276.9, 1202.9, 1180.9, 1136.5, 839.7, 801.6, 723.7, 695.9 $cm^{-1}$.

**HR-MS (ESI) m/z:** [Found $[M+H]^+$ 342.1445, Calc'd for $[C_{18}H_{20}N_3O_4]^+$ 342.1448

**4-(7-methoxy-2*H*-chromen-3-yl)-1-((methoxycarbonyl)(methyl)amino)-2,6-dimethylpyrimidinium trifluoroacetate (11):**

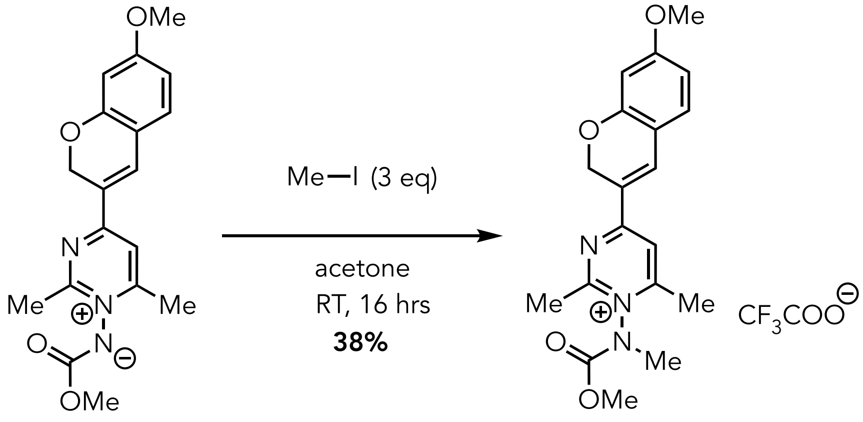


**S30** **11**

A scintillation vial (10 mL) equipped with a stir bar was charged with ylide **S30** (0.170 g, 0.498 mmol,1.0 eq.) and the vial was evacuated and filled with $N_2$. To the vial was added acetone (1.5mL) and the resulting mixture stirred under an inert atmosphere. To the stirring solution was added methyl iodide (93.0 μL, 1.49 mmol, 3 eq.) the resulting mixture was allowed to stir for 16 hours at room temperature. The resulting mixture was then concentrated using a rotary evaporator and the residue was washed with cold diethyl ether. The crude product was purified by HPLC Method A to yield **11** as rust-orange solid. (Yield: 0.088g, 38%)

**$^1$H NMR** (500 MHz, $CD_3CN$) δ: 8.15 (s, 1H), 7.86 (s, 1H), 7.35 (d, *J* = 8.5 Hz, 1H), 6.65 (d, *J* = 8.7 ,1H), 6.53 (s, 1H), 5.28 (s, 2H), 3.90 (s, 1.6H), 3.85 (s, 3H), 3.74 (s, 1.5H), 3.47 (s, 1.6H), 3.44 (s, 1.4H), 2.77(s, 3H), 2.61(s, 3H) ppm.

**$^{13}$C NMR** (126 MHz, $CD_3CN$) δ: 166.3, 165.5, 163.8, 161.7, 159.0, 154.3, 152.7, 140.4, 133.0, 122.9, 120.5, 116.8, 115.9, 115.4, 110.3, 101.8, 65.1, 56.3, 55.4, 37.7, 36.8, 30.5, 21.5, 18.7 ppm.

**$^{19}$FNMR** (471 MHz, $CD_3CN$) δ: -75.60 ppm.

**FT-IR:** 1733.8, 1687.3, 1588.8, 1536.4, 1436.4, 1338.6, 1277.4, 1204.4, 1180.5, 1133.4, 1028.5 cm$^{-1}$.

**HR-MS (ESI) m/z:** Found $[M]^+$ 356.1605, Calc'd for $[C_{19}H_{22}N_3O_4]^+$ 356.1605

**MP:** 86-92 °C

**4-(7-methoxy-2*H*-chromen-3-yl)-2,6-dimethyl-1-(methyl(((5-((3a*R*,4*S*,6a*S*)-2-oxohexahydro-1*H*-thieno[3,4-*d*]imidazol-4-yl)pentyl) -oxy)carbonyl)amino)pyrimidin-1-ium trifluoroacetate (11a):**

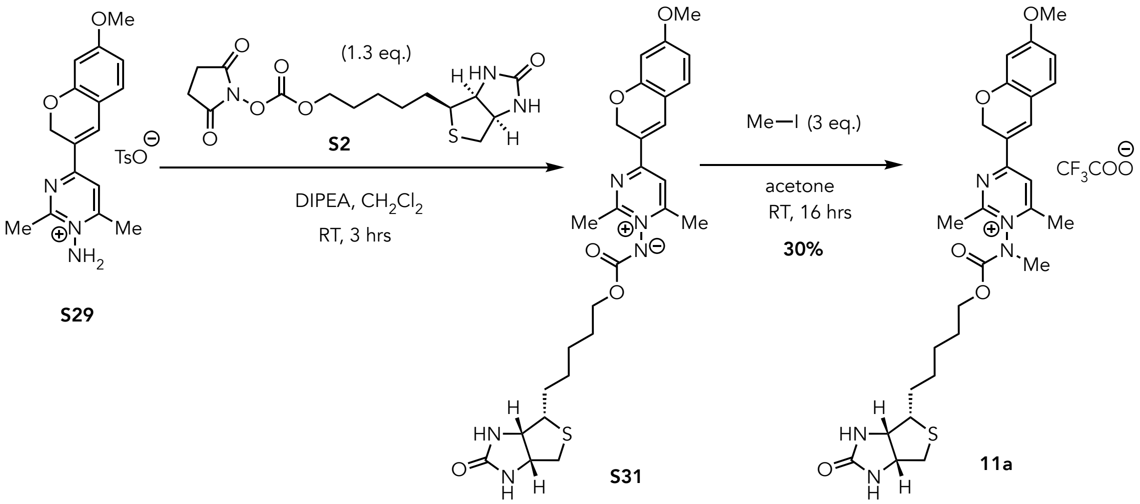


An oven-dried round bottomed flask equipped with a stir bar and sidearm inlet adapter was charged with 1-amino-4-(7-methoxy-2*H*-chromen-3-yl)-2,6-dimethylpyrimidinium tosylate salt **S29** (0.125 g, 0.274 mmol, 1.0 eq.) and **S2** (0.132 g, 0.356 mmol,1.3 eq.) and then the flask was evacuated and refilled with $N_2$. To the flask was added anhydrous $CH_2Cl_2$ (1.3 mL) *via* syringe and the resulting mixture was stirred. To the stirring solution was added dropwise *N*,*N*-diisopropylethylamine (0.140 mL, 0.822 mmol, 3.0 eq.) and the resulting solution was stirred vigorously at room temperature for 3 hours. The resulting mixture was concentrated using a rotary evaporator and the resulting residue was washed with ether and to yield crude residue containing **S31** that used for next step without further purification. The presence of **S31** was verified by LRMS.

**LR-MS (ESI) m/z:** Found $[M+H]^+$ 540.4, calcd for $[C_{27}H_{34}N_5O_5S]^+$ 540.2

A 10 mL scintillation vial, equipped with a stir bar, was charged with crude ylide **S31,** and the vial was evacuated and filled with $N_2$. To the vial was added anhydrous acetone (1.0 mL) and the resulting mixture stirred under an inert atmosphere. To the stirring solution was added methyl iodide (0.051 mL, 0.822 mmol, 3.0 eq.) and the resulting mixture was allowed to stir for 16 hours at room temperature. The resulting mixture was then concentrated using a rotary evaporator and the resulting residue was washed with cold diethyl ether. The crude product was purified by HPLC Method B to yield the desired product **11a** as rust orange solid. (Yield: 0.055 g, 30%)

**$^1$H NMR** (500 MHz, $CD_3CN$): δ 8.23-8.17 (m,1H), 7.97-7.91 (m, 1H), 7.40-7.36 (m, 1H), 6.67 (d, *J* = 10.4 Hz, 1H), 6.54 (s, 1H), 5.94 (s, 2H), 5.31 (s, 2H), 4.52-4.50 (m, 0.5H), 4.46-4.43 (m, 0.5H), 4.34-4.30 (m, 1.5H), 4.25-4.15 (m, 1.7H), 3.87 (s, 3H), 3.51 (s, 1.4H), 3.46 (s, 1.8H), 3.25-3.19 (m, 0.7H), 3.11-3.10 (m, 0.7H), 2.94-2.91 (m, 0.6H), 2.89-2.83 (m, 0.5H), 2.79 (s, 3H), 2.63 (s, 3H), 1.79-1.72 (m, 1.9H), 1.60-1.43 (m, 5H), 1.31-1.28 (m, 1.7H), ppm.

**$^{13}$C NMR** (126 MHz, $CD_3OD$): δ 167.6, 166.1, 162.9, 160.2, 154.8, 153.1, 141.7, 141.3, 133.6, 123.7, 121.0, 119.1, 117.1, 116.4, 115.2, 111.0, 102.4, 69.6, 65.7, 63.5, 61.6, 57.2, 56.4, 41.0, 37.8, 36.7, 29.9, 26.9, 21.6 ppm.

**$^{19}$FNMR** (471 MHz, $CD_3CN$): δ = -75.92 ppm.

**FT-IR:** 1679.8, 1588.0, 1535.7, 1439.1, 1331.2, 1279.2, 1205.3, 1139.6, 1019.6, 842.5, 801.5, 760.7, 724.1

**HR- MS (ESI) m/z:** Found $[M]^+$ 554.2434, Calc'd for $[C_{28}H_{36}N_5O_5S]^+$ 554.2432.

**MP:** 63-65°C

### 3. Protein labeling and mapping:

#### Labeling of Lysozyme with pyridinium and pyrimidinium salts (Probe 1-11):

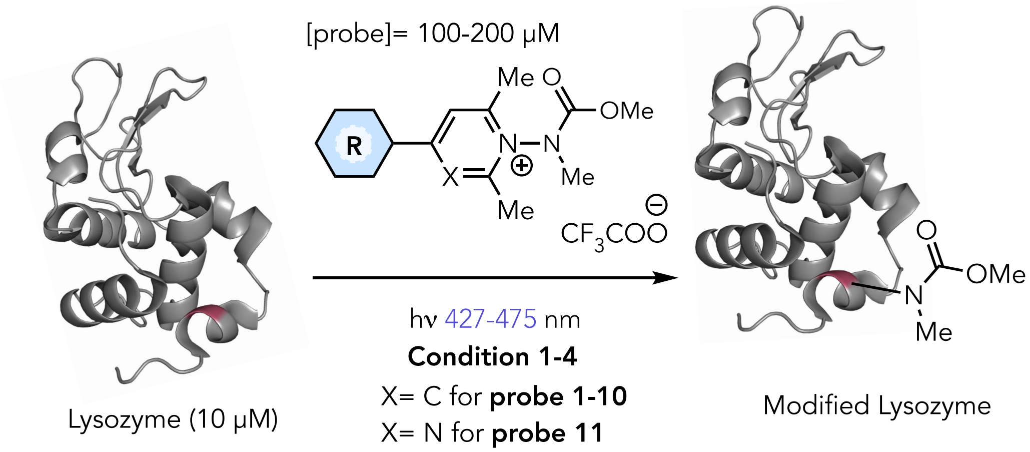


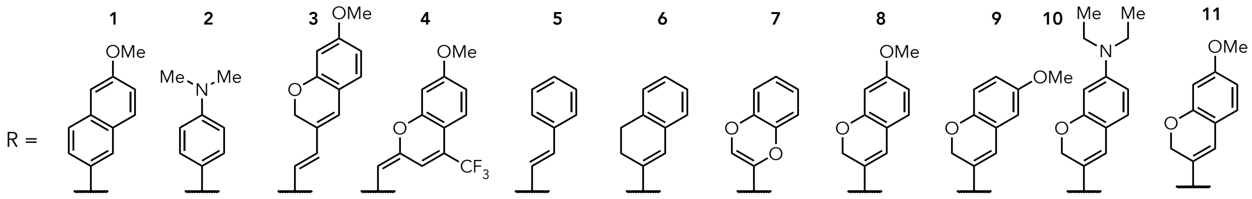


The labeling of lysozyme with the pyridinium and pyrimidinium salts (**1-11,** final concentration 100-200 µM) was screened under four distinct conditions, as outlined below.

**Condition 1:** [lysozyme]= 10 µM, [probe]= 200 µM, [GSH]=300 µM, buffer: 20 mM pH 6.9 $NH_4OAc$, irradiation time: 60 min.

**Condition 2:** [lysozyme]= 10 µM, [probe]= 100 µM, [GSH]=300 µM, buffer: 20 mM pH 6.9 $NH_4OAc$, irradiation time: 60 min.

**Condition 3:** [lysozyme]= 10 µM, [probe]= 200 µM, [GSH]=300 µM, buffer: 20 mM pH 7.4 $Na_2HPO_4$ , irradiation time: 60 min.

**Condition 4:** [lysozyme]= 10 µM, [probe]= 100 µM, [GSH]=300 µM, buffer: 20 mM pH 7.4 $Na_2HPO_4$, irradiation time: 60 min.

To a 2 mL Pyrex LC/MS vial (Thermo model # 03-391-39) were sequentially added solutions containing the lysozyme, glutathione, desired pyridinium or pyrimidinium salt (**1-11**), and respective buffer solution. The resulting solution was diluted to a final volume of 500 µL by adding deionized water and sparged using Nitrogen for a minimum of 30 minutes prior to irradiation. The actively sparging solution was subsequently placed to an EvoluChem™ PhotoRedox Box photoreactor, which was equipped with either a Kessil, PR160 427 nm (for probe **6**), PR160 456 nm (for probes **1-5** and **7-10**), or a 475 nm LED lamp (for probe **11**). Subsequently, the photoreactor was subjected to a 100% lamp intensity irradiation for a duration of 60 minutes, while active sparging was continuously maintained. The resultant reaction mixture was then analyzed directly via LC/MS.

Labelling outcomes were analyzed using LC/MS by either extracting ion chromatogram intensities (LC-MS Method A). The total values were then used to estimate percentages, which were determined based on the relative intensities of peaks or ion counts.
The following light source were used for protein photo-activation:

427 nm Light source = Kessil PR-160L-427 nm LED, 40W

456 nm Light source = Kessil PR-160L-456 nm LED, 40W

475 nm Light source = EvoluChem™ 475PF LED,18W

**LC-MS Method A**: solvents: A: 0.1% formic acid in $H_2O$ B: 0.1% formic acid in $CH_3CN$, method: 20-70% B over 5 min, 70-95% B over 1 min, hold 95% B for 2 min, 95-20% B over 0.5 min, then hold 20% B 1.5 min, column: Aeris™ 3.6 µm WIDEPORE XB-C18 20, LC Column 100 x 21 mm, m/z range: 500-2000, flow rate: 0.2 mL/min.

| | % Conversion | | | | |
|---|---|---|---|---|---|
| **Probe** | **Light wavelength** | **Condition 1** | **Condition 2** | **Condition 3** | **Condition 4** |
| **1** | 456 | 35 | 28 | 19 | 11 |
| **2** | 456 | 0 | 0 | 0 | 0 |
| **3** | 456 | 7 | 6 | 0 | 0 |
| **4** | 456 | 0 | 0 | 0 | 0 |
| **5** | 456 | 0 | 0 | 0 | 0 |
| **6** | 427 | 52 | 33 | 15 | 8 |
| **7** | 456 | 0 | 0 | 0 | 0 |
| **8** | 456 | 81 | 73 | 47 | 36 |
| **9** | 456 | 37 | 31 | 13 | 6 |
| **10** | 456 | 0 | 0 | 0 | 0 |
| **11** | 475 | 26 | 18 | 13 | 10 |

### MS-MS Analysis of Labelled Protein Conjugates

#### General Protocol for Protein Digestion with Trypsin:

The labelled protein solution was transferred to a 2 mL microcentrifuge tube, precipitated using ice-cold methanol: chloroform (4:1) mixture, and centrifuged at 20,000 g for 20 minutes at 4°C to isolate protein pellets. The protein pellets were washed with cold methanol (2x's), centrifuged, and then air-dried at RT. This protein pellets were resuspended in urea (240 µL, 10 M, 6 M final concentration), tris-buffer (140 µL, 50 mM, pH 8.0, 17.5 mM final concentration), and TCEP (20 µL, 100 mM, 5 mM final concentration). This solution was incubated for 30 minutes at 50°C. A freshly prepared solution of 2-iodoacetamide (25 µL, 250 mM, 14.7 mM final concentration) was subsequently added to the tube and incubated at room temperature for an additional 30 minutes in the dark. The reduced, Cys-capped labelled protein was then concentrated using an Amicon 10 kDa filter (final volume 40 µL). The concentrate was then diluted with urea (20 µL, 10 M, 1 M final concentration), tris base buffer (134 µL, 50 mM, pH 8.0, 33.5 mM final concentration), $CaCl_2$ (5 µL, 50 mM, 1.25 mM final concentration), and Trypsin/Lys-C (Promega) (1 µL, 0.5 µg/µL, 0.002 µg/µL final concentration) and incubated at 37°C for 20 hours. Formic acid (2.5 µL) was then added to the digested protein solution and dried on a SpeedVac (ThermoFisher Scientific). This tryptic peptide sample was desalted using Pierce™ peptide desalting spin columns (Thermo Catalog # 8952) according to manufacturer's protocol and the solution was dried on a SpeedVac.

Desalted peptides were redissolved in 0.1% FA in optima grade water (50 µL) and analyzed with a Orbitrap Exploris 240 mass-spectrometer (Thermo Fisher Scientific) using nLC-MS/MS Method.

**General LC-MS/MS Method:**
Two microliters of desalted digested peptide samples were injected into the PepMap Neo Trap column (P/N 174502, Thermo Fisher Scientific), and eluted on the PepMap RSLC C18, PN ES 900 HPLC column (3 µm, 100 Å, 75 µm x 15 cm) using the Vanquis Neo UHPLC system (Thermo Fisher Scientific) coupled to an Orbitrap Exploris 240 mass spectrometer (Thermo Fisher Scientific). Mobile phase A was 0.1% formic acid in $H_2O$ (optima grade), and mobile phase B was 0.1% formic acid, 80% acetonitrile (optima Grade) in $H_2O$ (Optima Grade). Chromatographic separation was achieved using a linear gradient starting with a flow rate of 350 nL $min^{-1}$ from 2% Buffer B (0.1% FA in MeCN) followed by incremental increase to 50% over 46 min, 50-90% B for 14 min, 90-95% B for 2 min, and hold at 95% B for 8 min. The eluted peptides were analyzed with a Orbitrap Exploris 240 mass-spectrometer (Thermo Fisher Scientific) equipped with nano-LC electrospray ionization source (applied voltage: 2.0 kV, and the ion transferred tube temperature maintained at 280 °C). Under the positive-ion mode, full-scan mass spectra were acquired over the m/z range from 380 to 2000 using the Orbitrap mass analyzer with 120k MS1 resolution, RF Lens 70%, 28 ms max injection time and 300% AGC. MS/MS fragmentation is performed in a data independent mode. MS2 features between 380-2000 amu with resolution of 15k, isolation window (m/z) 10, overlapping window set to (m/z) 1, maximum injection time 25 ms, a normalized HCD of 30%, and AGC of 100%.

**Peptide mapping and Data Analysis:**
All the DIA data of the digested protein was analyzed by FragPipe GUI v 23.0 computational Platform using MSFragger-DIA-NN Algorithm.[16] Basic search workflow was used to directly search the DIA data. The search was performed against the database (downloaded from Protein data bank, Lysozyme (from chicken egg) PDB ID: 1DPX, and alpha-chymotrypsinogen A (from bovine pancreas) PDB ID: 1YPH) with carbamidomethylated cysteine (+57.02146 Da) set as a static modification, oxidation of methionine (+15.9949 Da), acetylation of N-termini (+42.01106 Da), and *N*-methylcarbamate (+87.0320 Da) of tryptophan, histidine, tyrosine and phenylalanine set as dynamic modifications. Trypsin was specified as the proteolytic enzyme with up to two missed cleavage sites allowed. Closed search of probe modified sites was performed with precursor and fragment ion tolerances set to 10 ppm and 20 ppm, respectively. Peptide length was set to 5– 50, and peptide mass range was set to 400–5000. Search results were filtered with 1% FDR for peptide identification, using Percolator as part of the Philosopher toolkit (v5).

**Labeling of Lysozyme with Probe 8:**
Labelling of lysozyme with the Probe **8** was carried out as per the general labelling procedure under condition **A**, using the following solutions: lysozyme (35 µL, 145 µM, 10 µM final concentration), **8** (36 µL, 2.8 mM, 200 µM final concentration), $NH_4OAc$, (50 µL, 200 mM, pH 6.9, 20 mM final concentration), GSH (15 µL, 10.0 mM, 300 µM final concentration), and deionized water (364 µL). The resulting solution sparged using Nitrogen for a minimum of 30 minutes and irradiated at 456 nm for 60 min. The reaction mixture was directly analyzed with **LC-MS Method A** and was estimated to have proceeded with 81% conversion and a 5.3:1 (+1:+2) labelled ratio.

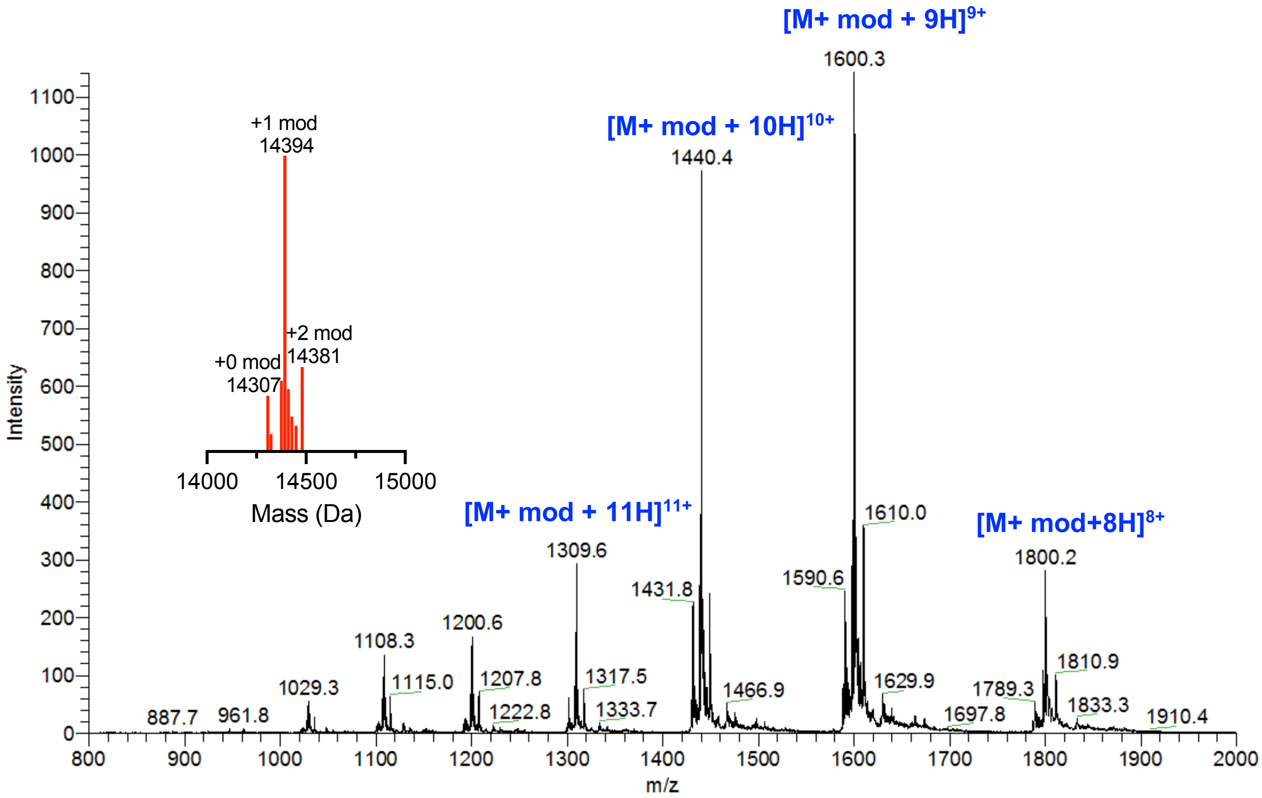


**Figure S1:** Mass spectrum of the *N*-methylcarbamate modified **lysozyme 8-conjugate** ($T_r$ 3.2-6.8 min).

**MS-MS Analysis of Lysozyme *N*-methycarbamate Modifications:**
nLC-MS/MS Method A analysis was performed on a lysozyme **8**-conjugate to know the site of modifications. Modifications were identified at **W108** and **W62**.

**W108**:

| | b | $b^{2+}$ | AA Sequence | y | $y^{2+}$ | |
|---|---|---|---|---|---|---|
| 1 | 114.09 | 57.55 | I | | | 15 |
| 2 | 213.16 | 107.08 | V | 1665.74 | 833.38 | 14 |
| 3 | 300.19 | 150.60 | S | 1566.68 | 783.84 | 13 |
| 4 | 415.22 | 208.11 | D | 1479.64 | 740.33 | 12 |
| 5 | 472.24 | 236.62 | G | 1364.62 | 682.81 | 11 |
| 6 | 586.28 | 293.65 | N | 1307.59 | 654.30 | 10 |
| 7 | 643.30 | 322.16 | G | 1193.55 | 597.28 | 9 |
| 8 | 790.34 | 395.67 | M | 1136.53 | 568.77 | 8 |
| 9 | 904.38 | 452.70 | N | 989.50 | 495.25 | 7 |
| 10 | 975.42 | 488.21 | A | 875.45 | 438.23 | 6 |
| 11 | 1248.53 | 624.77 | W* | 804.42 | 402.71 | 5 |
| 12 | 1347.60 | 674.30 | V | 531.30 | 266.16 | 4 |
| 13 | 1418.64 | 709.82 | A | 432.24 | 216.62 | 3 |
| 14 | 1604.72 | 802.86 | W | 361.20 | 181.10 | 2 |
| 15 | | | R | 175.12 | 88.06 | 1 |

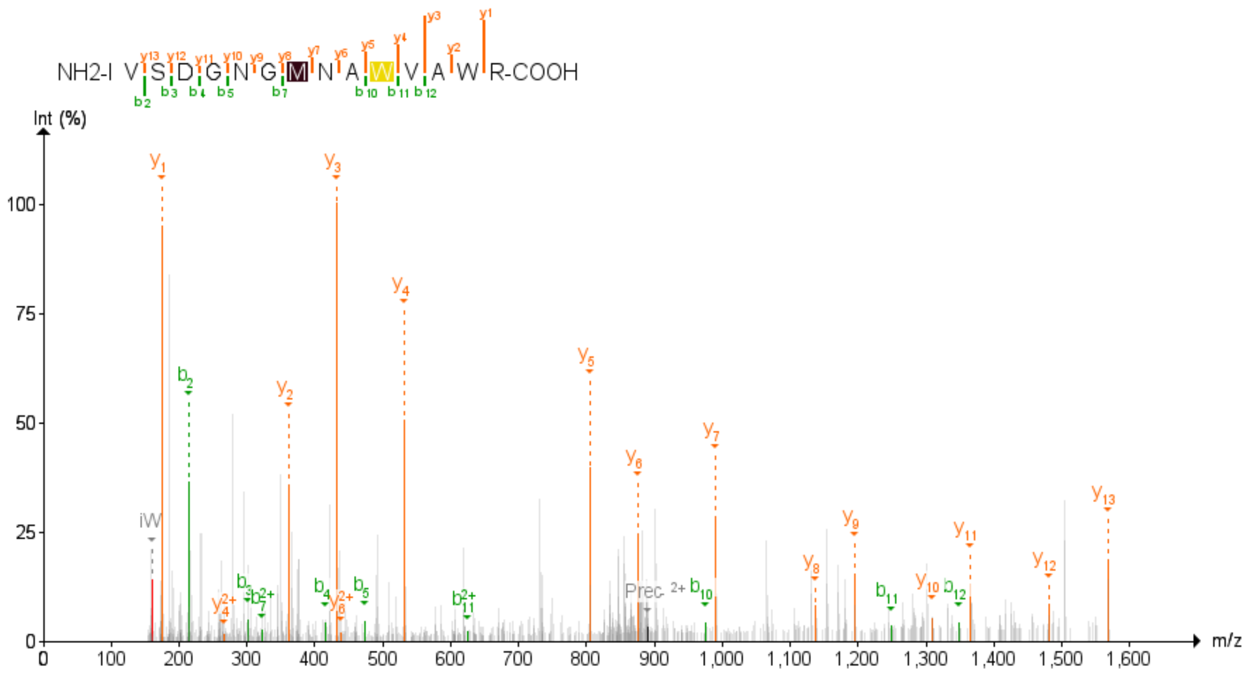


MS/MS data was obtained after analyzing the digested lysozyme **8**-conjugate sample according to the protocol outlined in nLC-MS/MS Method A

**W62**

| | b | $b^{2+}$ | AA Sequence | y | $y^{2+}$ | |
|---|---|---|---|---|---|---|
| 1 | 115.05 | 58.03 | N | | | 28 |
| 2 | 216.10 | 108.55 | T | 3199.46 | 1600.23 | 27 |
| 3 | 331.12 | 166.07 | D | 3098.41 | 1549.71 | 26 |
| 4 | 388.15 | 194.58 | G | 2983.39 | 1492.20 | 25 |
| 5 | 475.18 | 238.09 | S | 2926.36 | 1463.69 | 24 |
| 6 | 576.23 | 288.62 | T | 2839.33 | 1420.17 | 23 |
| 7 | 691.25 | 346.13 | D | 2738.29 | 1369.65 | 22 |
| 8 | 854.32 | 427.66 | Y | 2623.26 | 1312.13 | 21 |
| 9 | 911.34 | 456.17 | G | 2460.19 | 1230.60 | 20 |
| 10 | 1024.42 | 512.71 | I | 2403.17 | 1202.09 | 19 |
| 11 | 1137.51 | 569.26 | L | 2290.09 | 1145.55 | 18 |
| 12 | 1265.56 | 633.29 | Q | 2177.01 | 1089.01 | 17 |
| 13 | 1378.65 | 689.83 | I | 2048.95 | 1024.98 | 16 |
| 14 | 1492.69 | 746.85 | N | 1935.86 | 968.43 | 15 |
| 15 | 1579.72 | 790.37 | S | 1821.82 | 911.41 | 14 |
| 16 | 1735.82 | 868.42 | R | 1734.79 | 867.90 | 13 |
| 17 | 2008.94 | 1004.97 | W* | 1578.69 | 789.85 | 12 |
| 18 | 2195.02 | 1098.01 | W | 1305.58 | 653.29 | 11 |
| 19 | 2355.05 | 1178.03 | C | 1119.50 | 560.25 | 10 |
| 20 | 2469.09 | 1235.05 | N | 959.47 | 480.24 | 9 |
| 21 | 2584.12 | 1292.56 | D | 845.42 | 423.21 | 8 |
| 22 | 2641.14 | 1321.07 | G | 730.40 | 365.70 | 7 |
| 23 | 2797.24 | 1399.12 | R | 673.37 | 337.19 | 6 |
| 24 | 2898.29 | 1449.65 | T | 517.27 | 259.14 | 5 |
| 25 | 2995.34 | 1498.17 | P | 416.23 | 208.62 | 4 |
| 26 | 3052.36 | 1526.68 | G | 319.17 | 160.09 | 3 |
| 27 | 3139.39 | 1570.20 | S | 262.15 | 131.58 | 2 |
| 28 | | | R | 175.12 | 88.06 | 1 |

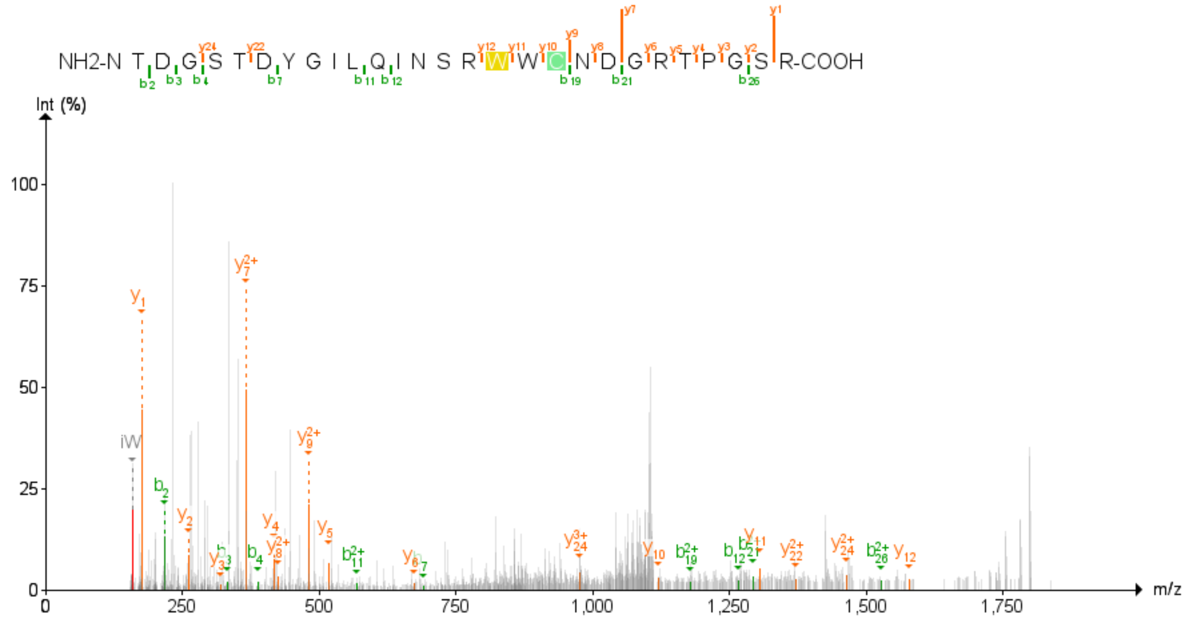


MS/MS data was obtained after analyzing the digested lysozyme **8**-conjugate sample according to the protocol outlined in nLC-MS/MS Method A

**Labeling of alpha-Chymotrypsinogen A(from bovine pancreas) with Probe 8:**
Labelling of alpha-Chymotrypsinogen with the Probe **8** was carried out as per the general labelling procedure under condition **A**, using the following solutions: alpha-Chymotrypsinogen (61 µL, 83 µM, 10 µM final concentration), **8** (36 µL, 2.8 mM, 200 µM final concentration), $NH_4OAc$, (50 µL, 200 mM, pH 6.9, 20 mM final concentration), GSH (15 µL, 10.0 mM, 300 µM final concentration), and deionized water (336 µL). The resulting solution sparged using Nitrogen for a minimum of 30 minutes and irradiated at 456 nm for 60 min. The reaction mixture was directly analyzed by **LC-MS Method A** and was estimated to have proceeded with 70% conversion and a 3.3:1 (+1:+2) labelled ratio.

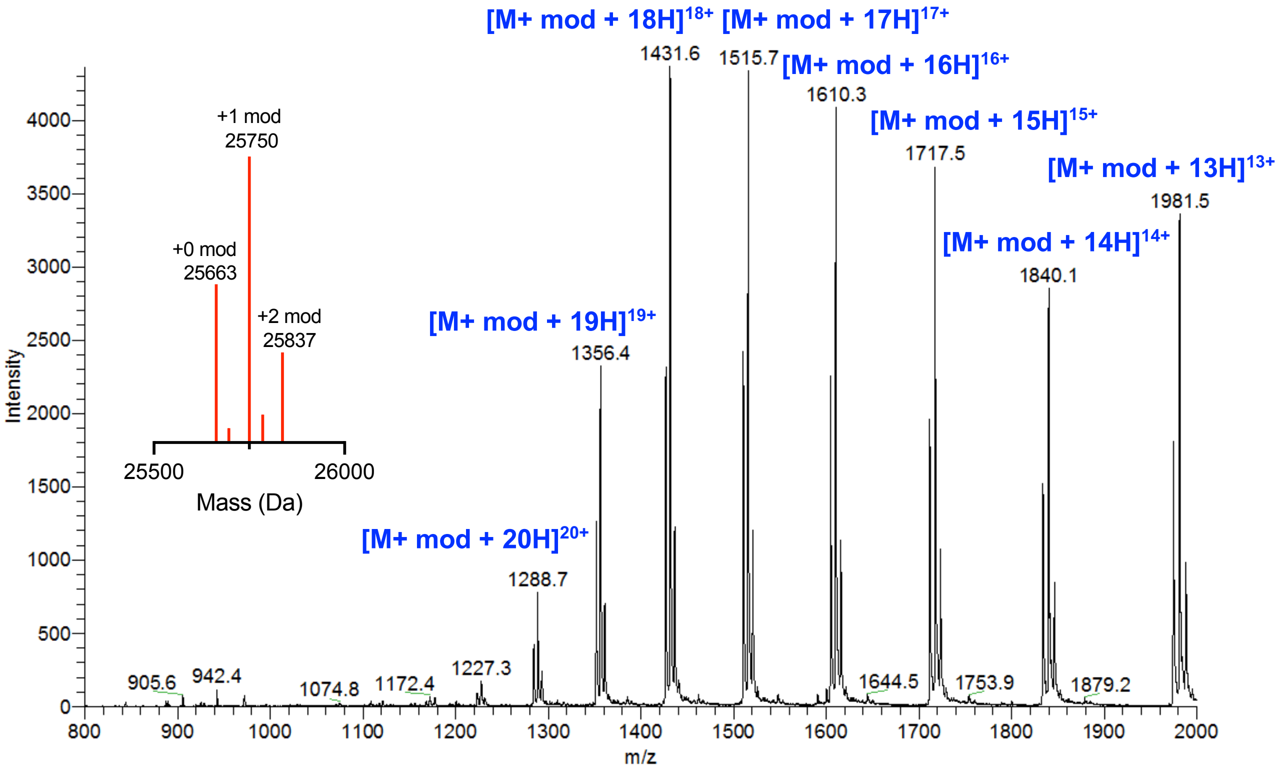


**Figure S2:** Mass spectrum of the N-methyl carbamate modified alpha-Chymotrypsinogen 8-conjugate ($T_r$ 4.4-6.8 min).

**MS-MS Analysis of alpha-Chymotrypsinogen A *N*-methycarbamate Modifications:**
nLC-MS/MS Method A analysis was performed on an alpha-Chymotrypsinogen A **8-**conjugate to know the site of modifications. Modifications were identified at **W237**.

**W237**:

| | b | b²⁺ | AA Sequence | y | y²⁺ | |
|---|---|---|---|---|---|---|
| 1 | 100.08 | 50.54 | V | | | 15 |
| 2 | 201.12 | 101.07 | T | 1615.84 | 808.43 | 14 |
| 3 | 272.16 | 136.58 | A | 1514.80 | 757.90 | 13 |
| 4 | 385.24 | 193.13 | L | 1443.76 | 722.38 | 12 |
| 5 | 484.31 | 242.66 | V | 1330.67 | 665.84 | 11 |
| 6 | 598.36 | 299.68 | N | 1231.61 | 616.31 | 10 |
| 7 | 871.47 | 436.24 | W* | 1117.56 | 559.29 | 9 |
| 8 | 970.54 | 485.77 | V | 844.45 | 422.73 | 8 |
| 9 | 1098.59 | 549.80 | Q | 745.38 | 373.20 | 7 |
| 10 | 1226.65 | 613.83 | Q | 617.33 | 309.17 | 6 |
| 11 | 1327.70 | 664.35 | T | 489.27 | 245.14 | 5 |
| 12 | 1440.78 | 720.90 | L | 388.22 | 194.61 | 4 |
| 13 | 1511.82 | 756.41 | A | 275.13 | 138.07 | 3 |
| 14 | 1582.86 | 791.93 | A | 204.10 | 102.55 | 2 |
| 15 | | | N | 133.06 | 67.03 | 1 |

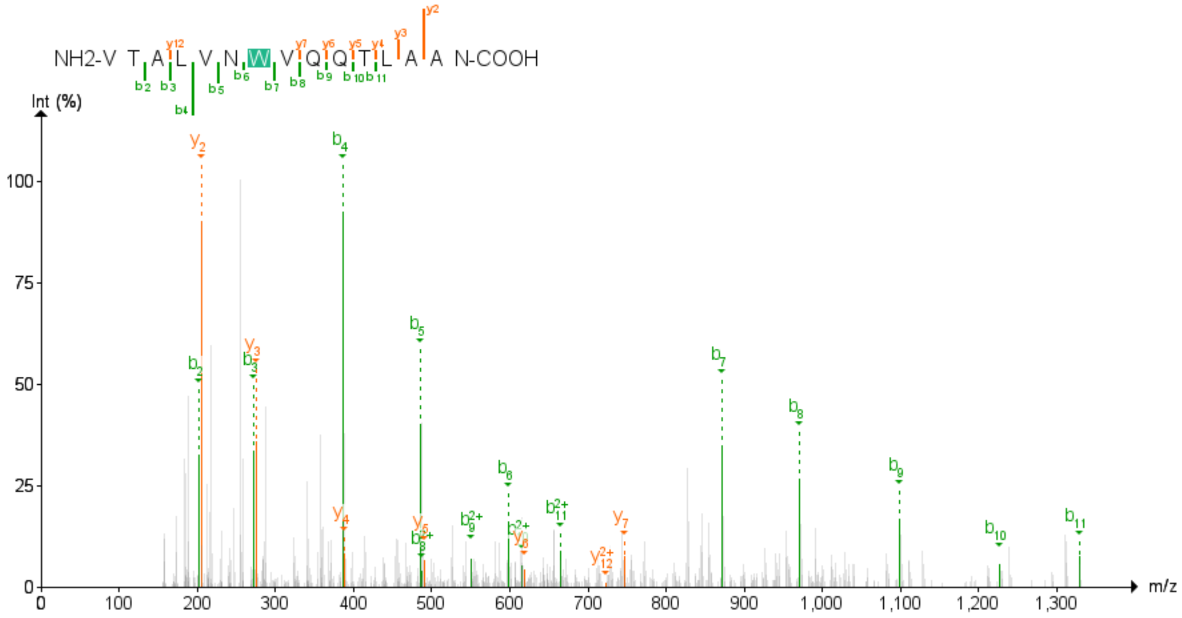


MS/MS data was obtained after analyzing the digested alpha-Chymotrypsinogen A conjugate sample according to the protocol outlined in nLC-MS/MS Method A.

### 4. Photophysical Properties of pyridinium and pyrimidinium probes

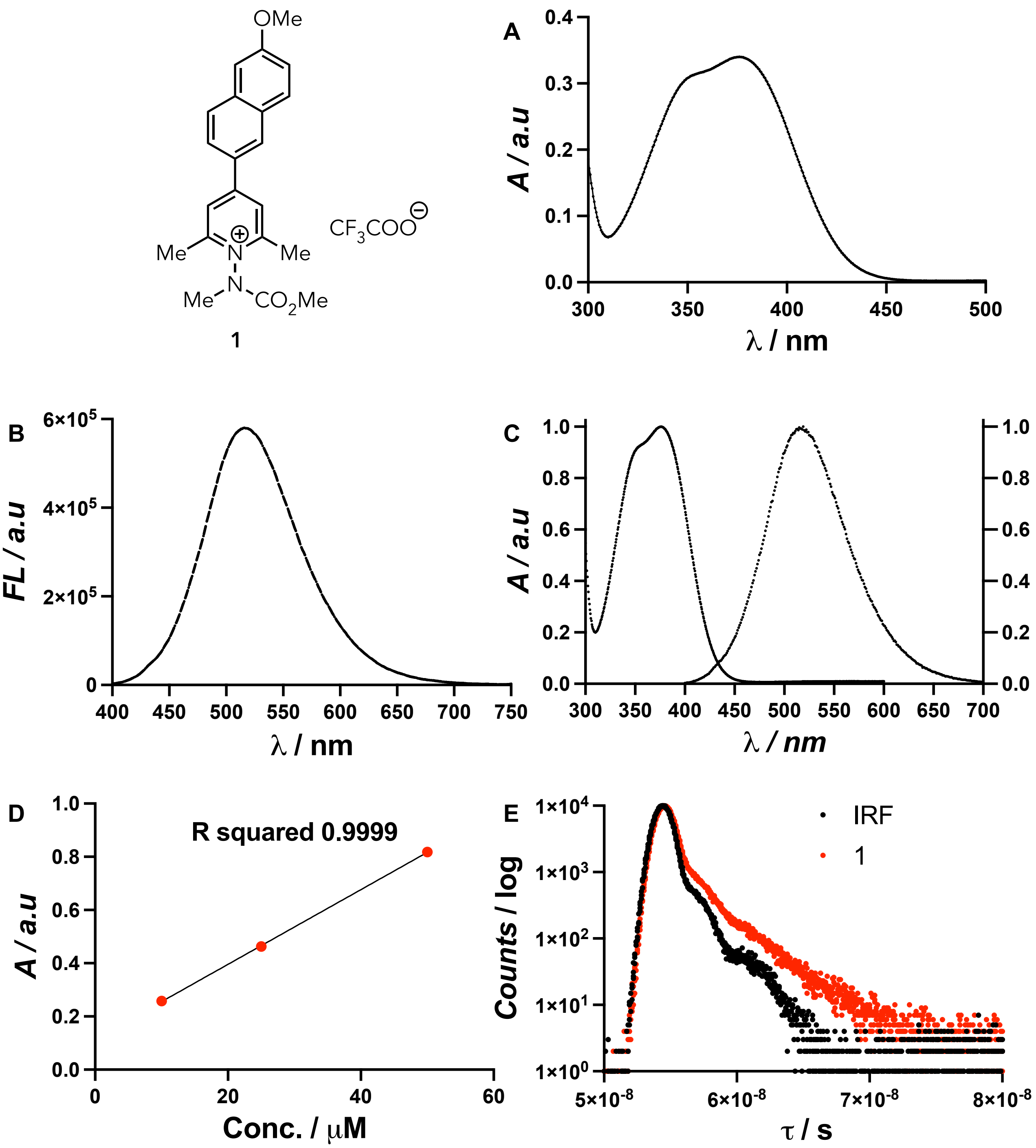


**Figure S3:** (A) UV-Vis absorption plot of **1** (20 μM) in phosphate buffer pH 7.4, (B) Fluorescence spectra of **1** (20 μM) in phosphate buffer pH 7.4 with excitation and emission slit width 2 nm, (C) Normalized absorption and emission spectra of **1** in phosphate buffer pH 7.4. (D) Absorbance vs Concentration (10, 25, and 50 μM) plot of **1** (in phosphate buffer pH 7.4) to measure the molar extinction coefficient ($\varepsilon = 14.0\times 10^3\ M^{-1}cm^{-1}$), (E) Fluorescence lifetimes of **1** in water ($\tau_1 = 0.23$ ns and $\tau_2 = 2.89$ ns, $\chi^2 = 1.19$) was measured using NanoLED-370 (peak wavelength 367 nm, bandpass = 5 nm), IRF = Instrument Response Function.

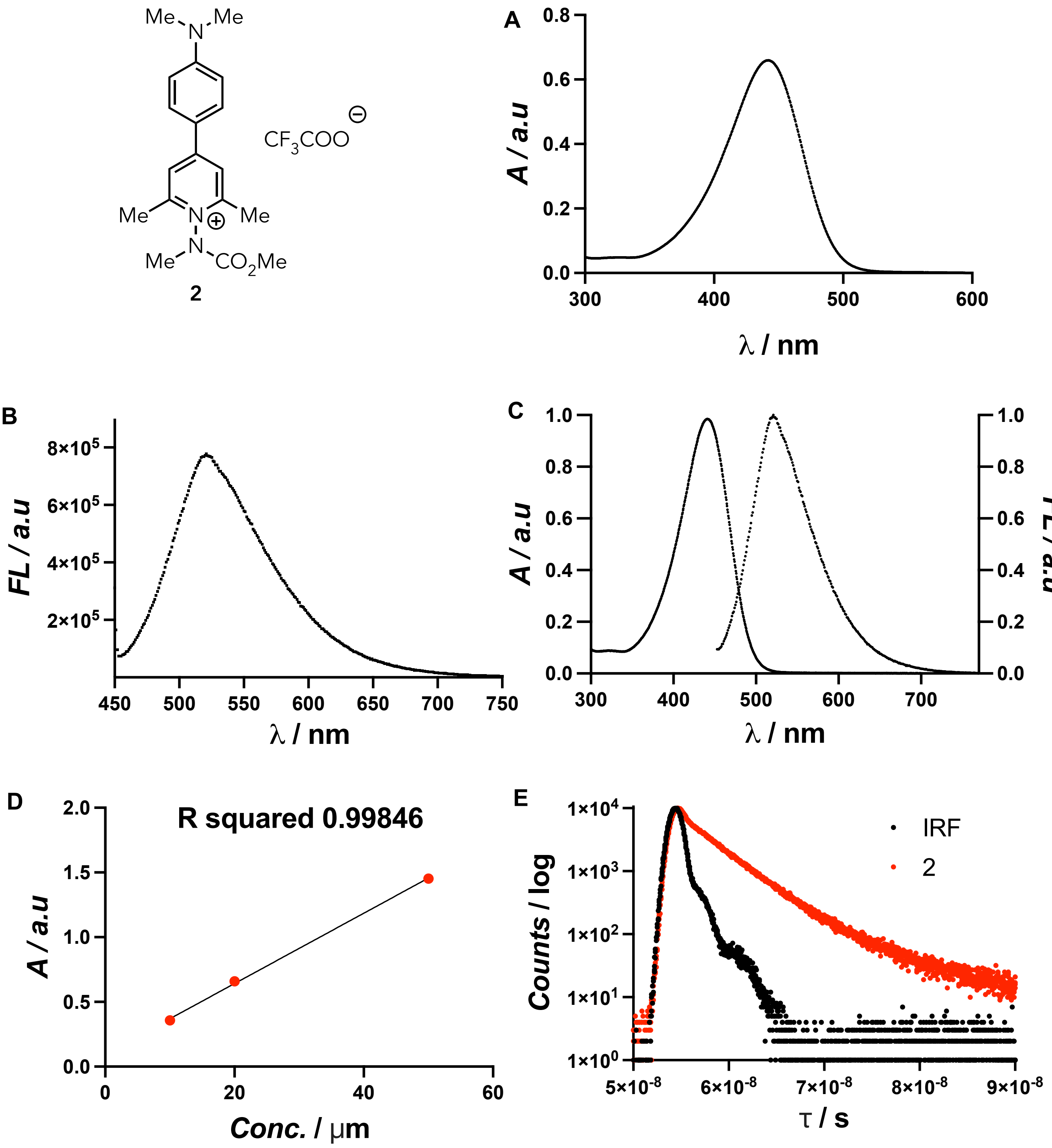


**Figure S4:** (A) UV-Vis absorption plot of **2** (20 µM) in phosphate buffer pH 7.4, (B) Fluorescence spectra of **2** (20 µM) in phosphate buffer pH 7.4 with excitation and emission slit width 2 nm, (C) Normalized absorption and emission spectra of **2** in phosphate buffer pH 7.4. (D) Absorbance vs Concentration (10, 20, and 50 µM) plot of **2** (in phosphate buffer pH 7.4) to measure the molar extinction coefficient ($\varepsilon = 27.1\times 10^{3}\ M^{-1}cm^{-1}$), (E) Fluorescence lifetimes of **2** in water ($\tau_1 = 3.12$ ns and $\tau_2 = 6.66$ ns, $\chi^2 = 1.07$) was measured using NanoLED-370 (peak wavelength 367 nm, bandpass = 5 nm), IRF = Instrument Response Function.

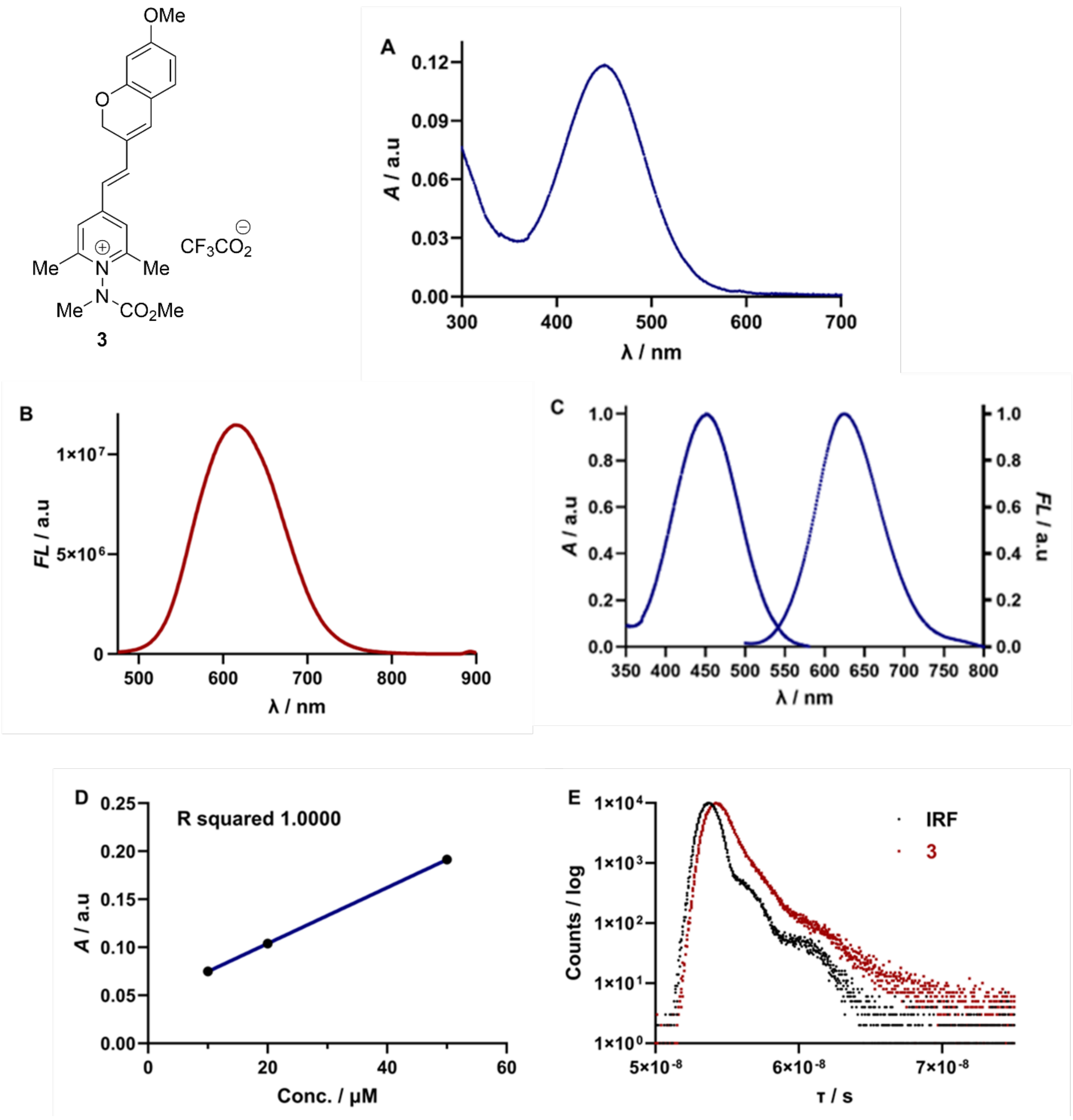


**Figure S5:** (A) UV-Vis absorption plot of **3** (20 μM) in phosphate buffer pH 7.4, (B) Fluorescence spectra of **3** (20 μM) in phosphate buffer pH 7.4 with excitation and emission slit width 5 nm, (C) Normalized absorption and emission spectra of **3** in phosphate buffer pH 7.4. (D) Absorbance vs Concentration (10, 20, and 50 μM) plot of **3** (in phosphate buffer pH 7.4) to measure the molar extinction coefficient ($\varepsilon = 10.6\times 10^3 M^{-1}cm^{-1}$), (E) Fluorescence lifetimes of **3** in water ($\tau_1 = 0.53$ ns and $\tau_2 = 2.74$ ns, $\chi^2 = 1.08$) was measured using NanoLED-370 (peak wavelength 367 nm, bandpass = 5 nm), IRF = Instrument Response Function.

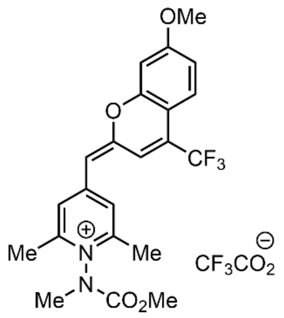


**4**

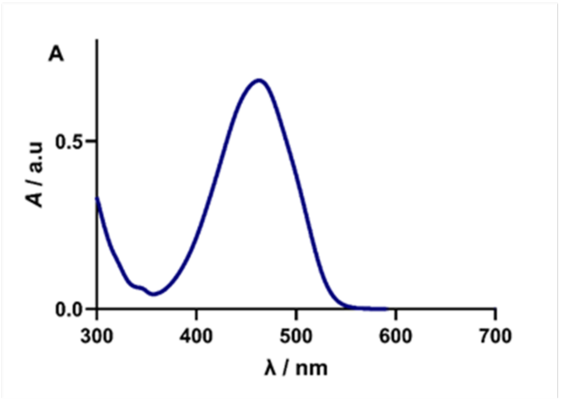


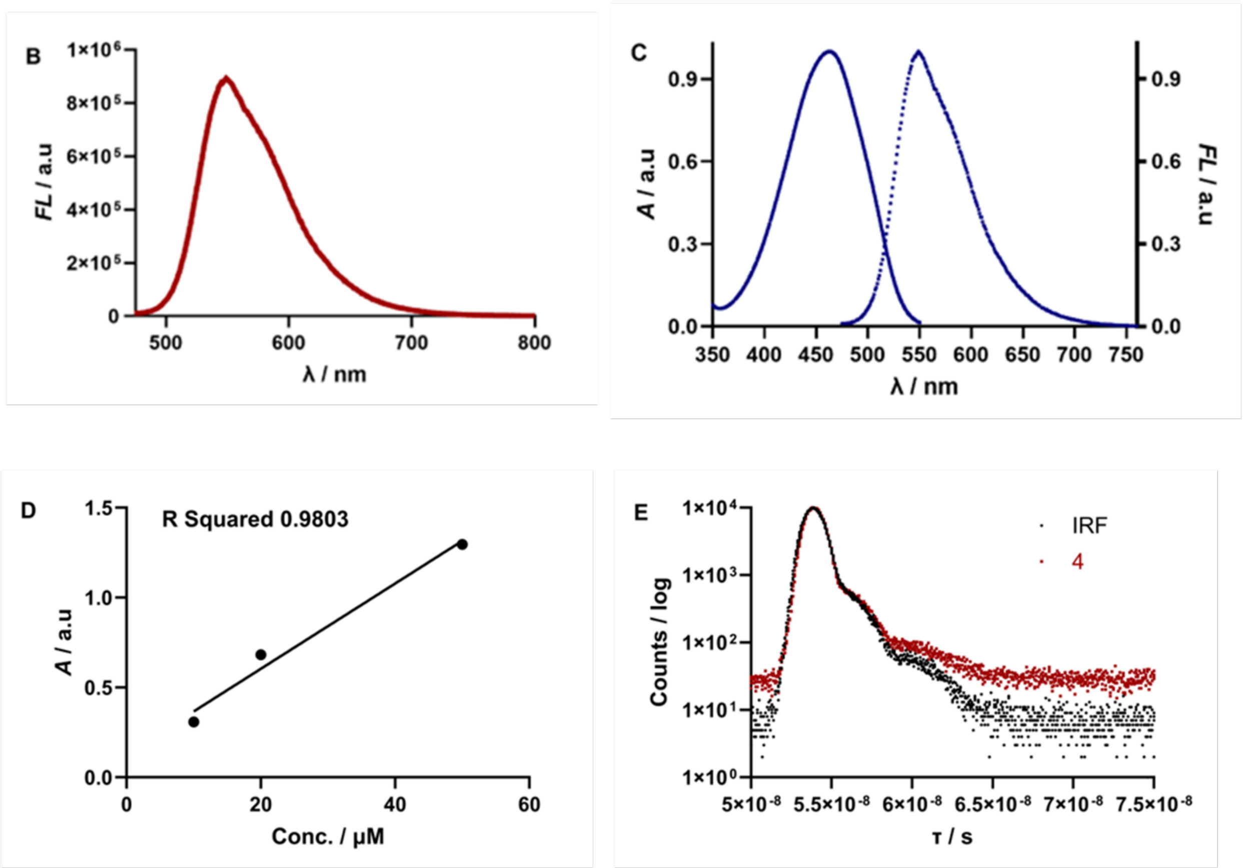


**Figure S6:** (A) UV-Vis absorption plot of **4** (20 μM) in phosphate buffer pH 7.4, (B) Fluorescence spectra of **4** (20 μM) in phosphate buffer pH 7.4 with excitation and emission slit width 2 nm, (C) Normalized absorption and emission spectra of **4** in phosphate buffer pH 7.4. (D) Absorbance vs Concentration (10, 20, and 50 μM) plot of **4** (in phosphate buffer pH 7.4) to measure the molar extinction coefficient ($\varepsilon = 23.7\times 10^{3}\,M^{-1}cm^{-1}$), (E) Fluorescence lifetimes of **4** in water ($\tau_1$ = 0.20 $\tau_2$ = 0.19 ns $\chi^2$ = 1.18) was measured using NanoLED-455 (peak wavelength 450 nm, bandpass = 5 nm), IRF = Instrument Response Function.

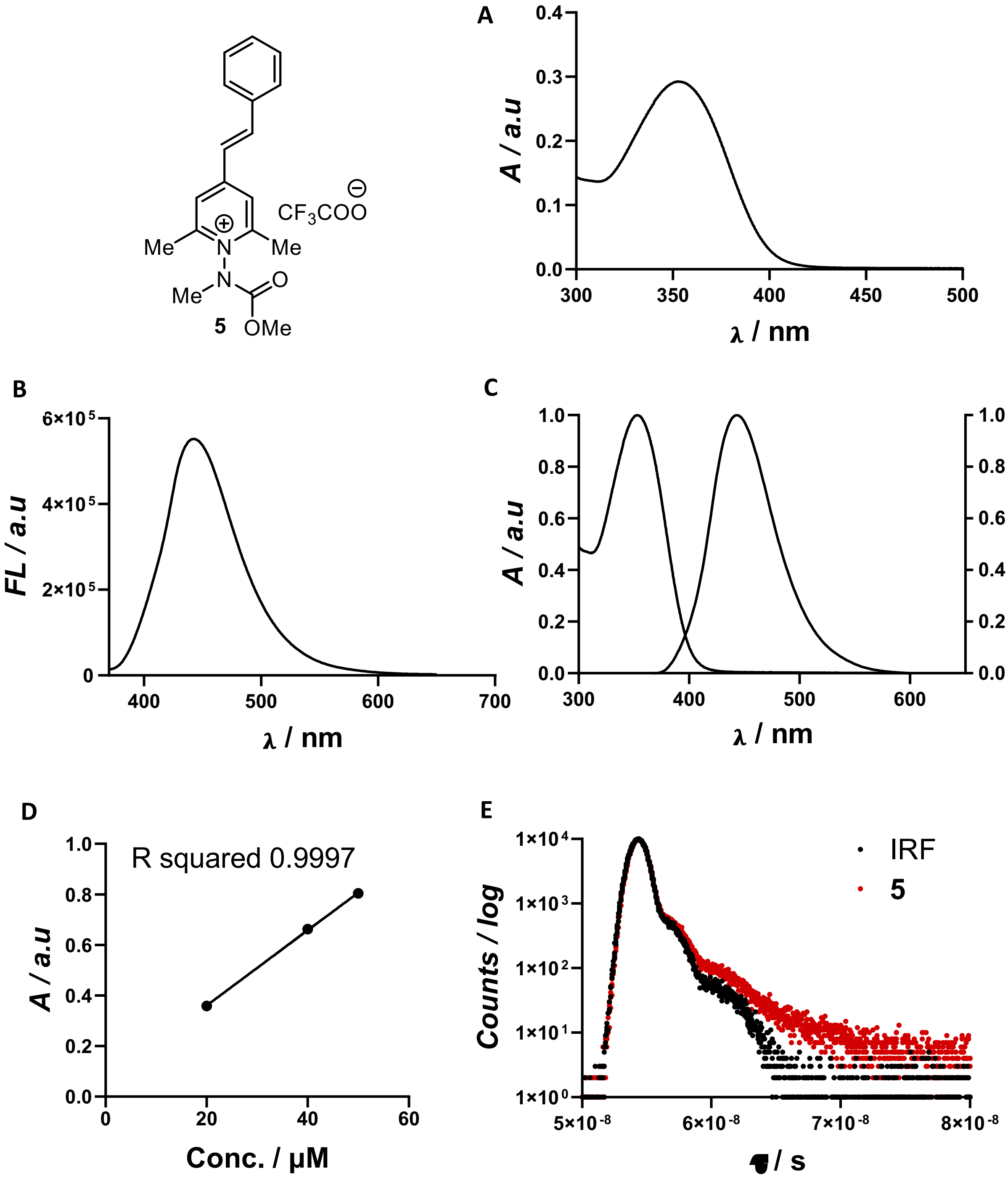


**Figure S7:** (A) UV-Vis absorption plot of **5** (20 μM) in phosphate buffer pH 7.5, (B) Fluorescence spectra of **5** (20 μM) in phosphate buff er pH 7.5 with excitation and emission slit width 5 nm, (C) Normalized absorption and emission spectra of **5** in phosphate buffer pH 7.5, (D) Absorbance vs Concentration (20, 40, and 50 μM) plot of **5** (in phosphate buffer pH 7.5) to measure the molar extinction coefficient ($\varepsilon = 14.8\times 10^3\ M^{-1}cm^{-1}$), (E) Fluorescence lifetimes of **5** in water ($\tau_1 = 4.62$ and $\tau_2 = 0.08$ ns, $\chi^2 = 1.20$) was measured using NanoLED-370 (peak wavelength 370 nm, bandpass = 8 nm), IRF = Instrument Response Function.

.

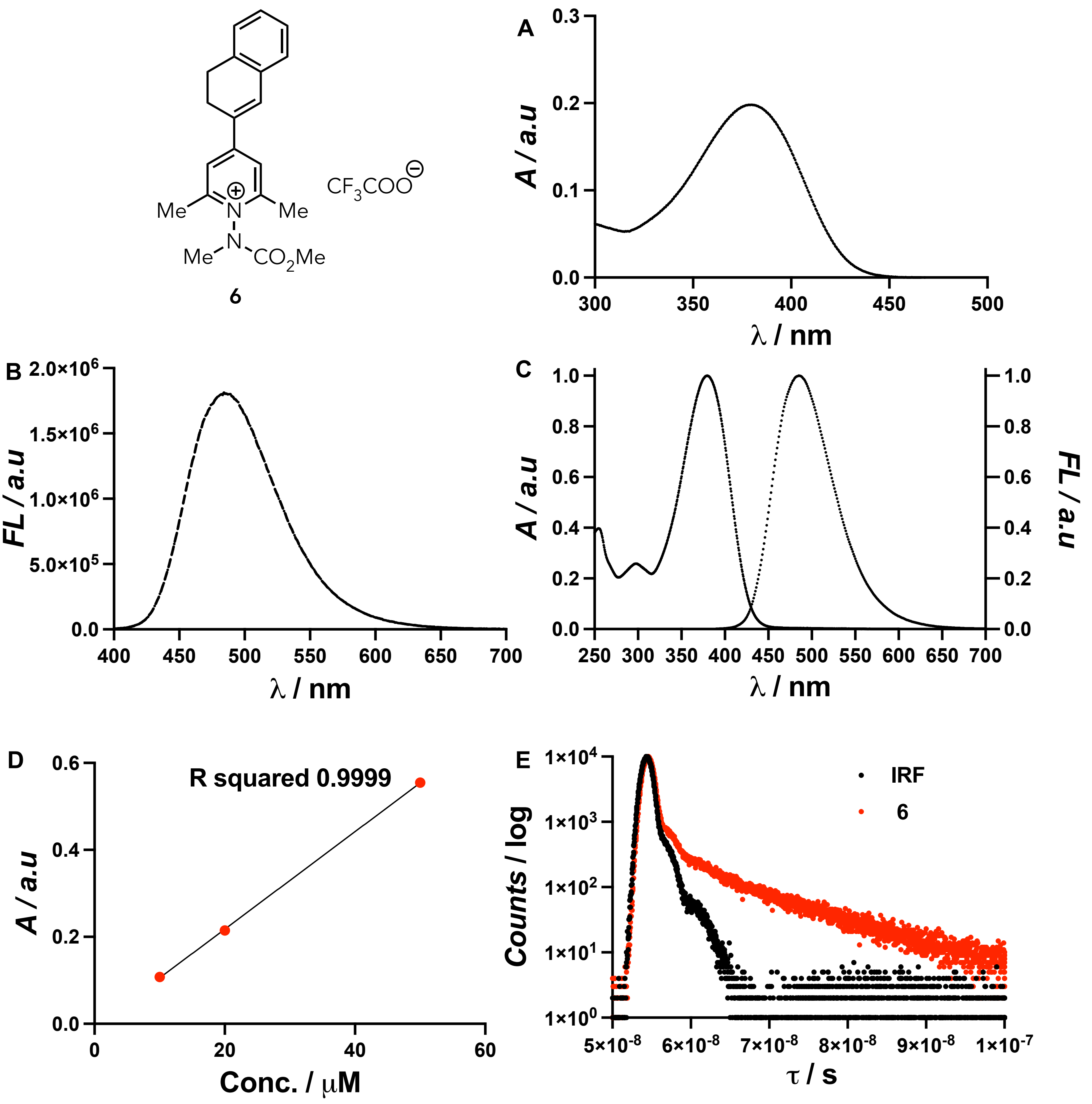


**Figure S8:** (A) UV-Vis absorption plot of **6** (20 μM) in phosphate buffer pH 7.4, (B) Fluorescence spectra of **6** (20 μM) in phosphate buffer pH 7.4 with excitation and emission slit width 3 nm, (C) Normalized absorption and emission spectra of **6** in phosphate buffer pH 7.4. (D) Absorbance vs Concentration (10, 20, and 50 μM) plot of **6** (in phosphate buffer pH 7.4) to measure the molar extinction coefficient ($\varepsilon = 11.2\times 10^3\ M^{-1}cm^{-1}$), (E) Fluorescence lifetimes of **6** in water ($\tau_1 = 0.10$ ns and $\tau_2 = 9.76$ ns, $\chi^2 = 1.07$) was measured using NanoLED-370 (peak wavelength 367 nm, bandpass = 5 nm), IRF = Instrument Response Function.

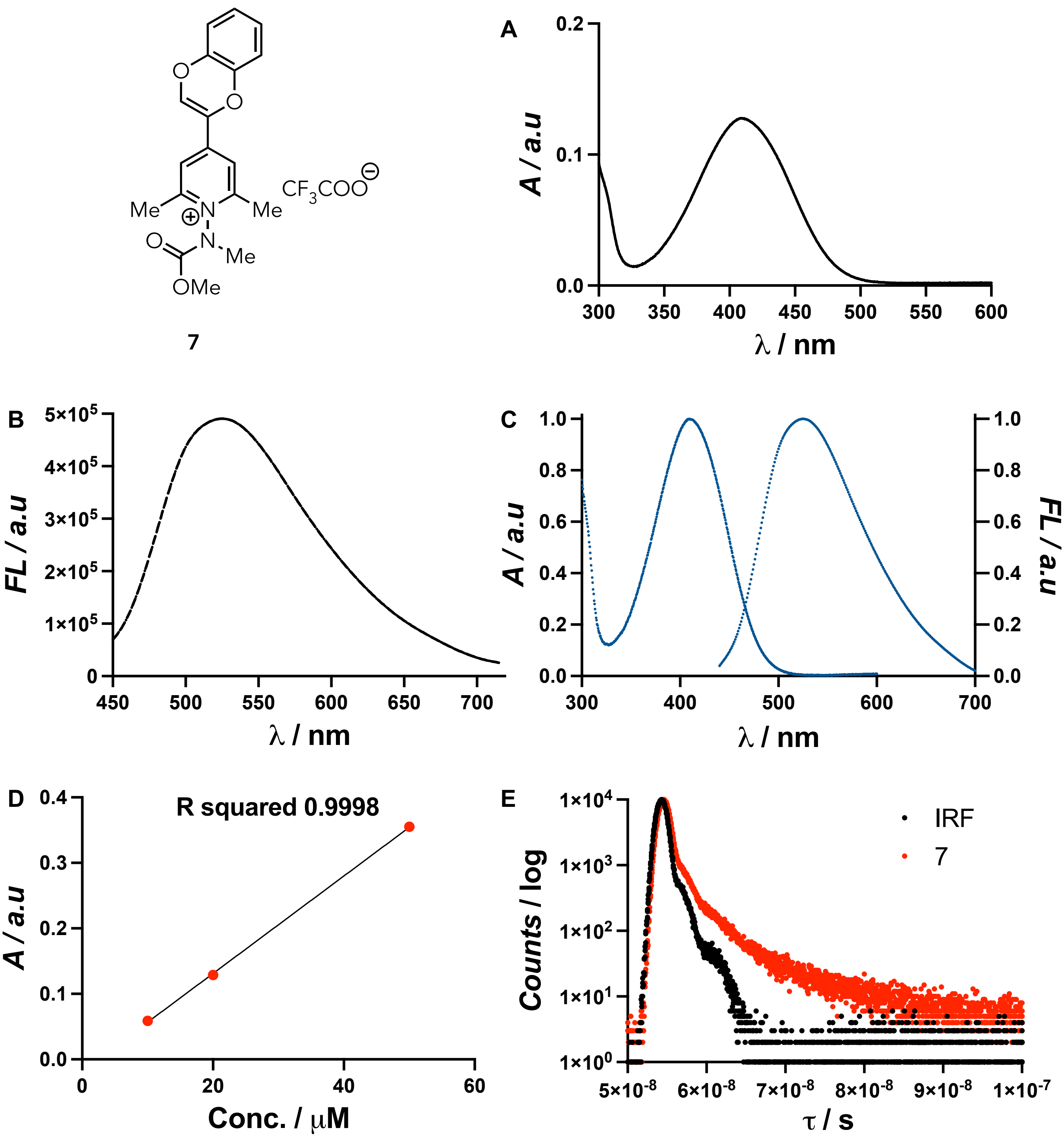


**Figure S9:** (A) UV-Vis absorption plot of **7** (20 μM) in phosphate buffer pH 7.4, (B) Fluorescence spectra of **7** (20 μM) in phosphate buffer pH 7.4 with excitation and emission slit width 2 nm, (C) Normalized absorption and emission spectra of **7** in phosphate buffer pH 7.4. (D) Absorbance vs Concentration (10, 20, and 50 μM) plot of **7** (in phosphate buffer pH 7.4) to measure the molar extinction coefficient ($\varepsilon = 7.28 \times 10^3$ $M^{-1}cm^{-1}$), (E) Fluorescence lifetimes of **7** in water ($\tau_1 = 0.28$ ns and $\tau_2 = 4.58$ ns, $\chi^2 = 1.32$) was measured using NanoLED-370 (peak wavelength 367 nm, bandpass = 5 nm), IRF = Instrument Response Function.

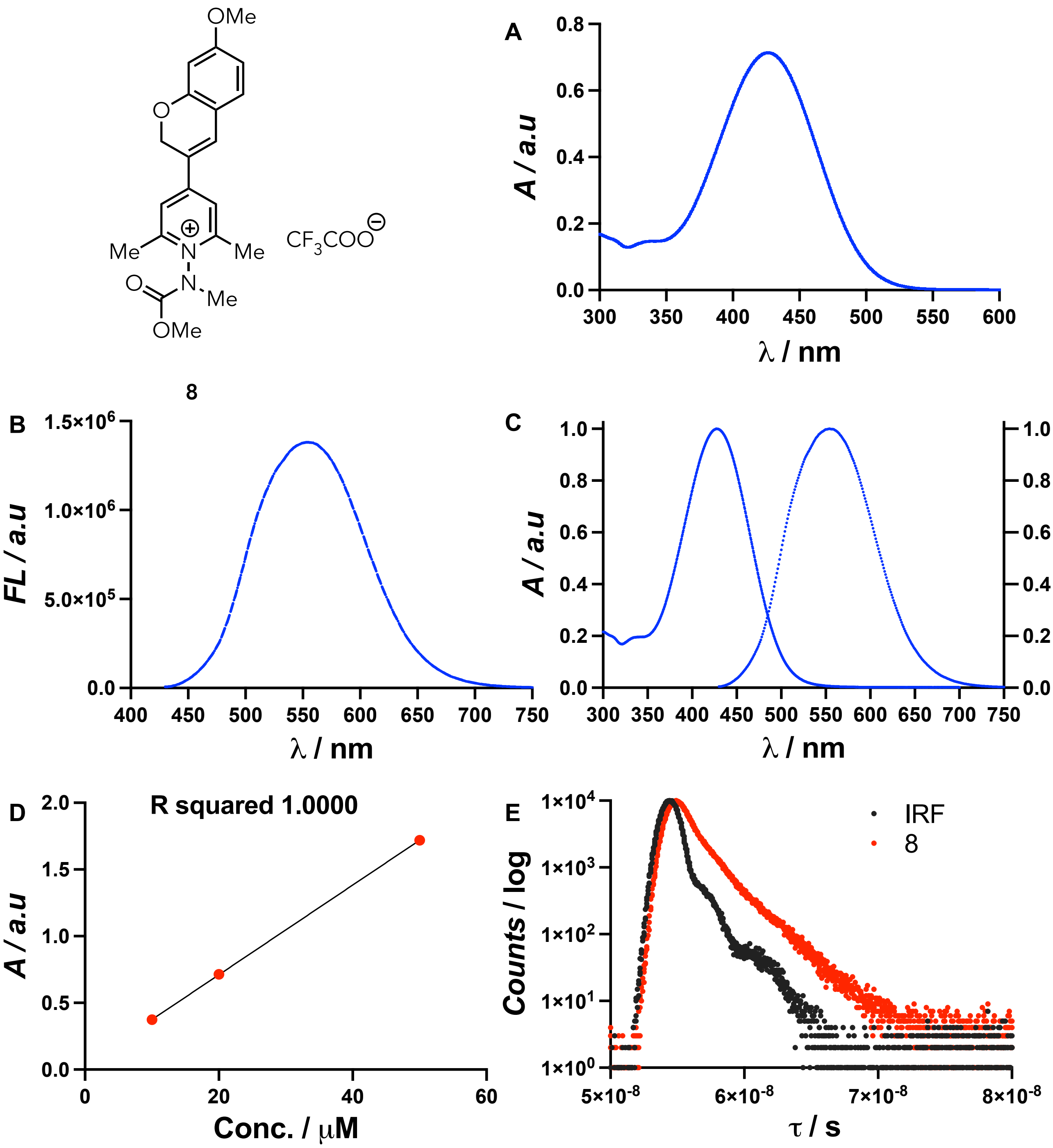


**Figure S10:** (A) UV-Vis absorption plot of **8** (20 μM) in phosphate buffer pH 7.4, (B) Fluorescence spectra of **8** (20 μM) in phosphate buffer pH 7.4 with excitation and emission slit width 2 nm, (C) Normalized absorption and emission spectra of **8** in phosphate buffer pH 7.4. (D) Absorbance vs Concentration (10, 20, and 50 μM) plot of **8** (in phosphate buffer pH 7.4) to measure the molar extinction coefficient ($\varepsilon = 33.6 \times 10^3\ M^{-1}cm^{-1}$), (E) Fluorescence lifetimes of **8** in water ($\tau_1 = 0.64$ ns and $\tau_2 = 2.06$ ns, $\chi^2 = 1.26$) was measured using NanoLED-370 (peak wavelength 367 nm, bandpass = 5 nm), IRF = Instrument Response Function.

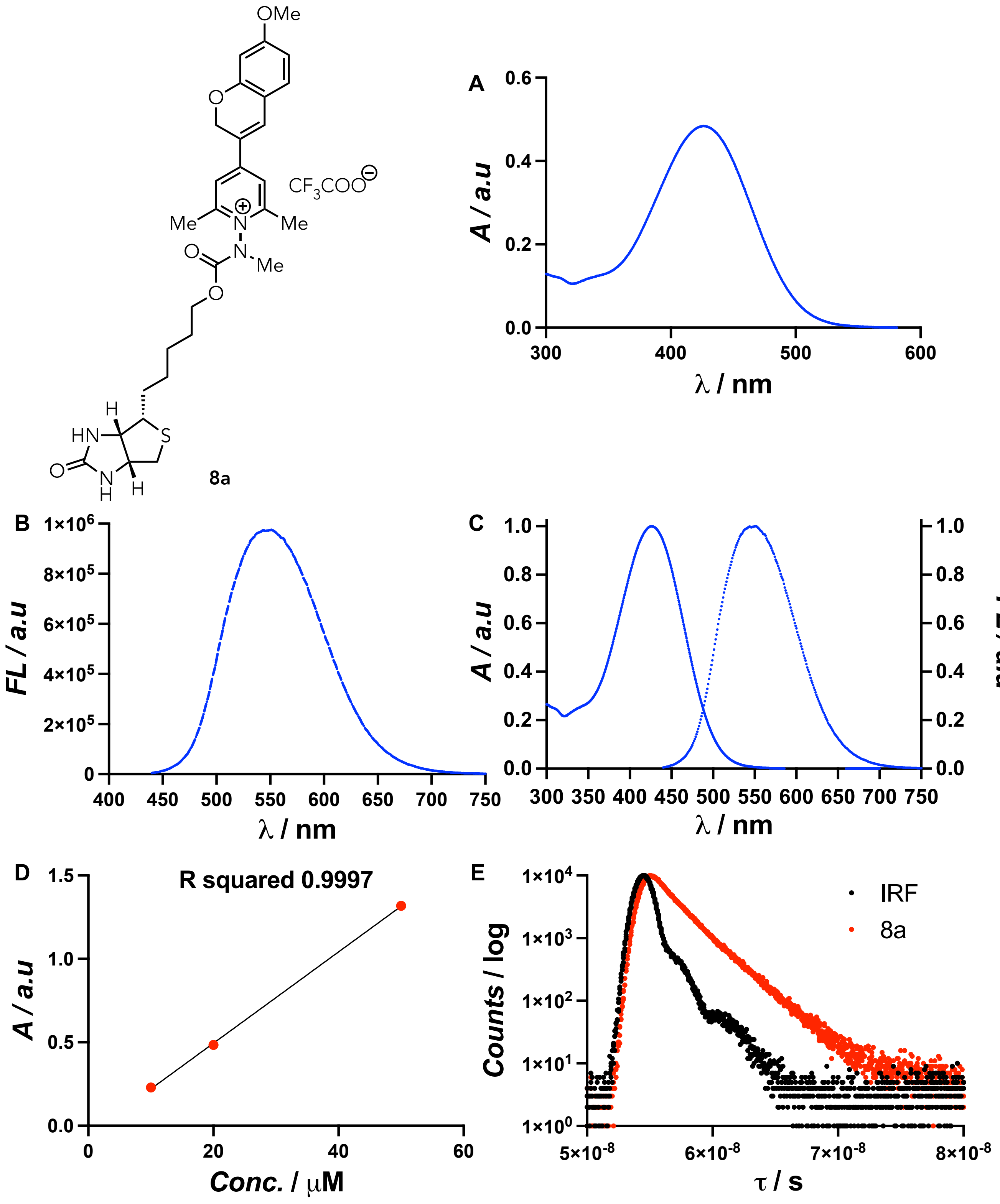


**Figure S11:** (A) UV-Vis absorption plot of **8a** (20 μM) in phosphate buffer pH 7.4, (B) Fluorescence spectra of **8a** (20 μM) in phosphate buffer pH 7.4 with excitation and emission slit width 1.5 nm, (C) Normalized absorption and emission spectra of **8a** in phosphate buffer pH 7.4. (D) Absorbance vs Concentration (10, 20, and 50 μM) plot of **8a** (in phosphate buffer pH 7.4) to measure the molar extinction coefficient ($\varepsilon$ = 27.5 × $10^3$ $M^{-1}cm^{-1}$), (E) Fluorescence lifetimes of **8a** in water ($\tau_1$ = 1.11 ns and $\tau_2$ = 2.55 ns, $\chi^2$ = 1.19) was measured using NanoLED-370 (peak wavelength 367 nm, bandpass = 5 nm), IRF = Instrument Response Function.

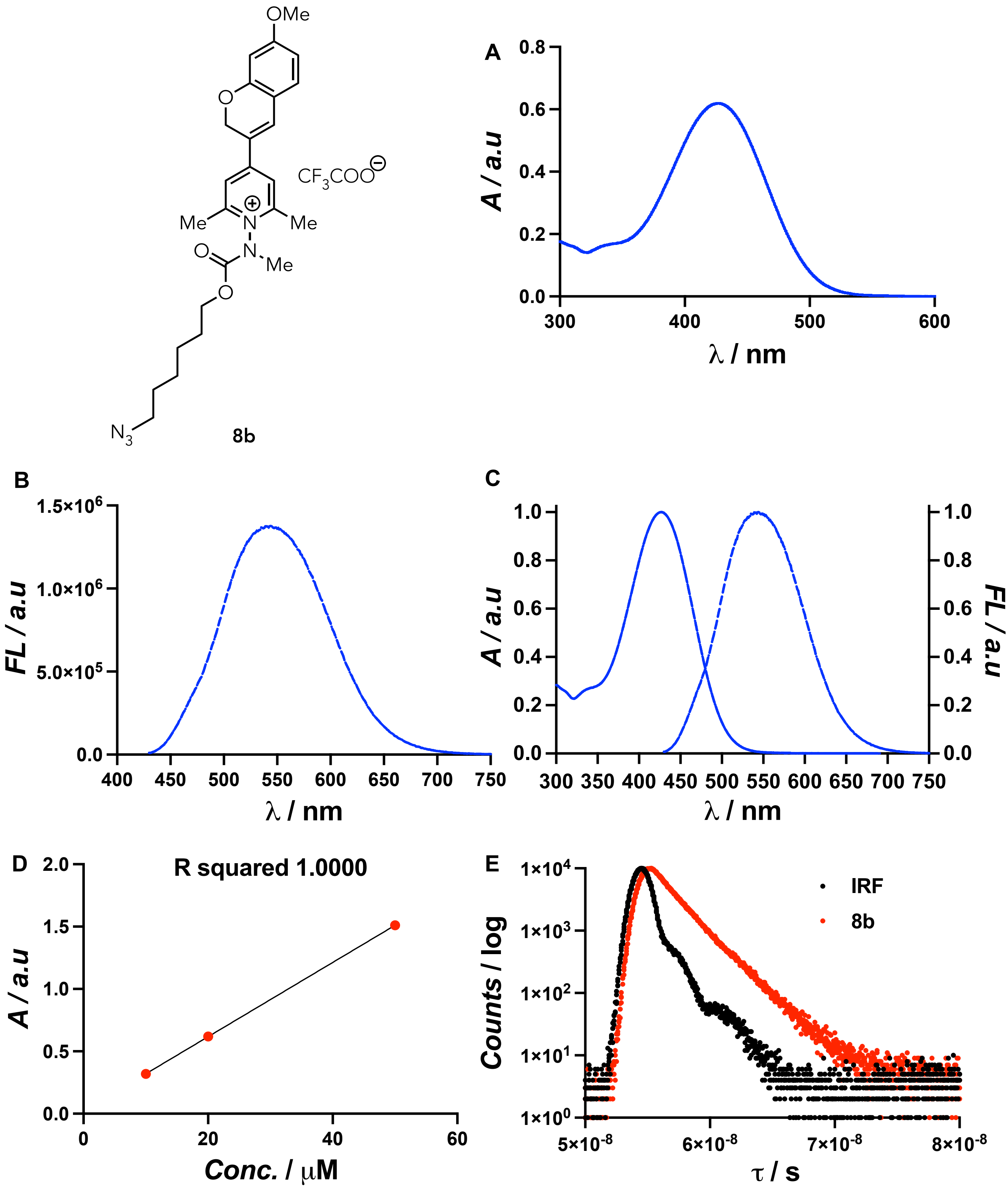


**Figure S12:** (A) UV-Vis absorption plot of **8b** (20 μM) in phosphate buffer pH 7.4, (B) Fluorescence spectra of **8b** (20 μM) in phosphate buffer pH 7.4 with excitation and emission slit width 2 nm, (C) Normalized absorption and emission spectra of **8b** in phosphate buffer pH 7.4. (D) Absorbance vs Concentration (10, 20, and 50 μM) plot of **8b** (in phosphate buffer pH 7.4) to measure the molar extinction coefficient ($\varepsilon = 29.8 \times 10^3$ $M^{-1}cm^{-1}$), (E) Fluorescence lifetimes of **8b** in water ($\tau_1 = 1.01$ ns, $\chi^2 = 1.00$) was measured using NanoLED-370 (peak wavelength 367 nm, bandpass = 5 nm), IRF = Instrument Response Function.

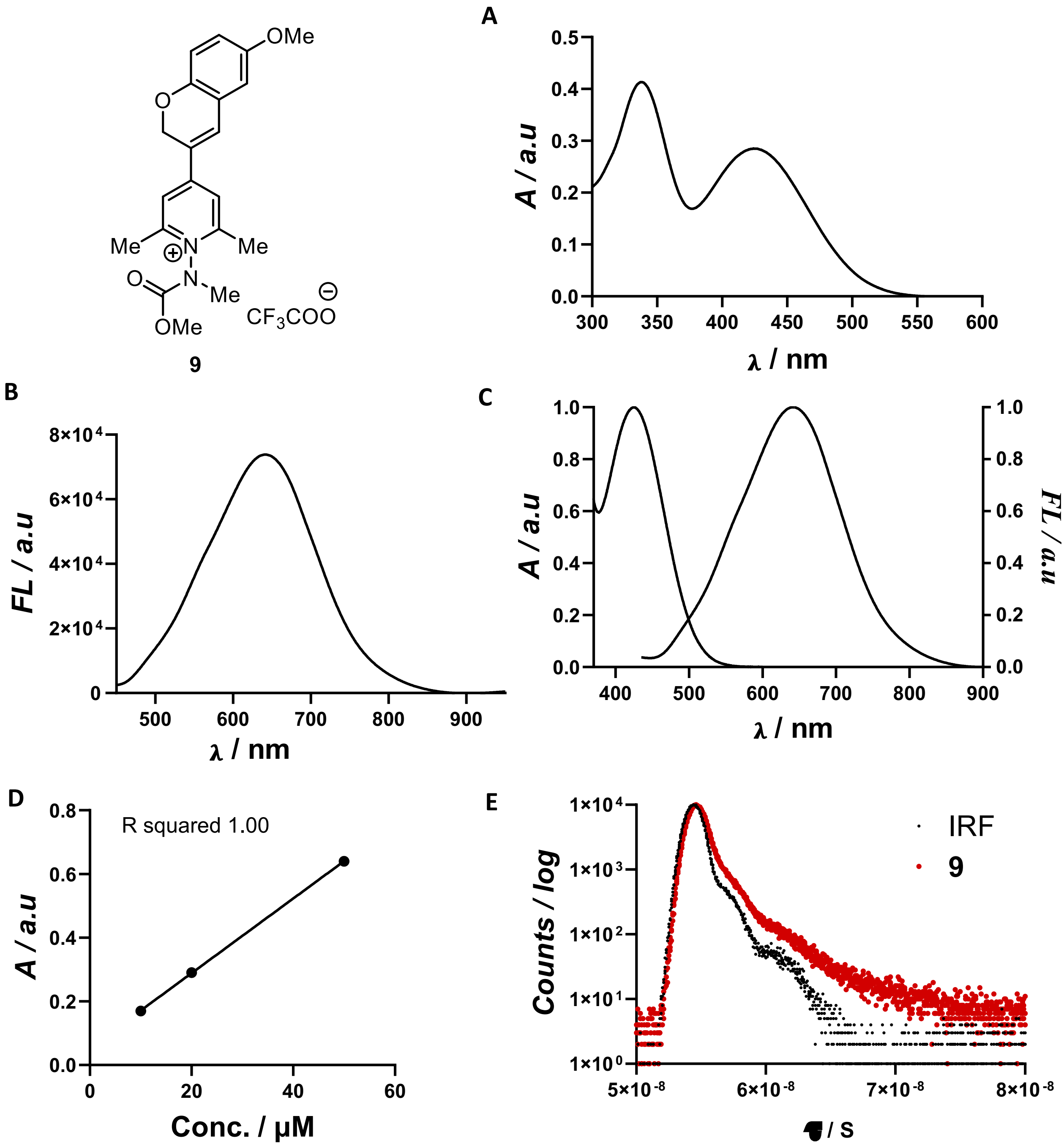


**Figure S13:** (A) UV-Vis absorption plot of **9** (20 μM) in phosphate buffer pH 7.5, (B) Fluorescence spectra of **9** (20 μM) in phosphate buffer pH 7.5 with excitation and emission slit width 4 nm, (C) Normalized absorption and emission spectra of **9** in phosphate buffer pH 7.5, (D) Absorbance vs Concentration (10, 20, and 50 μM) plot of **9** (in phosphate buffer pH 7.5) to measure the molar extinction coefficient ($\varepsilon$ = 11.7 × $10^3$ $M^{-1}cm^{-1}$), (E) Fluorescence lifetimes of **9** in water ($\tau_1$ = 0.32 and $\tau_2$ =2.41 ns, $\chi^2$ =1.29 ) was measured using NanoLED-370 (peak wavelength 367 nm, bandpass 8 nm), IRF= Instrument Response Function

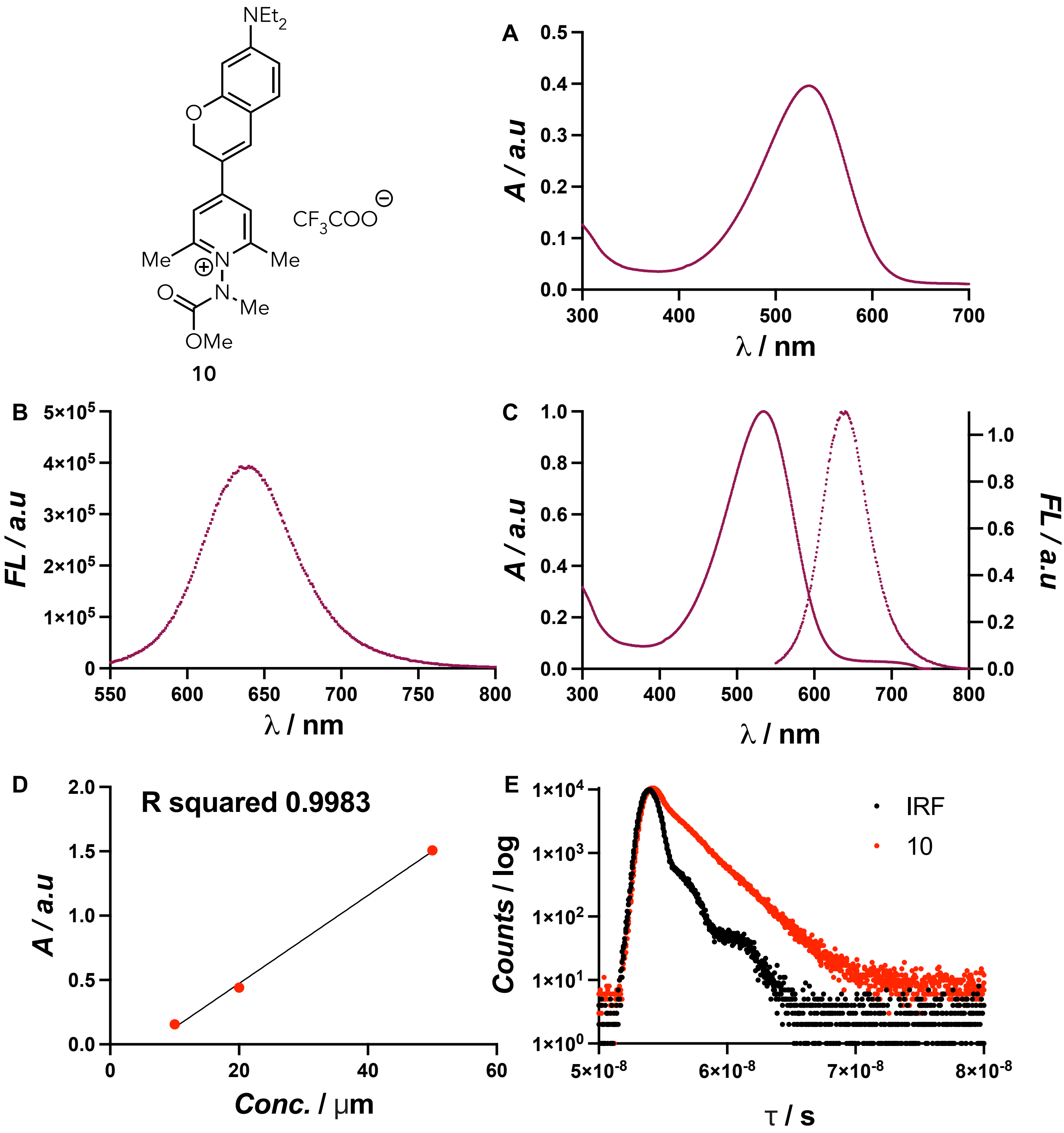


**Figure S14:** (A) UV-Vis absorption plot of **10** (20 μM) in phosphate buffer pH 7.4, (B) Fluorescence spectra of **10** (20 μM) in phosphate buffer pH 7.4 with excitation and emission slit width 3 nm, (C) Normalized absorption and emission spectra of **10** in phosphate buffer pH 7.4. (D) Absorbance vs Concentration (10, 20, and 50 μM) plot of **10** (in phosphate buffer pH 7.4) to measure the molar extinction coefficient ($\varepsilon$ = 34.2× $10^3$ $M^{-1}cm^{-1}$), (E) Fluorescence lifetimes of **10** in water ($\tau_1$ = 0.12 ns and $\tau_2$ = 1.95 ns, $\chi^2$ = 0.93) was measured using NanoLED-455 (peak wavelength 450 nm, bandpass = 5 nm), IRF = Instrument Response Function.

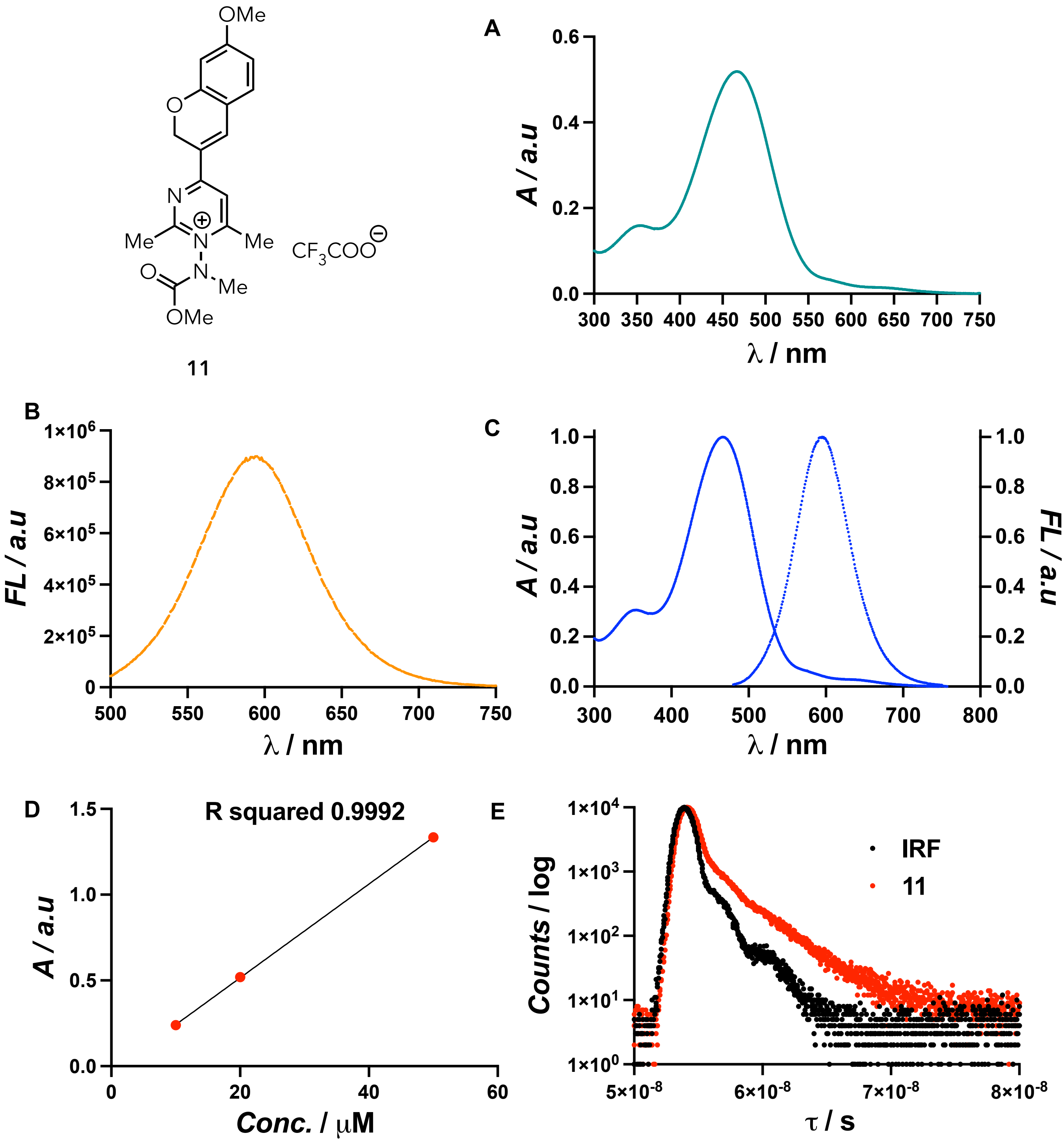


**Figure S15:** (A) UV-Vis absorption plot of **11** (20 μM) in phosphate buffer pH 3.5, (B) Fluorescence spectra of **11** (20 μM) in phosphate buffer pH 3.5 with excitation and emission slit width 2 nm, (C) Normalized absorption and emission spectra of **11** in phosphate buffer pH 3.5, (D) Absorbance vs Concentration (10, 20, and 50 μM) plot of **11** (in phosphate buffer pH 3.5) to measure the molar extinction coefficient ($\varepsilon$ = 27.3 × $10^3$ $M^{-1}cm^{-1}$), (E) Fluorescence lifetimes of **11** in phosphate buffer pH 3.5 ($\tau_1$ = 0.26 and $\tau_2$ = 2.94 ns, $\chi^2$ = 1.06) was measured using NanoLED-455 (peak wavelength 450 nm, bandpass = 5 nm), IRF = Instrument Response Function.

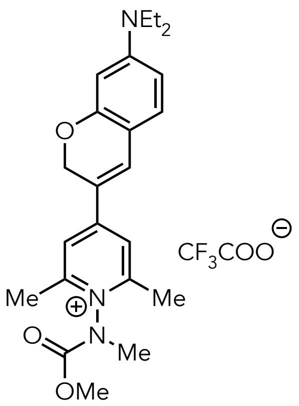

**10**

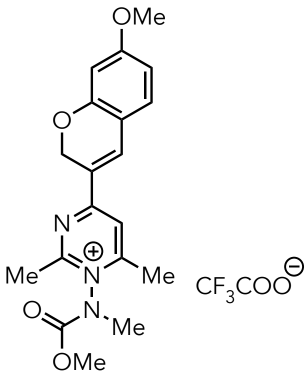

**11**

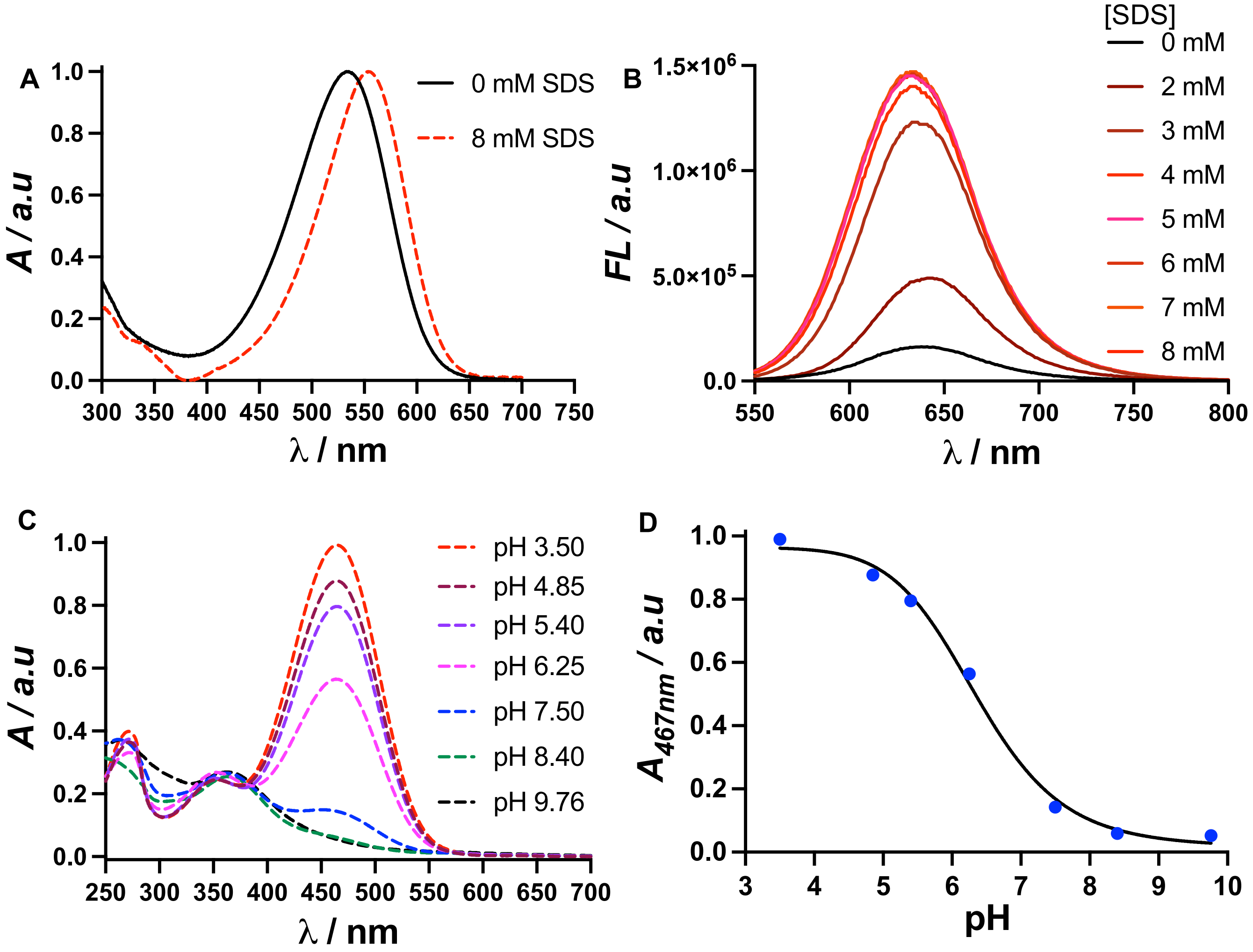


**Figure S16:** (A) A bathochromic shift in the UV-Vis absorption and (B) Turn-on effect in fluorescence intensity of **10** (20 μM) was observed in phosphate buffer (pH 7.4) with varying concentrations (0-8 mM) of sodium dodecyl sulfate (SDS). (C) A pH-dependent UV-Vis absorption plot of **11** in 20 mM phosphate buffer. All absorption data were collected after a 30-minute equilibration period. (D) The sigmoidal plot illustrates the pH-dependent changes in the absorption of **11** at 467 nm in phosphate buffer. The pKa of **11** was determined to be 6.4.

**5. Live cell imaging with pyridinium and pyrimidinium probes using confocal laser scanning microscopy:**

Confocal laser scanning microscopic (CLSM) images were captured on a Zeiss LSM880, AxioObserver, Plan-Apochromat 63x/1.40 OIL DIC M27 objective using 405, 488, 561, and 633 nm lasers. For live cell CLSM imaging HeLa cells (cell viability > 95%) at a density of ~$1\times10^5$ cells/mL were separately seeded in 35 mm glass bottom confocal dish (MATTEK catalogue no. P35G-1.5-14-C) and allowed to get 70-80% confluency in DMEM for 24 h at 37 °C in an air-jacketed 5% $CO_2$ incubator. Afterward washing with 1× DPBS(2x's) live cells were incubated with incubated first with 200 nM MitoTracker Red FM (Invitrogen, Cat. no. M22425) for 30 min at 37°C in 5% $CO_2$ environment and then the media was removed carefully and washed gently with 1× DPBS (3x's). Subsequently, the cells were incubated with 10 µM Probe **8** (stock concentration 1 mM in water), at 37°C in 5% $CO_2$ environment for 20 min. Finally, the media was replaced by imaging solution and without any further wash CLSM images were captured on a Zeiss LSM880 inverted confocal microscope using ZEN (Black edition) application software.
Following the same procedure mentioned above HeLa cells were incubated with 10 µM Probe **10** for 15 min at 37°C in 5% $CO_2$ environment. Subsequently, the incubation media was removed, and Live cell imaging solution was added to the dish and images were captured on a Zeiss LSM880 inverted confocal microscope without wash.

To monitor the cellular uptake and localization of Probe **11** live cell colocalization experiment was carried with LysoTracker Deep red (Invitrogen, Cat. no. L12692). HeLa cells (cell viability > 95%) at a density of ~$1\times10^5$ cells/mL were separately seeded in 35 mm glass bottom confocal dish and allowed to get 80% confluency in DMEM for 24 h at 37°C in an air-jacketed 5% $CO_2$ incubator. Cells were washed with 1× DPBS(2x's) and incubated with 10 µM Probe **11** (stock concentration 1 mM in water), for variable time (1h, 4h and 24h). Incubation media was removed, and without any wash cells were incubated with 75 nM LysoTracker Deep red for 15 min at 37°C in 5% $CO_2$ environment. at 37°C in 5% $CO_2$ environment. Finally, the media was replaced with Live Cell imaging solution (Invitrogen, Cat. no. A59688DJ), and CLSM images were acquired on a Zeiss LSM880 inverted confocal microscope without any further washing.

**Real-time Live cell imaging with Probe 11:** For real-time live cell confocal imaging HeLa cells were incubated with **11** following the identical protocol mentioned previously and without further wash the images were acquired over the duration of 3 min 11 sec at laser excitation wavelength of 488 nm (Argon ion laser, 25 mW), with a constant intensity of 5%.

All the CLSM images and movies were processed by ZEN (blue edition) software. Twenty real-time image frames were analyzed with ImageJ (version 1.54p) to generate photobleaching plot.

It's worth to mention that throughout the live cell CLSM imaging procedure 37 °C and 5% $CO_2$ atmosphere was maintained.

For **Probe 8**: laser excitation wavelength = 405 nm (detection range of emission wavelength 475-565 nm); MitoTracker Red FM: laser excitation wavelength = 561 nm (detection range of emission wavelength 580-700 nm); **Probe 10**: laser excitation wavelength = 561 nm, (detection range of emission wavelength 575-700 nm). **Probe 11**: laser excitation wavelength = 488 nm, (detection range of emission wavelength 501-625 nm); LysoTracker Deep Red: laser excitation wavelength = 633 nm (detection range of emission wavelength 640-725 nm).

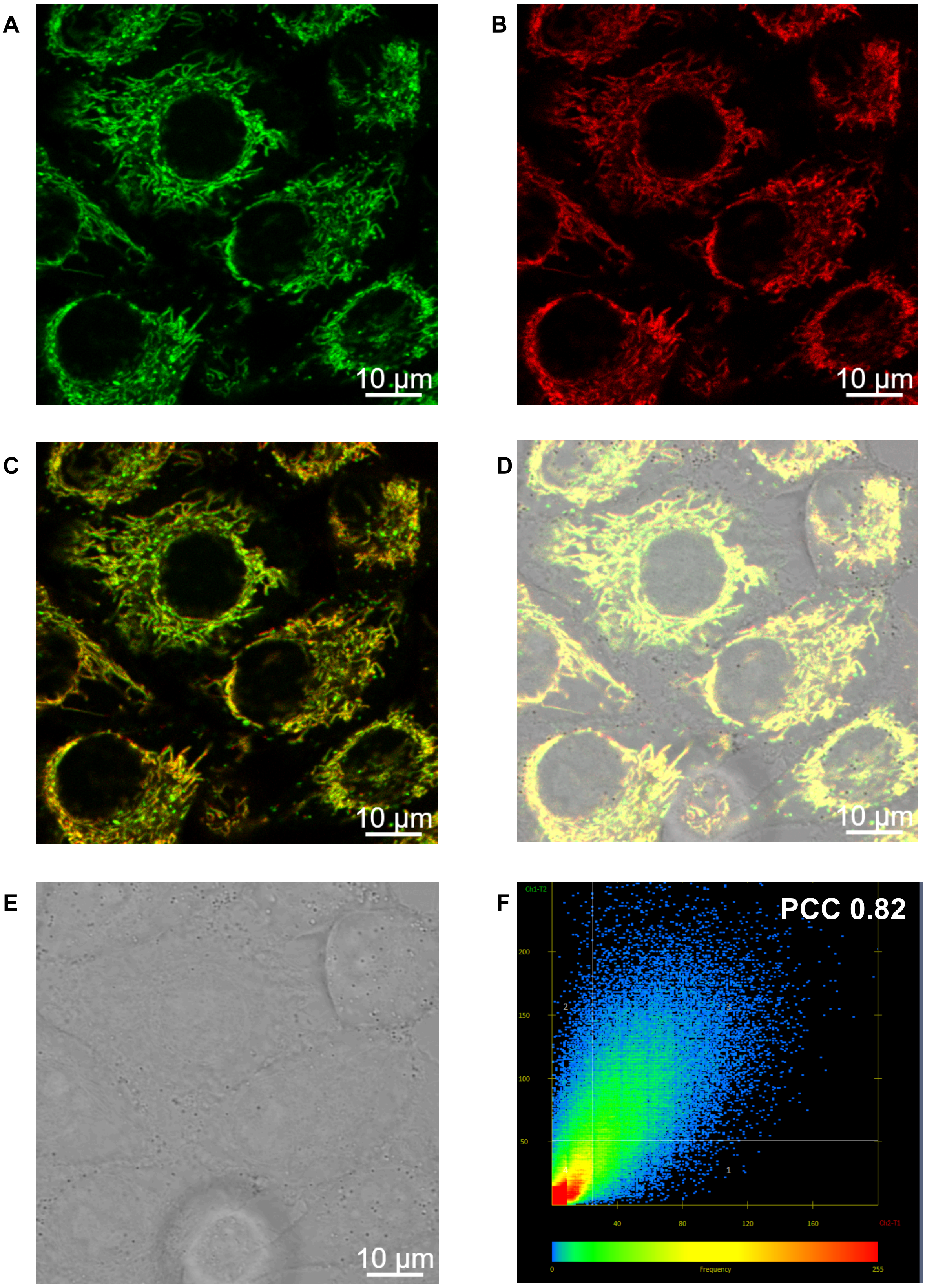


**Figure S17:** Live cell confocal laser scanning microscopic images of **8** ($\lambda_{ex}$=405 nm) colocalized with MitoTracker Red FM ($\lambda_{ex}$=561 nm) in living HeLa cells. (A) **8** (10 μM), (B) MitoTracker Red FM (200 nM), (C) Merged, (D) Merged Brightfield, (E) Brightfield and (F) PCC plot showing colocalization in mitochondria with PCC=0.82.

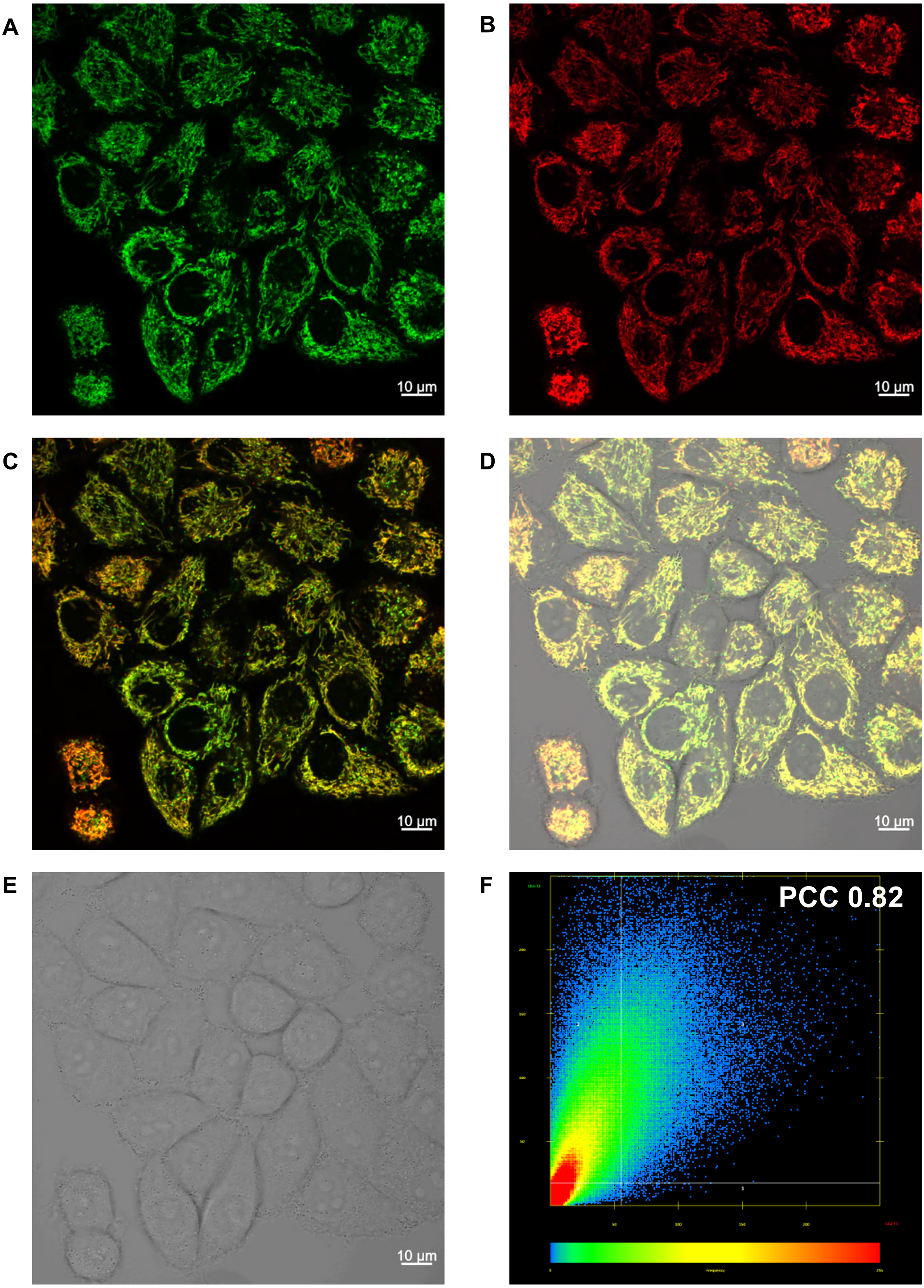


**Figure S18:** Live cell confocal laser scanning microscopic images of **8** ($\lambda_{ex}$=405 nm) colocalized with MitoTracker Red FM ($\lambda_{ex}$=561 nm) in living HeLa cells. (A) **8** (10 µM), (B) MitoTracker Red FM (200 nM), (C) Merged, (D) Merged Brightfield, (E) Brightfield and (F) PCC plot showing colocalization in mitochondria with PCC=0.82.

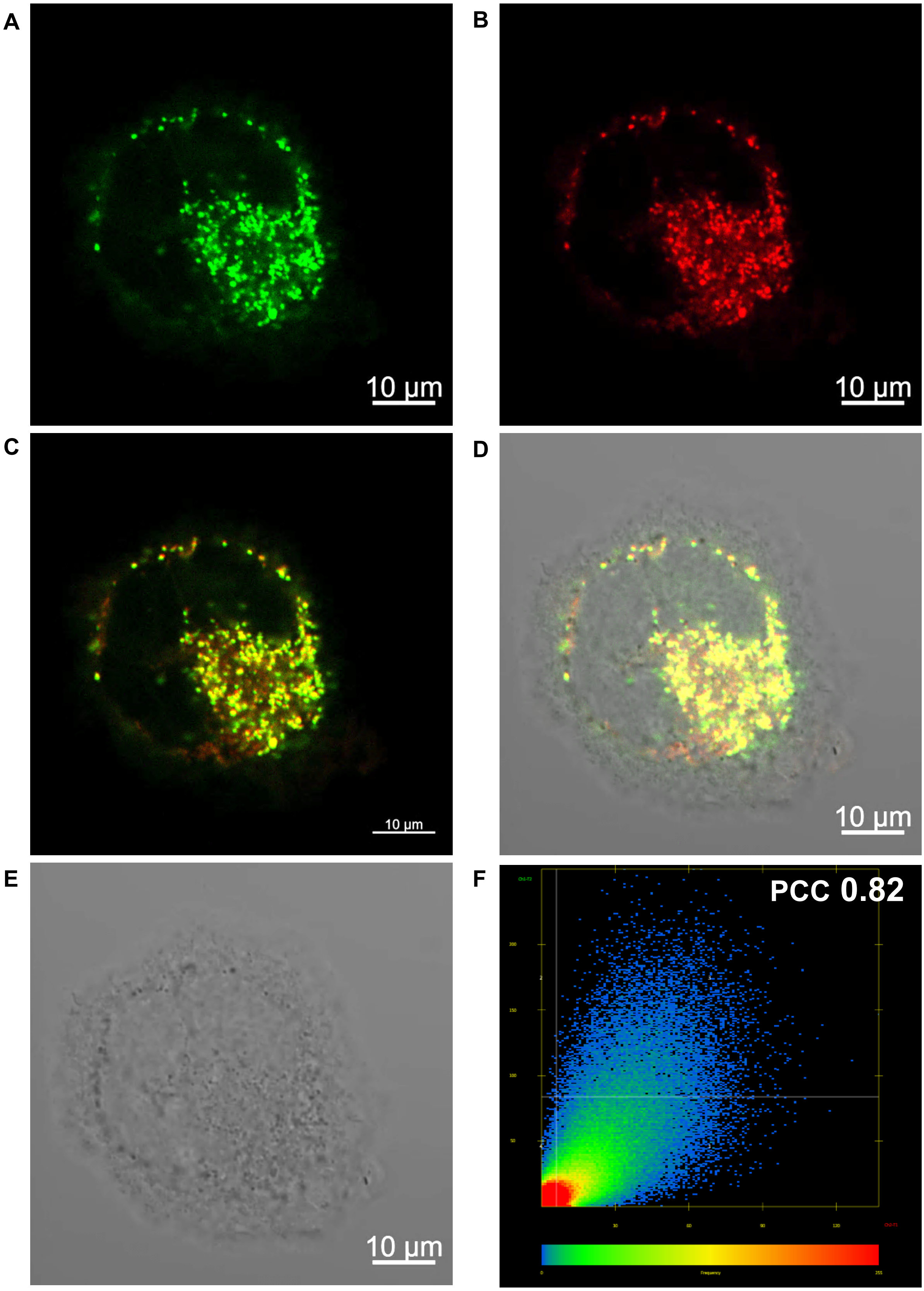


**Figure S19:** Live cell confocal laser scanning microscopic images of **11** ($\lambda_{ex}$=488 nm) colocalized with LysoTracker Deep Red ($\lambda_{ex}$=633 nm) in living HeLa cells. (A) Wash-free live cell confocal imaging using **11** (10 µM for 4h incubation), (B) LysoTracker Deep Red (75 nM, 15 min incubation), (C) Merged, (D) Merged Brightfield, (E) Brightfield and (F) PCC plot showing colocalization in lysosome with PCC=0.82.

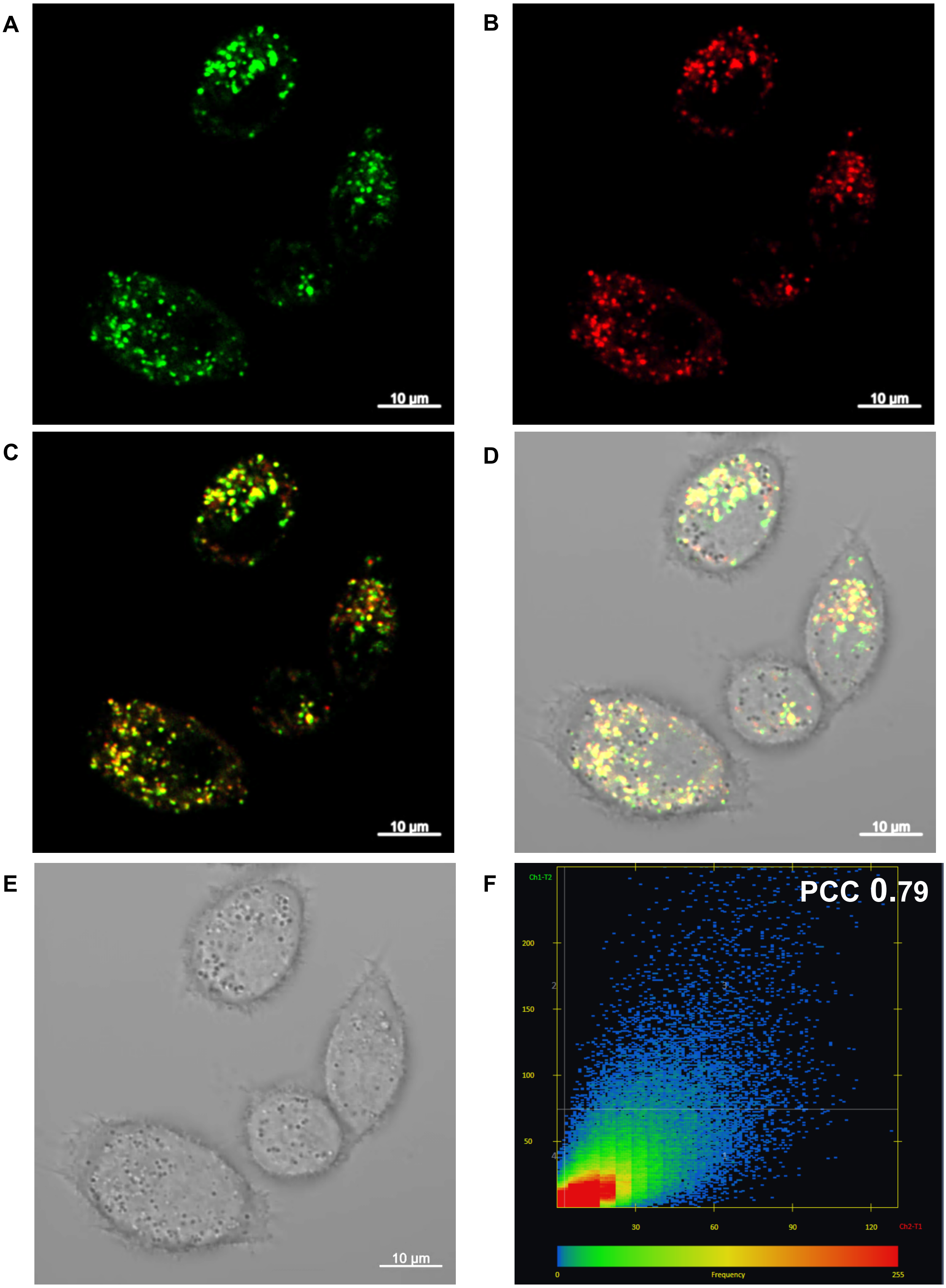


**Figure S20:** Live cell confocal laser scanning microscopic images of **11** ($\lambda_{ex}$=488 nm) colocalized with LysoTracker Deep Red ($\lambda_{ex}$=633 nm) in living HeLa cells. (A) Wash-free live cell confocal imaging using **11** (10 $\mu$M for 4h incubation), (B) LysoTracker Deep Red (75 nM, 15 min incubation), (C) Merged, (D) Merged Brightfield, (E) Brightfield and (F) PCC plot showing colocalization in lysosome with PCC=0.79.

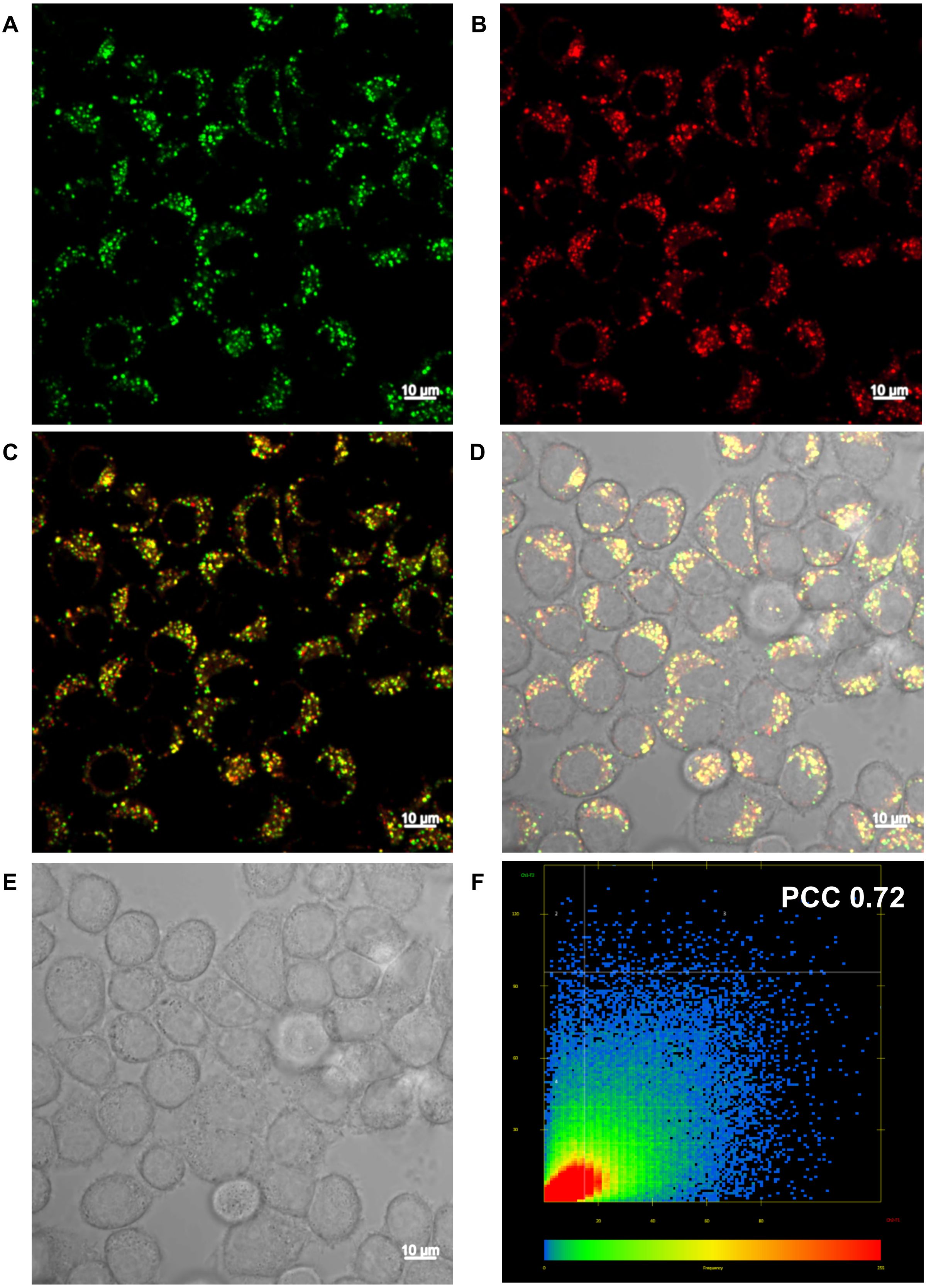


**Figure S21:** Live cell confocal laser scanning microscopic images of **11** ($\lambda_{ex}$=488 nm) colocalized with LysoTracker Deep Red ($\lambda_{ex}$=633 nm) in living HeLa cells. (A) Wash-free live cell confocal imaging using **11** (10 μM for 1h incubation), (B) LysoTracker Deep Red (75 nM, 15 min incubation), (C) Merged, (D) Merged Brightfield, (E) Brightfield and (F) PCC plot showing colocalization in lysosome with PCC=0.72.

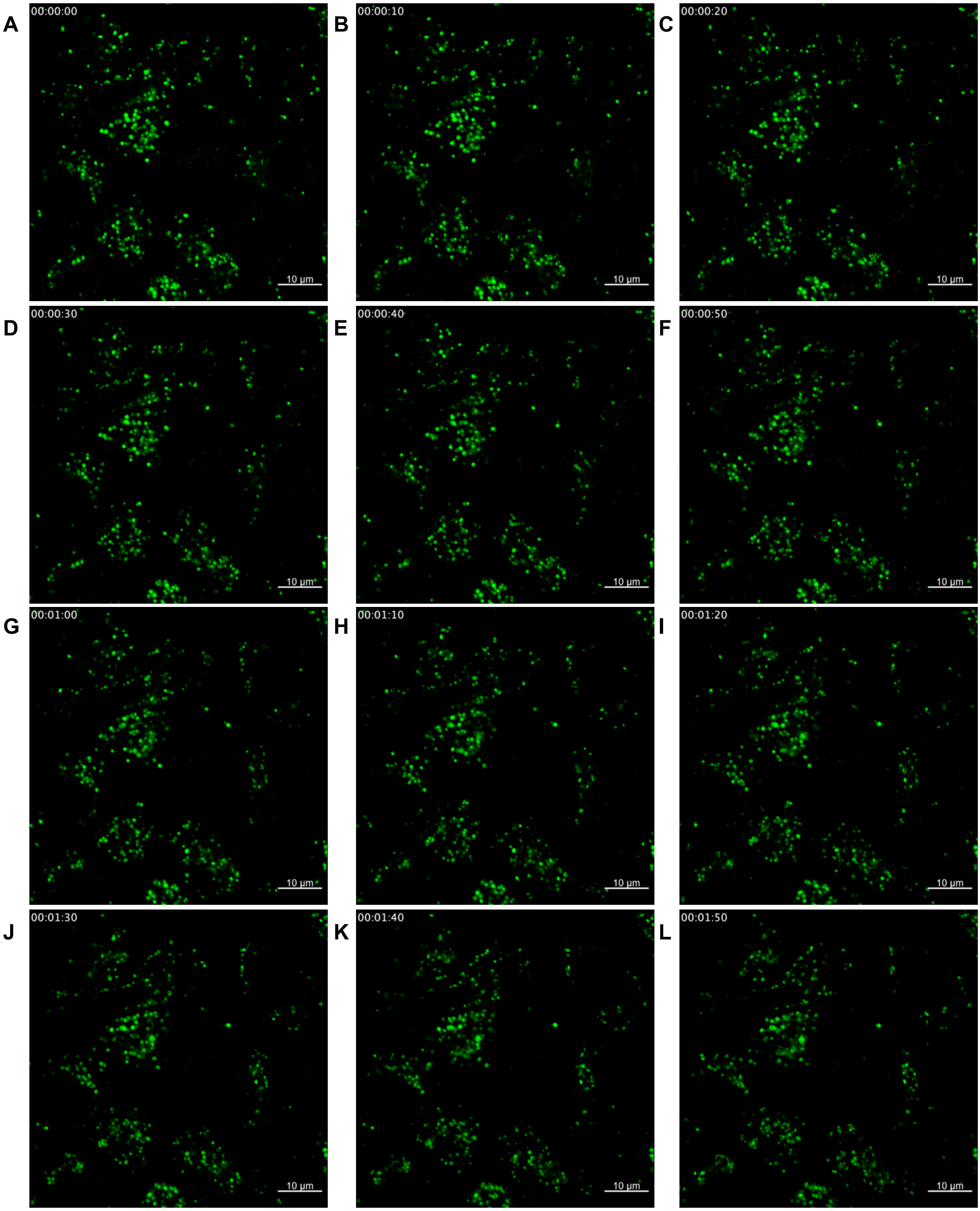
A
00:00:00
10 µm
B
00:00:10
10 µm
C
00:00:20
10 µm
D
00:00:30
10 µm
E
00:00:40
10 µm
F
00:00:50
10 µm
G
00:01:00
10 µm
H
00:01:10
10 µm
I
00:01:20
10 µm
J
00:01:30
10 µm
K
00:01:40
10 µm
L
00:01:50
10 µm

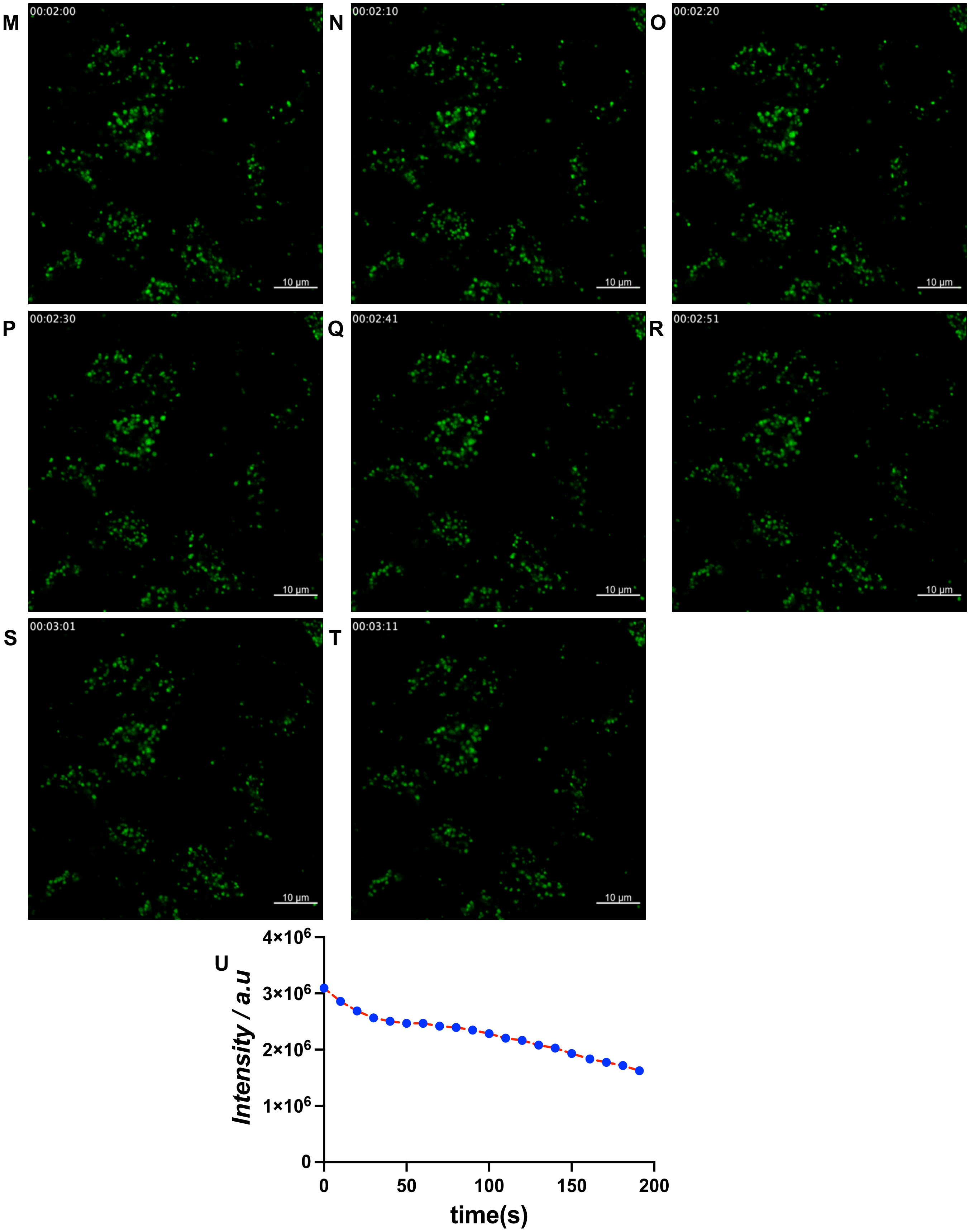


**Figure S22:** (A-T) Real-time wash free HeLa living cell confocal laser scanning microscopic images of **11** (10 μM for 1h incubation) at $\lambda_{ex}$=488 nm (Argon ion laser, 25 mW), with 5% laser intensity over the period of 3 min 11 sec (frame intervals 10.07 sec); (U) Photobleaching profile of **11** at $\lambda_{ex}$=488 nm (Argon ion laser, 25 mW), with a laser intensity of 5% over the period of 3 min 11 sec.

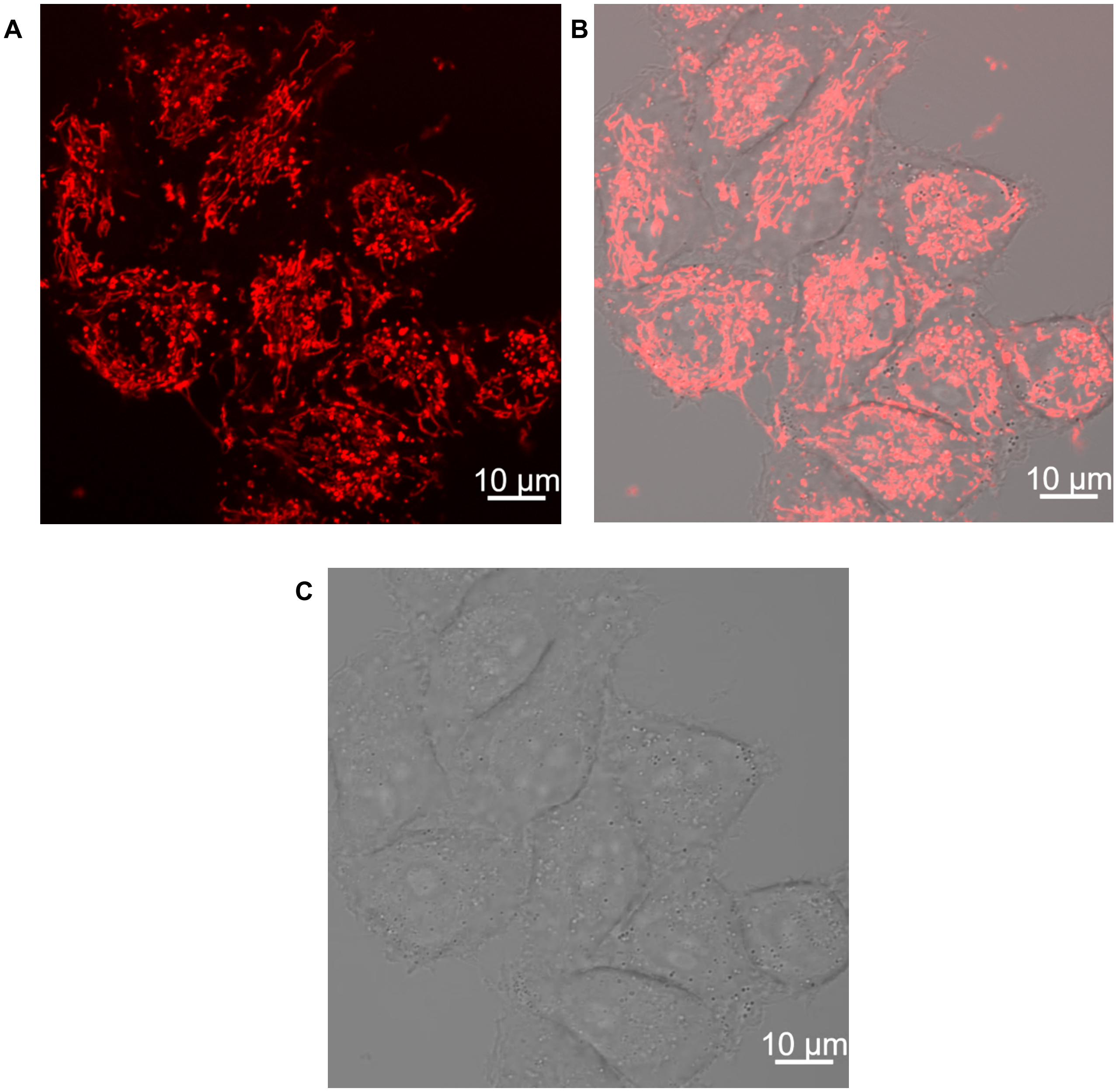


**Figure S23:** Live cell confocal laser scanning microscopic images using **10** ($\lambda_{ex}$=561 nm) under wash-free condition in living HeLa cells. (A) **10** (10 μM for 15 min incubation), (B) Merged Brightfield, (C) Brightfield.

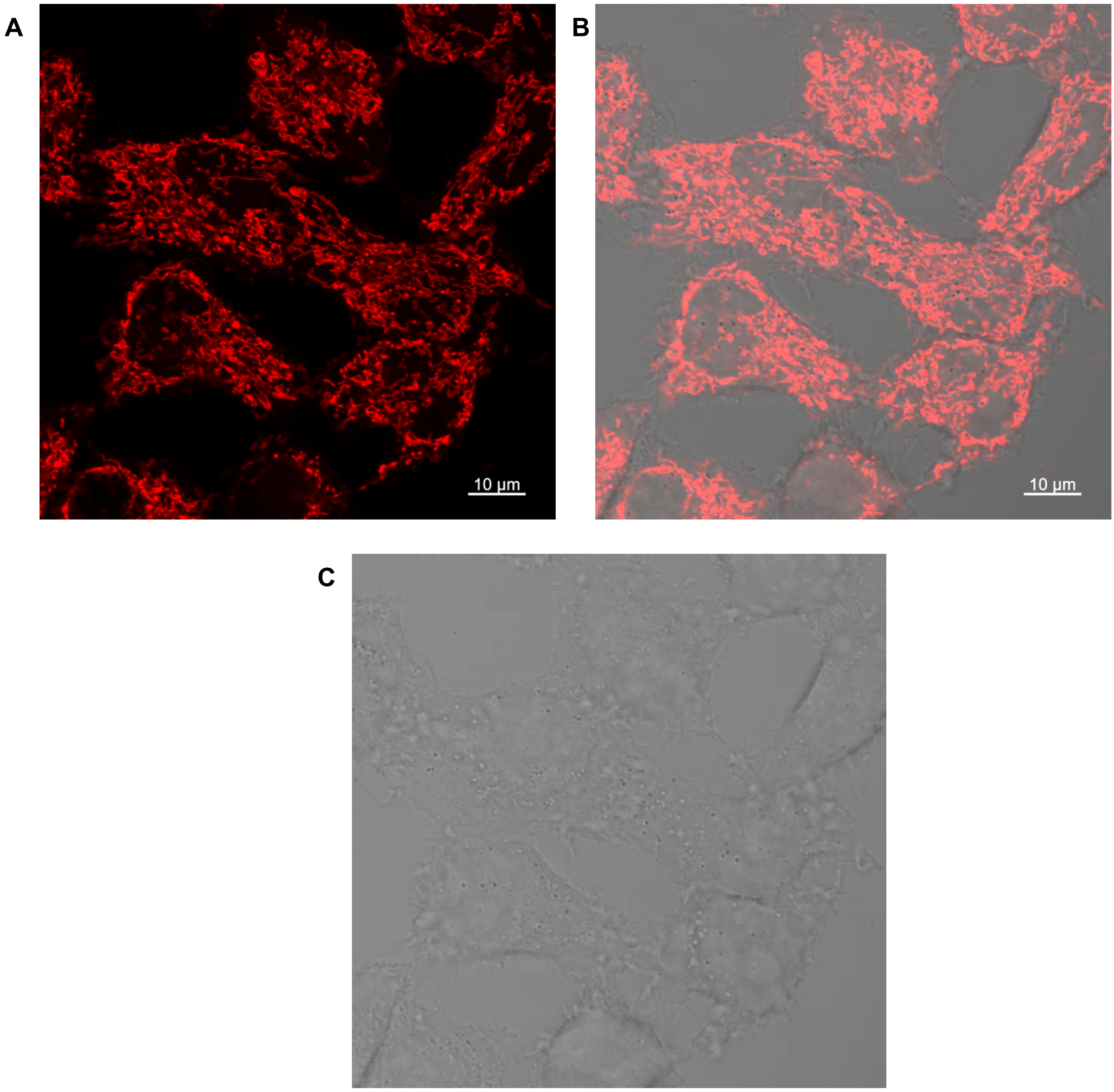


**Figure S24:** Live cell confocal laser scanning microscopic images using **10** ($\lambda_{ex}$=561 nm) under washing condition in living HeLa cells. (A) **10** (10 μM for 15 min incubation), (B) Merged Brightfield, (C) Brightfield.

### 6. Chemo-proteomic Profiling Analysis, Procedures, and Validation

#### In HEK293T cell lysate biotinylated pyridinium/pyrimidinium salts treatment for protein-level validation:

Light sources used for HEK293T cell lysate photo-activation:

456 nm Light source = Kessil PR-160L-456 nm LED, 40W

475 nm Light source = EvoluChem 475PF LED,18W

525 nm Light source = Kessil PR-160L-525 nm LED,40W

**Lysate generation:**
Human embryonic kidney cells (HEK293T) were cultured in 100 mm dishes (Thermo Scientific, BioLite Cell Culture Treated Dishes) containing Dulbecco's Modified Eagle's Medium (DMEM, Genesee Scientific) supplemented with 10% Fetal Bovine Serum (Gibco) and 1% Penicillin-Streptomycin (Sigma-Aldrich). Cells were maintained at 37°C in the incubator with routine passage of 5% $CO_2$ in a humidified atmosphere until ~ 90 % confluent. The cells were collected by rinsing the plate with ice-cold DPBS (2x 1 mL) followed by scraping. Cells were transferred to 1.5 mL microcentrifuge tubes and pelleted by centrifugation at 4 °C (5 min, 500g). The pellets were resuspended in lysis buffer containing 50 mM HEPES pH 7.5, 1.5 mM $MgCl_2$, 1% SDS, 0.8% NP-40, and 1x protease inhibitors (Thermo Scientific) and placed in the ice for 30 minutes followed by sonication (QSonica) at 20% amplitude, 2 seconds ON, 3 seconds OFF, for 30 seconds. Cell debris was discarded through centrifugation at 20,000 g for 20 minutes at 4°C. The supernatant was subsequently transferred to a clean microcentrifuge tube, and the protein concentration was determined using the Pierce™ 660nm Protein Assay Kit (Thermo Scientific).

**General procedure for HEK293T cell lysate labelling:**
In 200 µL flat bottom insert vials (Fisher Scientific), HEK293T cell lysates (100 µg, 2 mg/mL) were separately treated with biotinylated pyridinium/pyrimidinium salts **8a**, **9a**, or **11a** (0,1, 10, and 50 µM) in 20 mM phosphate buffer, pH 7.4,incubated for 30 minutes followed by irradiation for 20 minutes in a Hepatochem PhotoredoxBox$^{TM}$ reactor using the conditions and wavelengths specified below*. The reaction mixtures were transferred to 2 mL microcentrifuge tube, precipitated using ice-cold methanol: chloroform (4:1) mixture and centrifuged at 20,000 g for 20 minutes at 4°C to isolate protein pellets. The protein pellets were washed with cold methanol(2x's), centrifuged, and then air-dried at RT. The protein pellets were resuspended in 50 µL Laemmli buffer (2x) containing 10 mM DTT and heated at 55°C for 30 minutes. Samples were then subjected to Western blot analysis, as outlined in the protocol for western blot analysis,.
*Controls with probe incubation but no irradiation were allowed to stand under ambient conditions, while all other experimental conditions remained unchanged.

**Conditions for establishing probe lysate labelling efficiency:**

**1a**:
[**1a**]= 0, 1, 10, 50 µM
hν source: 456 nm LED.
Controls: 0 µM probe + hν, 50 µM **1a** no hν

**8a**:
[**8a**]= 0, 1, 10, 50 µM
hν source: 456 nm LED.

Controls: 0 μM probe + hν, 50 μM **8a** no hν

**9a**:
[**9a**]= 0, 1, 10, 50 μM
hν source: 456 nm LED.
Controls: 0 μM probe + hν, 50 μM **9a** no hν

**11a**:
[**11a**]= 0, 1, 10, 50 μM
hν source: 475 nm LED.
Controls: 0 μM probe + hν, 50 μM **11a** no hν

**Conditions for establishing wavelength dependent labelling with probes:**

Probes used: **8a**, **9a**, **11a**
[probe]= 50 μM

Wavelengths:
456 nm
475 nm
525 nm

**Protocol for Western blot analysis**: Standard immunoblotting procedures were employed. Proteins were separated on 4-20% Mini-Protean® TGM™ Precast Gels (Bio-Rad, Cat. no. 4561094) at 100V for 90 min, followed by transfer onto 0.45 μm nitrocellulose membranes (Bio-Rad, Cat. no. 1620117) at 100V for 1 h using the Mini Trans-Blot® Cell (Bio-Rad). The membranes were subsequently washed with 1x PBST for 5 min and blocked with EveryBlot Blocking Buffer (Bio-Rad, Cat. no. 12010020) at 4 °C for 16 h. Finally, the membranes were incubated with HRP-linked Streptavidin (1:15000) (Abcam) at 4 °C for 16 h. After washing the membrane (6x's each for 5 min) with 1x PBST it was incubated with the Clarity™ Western ECL Substrate (Bio-Rad, Cat. no. 1705060) for 2 min in dark with gentle shaking. The membrane was visualized with UVP ChemStudio PLUS Imaging Systems, analytic jena (Avantor®).

A

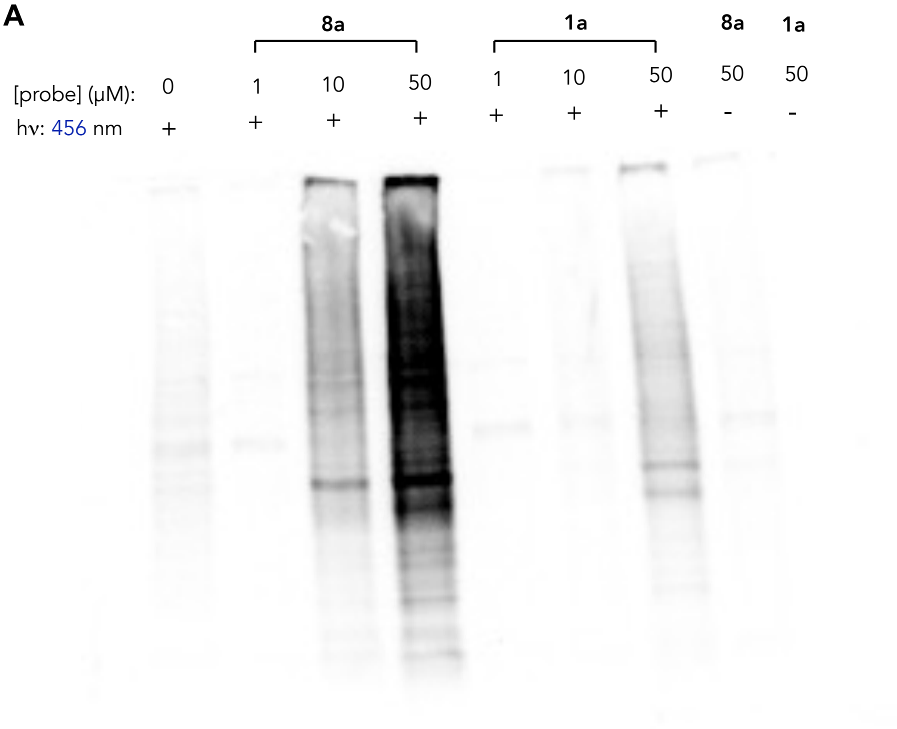


B

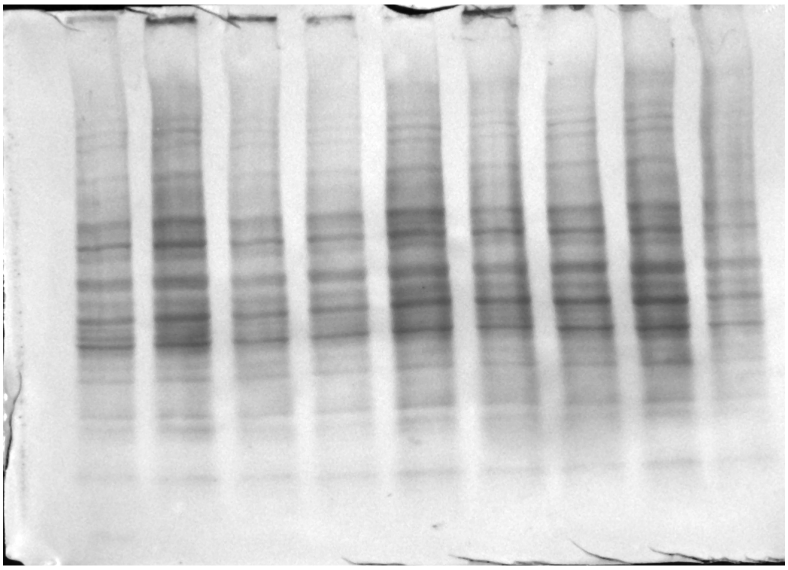

**Figure S25: (A)** Western blot analysis of proteome HEK293T cell lysate profiling with different concentration (1, 10, and 50μM) of probe **1a** and **8a** at 456 nm wavelength of light irradiation for 20 minutes and **(B)** Corresponding Coomassie stain.

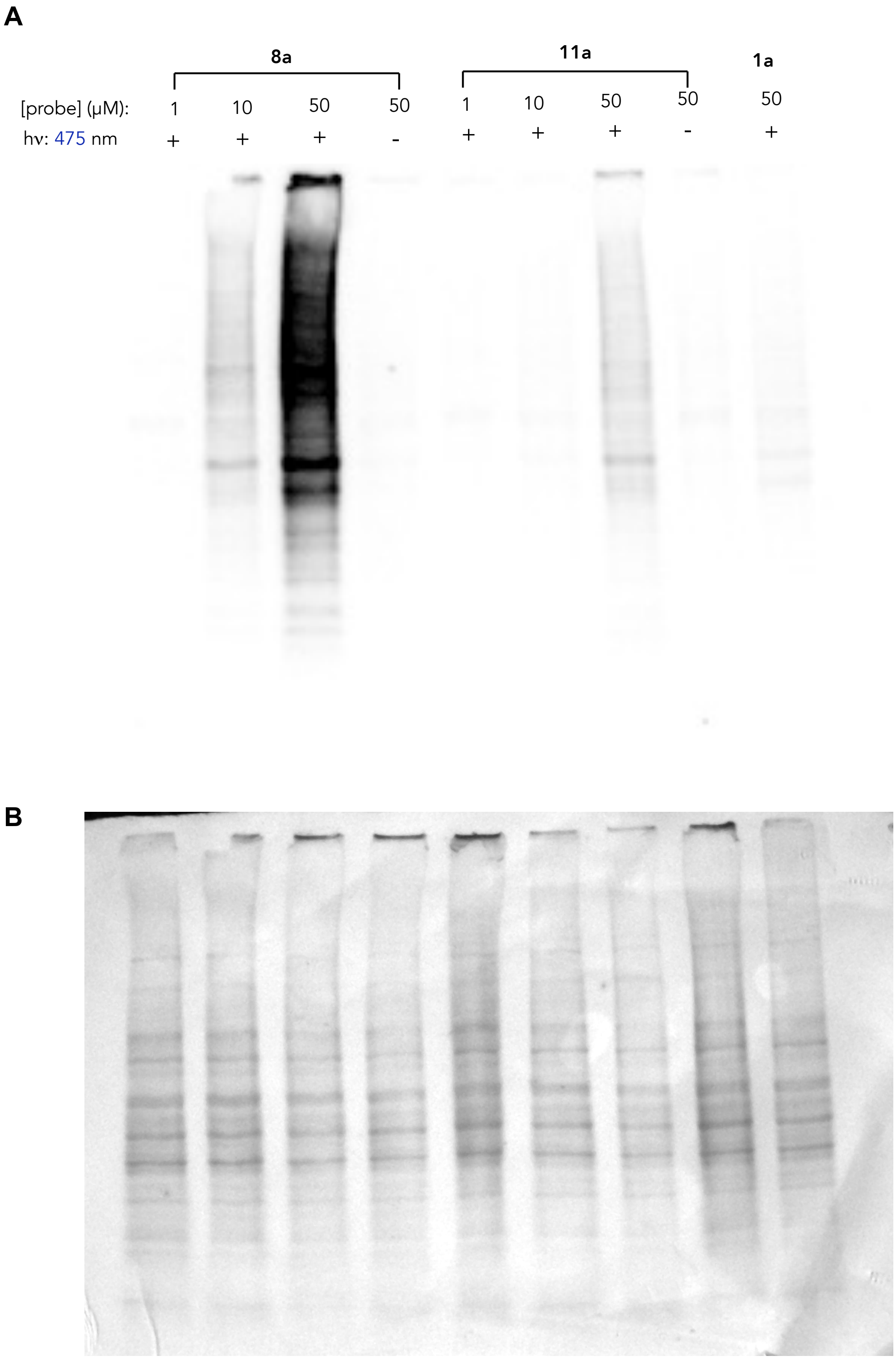


**Figure S26: (A)** Western blot analysis of proteome HEK293T cell lysate profiling with different concentration (1, 10, and 50μM) of probe **1a, 8a** and **11a** at 475 nm wavelength of light irradiation for 20 minutes and **(B)** Corresponding Coomassie stain.

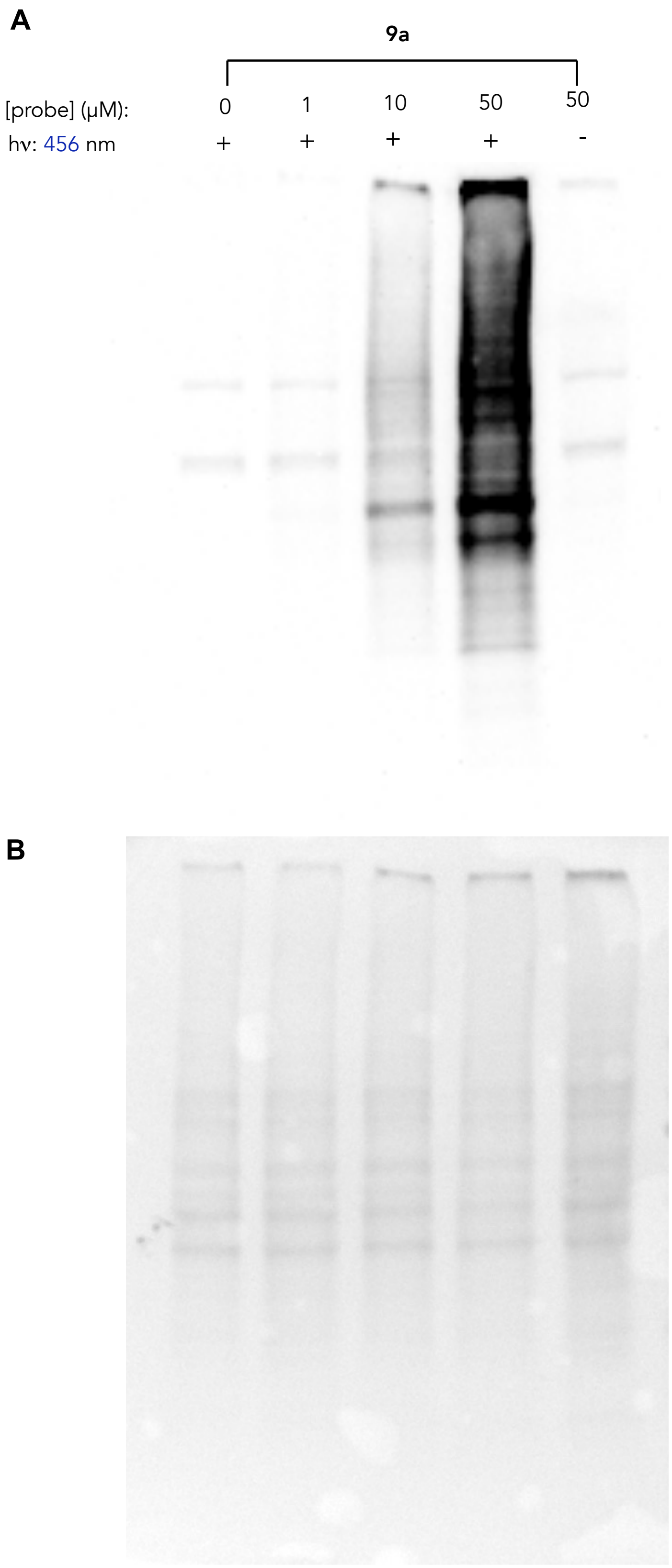


**Figure S27: (A)** Western blot analysis of proteome HEK293T cell lysate profiling with different concentration (1, 10, and 50μM) of probe **9a** at 456 nm wavelength of light irradiation for 20 minutes and **(B)** Corresponding Coomassie stain.

A

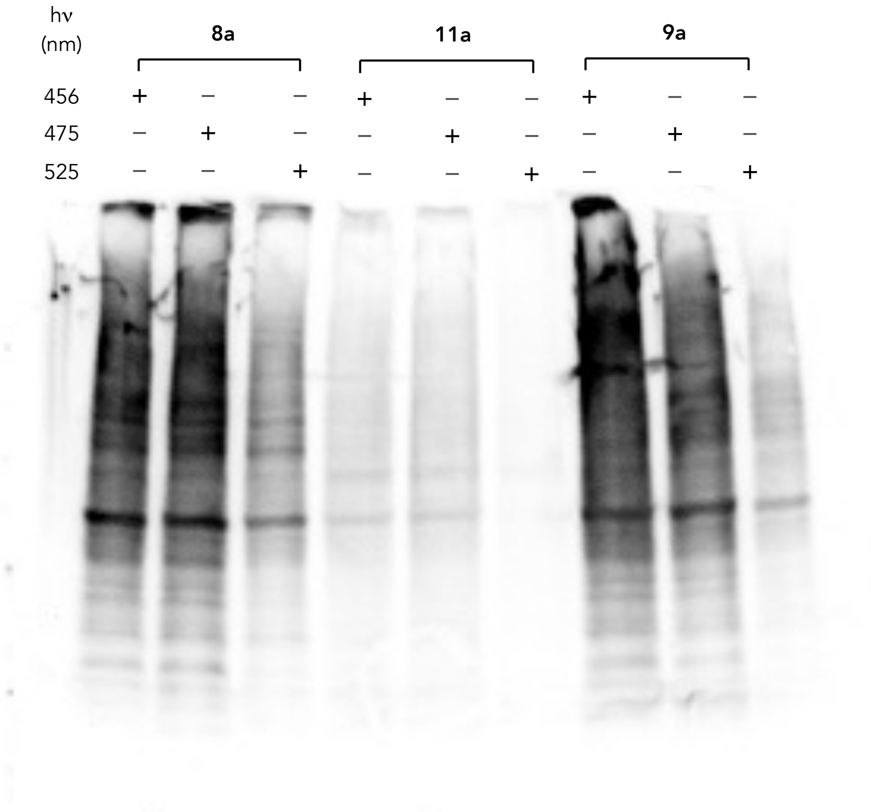


B

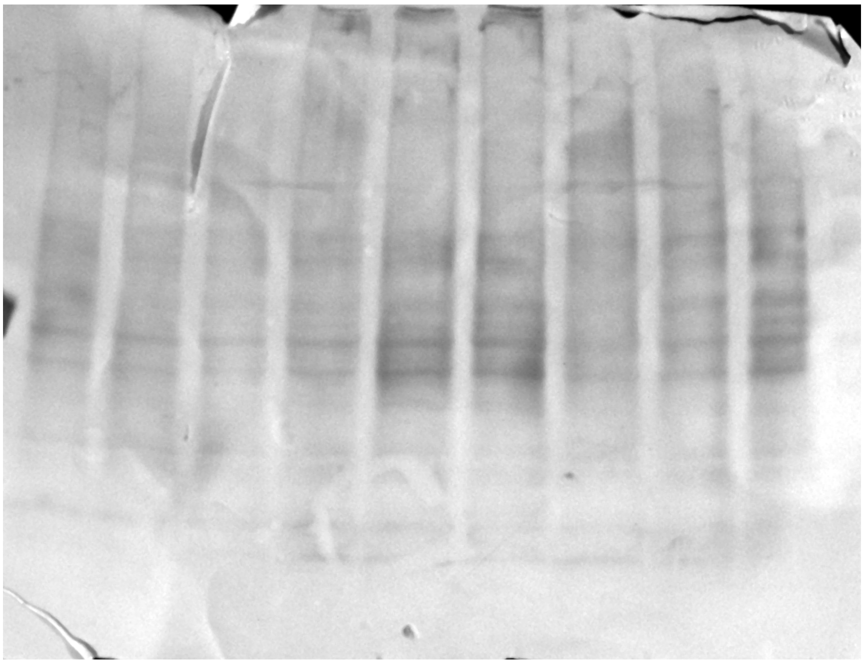

**Figure S28: (A)** HEK293T cell lysate proteome profiling with probe **8a, 9a** and **11a** (concentration 50μM) by irradiating at 456, 475 and 525 nm wavelength of light for 20 minutes and **(B)** Corresponding Coomassie stain.

**Cell culture and *in-situ* probe 8b treatment**

Human cervix adenocarcinoma epithelial cells (HeLa) were grown in T75 flasks (Nunc™ Easy Flask™ Cell Culture Flasks, Thermo Scientific) containing Dulbecco's Modified Eagle's Medium (DMEM, Genesee Scientific) supplemented with 10% Fetal Bovine Serum (Gibco) and 1% Penicillin (10000 units/mL)-Streptomycin (10 mg/mL) (Sigma-Aldrich). All cells were maintained at 37°C in the incubator with routine passage of 5% $CO_2$ in a humidified atmosphere.

For *in situ* labeling, HeLa cells were grown in T75 flask to 90% confluence (cell viability >95%), washed with 1x DPBS(Gibco), treated with 100x stocks of the probe 8b in DMEM media to generate desired final concentrations (0 and 100 μM) in 0.2% DMSO. HeLa cells were incubated for 1 hour at 37°C with 5% $CO_2$ followed by photo-activation at 530 nm (LEDA-530, Product code: E50.100.530, AMUZA) for 20 minutes at 4°C. As a positive control, another T75 culture flask was prepared where Hela cells were incubated with 100 μM of probe 8b for 1h at 37°C with 5% $CO_2$ but were not exposed to light irradiation. In all the cases, cells were harvested using a cell scraper, washed three times with cold 1x DPBS (Gibco), centrifuged at 500g for 5 minutes at 4°C to isolate cell pellets, and the pellets were stored at -80°C until further use.

**In situ probe 8b sample processing for protein-level characterization**:

Cell pellets were resuspended in 50 mM HEPES pH 7.5, 1.5 mM MgCl2, 1% SDS, 0.8% NP-40, and 1x protease inhibitors (Thermo Scientific) and kept in the ice for 30 minutes and lysed by sonication (QSonica) at 20% amplitude, 2 seconds ON, 3 seconds OFF, for 30 seconds. Cell debris was discarded through centrifugation at 20,000 g for 20 minutes at 4°C. The supernatant was subsequently transferred to a clean microcentrifuge tube, and the protein concentration was determined using the Pierce™ 660nm Protein Assay Kit (Thermo Scientific).

The following table presents the three different sets of conditions used in live HeLa cell labeling experiments for protein-level characterization:

**Table 1:**

| Condition | [8a] (μM) | Incubation period (min) | wavelength of light (nm) | Irradiation time (min) |
|---|---|---|---|---|
| Negative Control | 0 | 60 | 530 | 20 |
| Positive Control | 100 | 60 | - | - |
| Treated | 100 | 60 | 530 | 20 |

500 μL protein samples (2mg /mL) from each condition were taken in microcentrifuge tubes for CuAAC reactions with 100 μM of the acid cleavable DADPS Biotin Alkyne (Click Chemistry Tools, stock solution 10 mM in DMSO, CCT-1331, Vector Laboratories), 250 μM CuSO4 (stock solution 25 mM in water), 500 μM BTTP (stock solution 50 mM in 9:1 DMSO:water) (Click Chemistry Tools, CCT-1414, Vector Laboratories), and 2.5 mM sodium ascorbate (stock solution 250 mM in water) and incubated for 1 hour at RT in the dark. Chilled methanol: chloroform (4:1) precipitation was used to terminate the click reaction and to remove excess biotin reagent: 800 μL ice-cold methanol, 200 μL chloroform and 300 μL ice-cold water was added, and the tube was vortexed. Subsequently, the precipitate was centrifuged at 20,000g for 15 minutes at 4 °C and the top and bottom layers were discarded carefully to obtain the protein pellets. The pellets were suspended in 500 μL chilled methanol with brief vortexed,

tube was centrifuged at 20,000g for 5 minutes at 4 °C and the supernatant was discarded. This methanol washing step was repeated. The protein pellets were air-dried at RT and resuspended in 400 µL 1x PBS containing 2.5% SDS via sonication. 1x PBS (4 mL) was added to adjust the SDS concentration to ~ 0.2%. NeutrAvidin UltraLink Resin (100 µL, 50% slurry, Thermo Fisher Scientific) was washed with 1x PBS (3 x 0.5 mL). Resin beads were resuspended in 500 of 1x PBS and subsequently transferred to LoBind tubes (Eppendorf) containing the protein solution. The tubes containing the resin beads and protein were rotated for 2 hours at RT and then at 4 °C overnight. Beads were transferred to a MobiCol column (Boca Scientific), harvested by centrifugation at 2000g for 3 minutes and the supernatant was discarded. The beads were washed at RT with 3 x 1 mL of each of the following solutions: (i) 0.2% SDS in PBS; (ii) 1M NaCl in PBS; (iii) freshly prepared 6M urea in PBS; (iv) PBS. For each wash, the beads were rotated for 5 min at RT. Prior to the final washing cycle, the beads were transferred to the Screw Cap Pierce Spin Column (Thermo Fisher Scientific). The beads were resuspended in 0.5 mL freshly prepared 6M urea in PBS. DTT (5 µL, 1M stock solution in water, final concentration 10 mM) was added, and the tube was rotated at 55 °C °C for 30 min. A freshly prepared solution of 2-iodoacetamide (25 µL, stock solution of 500 mM in PBS, final concentration 25 mM) was subsequently added to the tube and rotated at room temperature for an additional 30 minutes in the dark. Beads were harvested by centrifugation at 3000g for 2 min and the flow-through was discarded. Beads were washed with PBS (3 x 0.5 mL) and 50 mM Tris-HCl buffer pH 8.0 (3 x 0.5 mL). Beads were resuspended in 50 mM Tris-HCl buffer pH 8.0 (300 µL), treated with RapiGest to a final concentration of 0.05% (v/v), and incubated with 2.5 µg Trypsin/Lys-C (Promega) at 37 °C for 16 hours. The MobiCol column (Boca Scientific) was placed in a clean 2 mL Protein LoBind Tube (Eppendorf) and centrifuged at 3000g for 2 minutes to collect the flow-through. For all the samples, the DADPS linked bound peptides were subsequently cleaved from the resin by incubating with 200 µL of 10% formic acid in water for 30 minutes (2x) and flow-through was collected. All the flow-through fractions were combined and dried on a SpeedVac (Thermo Fisher Scientific).
The digested peptide samples were fractionated using a fractionation kit (Pierce™ high pH Reversed-Phase Fractionation Kit, Thermo Fisher Scientific, Cat. no. 84868) according to manufacturer's instructions. The peptide fractions were eluted from the spin column with consecutive solutions of 0.1% triethylamine combined with MeCN (5-50% MeCN). The fractions were dried on a SpeedVac, and stored at -80 °C until ready for mass spectrometer injection.

**Quantitative proteomics using two-dimensional nano liquid chromatography-tandem mass spectrometry (nLC-MS/MS):**

**LC-MS/MS analysis for DIA Method:**
Fractionated digested peptides samples were resuspended in 30 µL of 0.1% FA in Optima grade $H_2O$. Two microliters of each fractionated sample were injected into the PepMap Neo Trap column (P/N 174502, Thermo Fisher Scientific), and eluted on the PepMap RSLC C18, PN ES 900 HPLC column (3 µm, 100 Å, 75 µm x 15 cm) using the Vanquish Neo UHPLC system (Thermo Fisher Scientific) coupled to an Orbitrap Exploris 240 mass spectrometer (Thermo Fisher Scientific). Mobile phase A was 0.1% formic acid in $H_2O$ (optima grade), and mobile phase B was 0.1% formic acid, 80% acetonitrile (optima Grade) in $H_2O$ (Optima Grade). Chromatographic separation was achieved using a linear gradient starting with a flow rate of 350 nL min$^{-1}$ from 2% Buffer B (0.1% FA in 80% MeCN) followed by incremental increase to 50% over 46 min, 50-90% B for 14 min, 90-95% B for 2 min, and hold at 95% B for 8 min. The eluted peptides were analyzed with a Orbitrap Exploris 240 mass-spectrometer (Thermo Fisher Scientific) equipped with nano-LC electrospray ionization source (applied voltage: 2.0 kV, and the ion transferred tube temperature maintained at 280 °C). Under the positive-ion mode, full-scan mass spectra were acquired over the m/z range from 380 to 2000 using the Orbitrap mass analyzer with 120k MS1 resolution, RF Lens 70%, 28 ms max injection time and 300% AGC. MS/MS fragmentation is performed in a data independent mode. MS2 features between 380-2000 amu with resolution of 15k, isolation window (m/z) 10,

overlapping window set to (m/z) 1, maximum injection time 25 ms, a normalized HCD of 30%, and AGC of 100%.

**DIA data analysis:**

All the DIA data was analyzed and quantified by FragPipe GUI v 23.0 computational Platform using MSFragger-DIA-NN Algorithm [FragPipe GUI v16.0 with MSFragger (version 4.3), Philosopher (version 5.1.1), DIA-NN (version 1.8.2 beta 8), IonQuant (version 1.11.11) and EasyPQP (version 0.1.52)].[16] DIA_SpecLib_Quan close search workflow was used to directly search and quantify the DIA data.[16] The search results were processed by MSBooster for deep learning-based score calculation. Percolator was used for rescoring and error probability calculation, while ProteinProphet was used for protein inference. Philosopher was used for FDR filtering, and EasyPQP was used for spectral library building. For the protein identification and quantitative data, the search was performed against the Human Uniprot proteome database (UP000005640) with carbamidomethylated cysteine (+57.02146 Da) set as a static modification, oxidation of methionine (+15.9949 Da) and acetylation of N-termini (+42.01106 Da) set as dynamic modifications. Trypsin was specified as the proteolytic enzyme with up to two missed cleavage sites allowed. Closed search of probe modified sites was performed with precursor and fragment ion tolerances set to 10 ppm and 20 ppm, respectively. Peptide length was set to 5– 50, and peptide mass range was set to 500–5000. Search results were filtered with 1% FDR for peptide and protein identification and quantification, using Percolator as part of the Philosopher toolkit (v5). For the downstream analysis, peptide-protein quantification, DIA-NN output file was analyzed by FragPipe-Analyst web application. High confidence protein hits are defined as those with a protein p-value (Prob) < 0.05 found in the experimental samples but do not appear in the lists of control samples.

**Statement on proteomics data accessibility:** Mass spectrometry datasets will be deposited upon acceptance of this work for publication.

**In situ probe 8b sample processing for protein-level validation using Western blot analysis:**

To validate live cell labelling efficiency of the probe 8b, 400 μg protein (2mg/ mL) isolated from the three distinct live cell labelling experiments (**Table 3**) was subjected to a click reaction following the previously mentioned procedure. The three different conditions under which live cell labeling experiments were conducted are tabulated below:

**Table 2:**

| Condition | [8a] (μM) | Incubation period (min) | wavelength of light (nm) | Irradiation time (min) |
|---|---|---|---|---|
| 1 | 100 | 60 | N/A | 0 |
| 2 | 50 | 60 | 530 | 20 |
| 3 | 100 | 60 | 530 | 20 |

The protein pellets obtained after the click reaction were air-dried at RT and subsequently dissolved in 200 μL Laemmli buffer (2x) containing 10 mM DTT and heated at 55°C for 30 minutes. Samples were then subjected to Western blot, as outlined in the protocol for Western blot analysis.

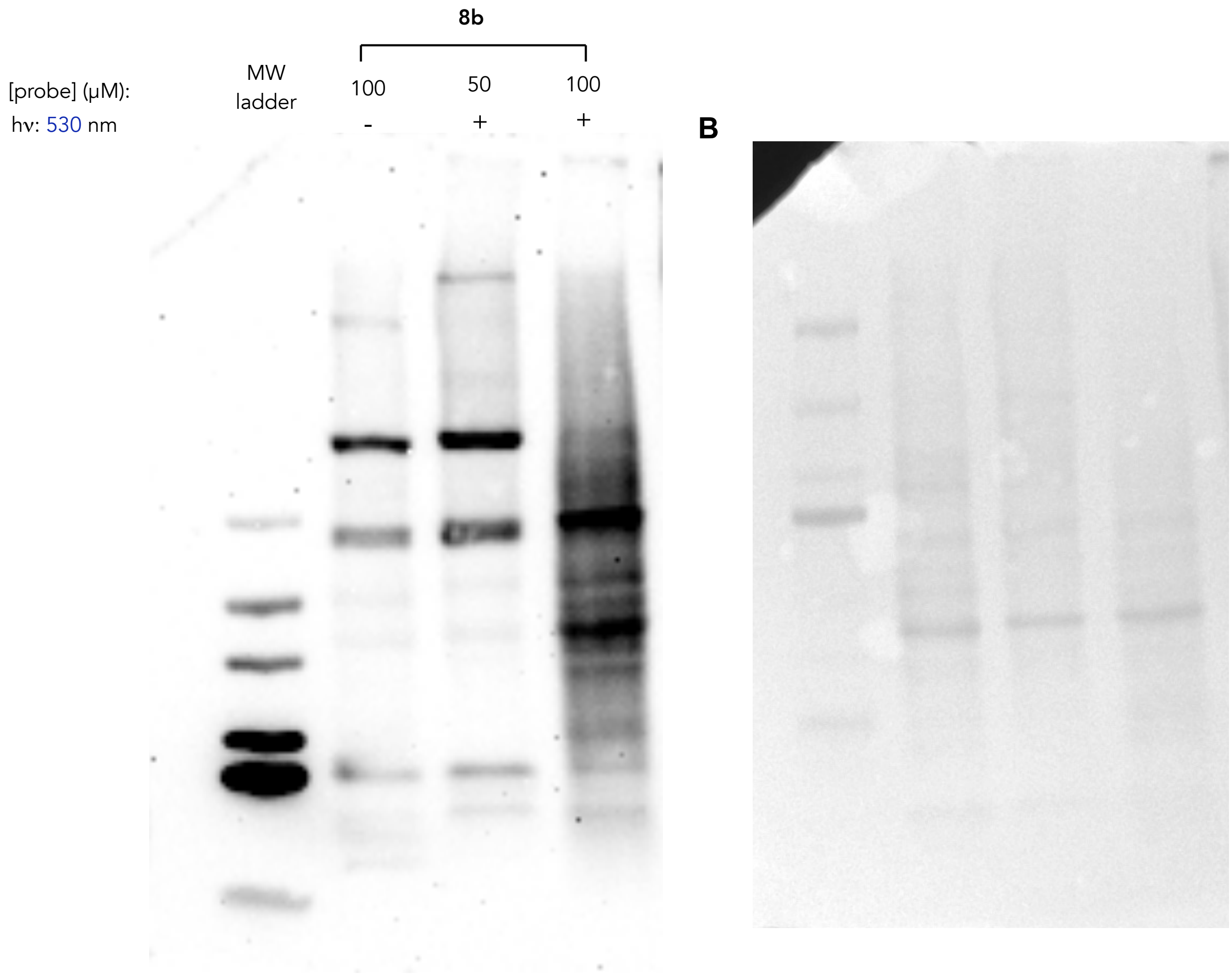


**Figure S29:** Validation of live cell photolabeling by proteome profiling of HeLa live cell with probe **8b** at 530 nm wavelength for 20 minutes.

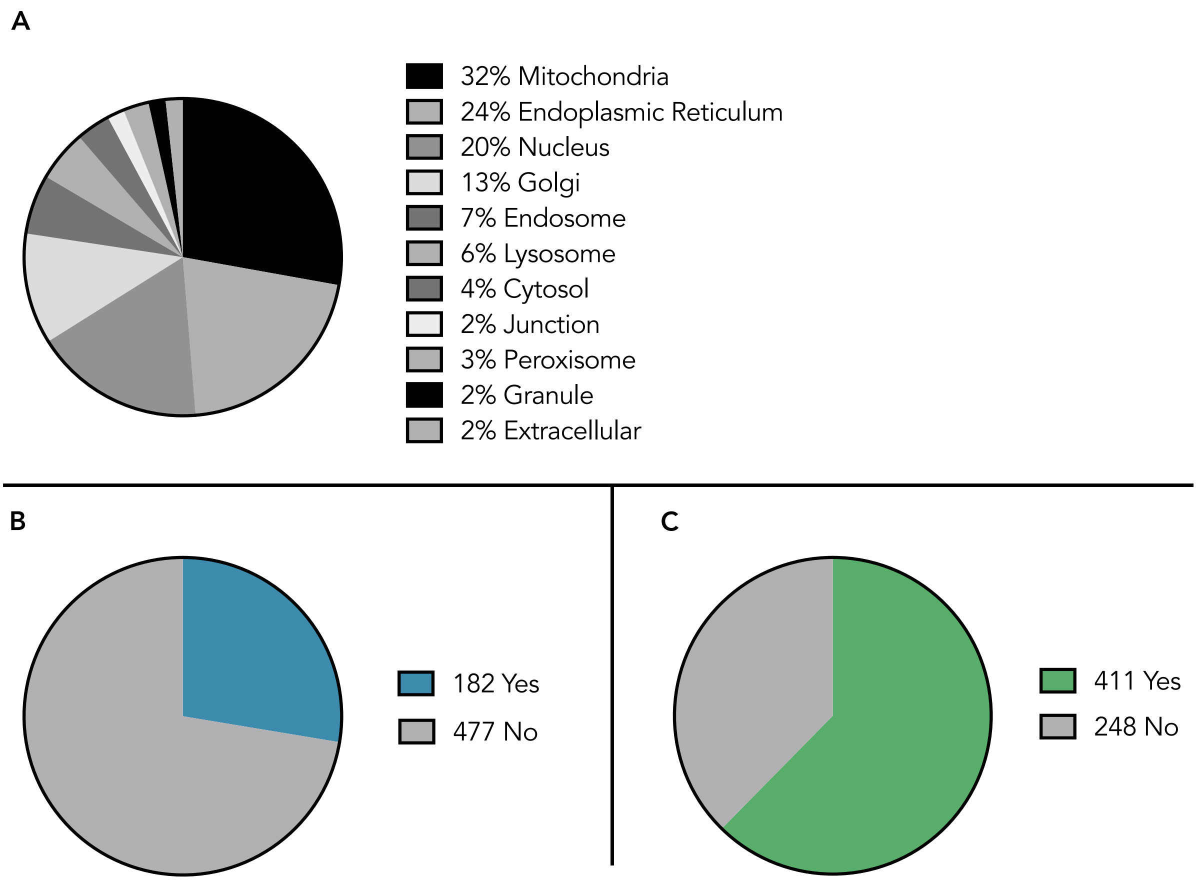


**Figure S30.** UniProt analyses of enriched proteins. (**A**) UniProt Subcellular localization analyses. For proteins with multiple assigned locations, each assignment counts as one for each component. (**B**) DrugBank associations of enriched proteins from UniProt. (**C**) Membrane association of enriched proteins from UniProt.

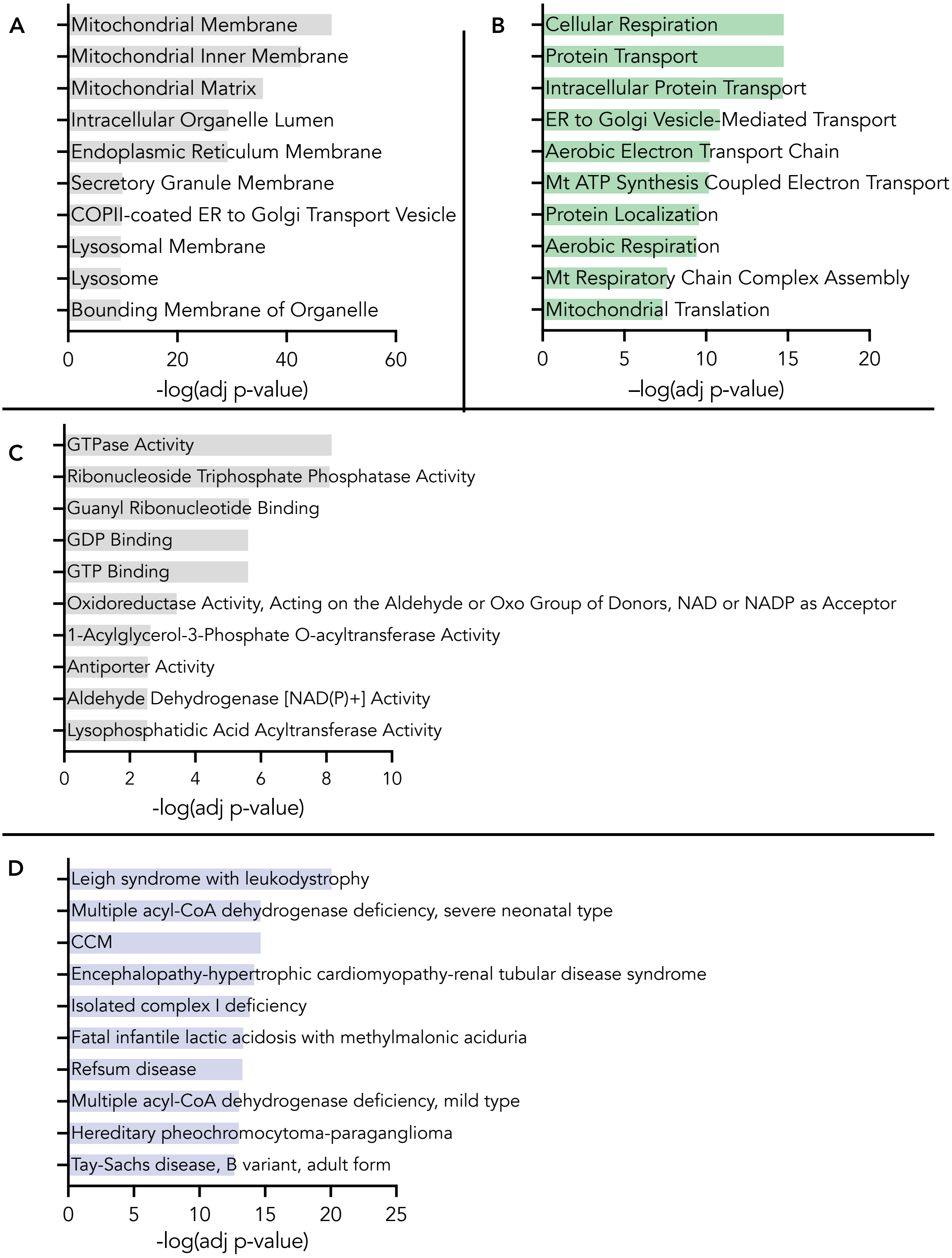


**Figure S31**. Gene Ontology analysis of enriched proteins from Enrichr of (**A**) Cellular componentes (**B**) Biological Processes. (**C**) Molecular function and (**D**) Orphanet Disease associations from Enrichr.

### 8. NMR Spectra

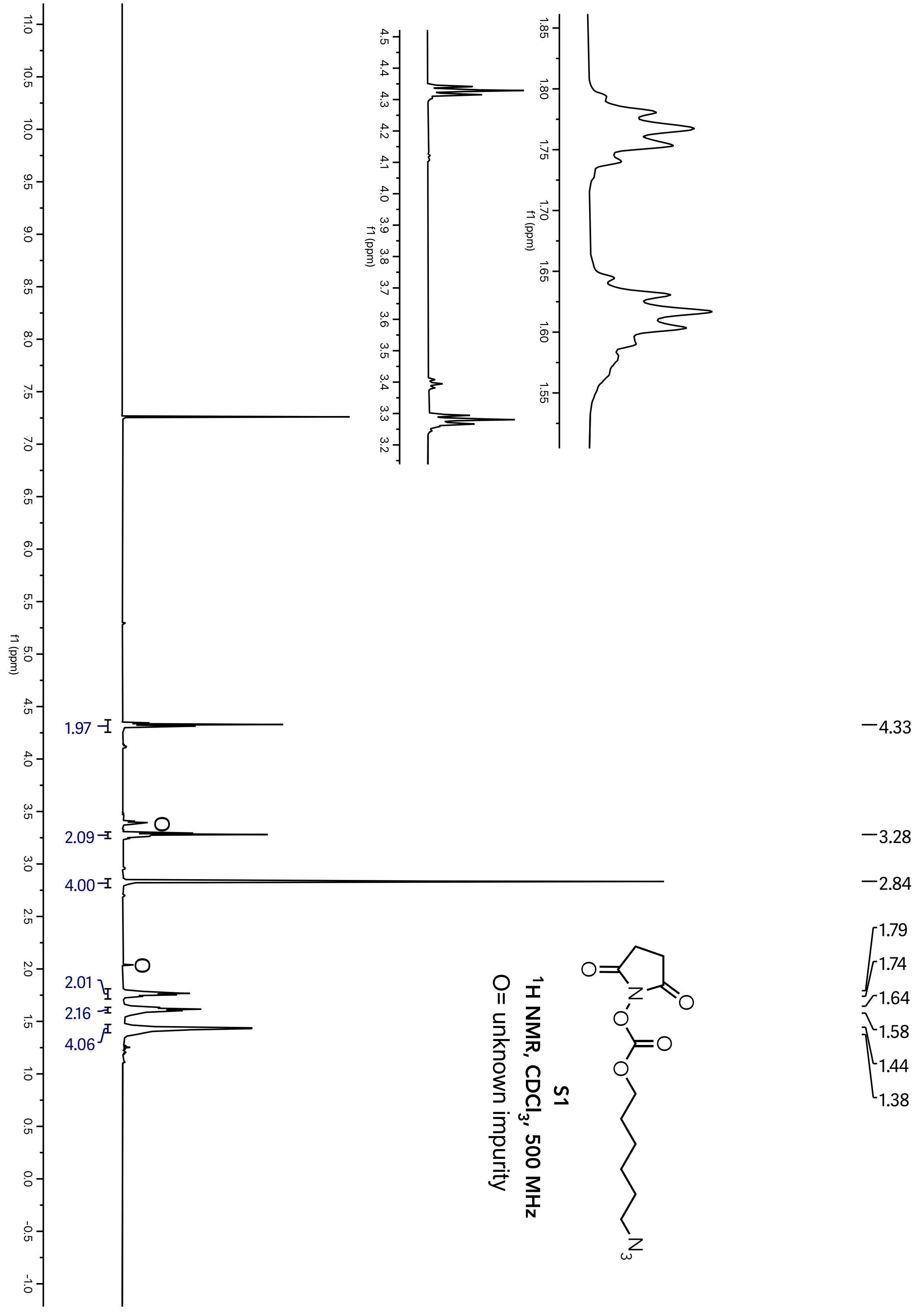

**S1**

$^{13}C$ NMR, $CDCl_3$, 126 MHz

O= unknown impurity

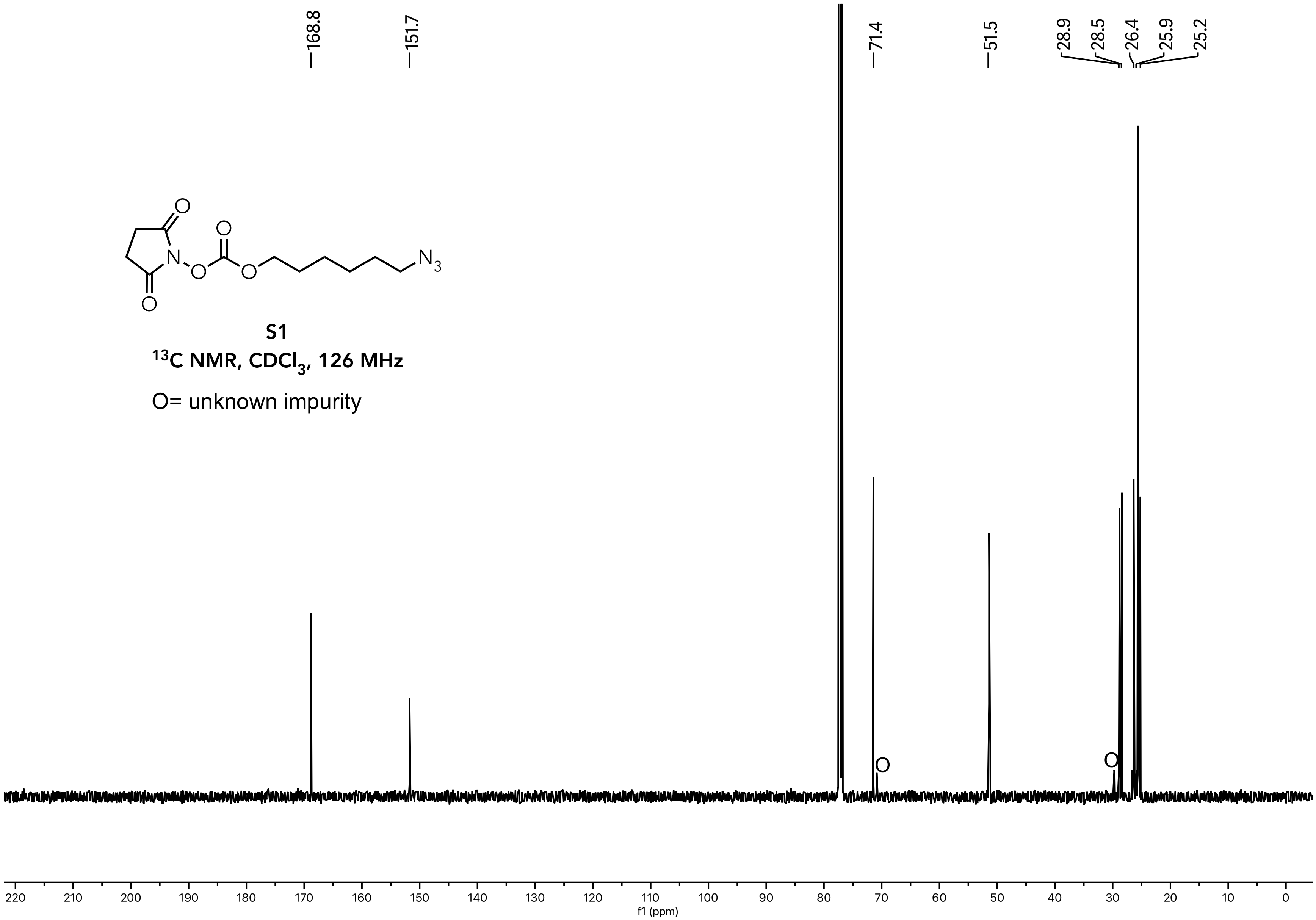

**S2**

**$^1$H NMR, $CDCl_3$, 500 MHz**

O= unknown impurity

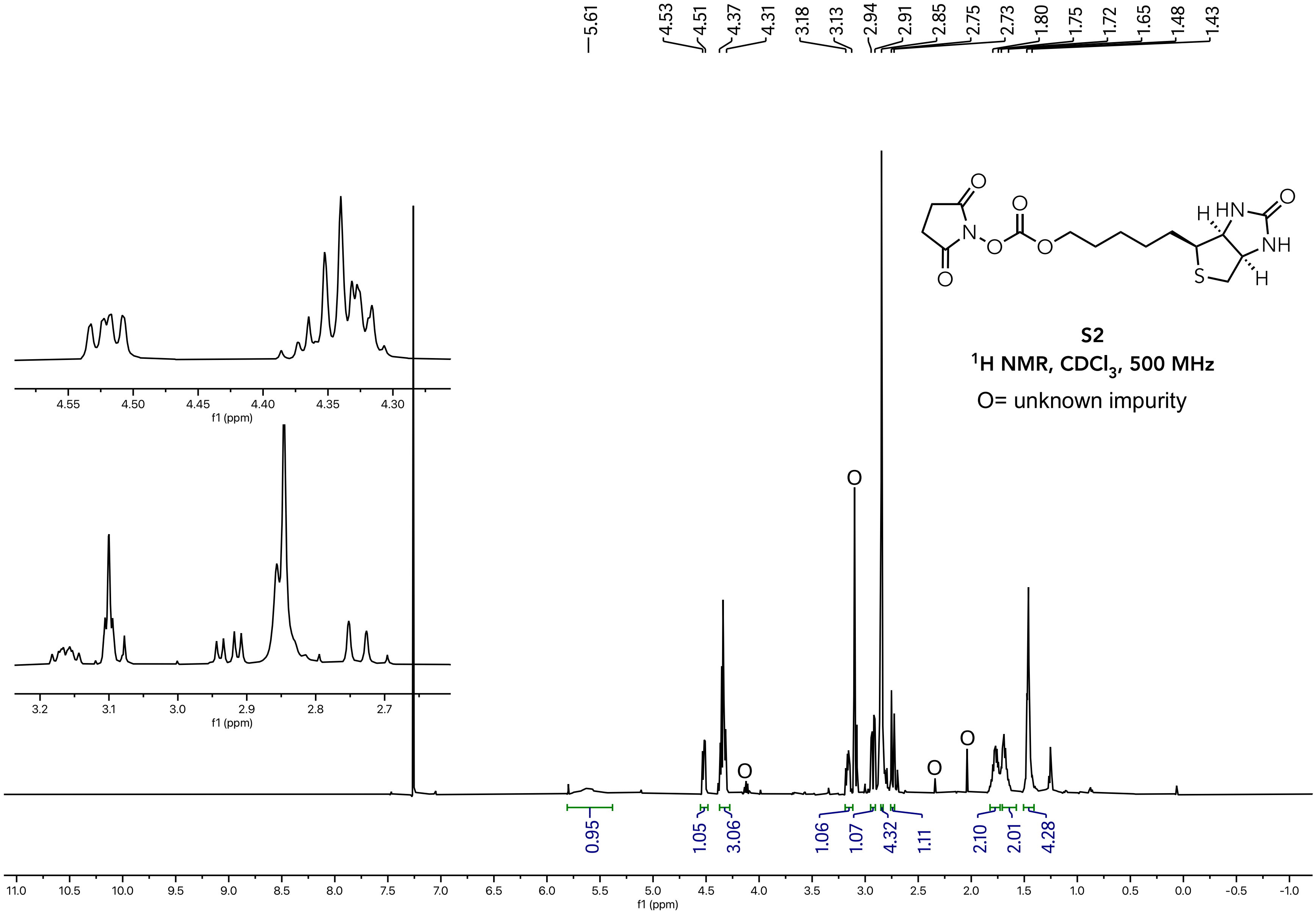

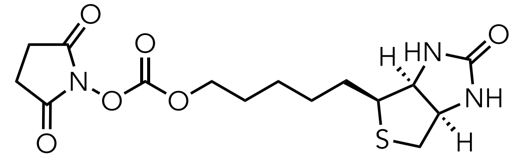

**S2**

$^{13}$C NMR, $CDCl_3$, 126 MHz

O= unknown impurity

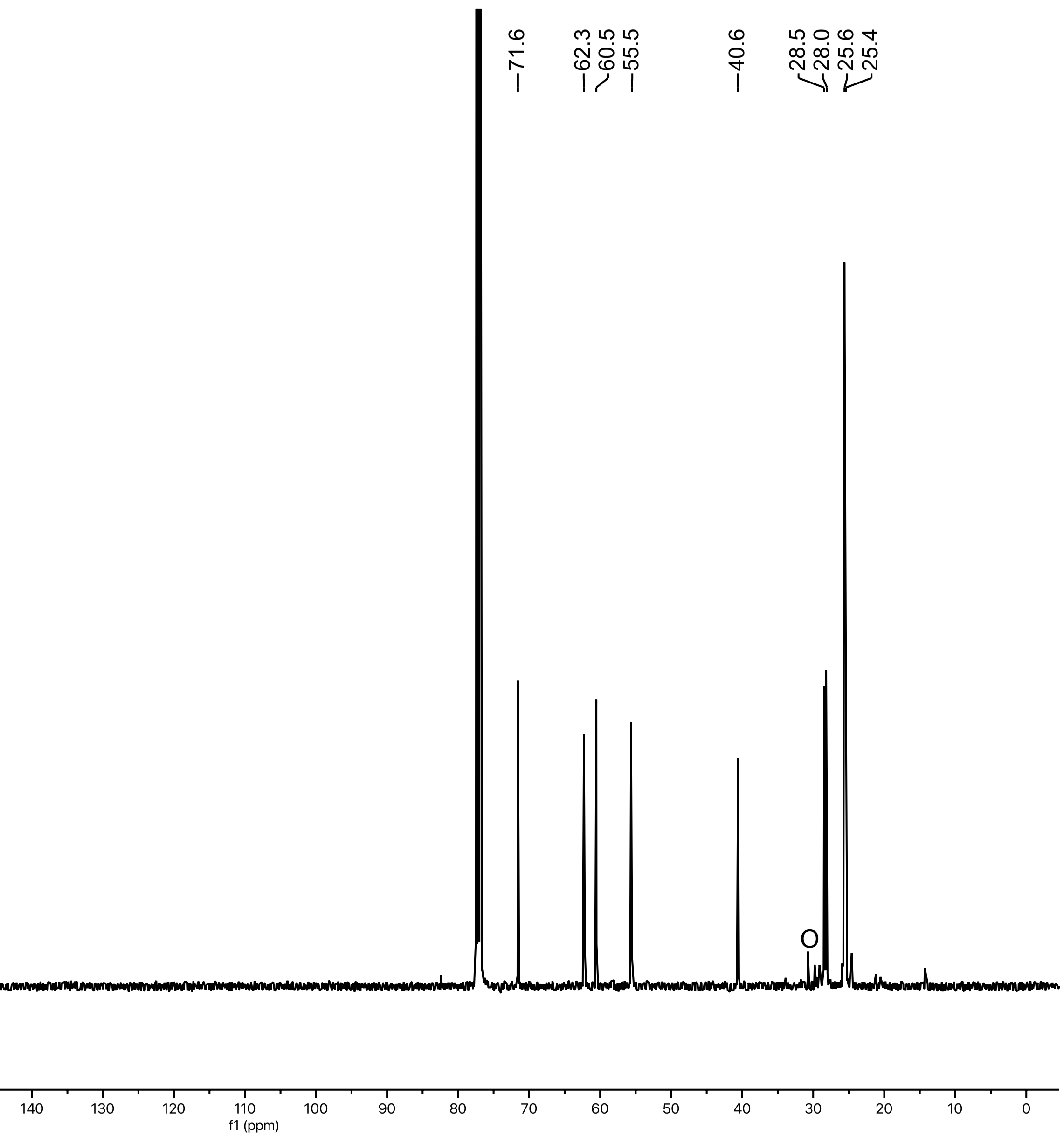

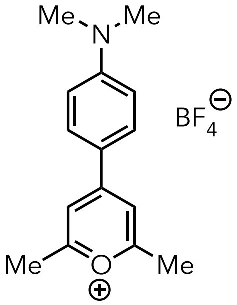


**S3**

**$^1$H NMR, $CD_3CN$, 500 MHz**

O= unknown impurity

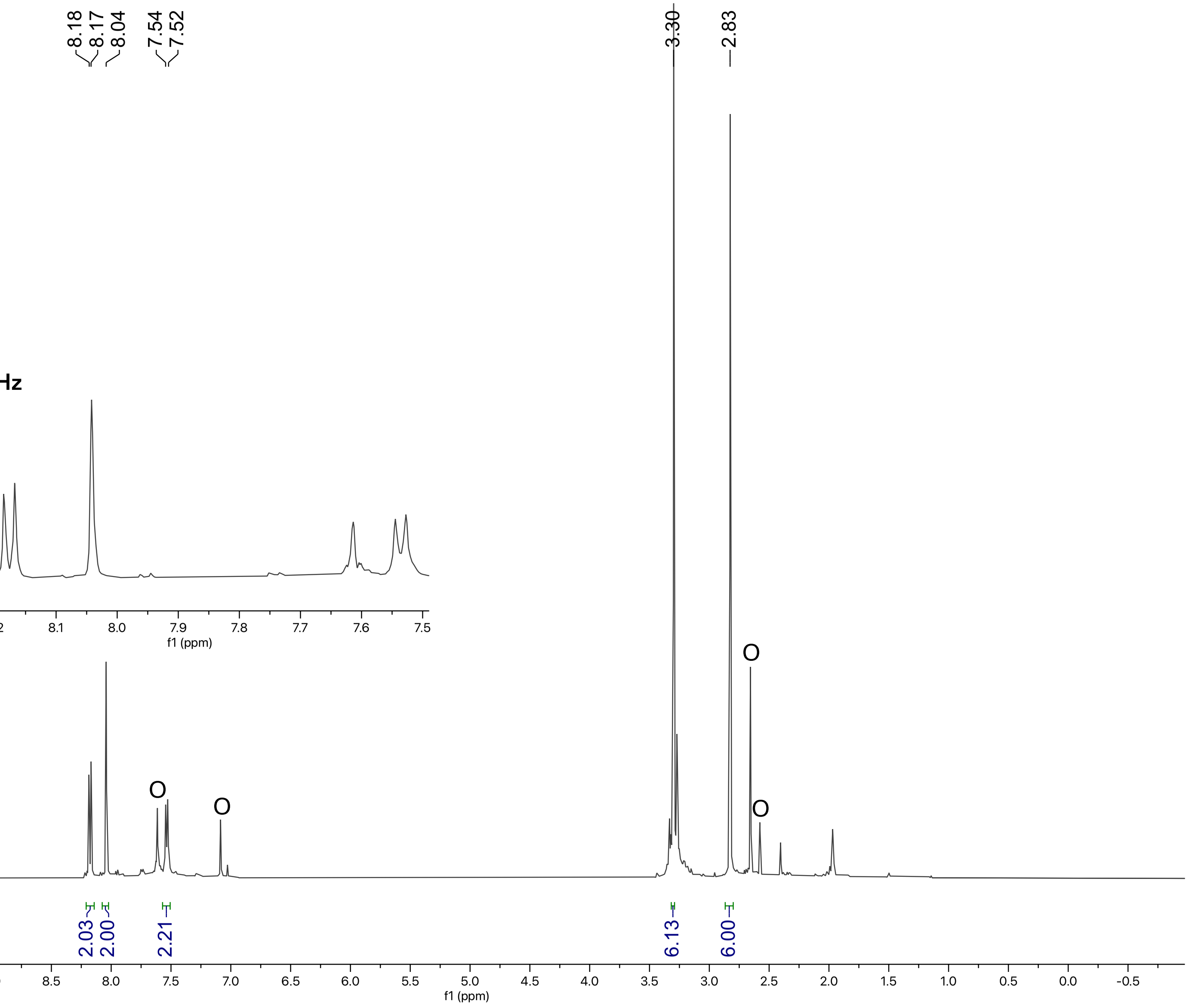

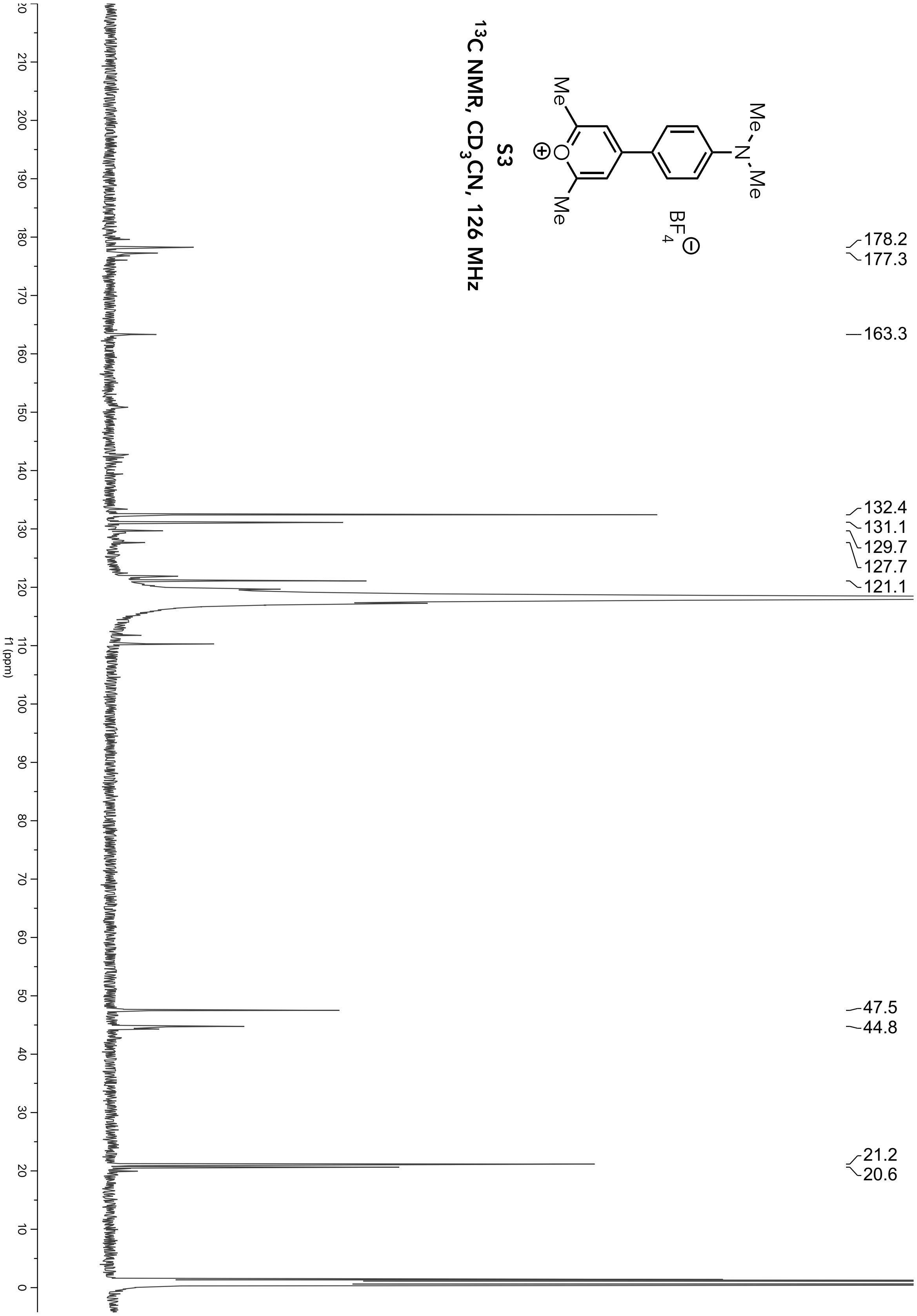

S3
13C NMR, CD3CN, 126 MHz
Me
N
Me
BF4
O
Me
Me
178.2
177.3
163.3
132.4
131.1
129.7
127.7
121.1
47.5
44.8
21.2
20.6
f1 (ppm)

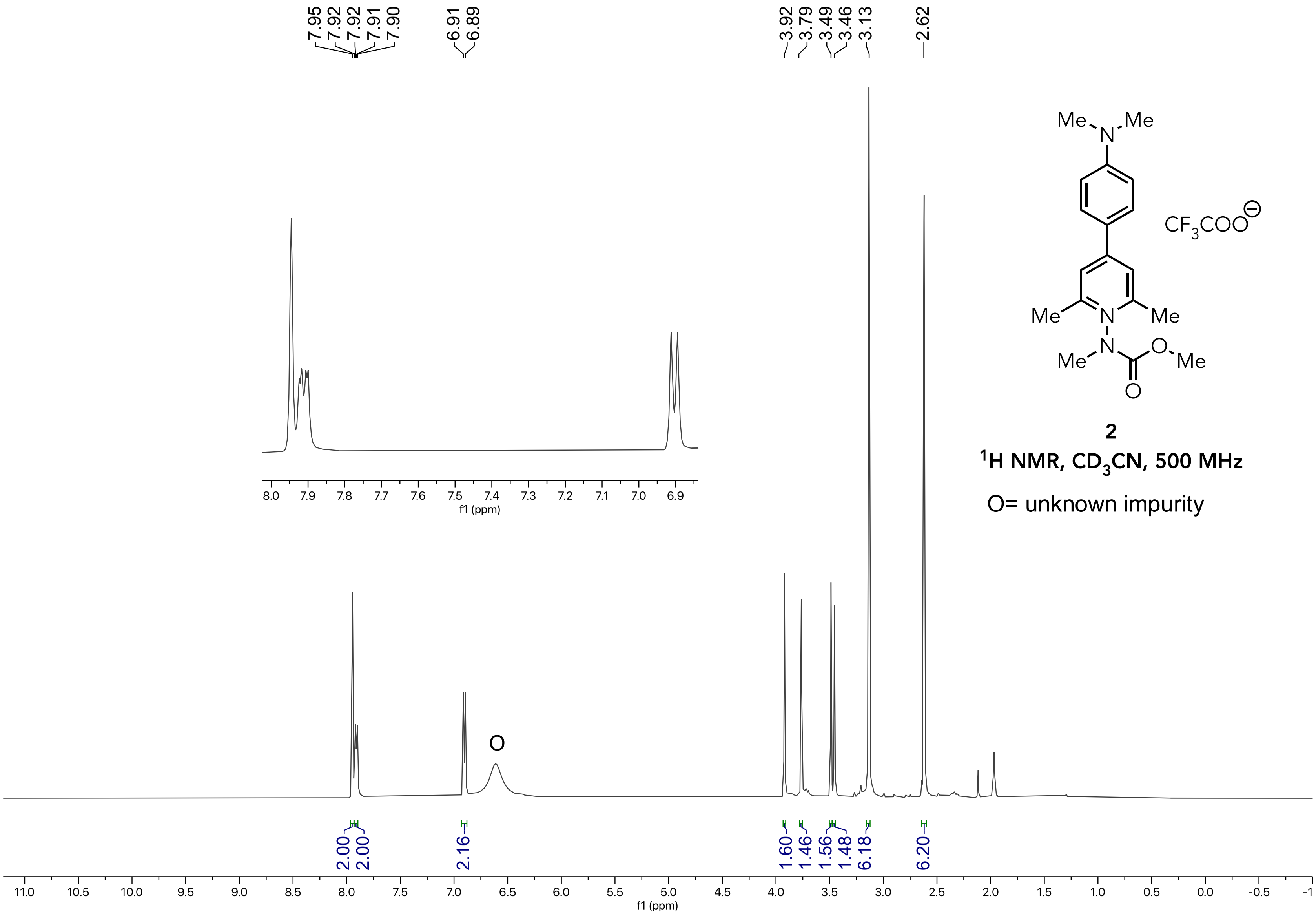


**2**

**$^1$H NMR, $CD_3CN$, 500 MHz**

O= unknown impurity

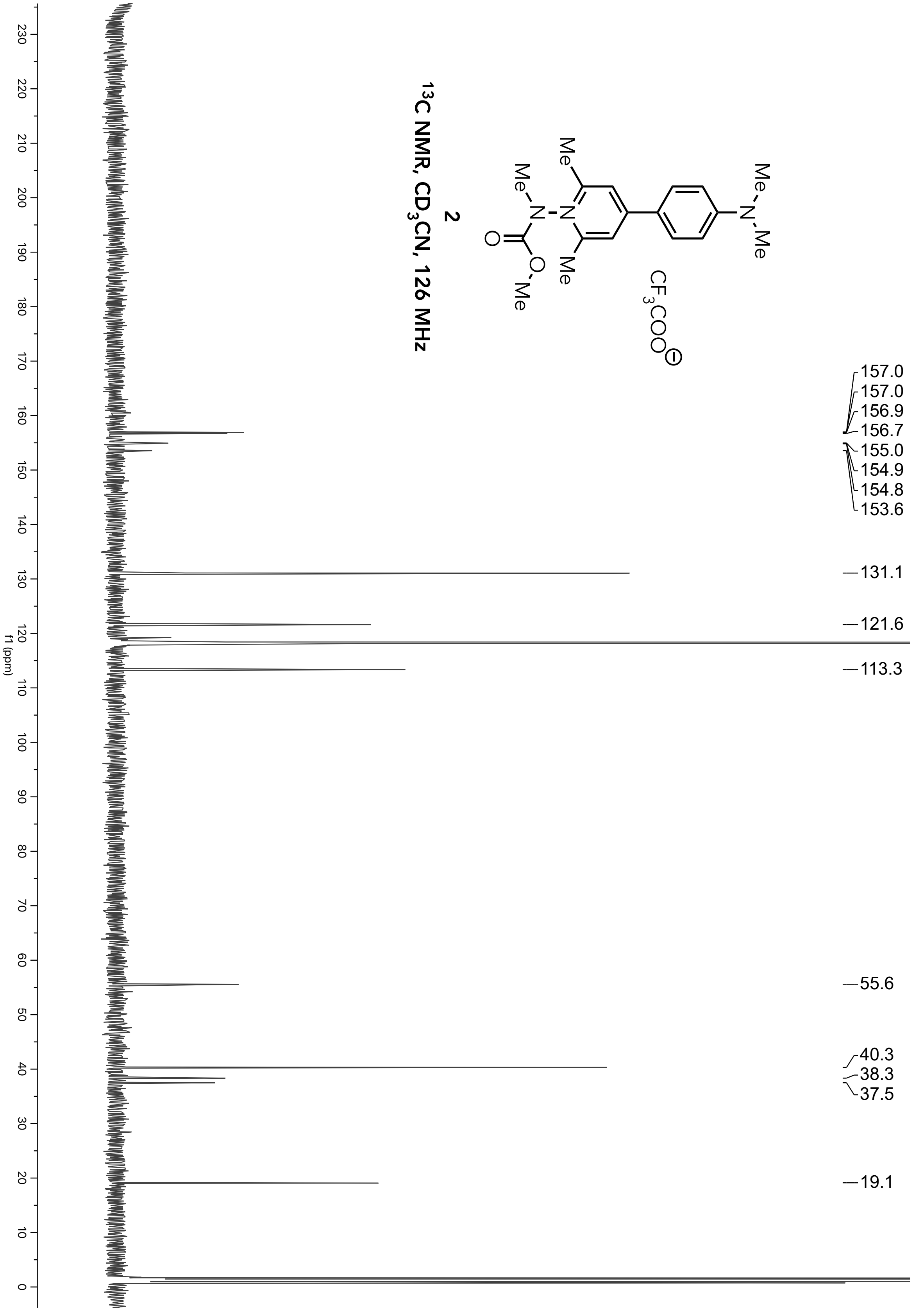
2
13C NMR, CD3CN, 126 MHz
CF3COO
157.0
157.0
156.9
156.7
155.0
154.9
154.8
153.6
131.1
121.6
113.3
55.6
40.3
38.3
37.5
19.1
f1 (ppm)

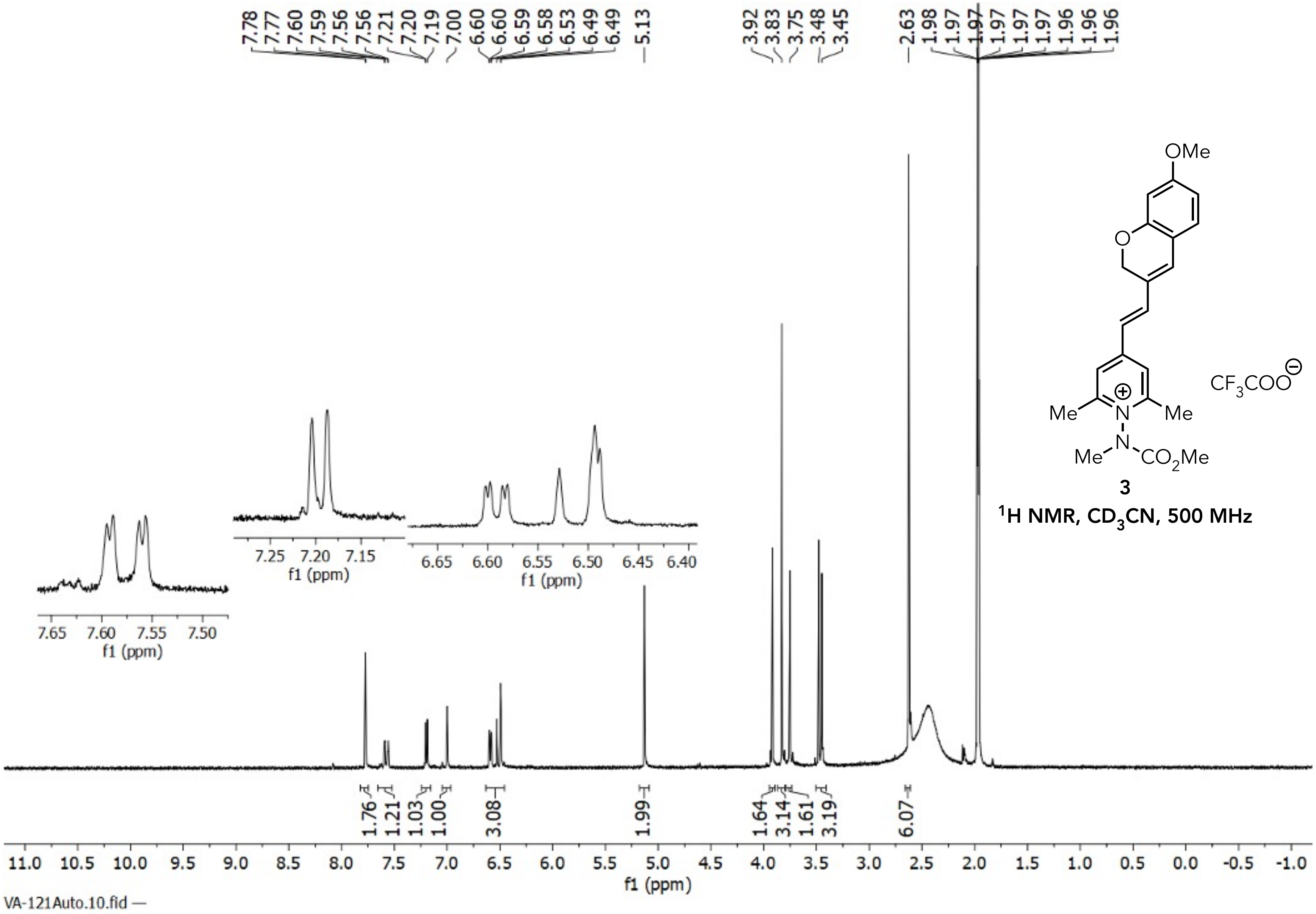


**3**

$^1$H NMR, $CD_3CN$, 500 MHz

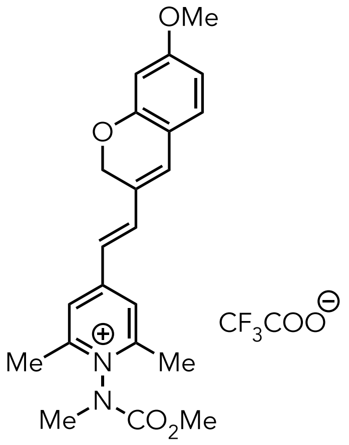


3

$^{13}C$ NMR, $CD_3CN$, 126 MHz

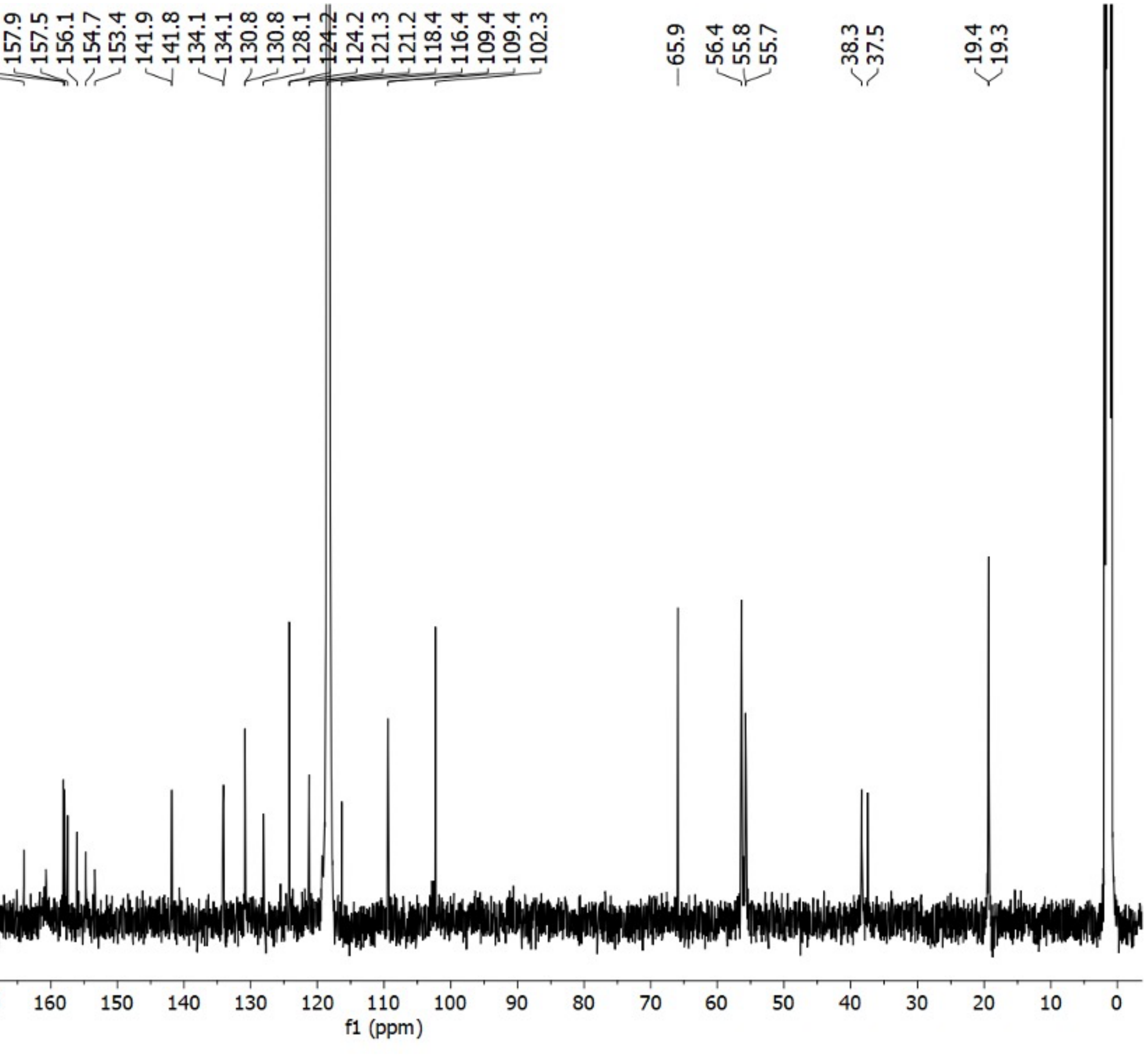

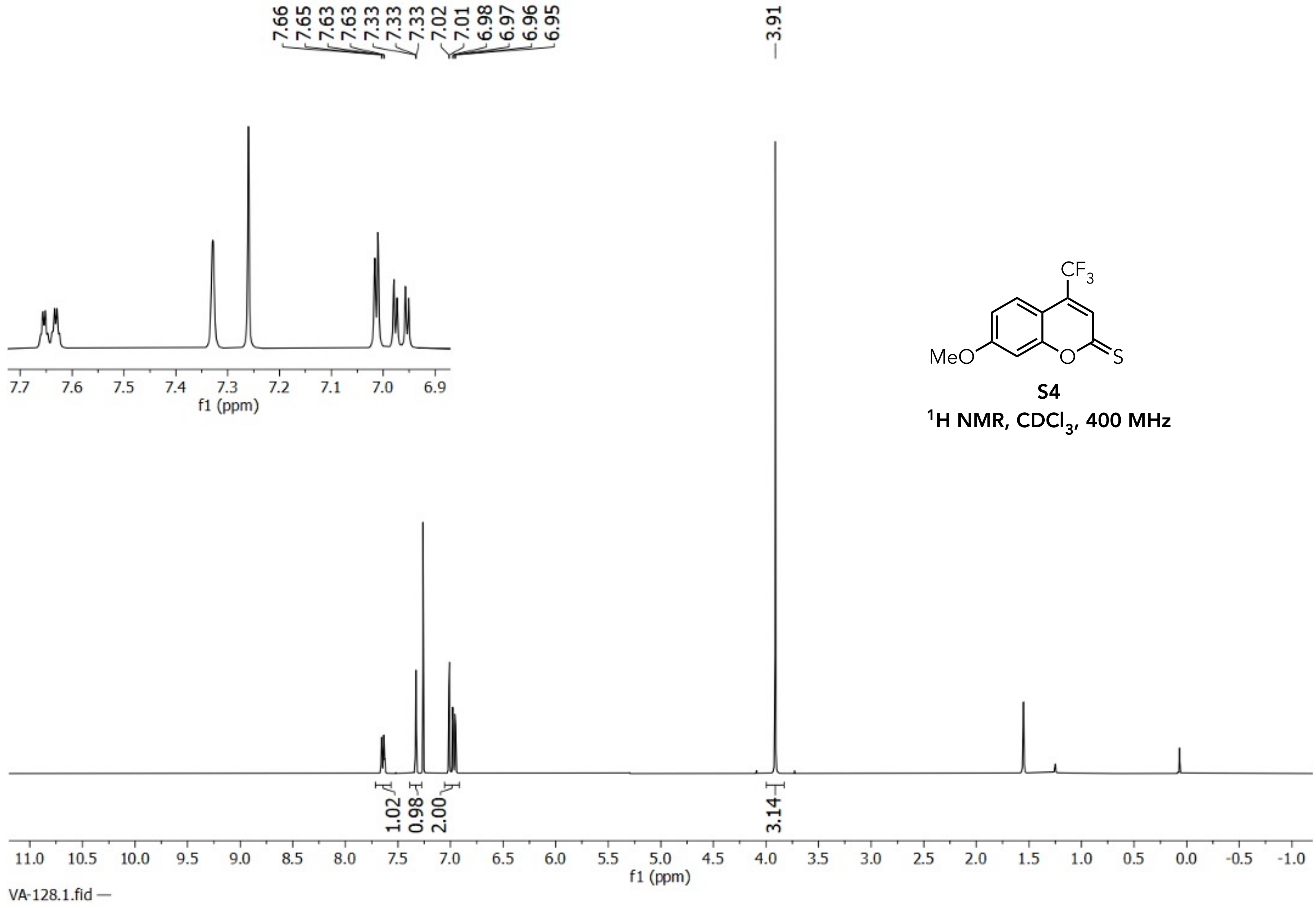


**S4**

$^1H$ NMR, $CDCl_3$, 400 MHz

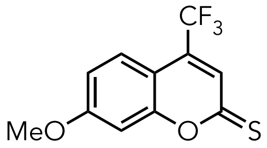


**S4**

**$^{13}C$ NMR, $CDCl_3$, 126 MHz**

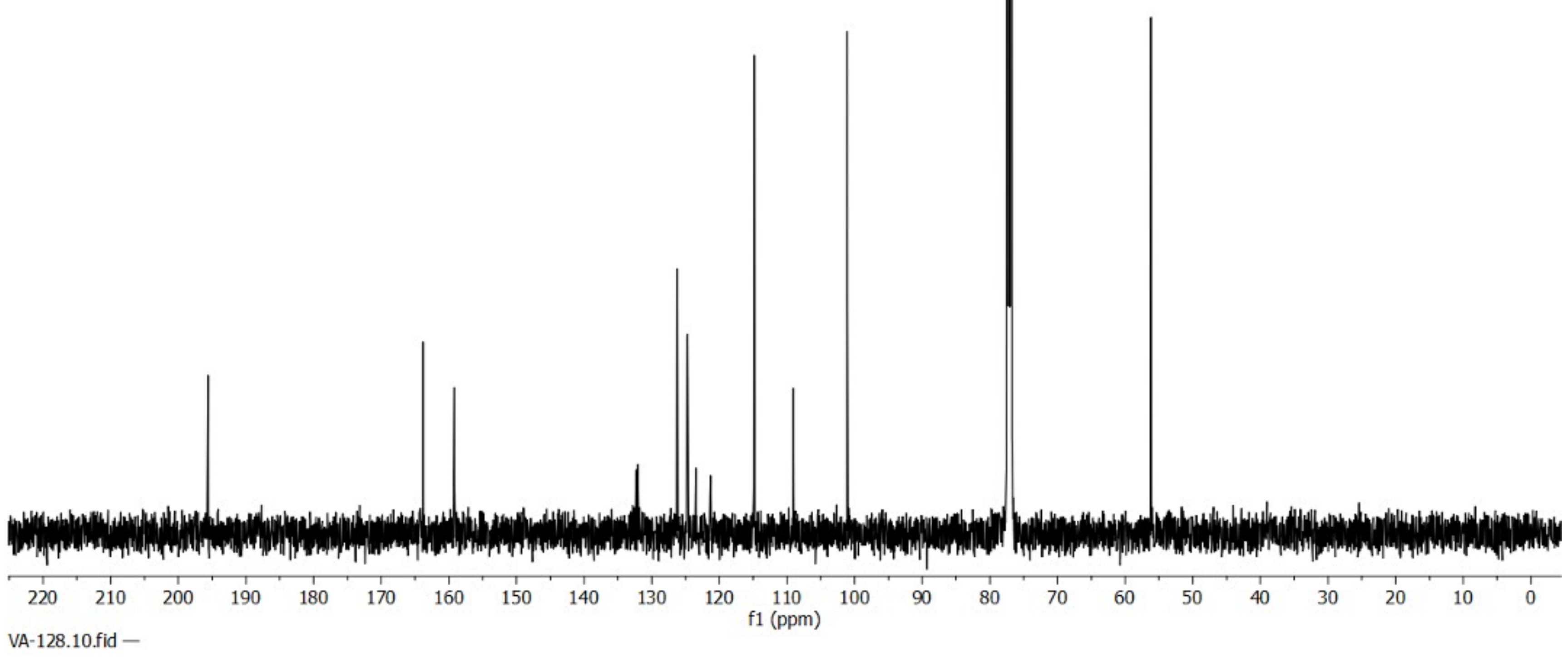

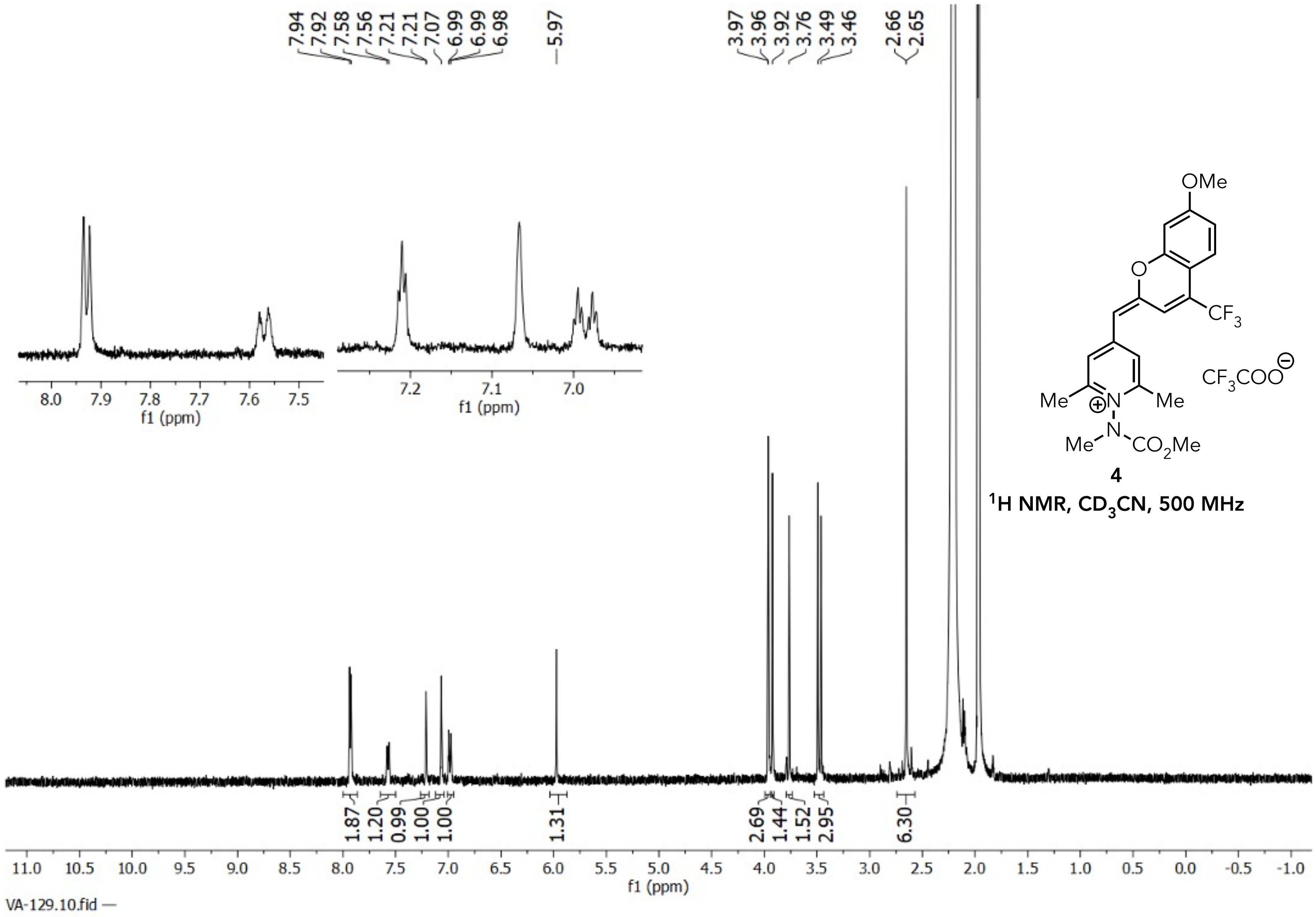


**4**

$^1$H NMR, $CD_3CN$, 500 MHz

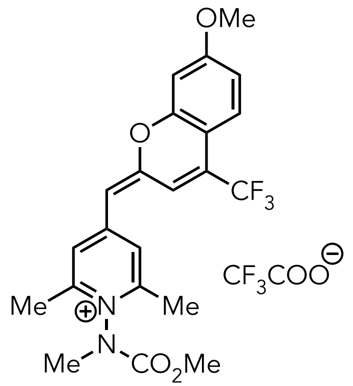


**4**

**$^{13}C$ NMR, $CD_3CN$, 126 MHz**

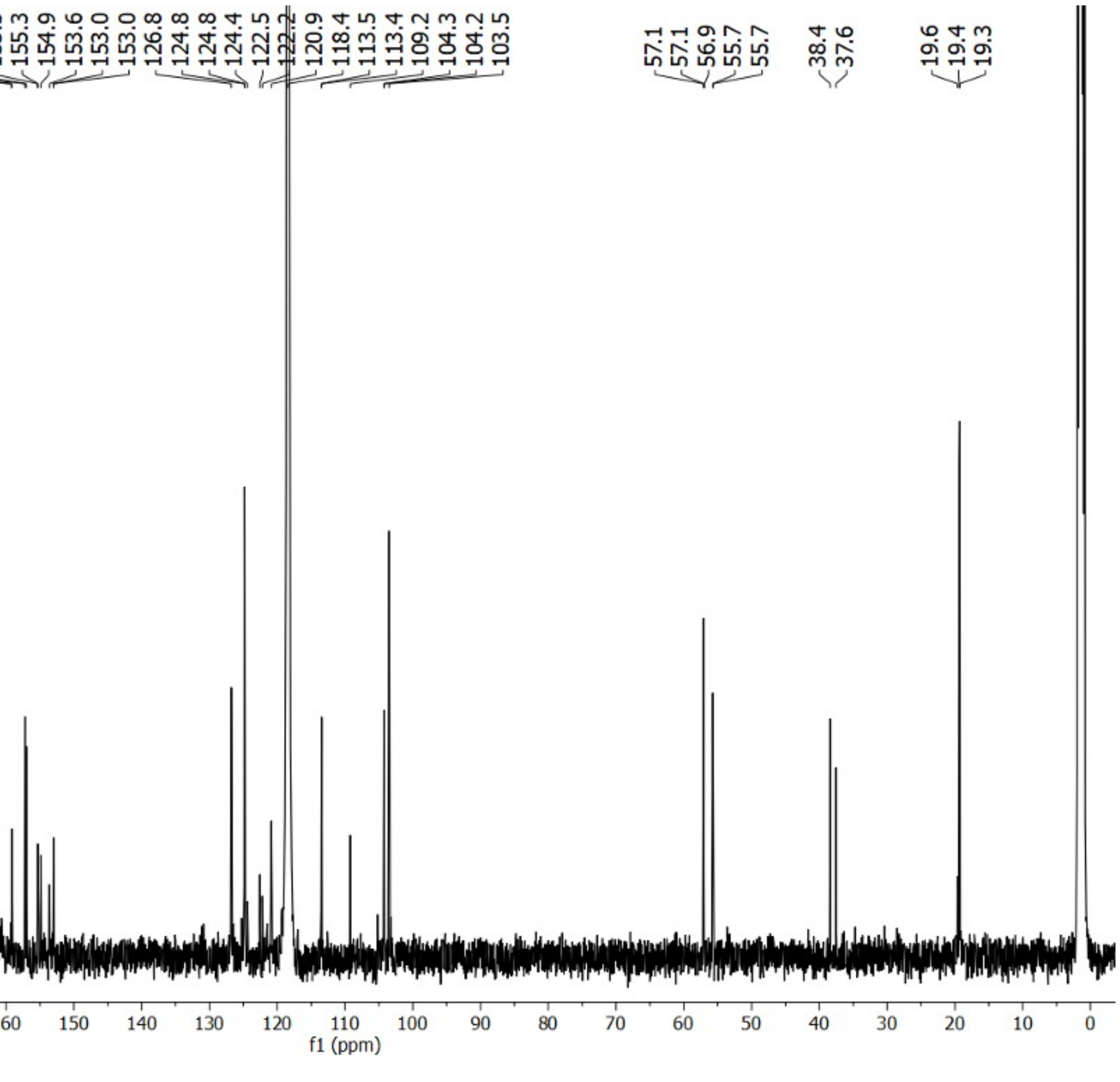

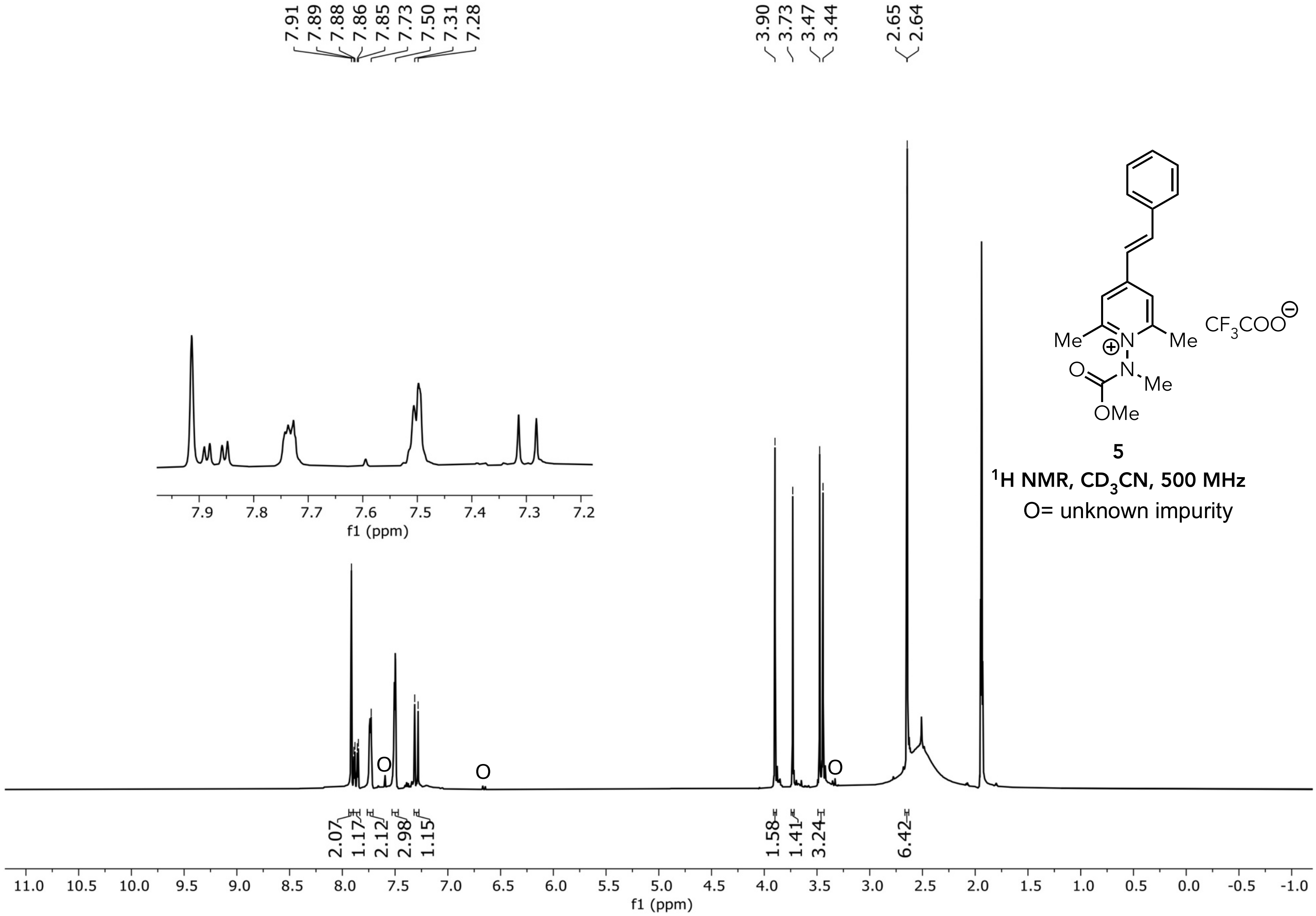


**5**

**$^1$H NMR, $CD_3CN$, 500 MHz**

O= unknown impurity

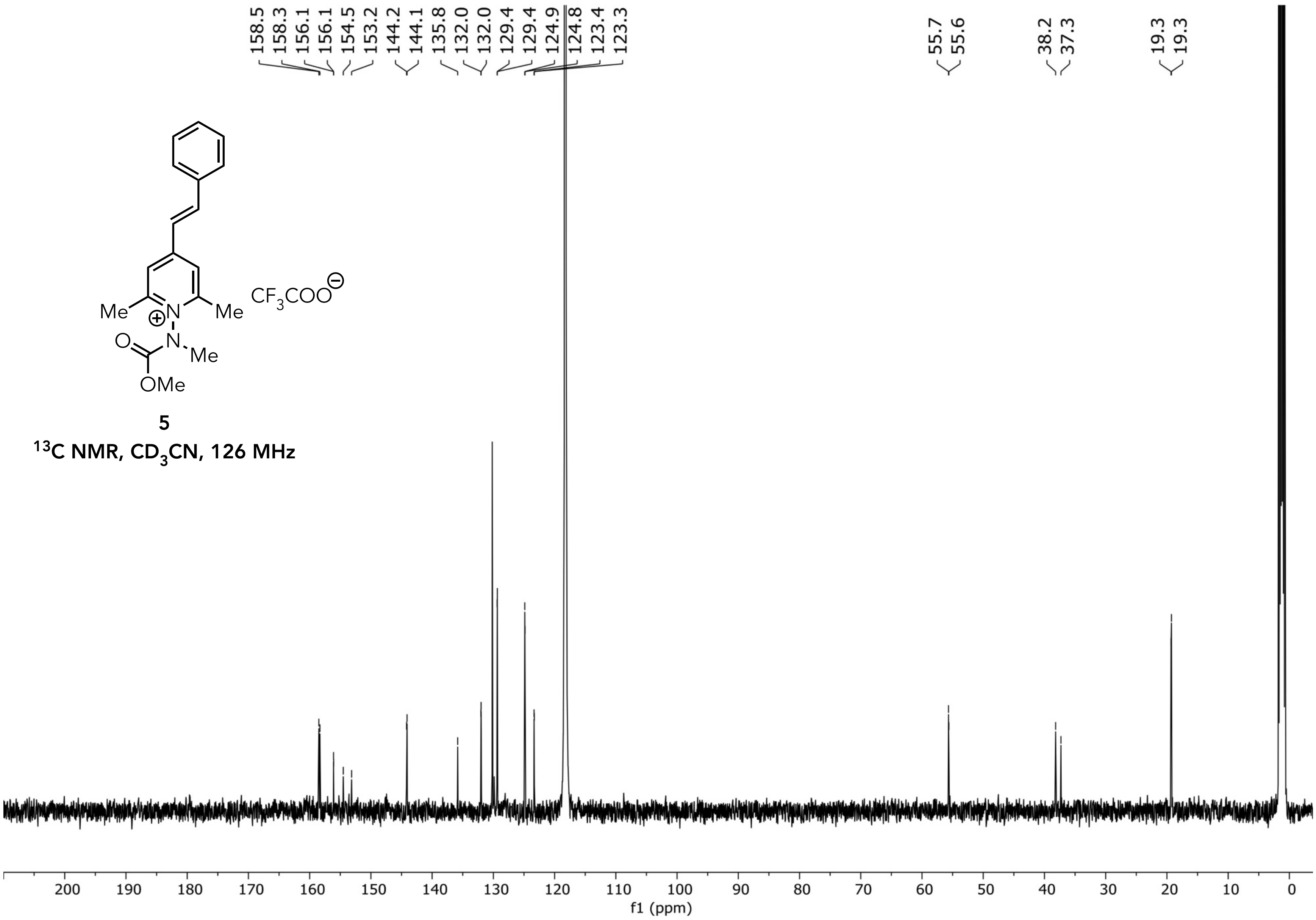


**5**

**$^{13}$C NMR, $CD_3CN$, 126 MHz**

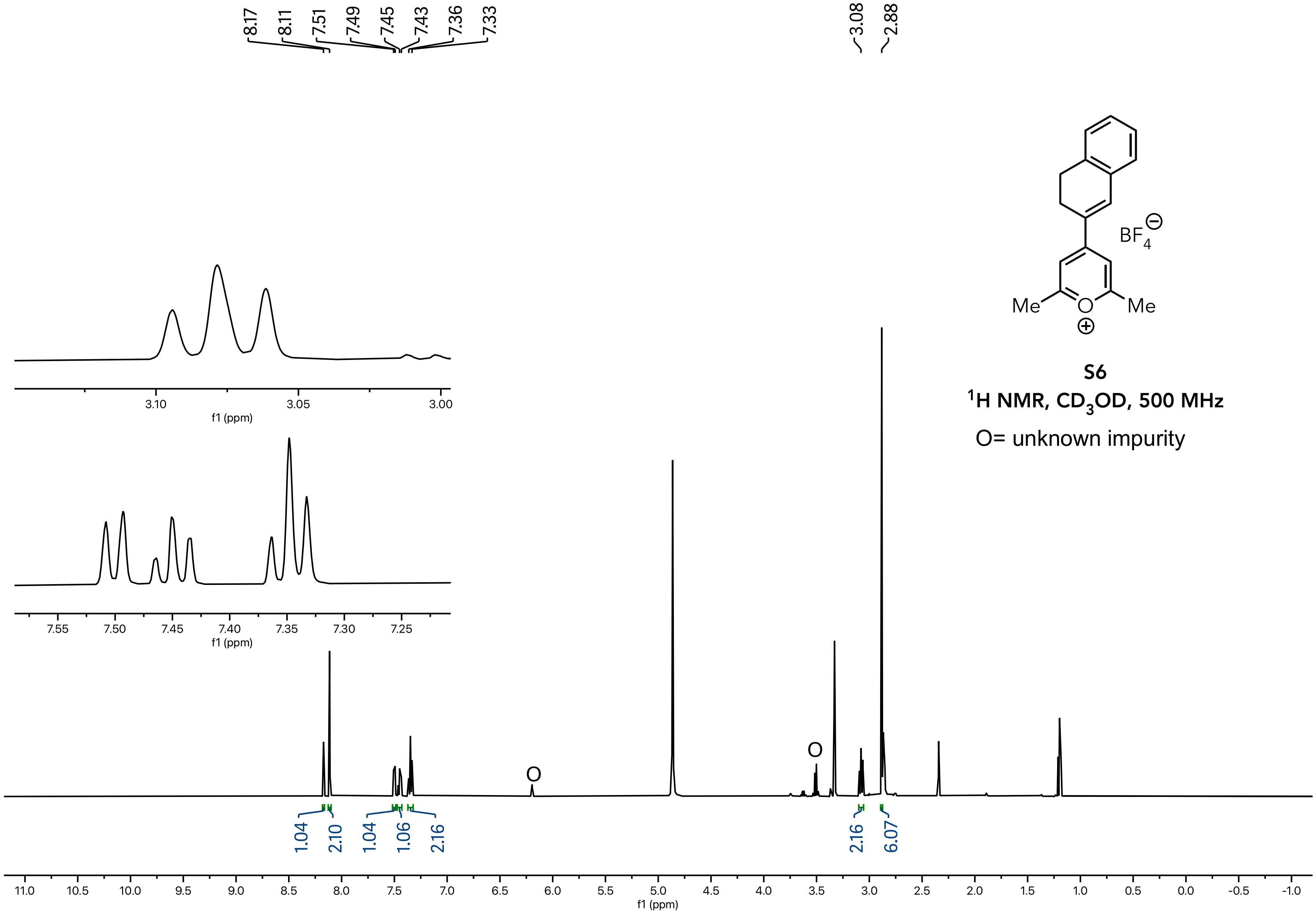


**S6**

**$^1$H NMR, $CD_3OD$, 500 MHz**

O= unknown impurity

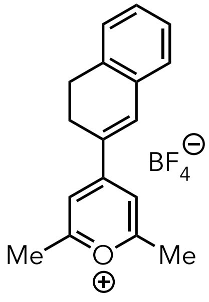


**S6**

**$^{13}$C NMR, $CD_3OD$, 126 MHz**

O= unknown impurity

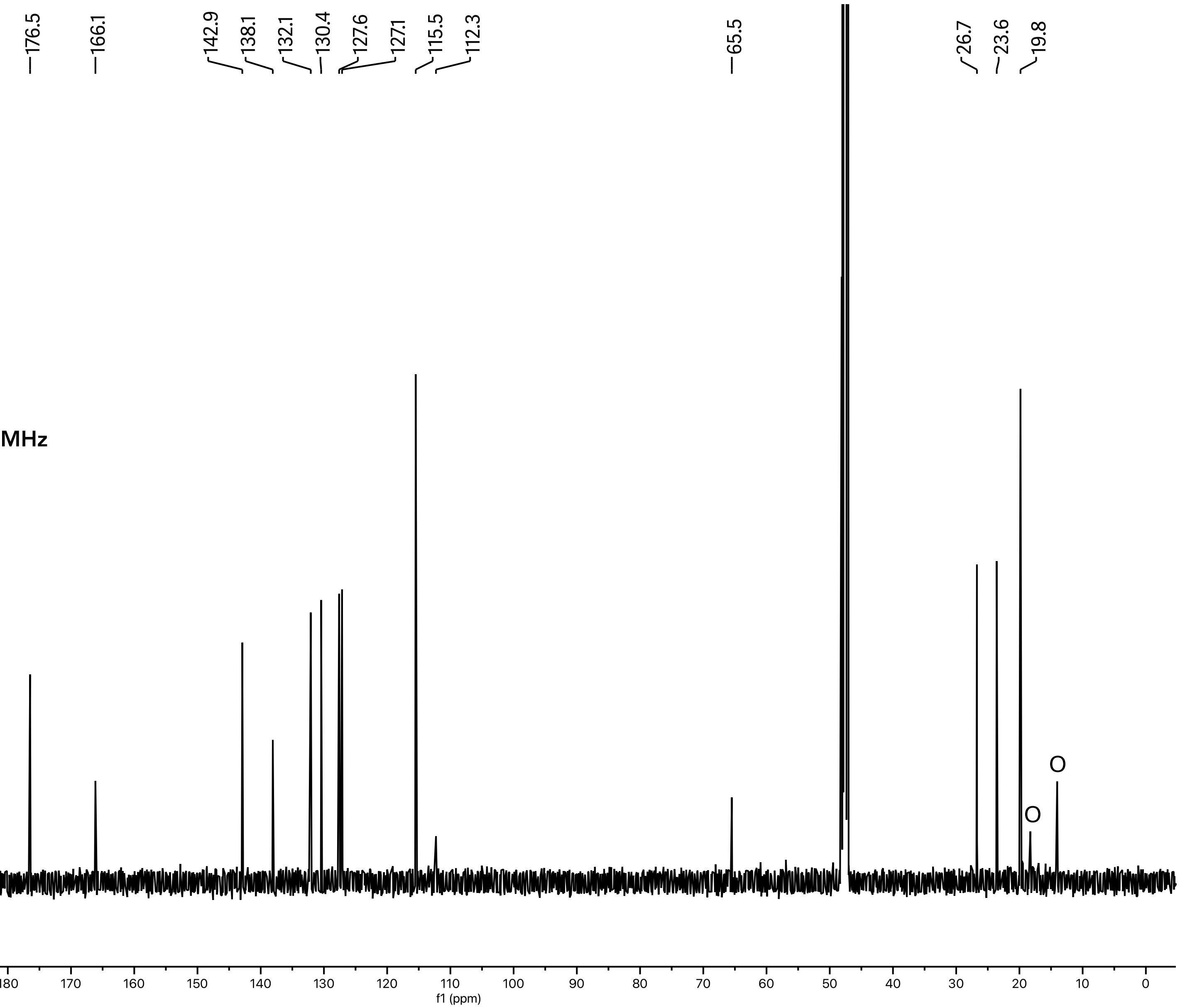

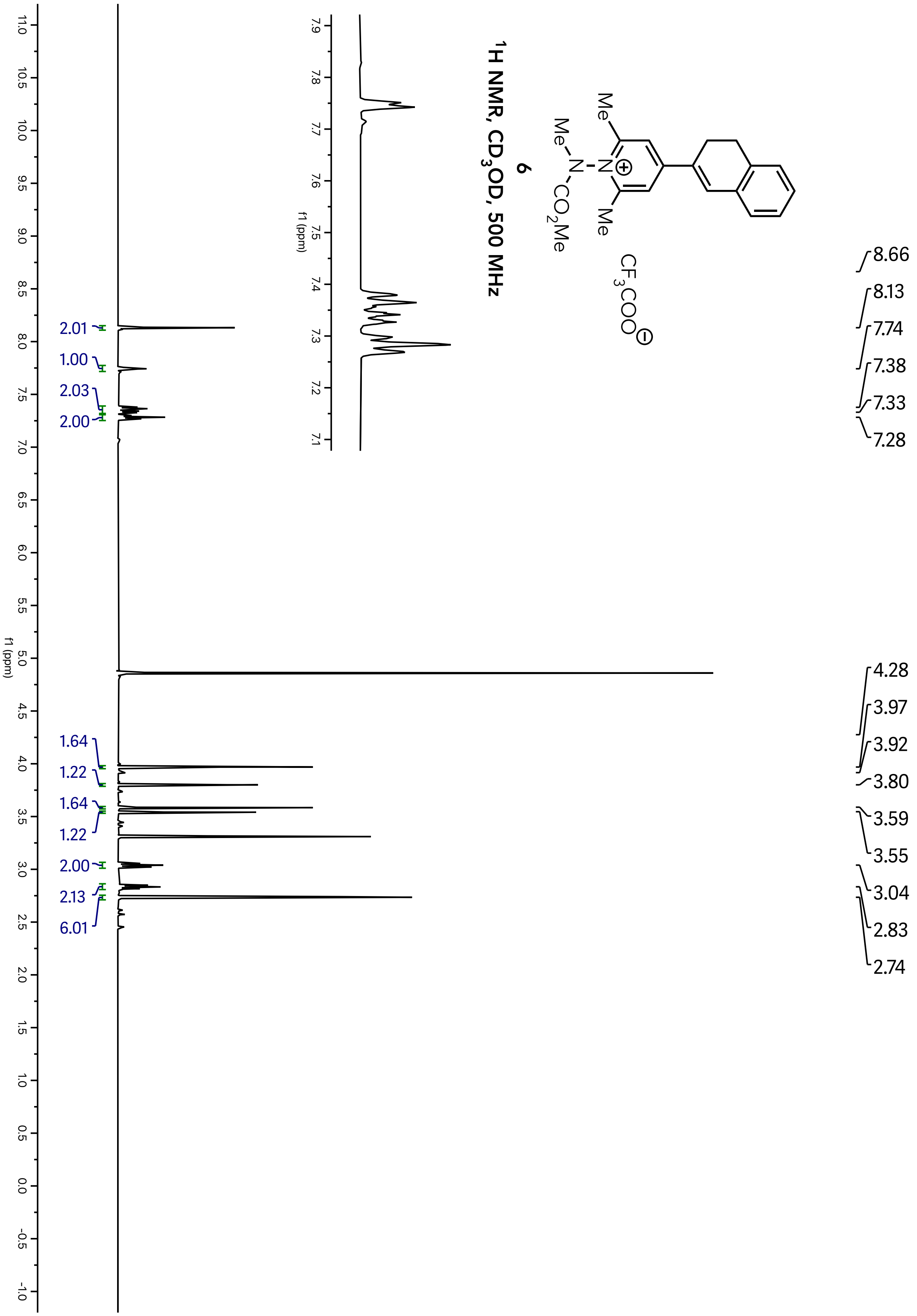

6
1H NMR, CD3OD, 500 MHz
CF3COO
8.66
8.13
7.74
7.38
7.33
7.28
4.28
3.97
3.92
3.80
3.59
3.55
3.04
2.83
2.74
2.01
1.00
2.03
2.00
1.64
1.22
1.64
1.22
2.00
2.13
6.01
f1 (ppm)

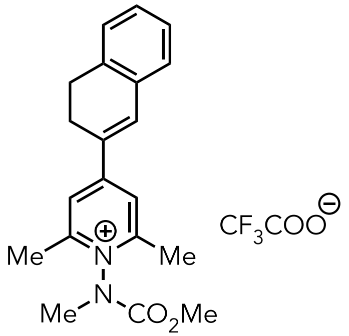


**6**

**$^{13}$C NMR, $CD_3OD$, 126 MHz**

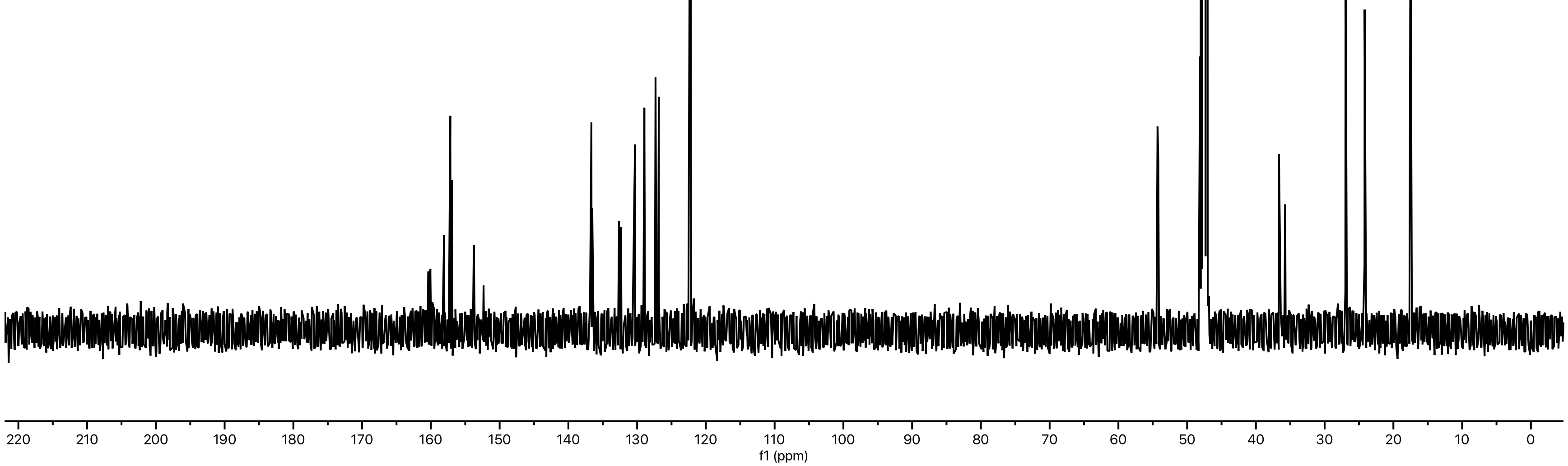

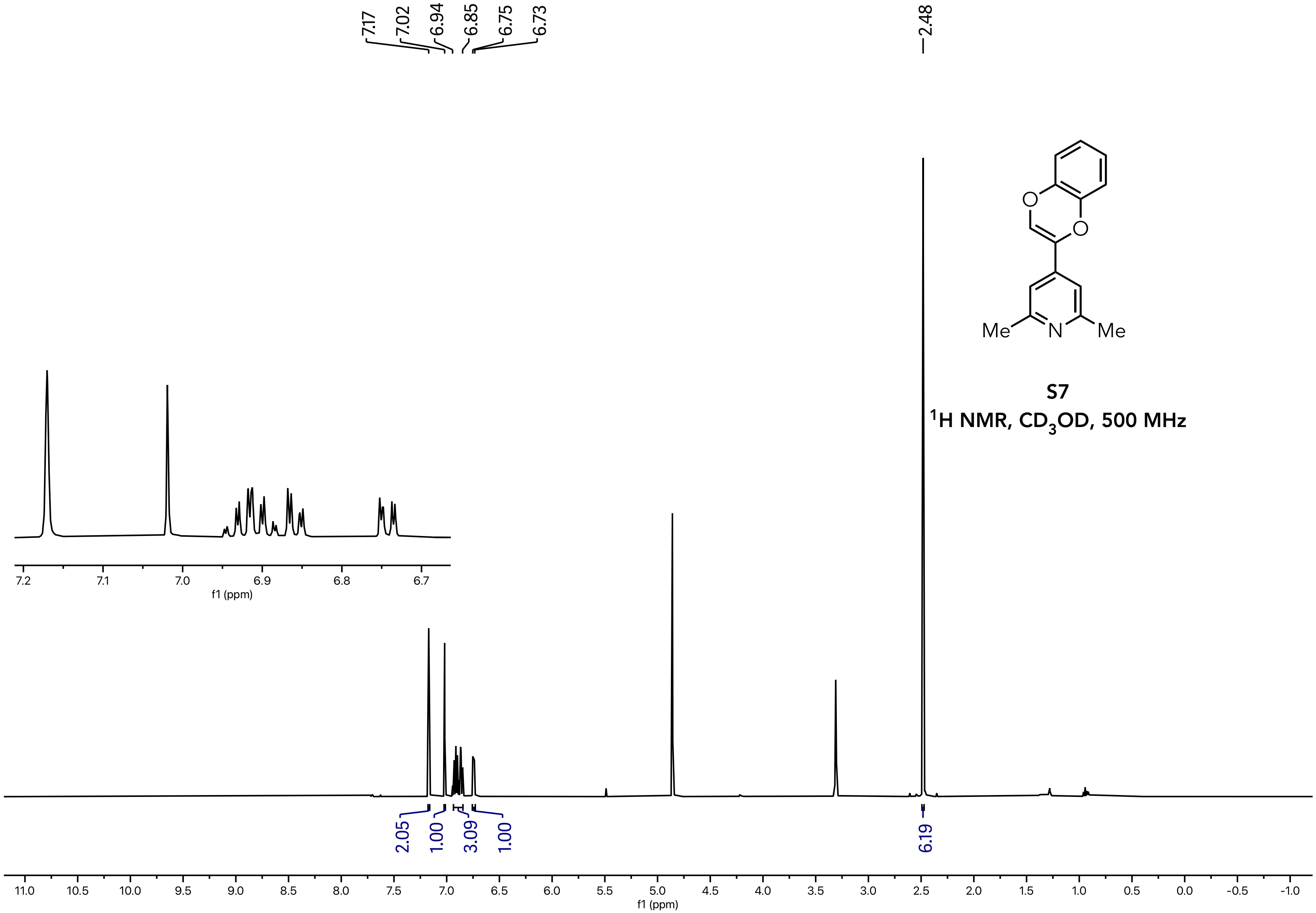


**S7**

**$^1$H NMR, $CD_3OD$, 500 MHz**

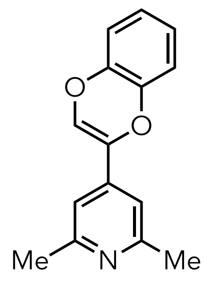


**S7**

$^{13}C$ NMR, $CD_3OD$, 126 MHz

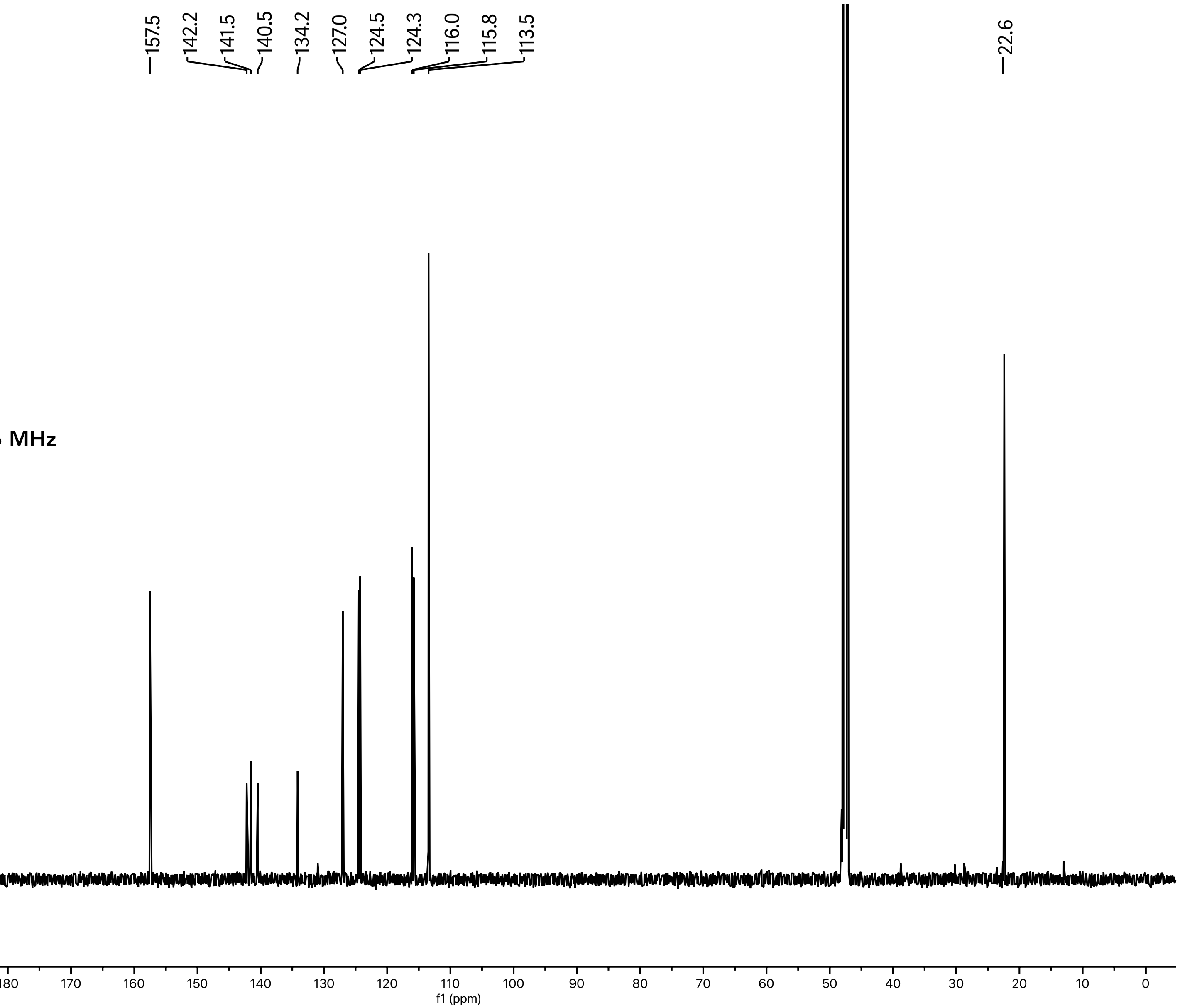

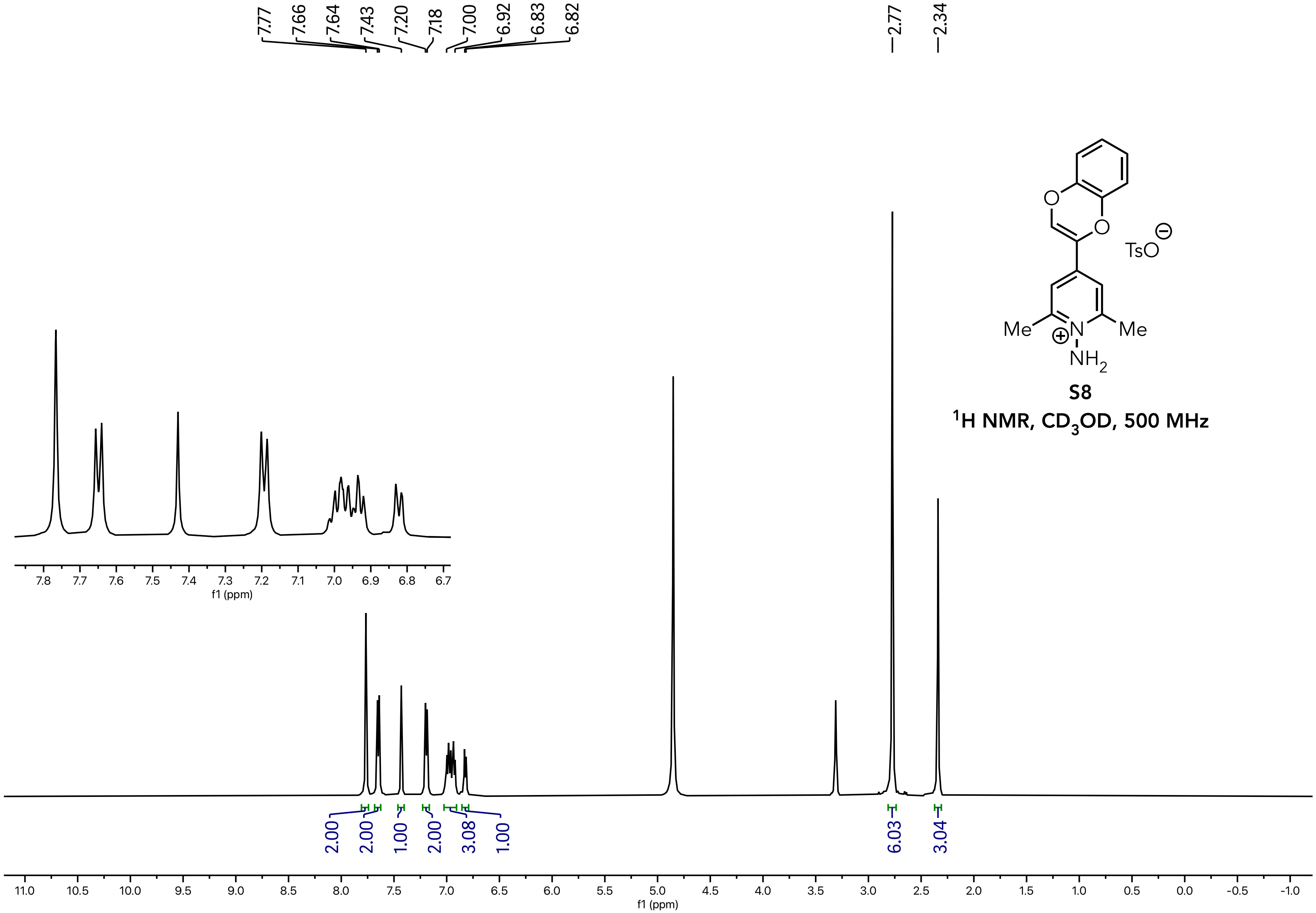


**S8**

**$^1$H NMR, $CD_3OD$, 500 MHz**

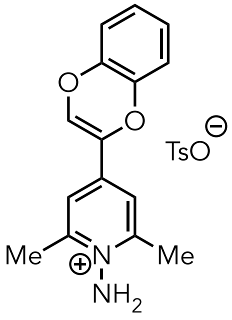


**S8**

**$^{13}C$ NMR, $CD_3OD$, 126 MHz**

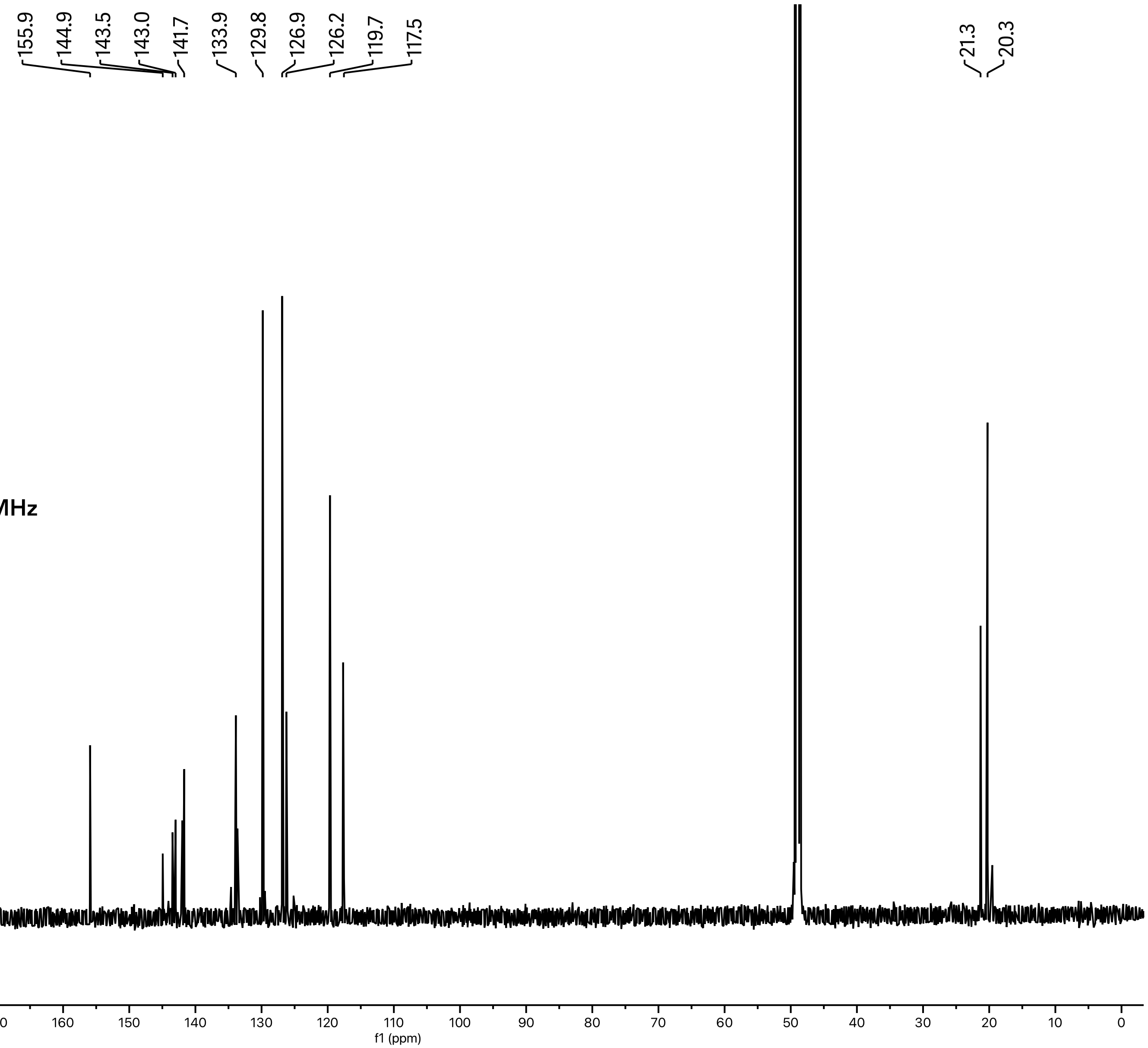

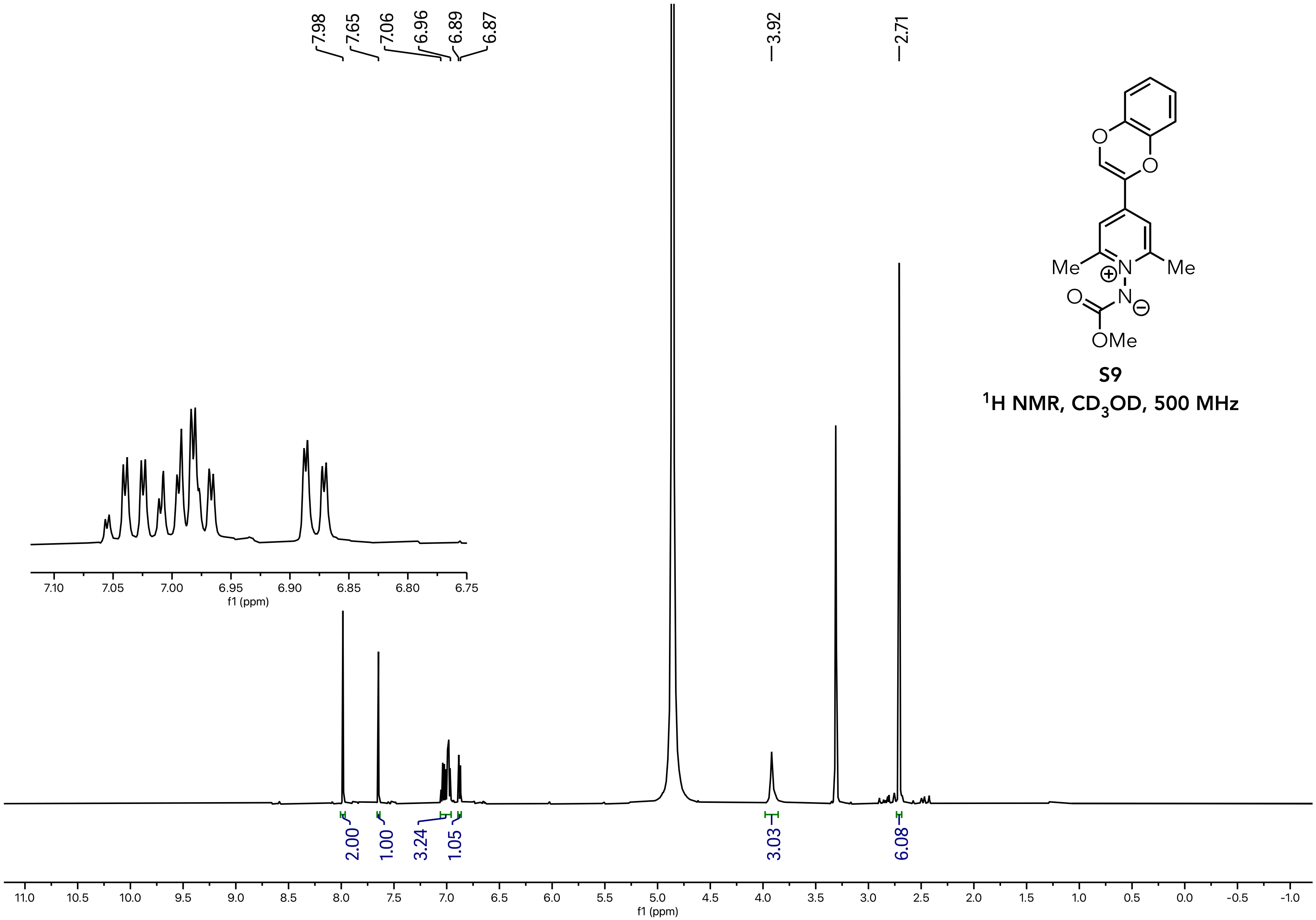


**S9**

**$^1$H NMR, $CD_3OD$, 500 MHz**

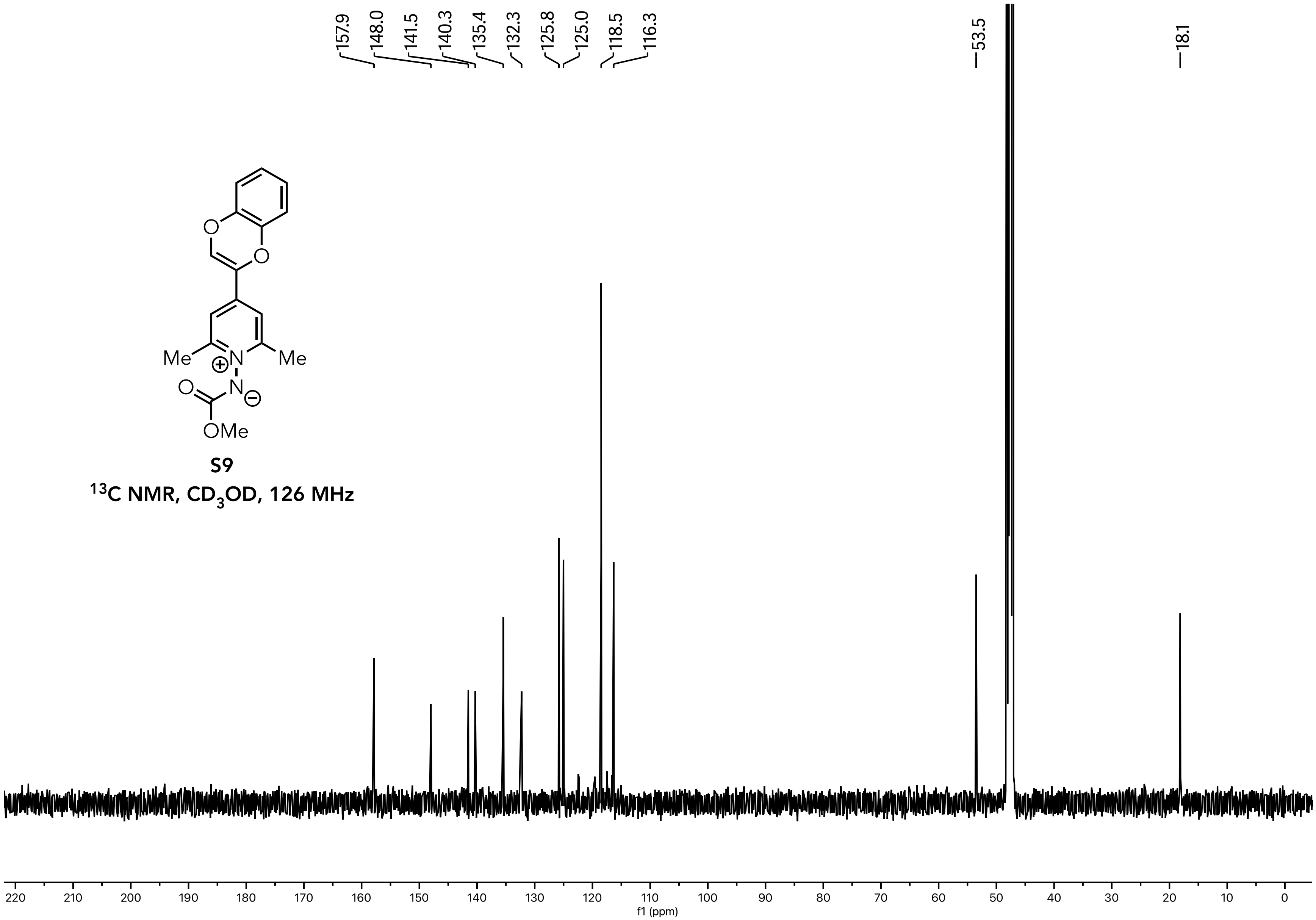


**S9**

$^{13}C$ NMR, $CD_3OD$, 126 MHz

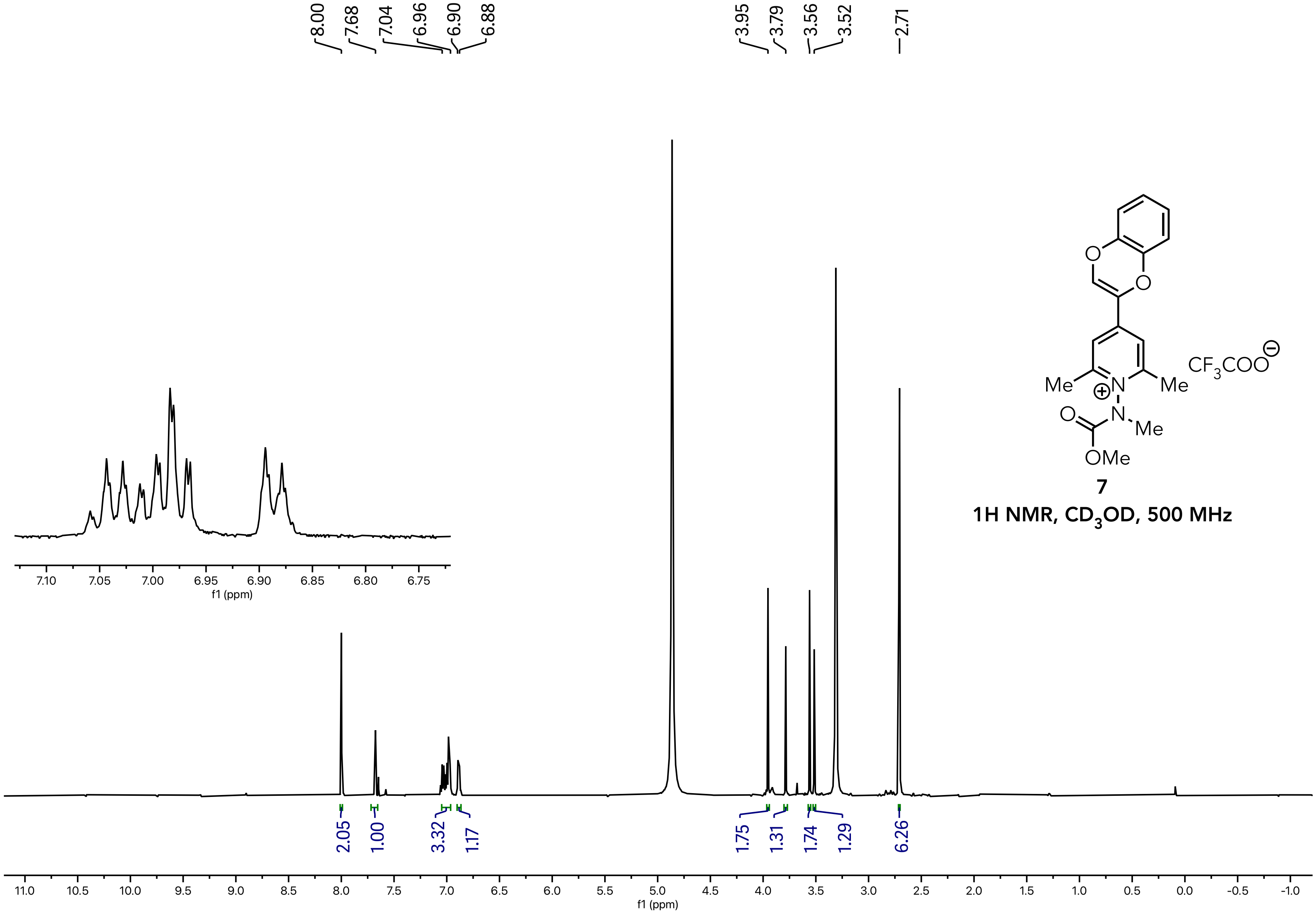


**7**

**1H NMR, $CD_3OD$, 500 MHz**

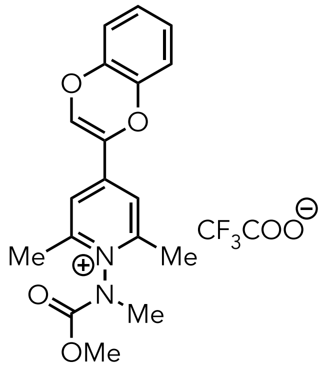


**7**

**$^{13}$C NMR, $CD_3OD$, 126 MHz**

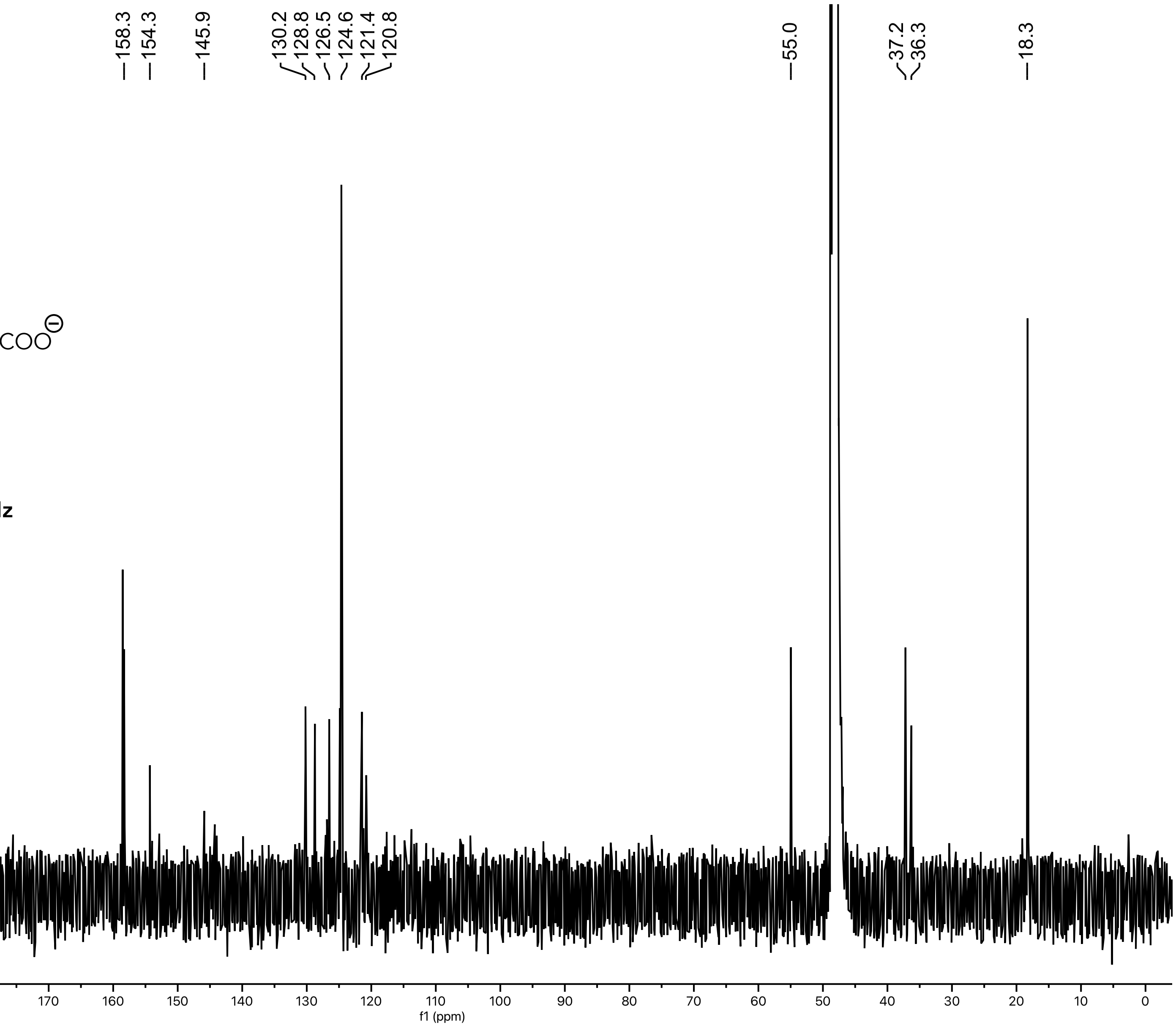

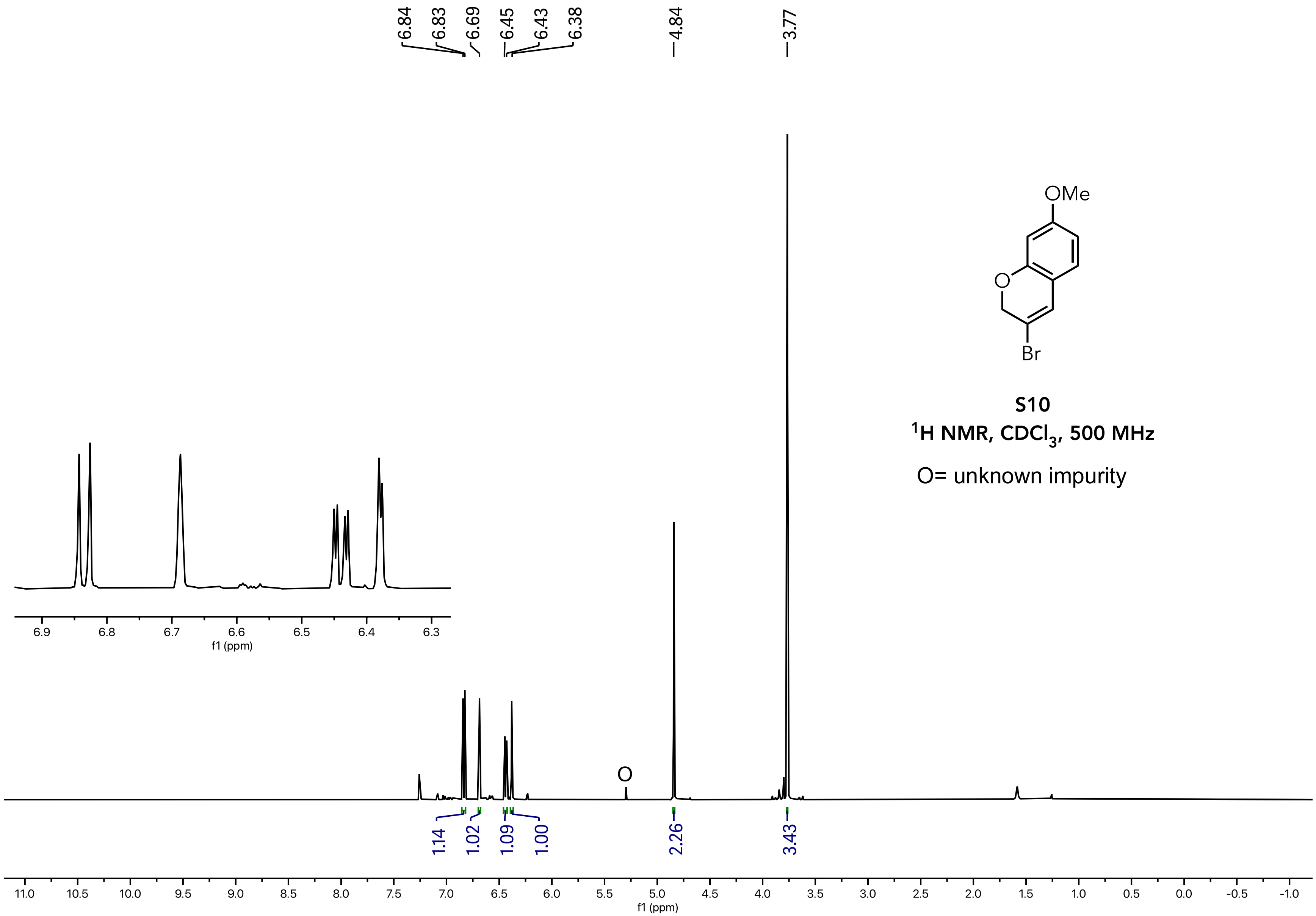


**S10**

**$^{1}H$ NMR, $CDCl_3$, 500 MHz**

O= unknown impurity

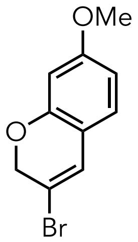


**S10**

**$^{13}C$ NMR, $CD_3OD$, 126 MHz**

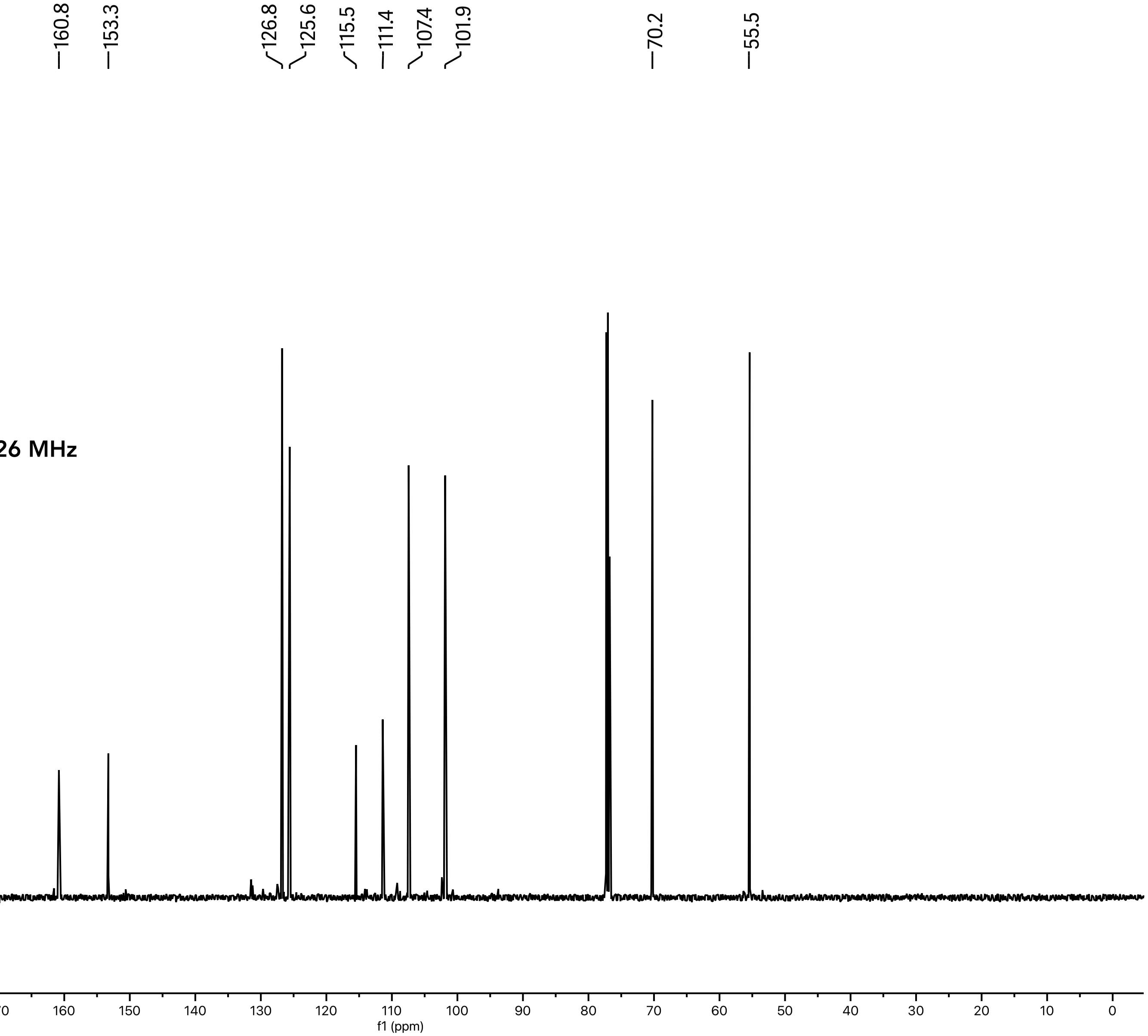

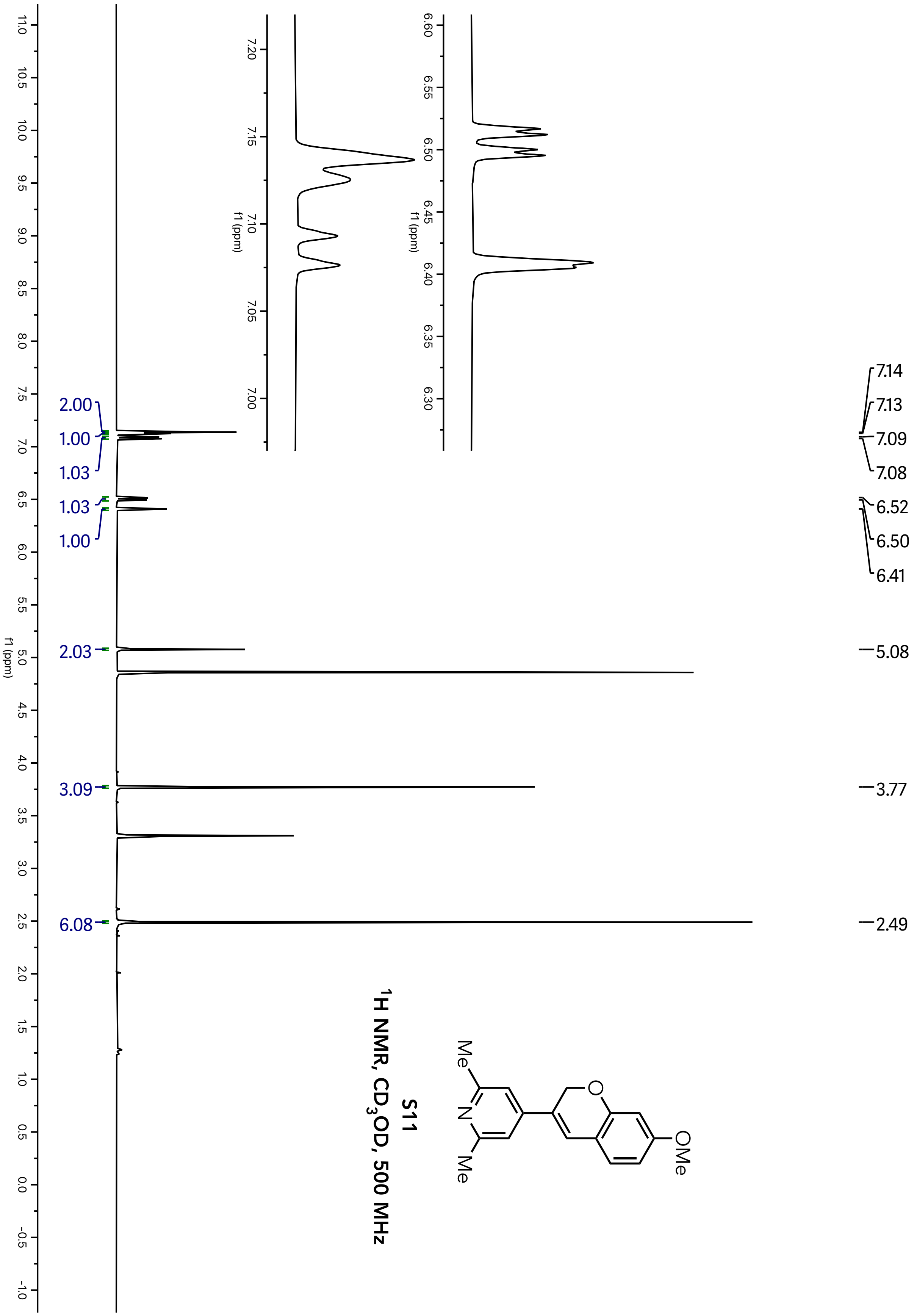
S11
1H NMR, CD3OD, 500 MHz
OMe
Me
N
Me
7.14
7.13
7.09
7.08
6.52
6.50
6.41
5.08
3.77
2.49
2.00
1.00
1.03
1.03
1.00
2.03
3.09
6.08
f1 (ppm)

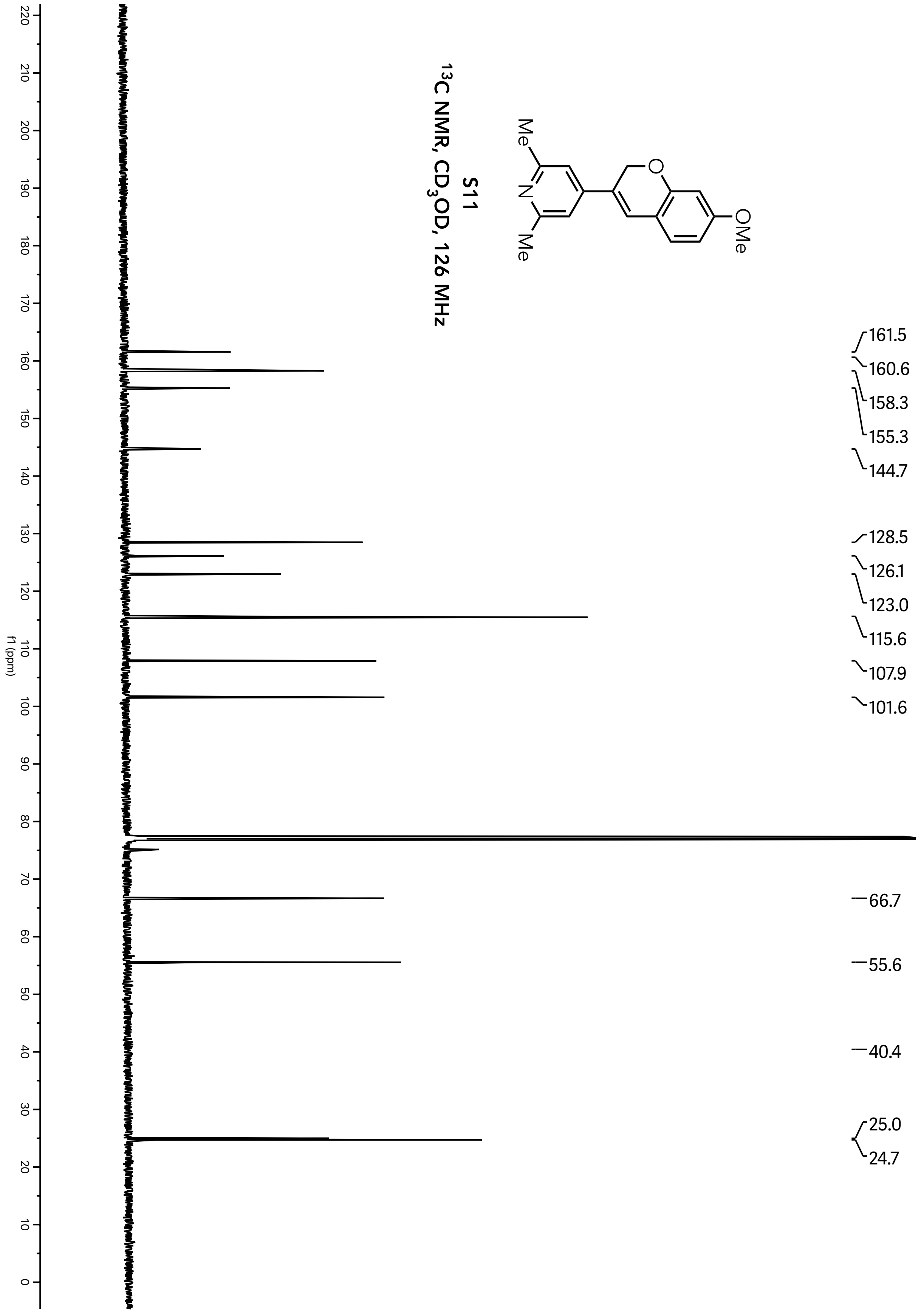
S11
13C NMR, CD3OD, 126 MHz
OMe
Me
N
Me
O
161.5
160.6
158.3
155.3
144.7
128.5
126.1
123.0
115.6
107.9
101.6
66.7
55.6
40.4
25.0
24.7
f1 (ppm)

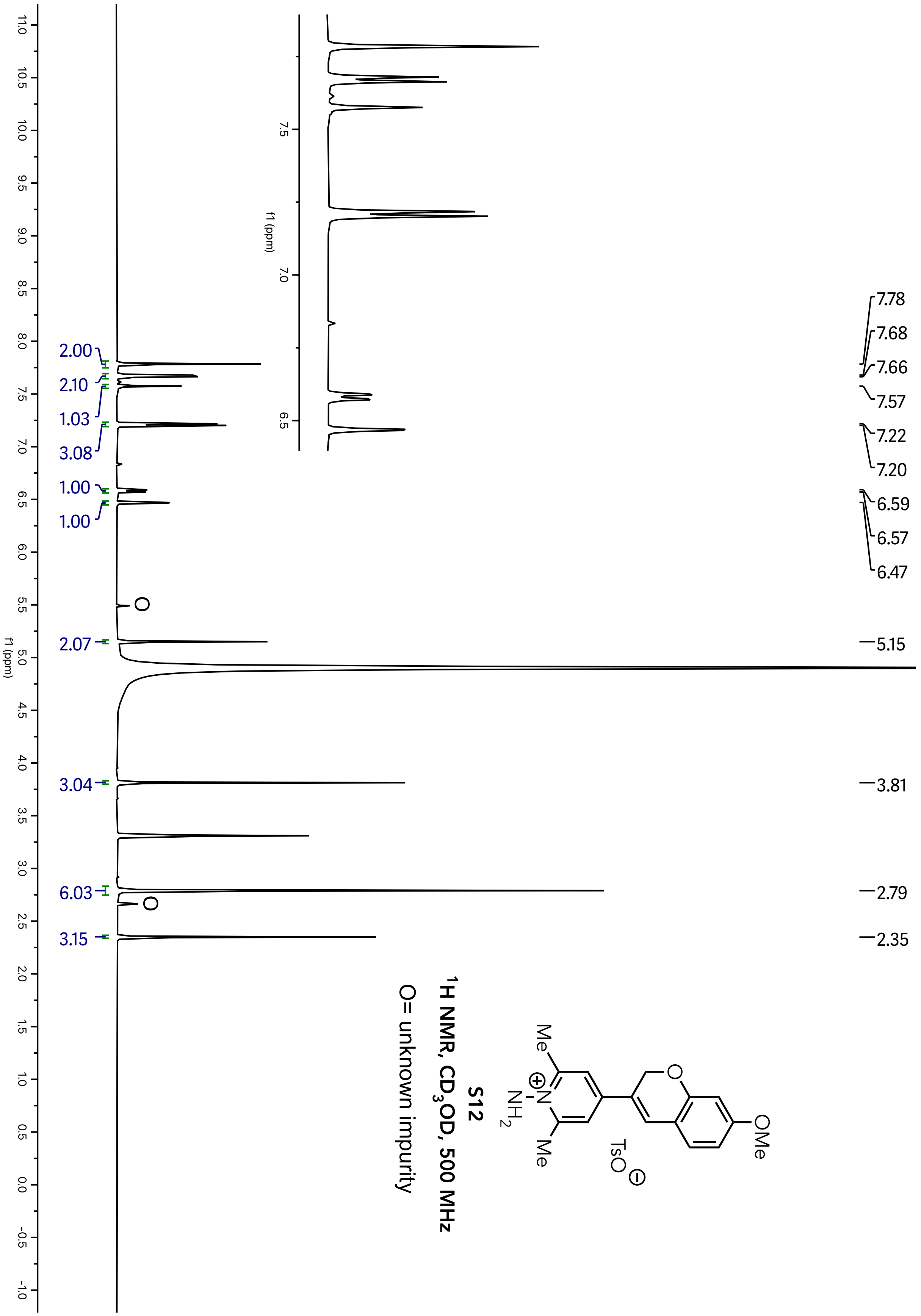
S12
1H NMR, CD3OD, 500 MHz
O= unknown impurity
Me
OMe
TsO
NH2
f1 (ppm)
7.78
7.68
7.66
7.57
7.22
7.20
6.59
6.57
6.47
5.15
3.81
2.79
2.35
2.00
2.10
1.03
3.08
1.00
1.00
2.07
3.04
6.03
3.15

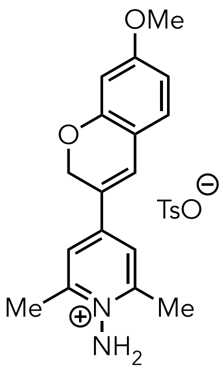


**S12**

**$^{13}C$ NMR, $CD_3OD$, 126 MHz**

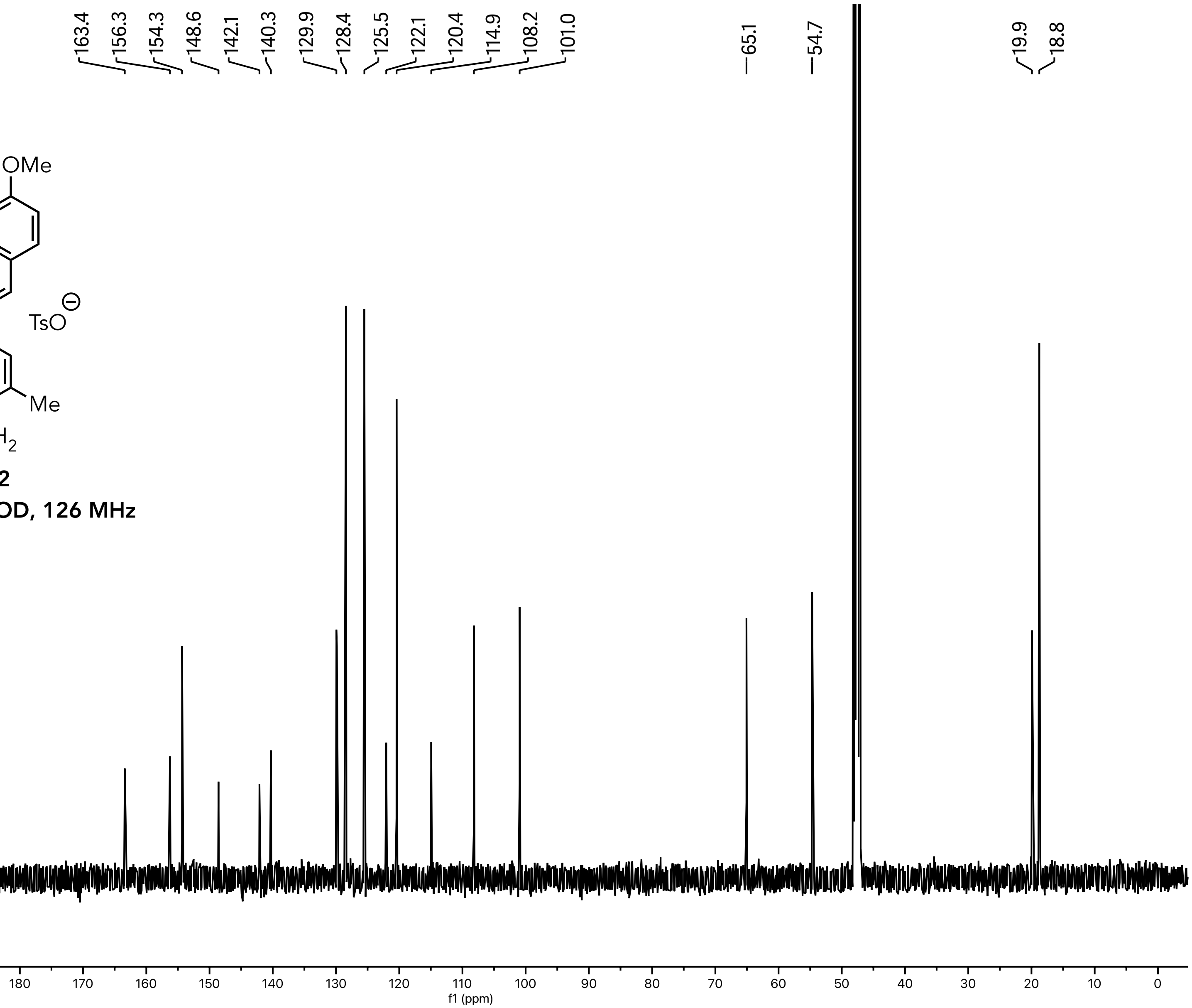

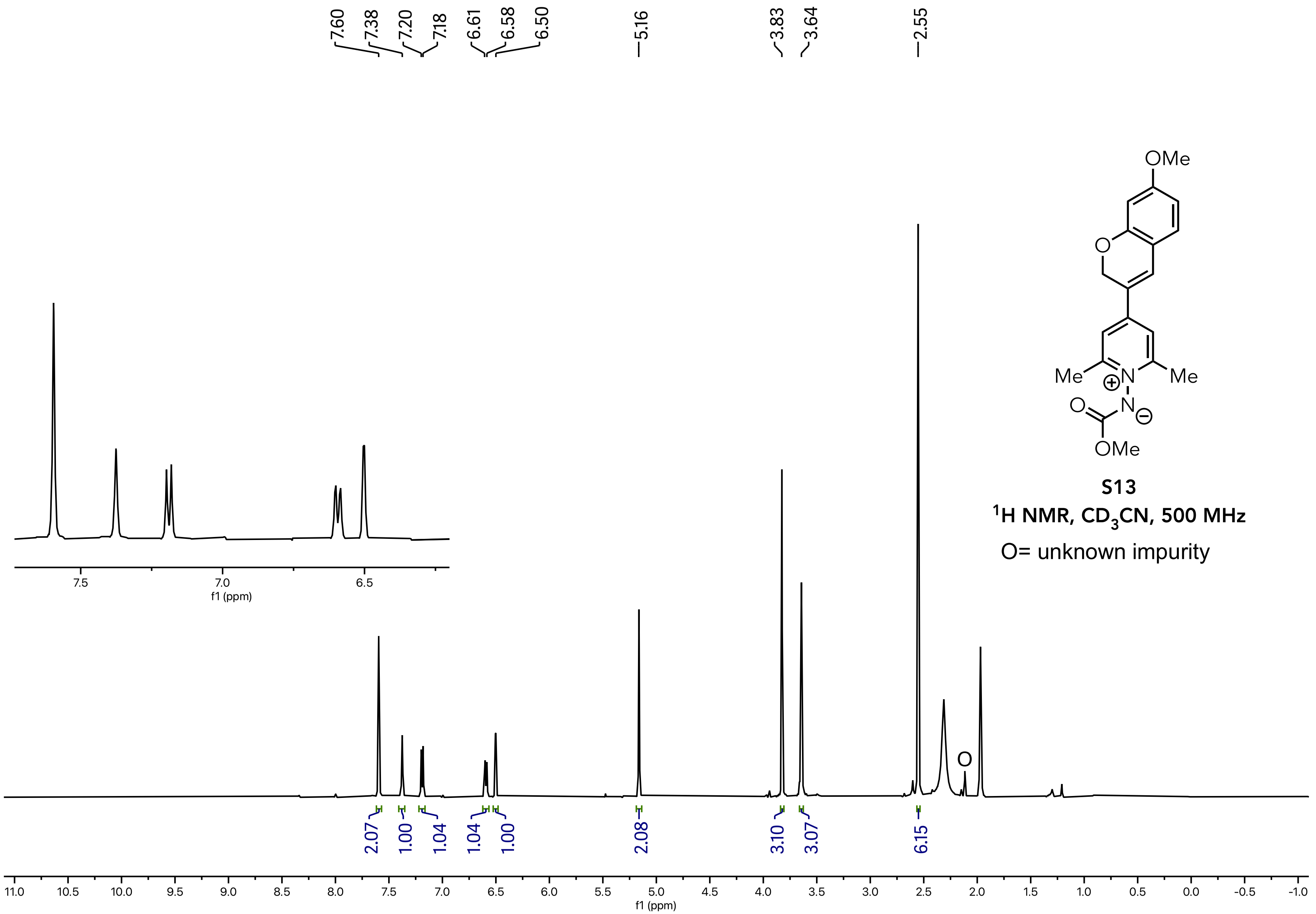


**S13**

$^1$H NMR, $CD_3CN$, 500 MHz

O= unknown impurity

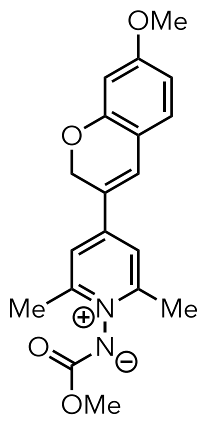


**S13**

**$^{13}C$ NMR, $CD_3CN$, 126 MHz**

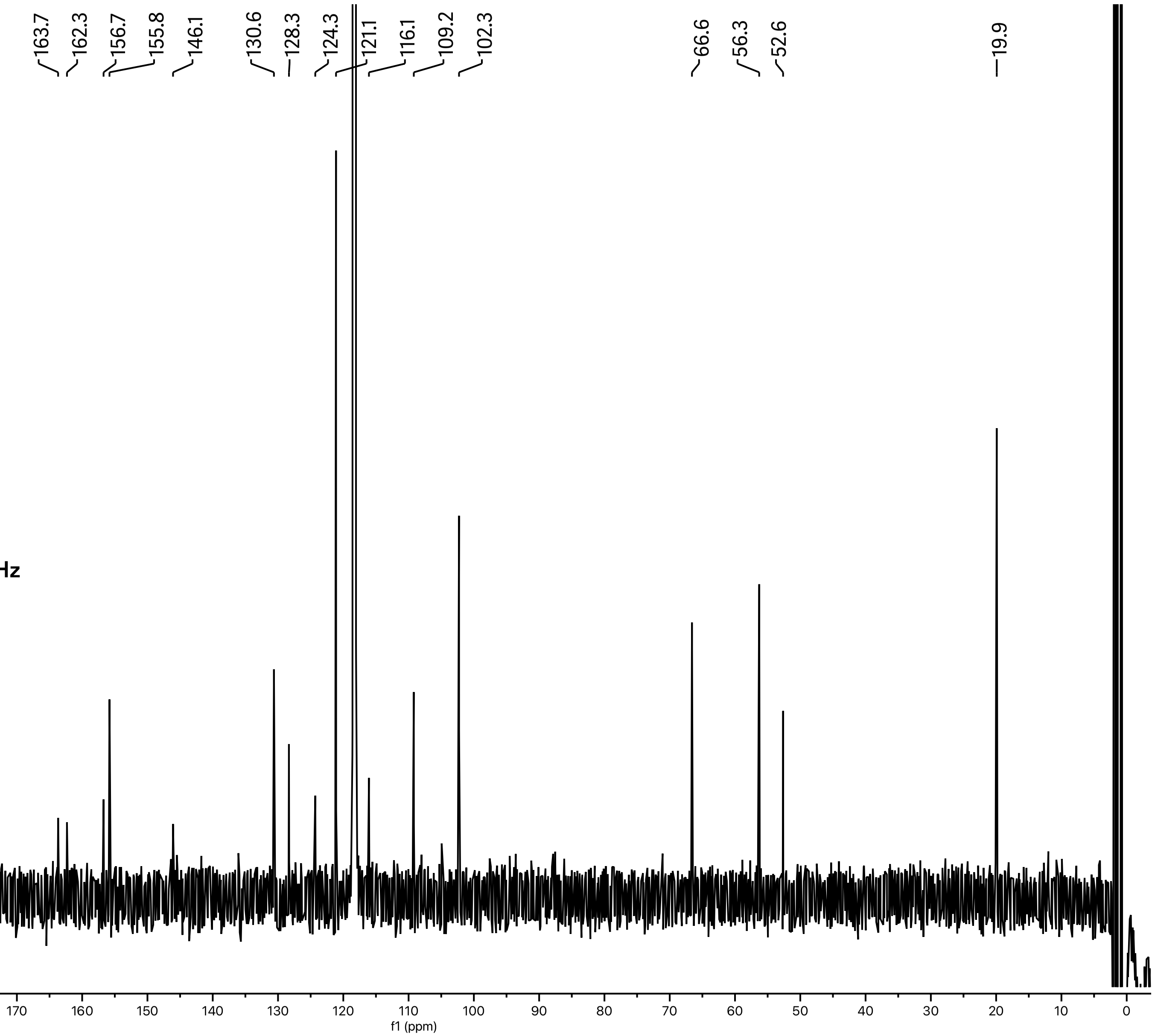

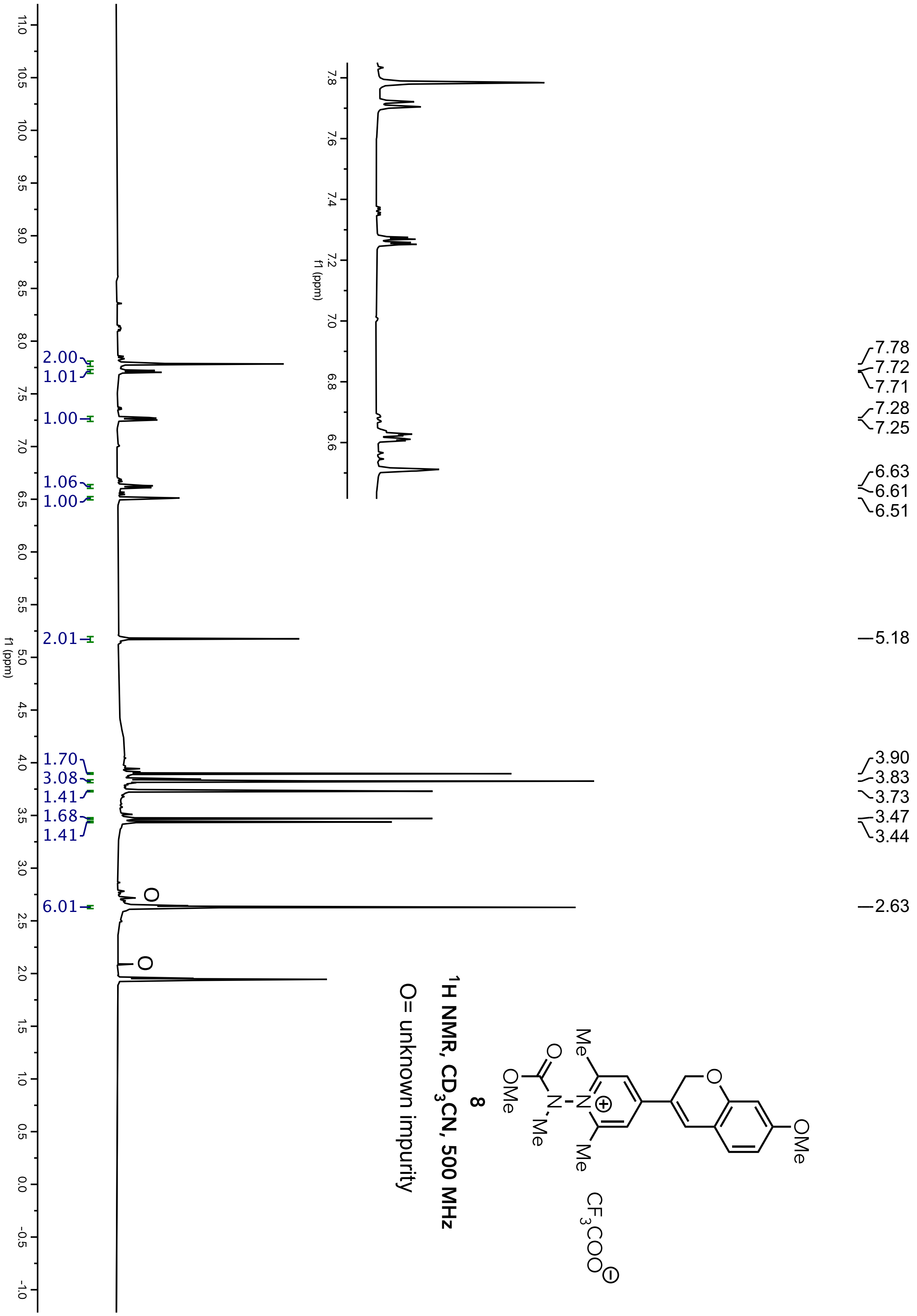
8
$^1$H NMR, $CD_3CN$, 500 MHz
O= unknown impurity
7.78
7.72
7.71
7.28
7.25
6.63
6.61
6.51
5.18
3.90
3.83
3.73
3.47
3.44
2.63
2.00
1.01
1.00
1.06
1.00
2.01
1.70
3.08
1.41
1.68
1.41
6.01
f1 (ppm)
OMe
Me
O
N
Me
CF3COO

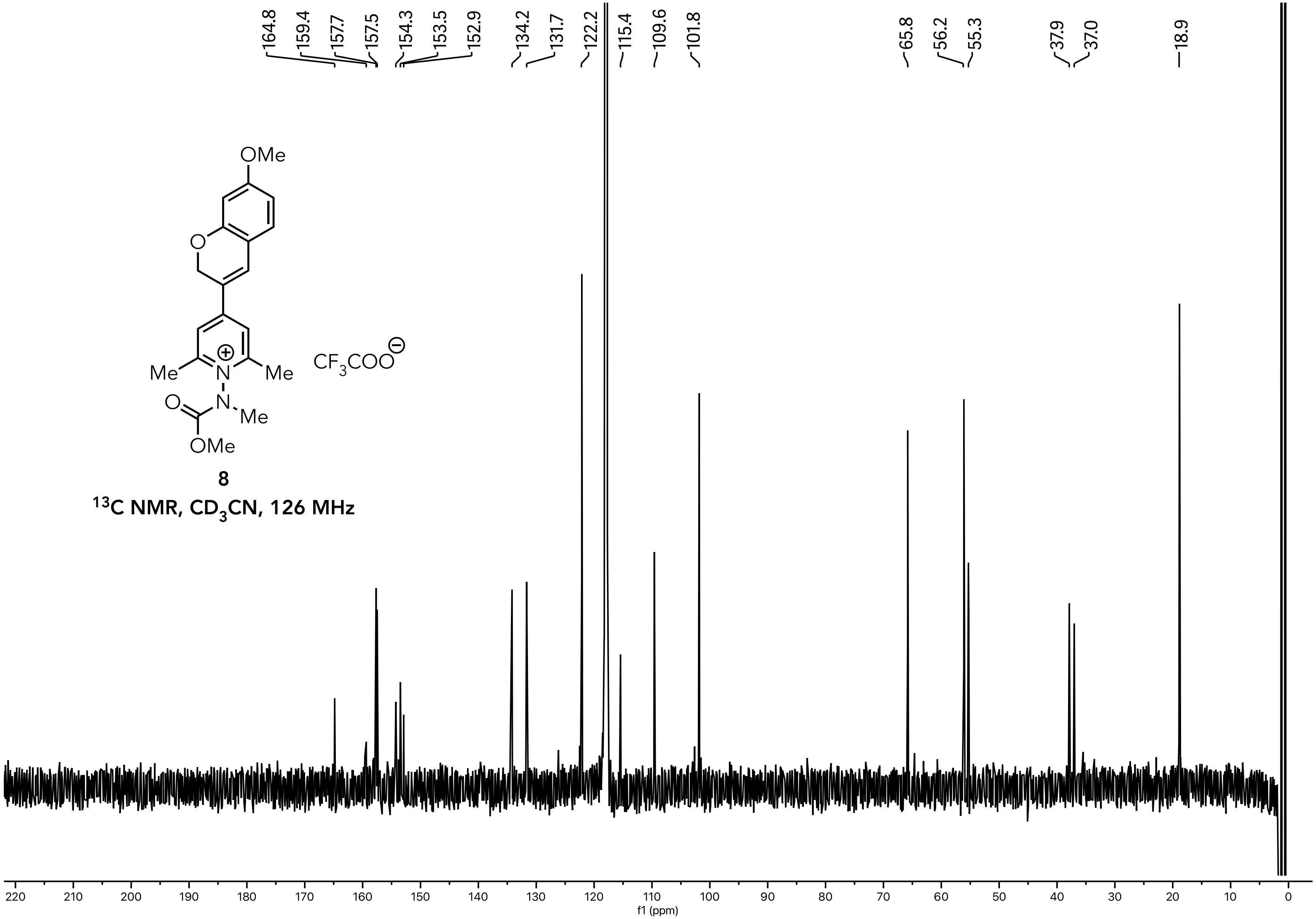


**8**

**$^{13}$C NMR, $CD_3CN$, 126 MHz**

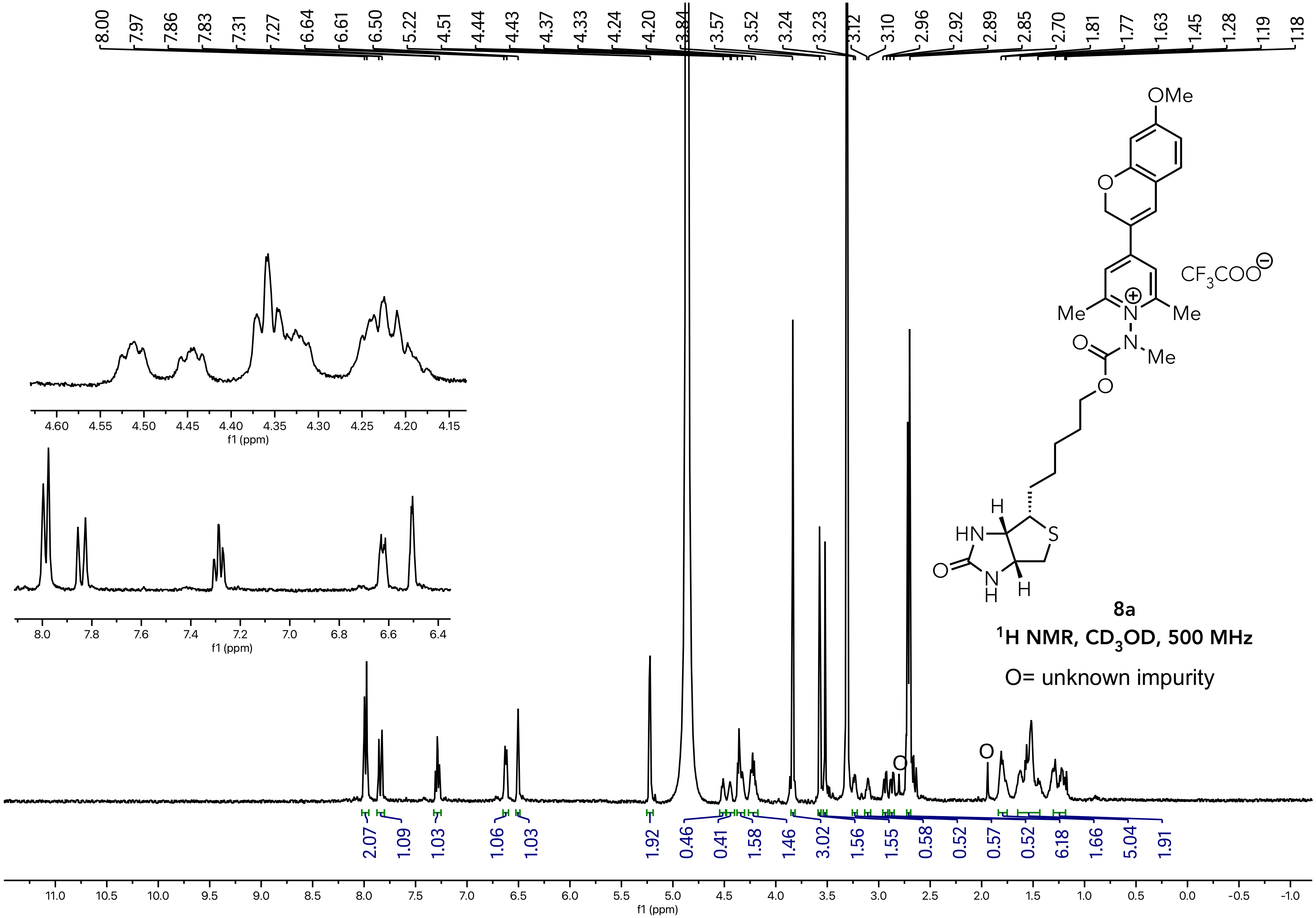

**8a**

**$^{1}$H NMR, $CD_3OD$, 500 MHz**

O= unknown impurity

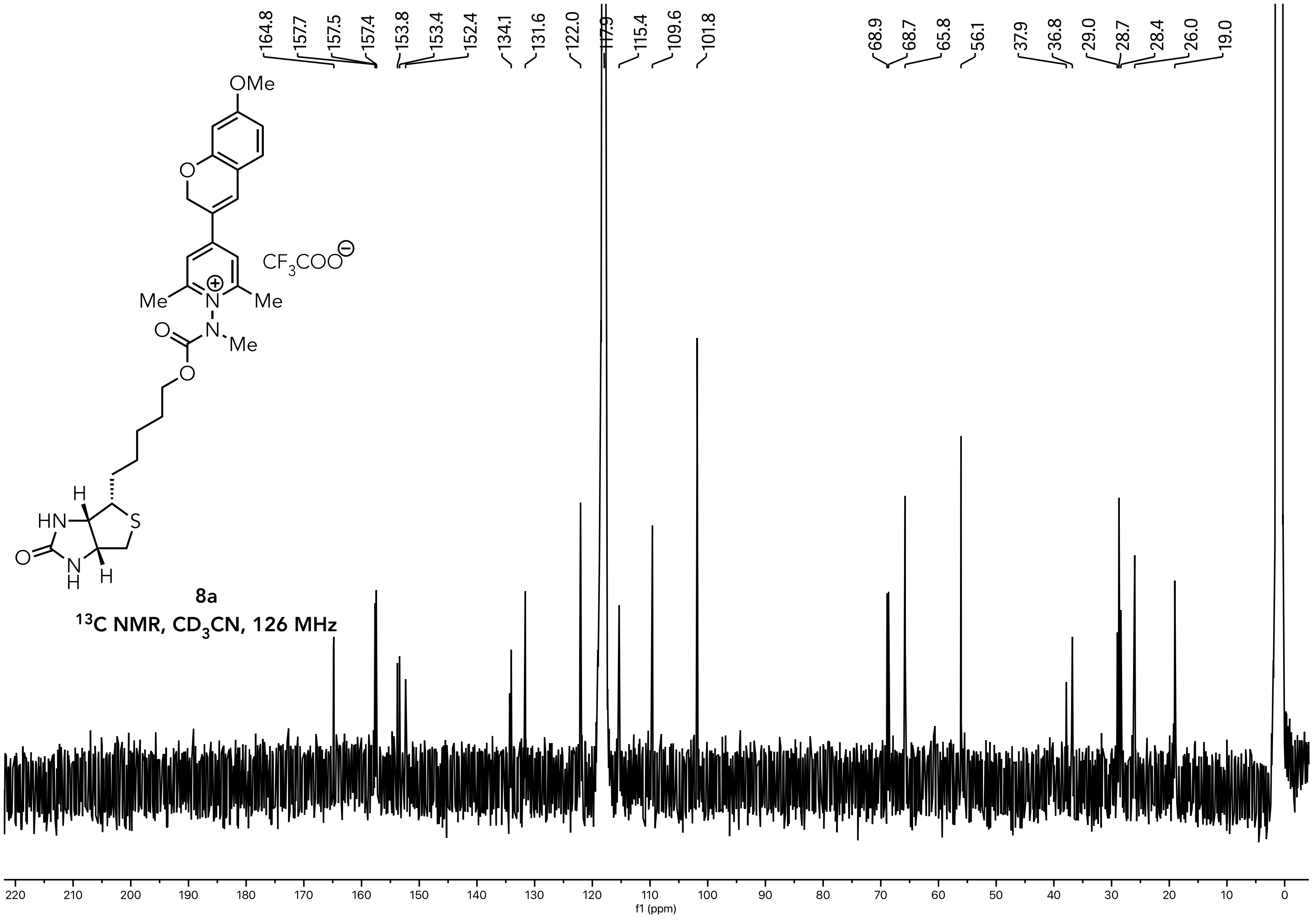


**8a**

**$^{13}$C NMR, $CD_3CN$, 126 MHz**

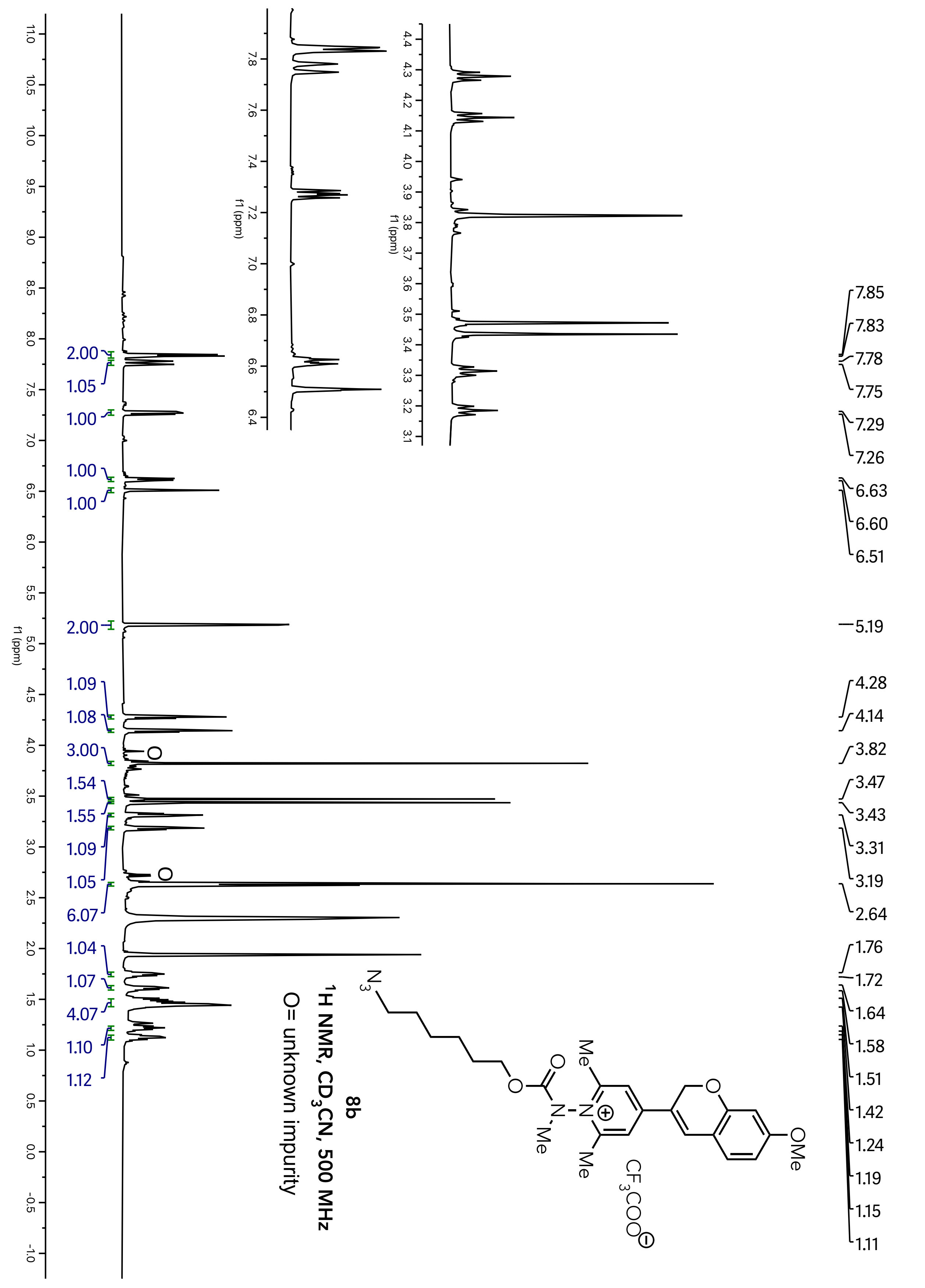
8b
$^1$H NMR, $CD_3CN$, 500 MHz
O= unknown impurity
f1 (ppm)
7.85 7.83 7.78 7.75 7.29 7.26 6.63 6.60 6.51 5.19 4.28 4.14 3.82 3.47 3.43 3.31 3.19 2.64 1.76 1.72 1.64 1.58 1.51 1.42 1.24 1.19 1.15 1.11
2.00 1.05 1.00 1.00 1.00 2.00 1.09 1.08 3.00 1.54 1.55 1.09 1.05 6.07 1.04 1.07 4.07 1.10 1.12
OMe
Me
CF3COO

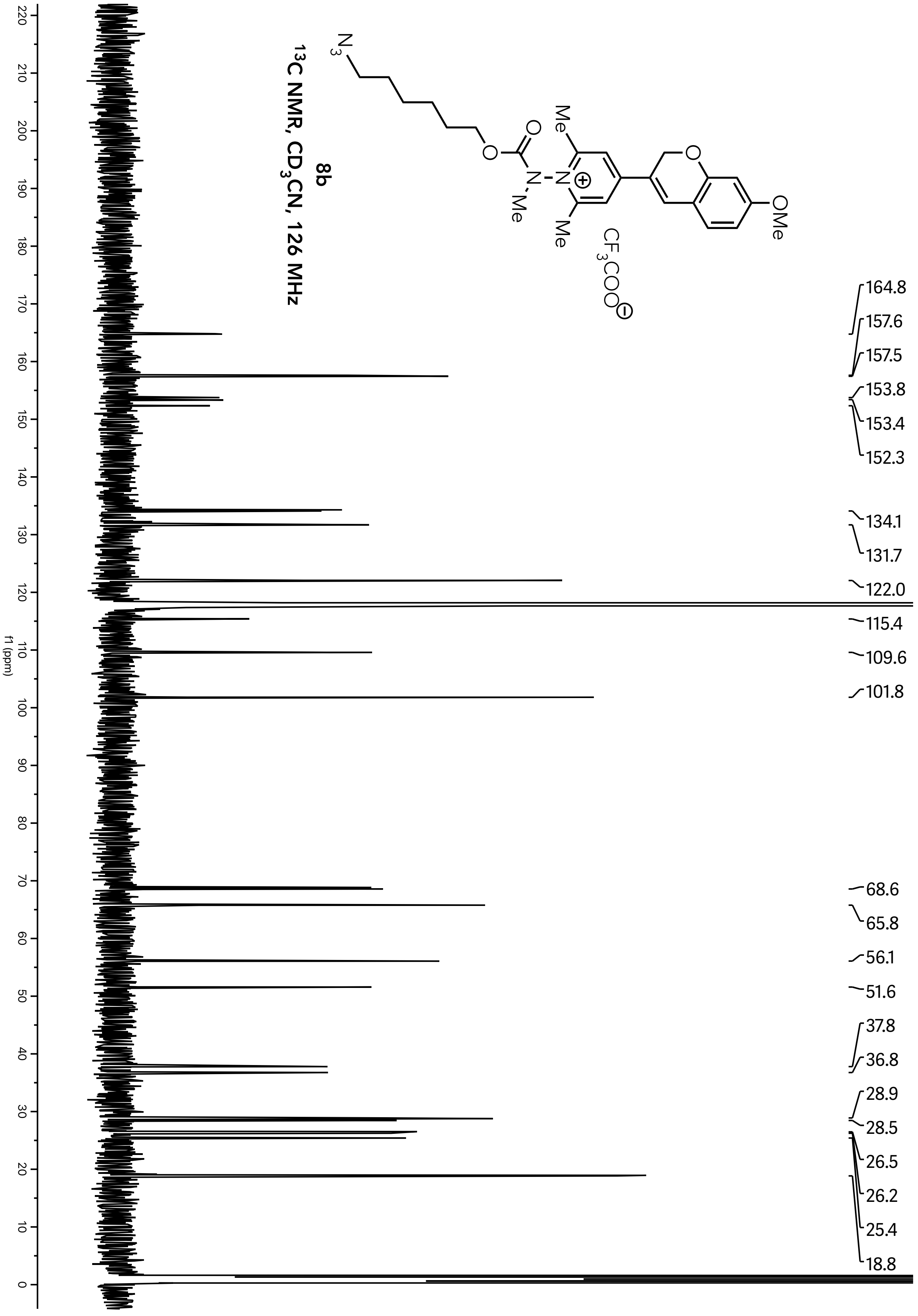
8b
13C NMR, CD3CN, 126 MHz
164.8
157.6
157.5
153.8
153.4
152.3
134.1
131.7
122.0
115.4
109.6
101.8
68.6
65.8
56.1
51.6
37.8
36.8
28.9
28.5
26.5
26.2
25.4
18.8
f1 (ppm)

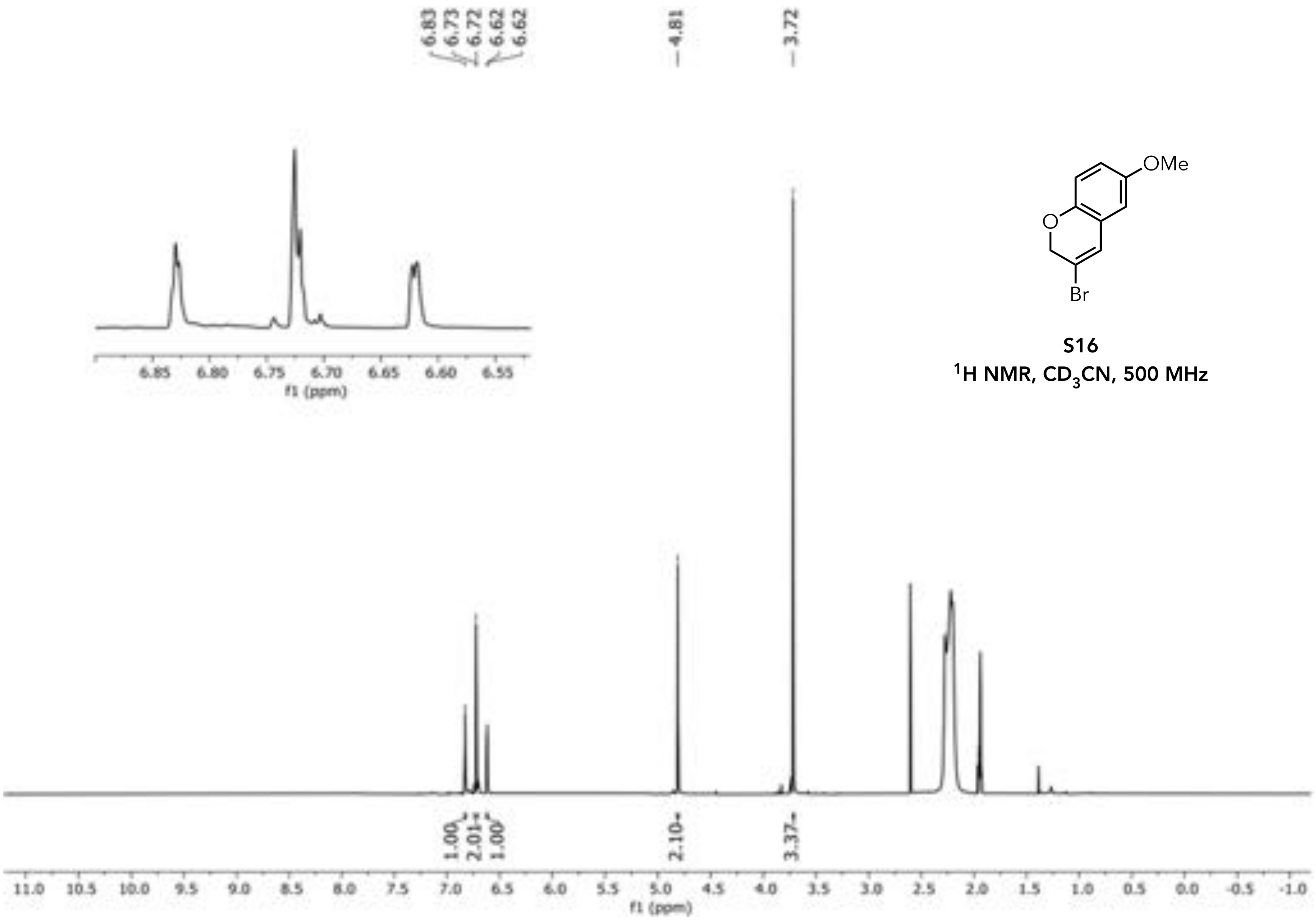


**S16**
**$^1H$ NMR, $CD_3CN$, 500 MHz**

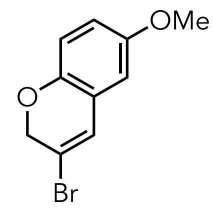


**S16**

$^{13}C$ NMR, $CD_3CN$, 126 MHz

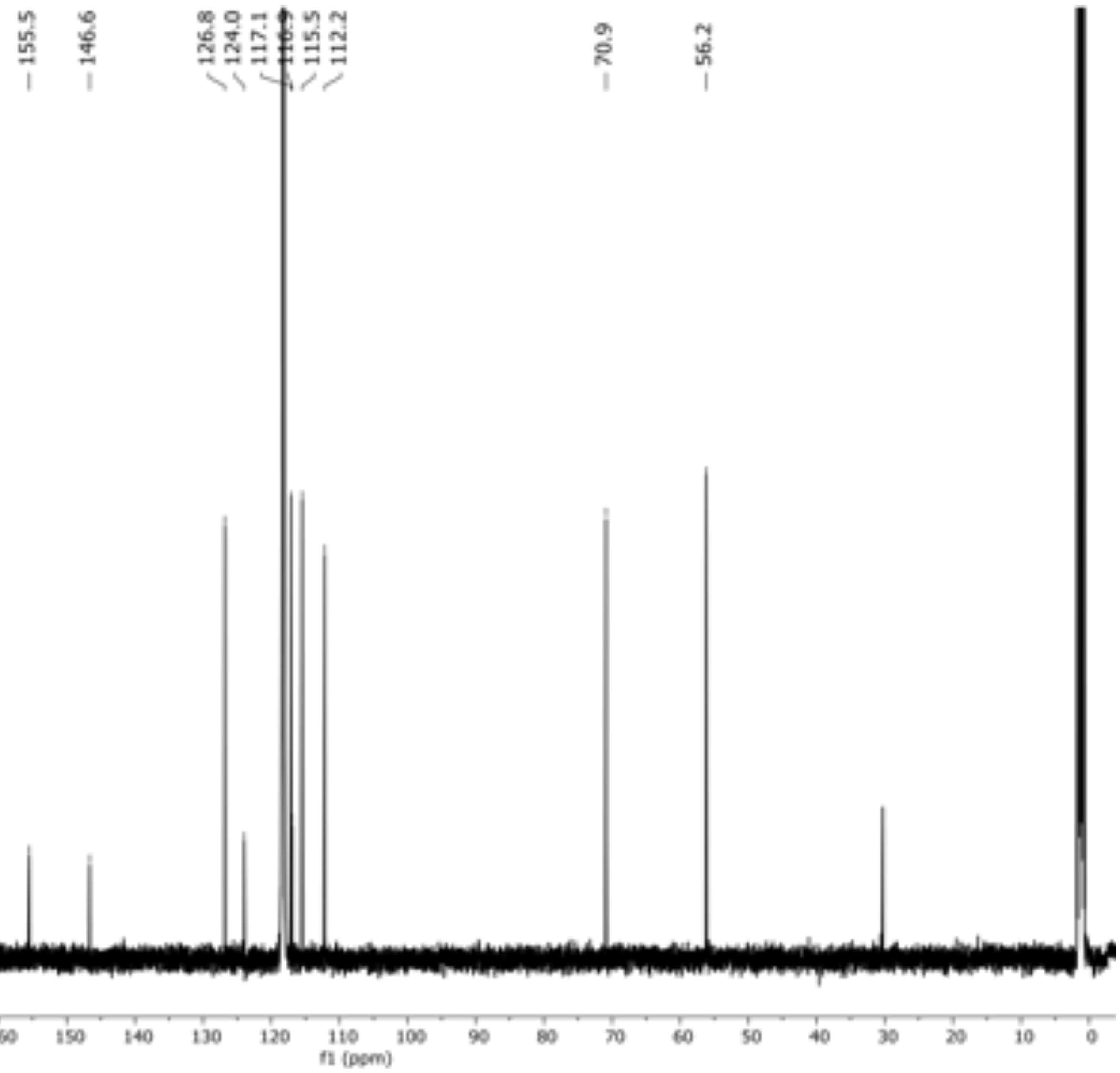

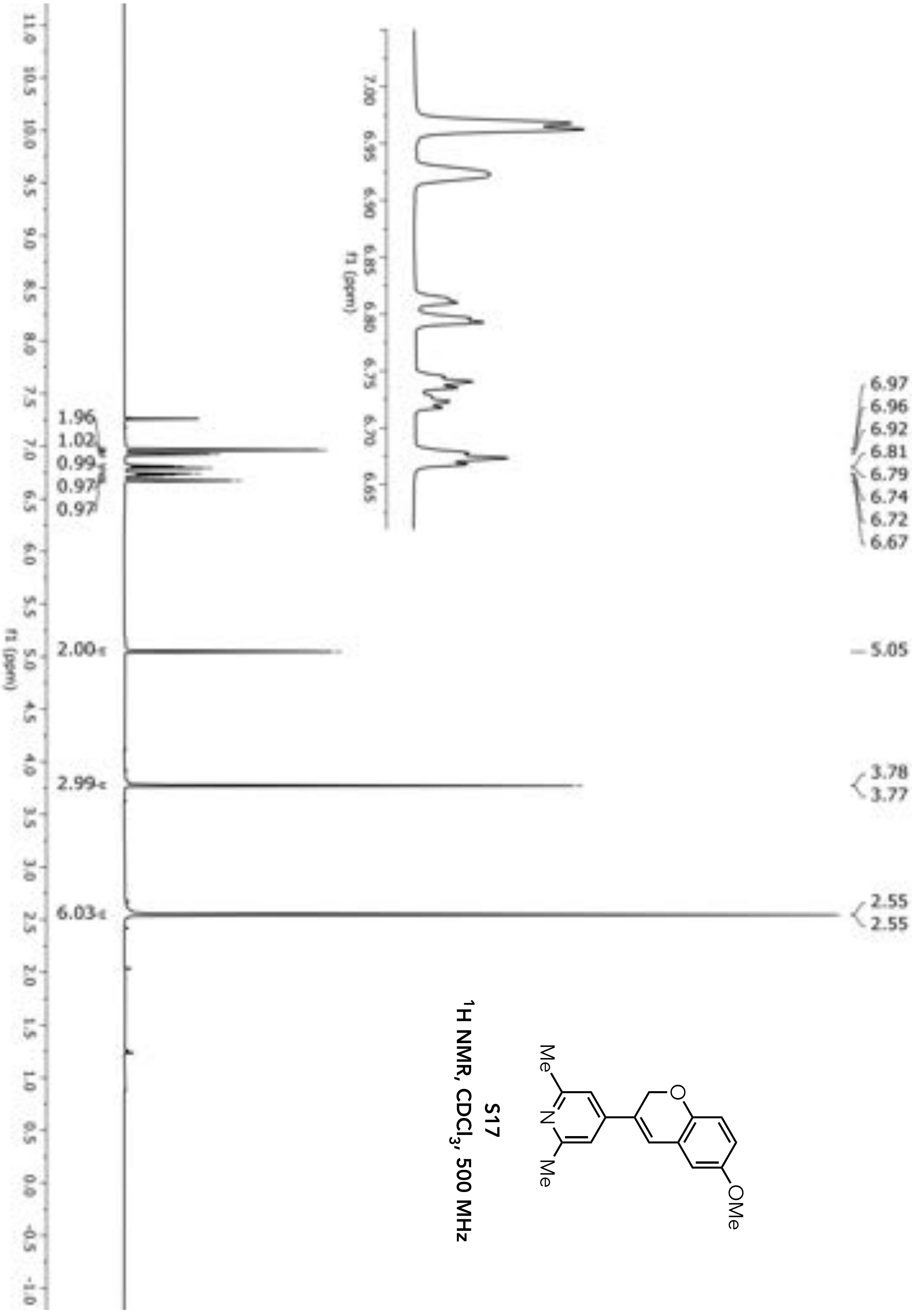


**S17**
**$^1$H NMR, $CDCl_3$, 500 MHz**

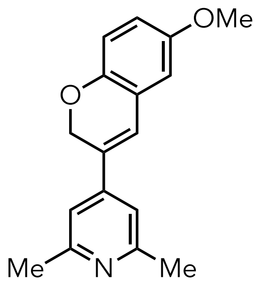


**S17**

**$^{13}C$ NMR, $CDCl_3$, 126 MHz**

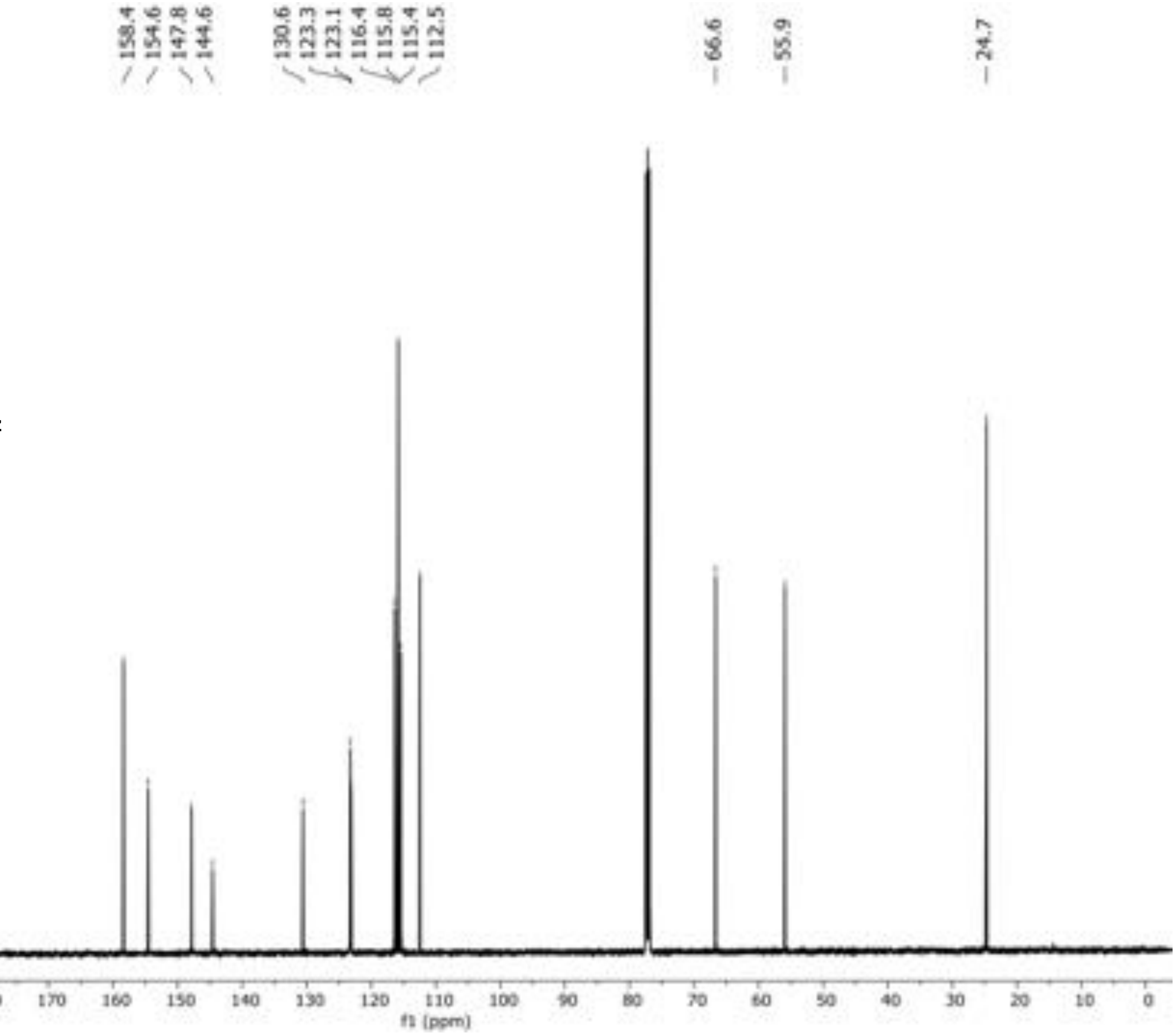

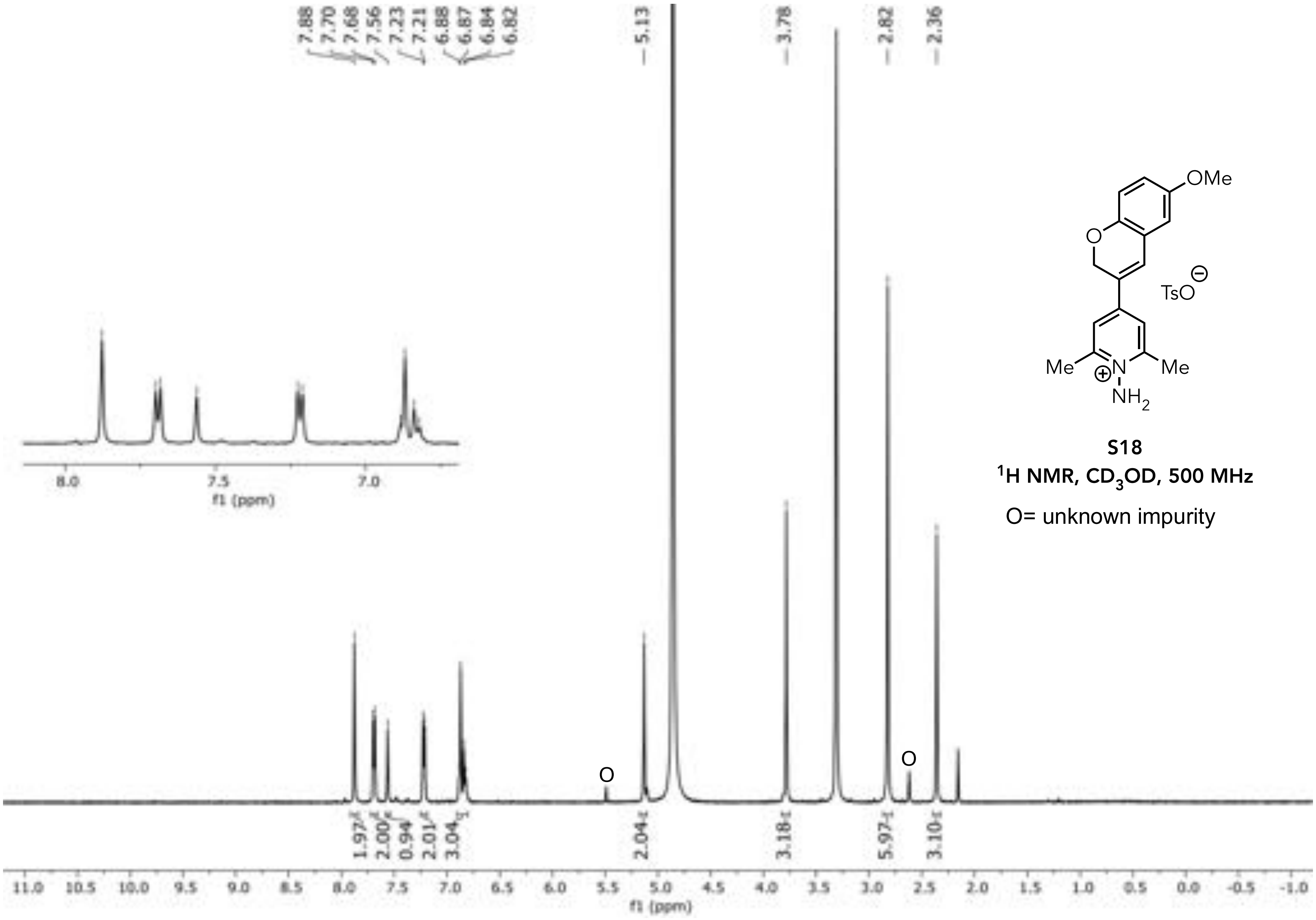


**S18**

$^1$H NMR, $CD_3OD$, 500 MHz

O= unknown impurity

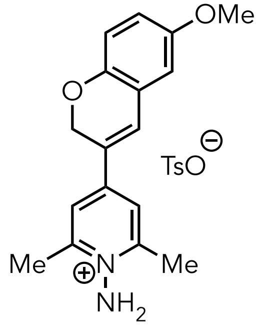


**S18**

$^{13}$CH NMR, $CD_3OD$, 126 MHz

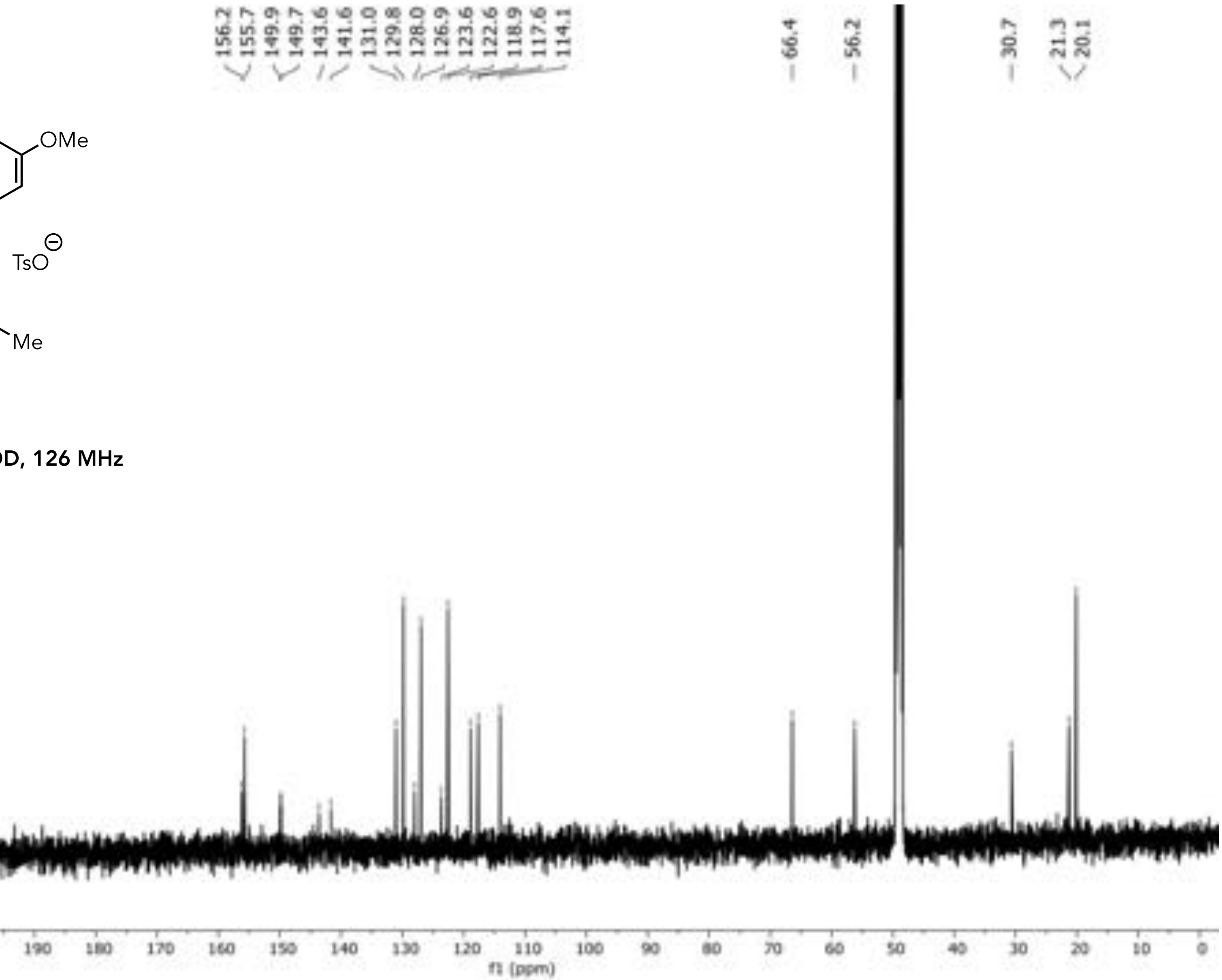

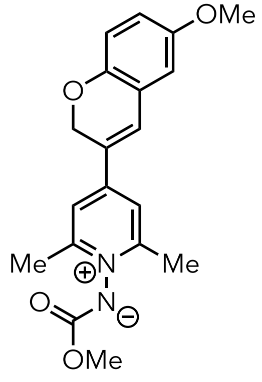


**S19**

**$^{1}H$ NMR, $CD_3CN$, 500 MHz**

O= unknown impurity

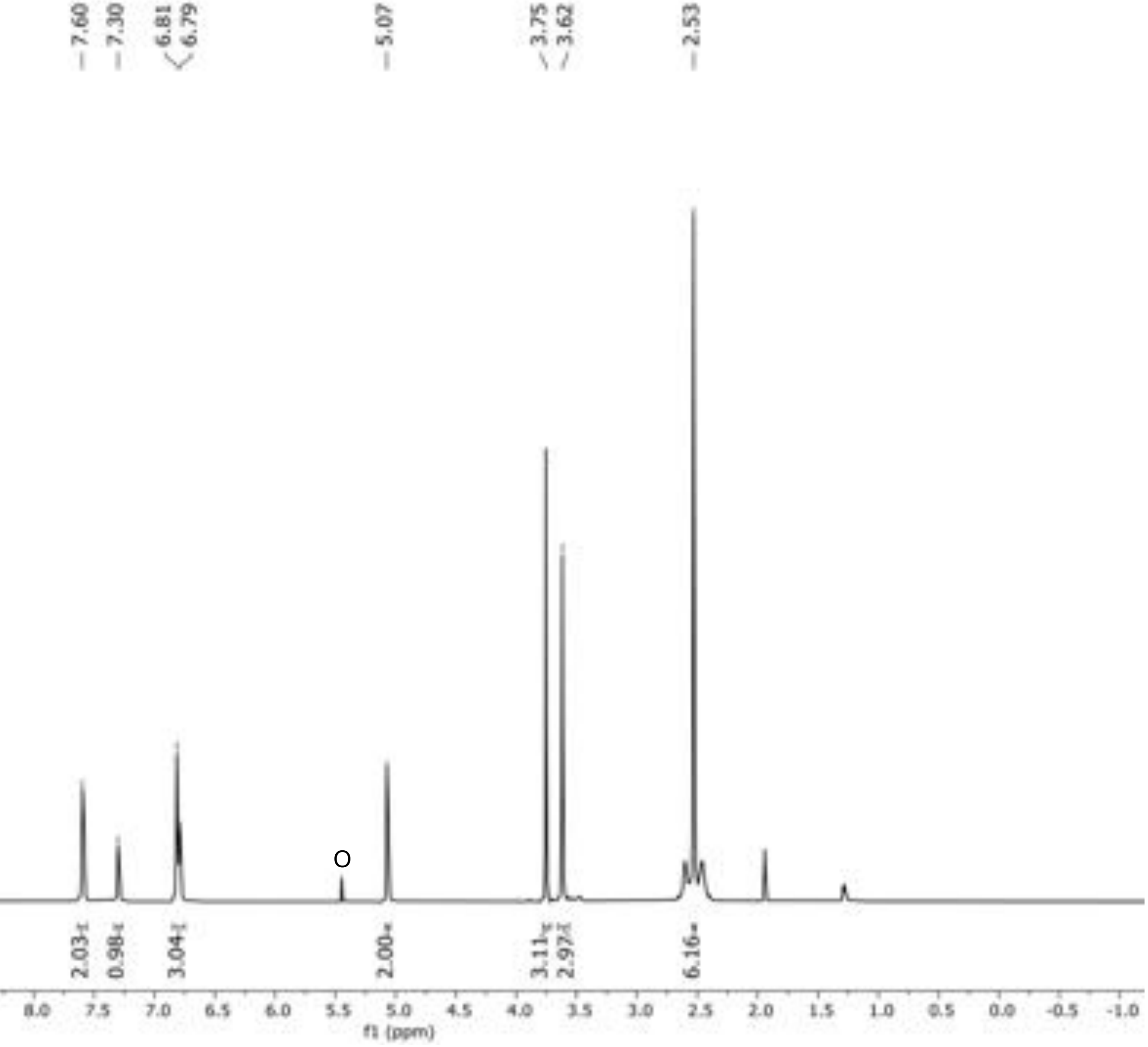

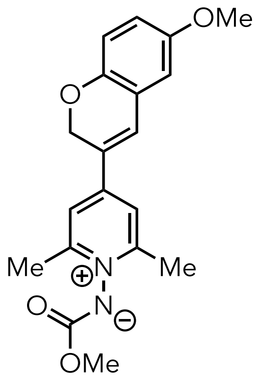


**S19**

**$^{13}C$ NMR, $CD_3CN$, 126 MHz**

O= unknown impurity

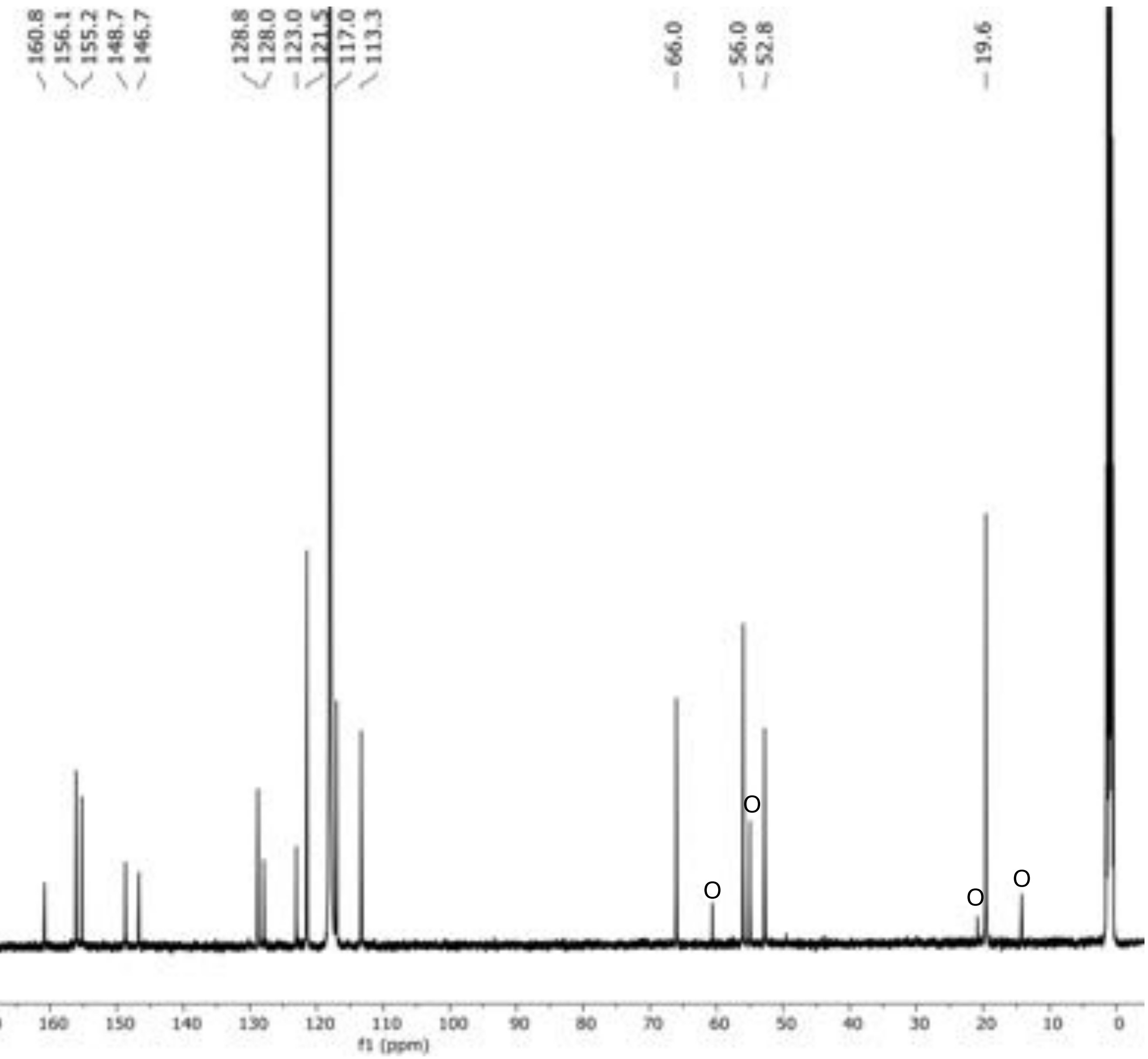

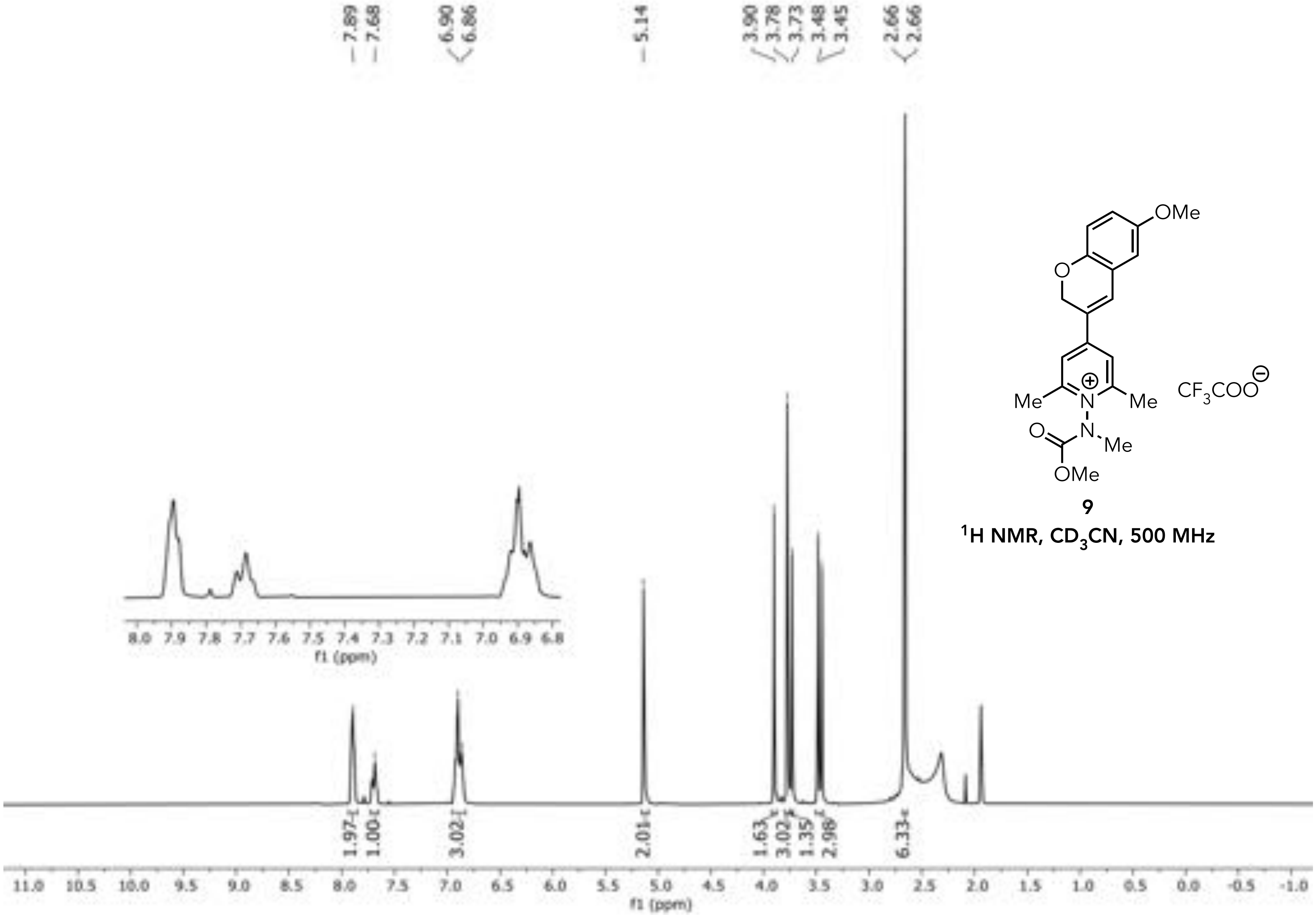


9

$^1H$ NMR, $CD_3CN$, 500 MHz

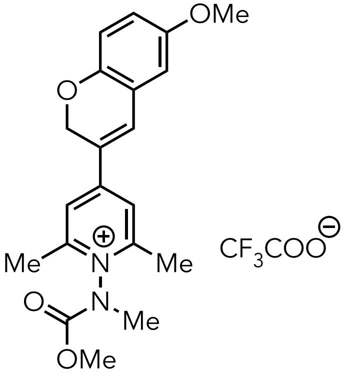

**9**

**$^{13}C$ NMR, $CD_3CN$, 126 MHz**

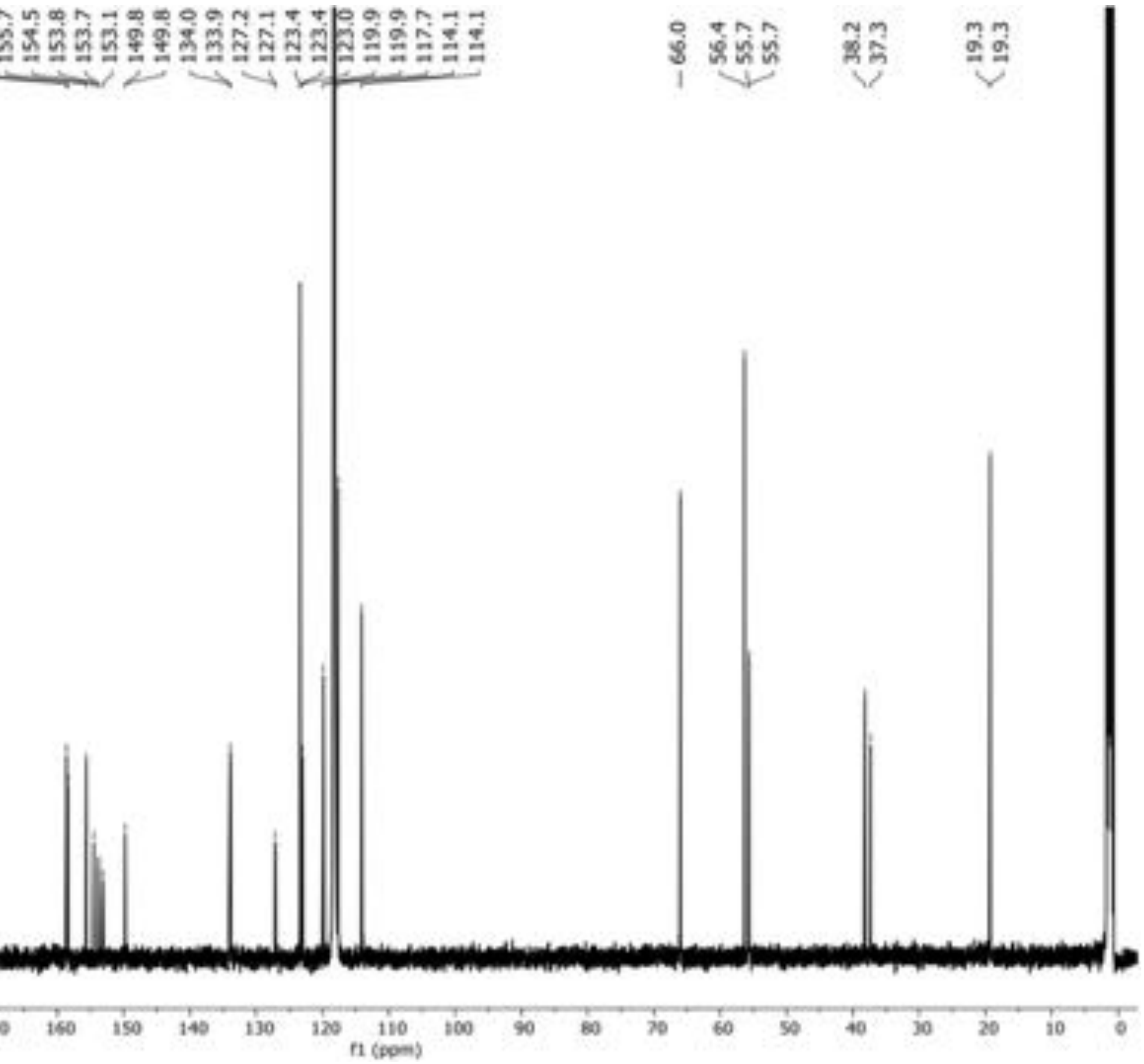

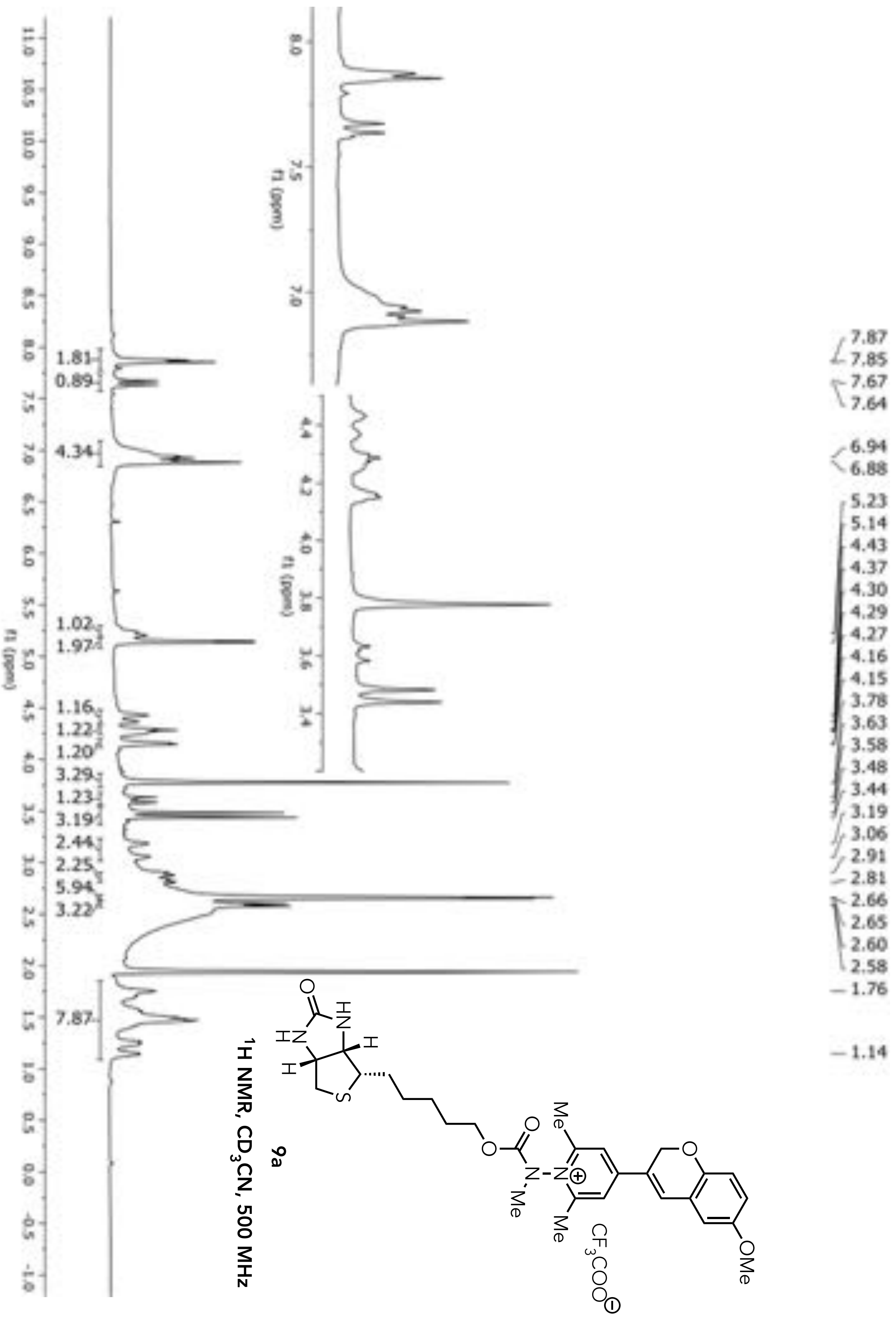


**9a**

$^1$H NMR, $CD_3CN$, 500 MHz

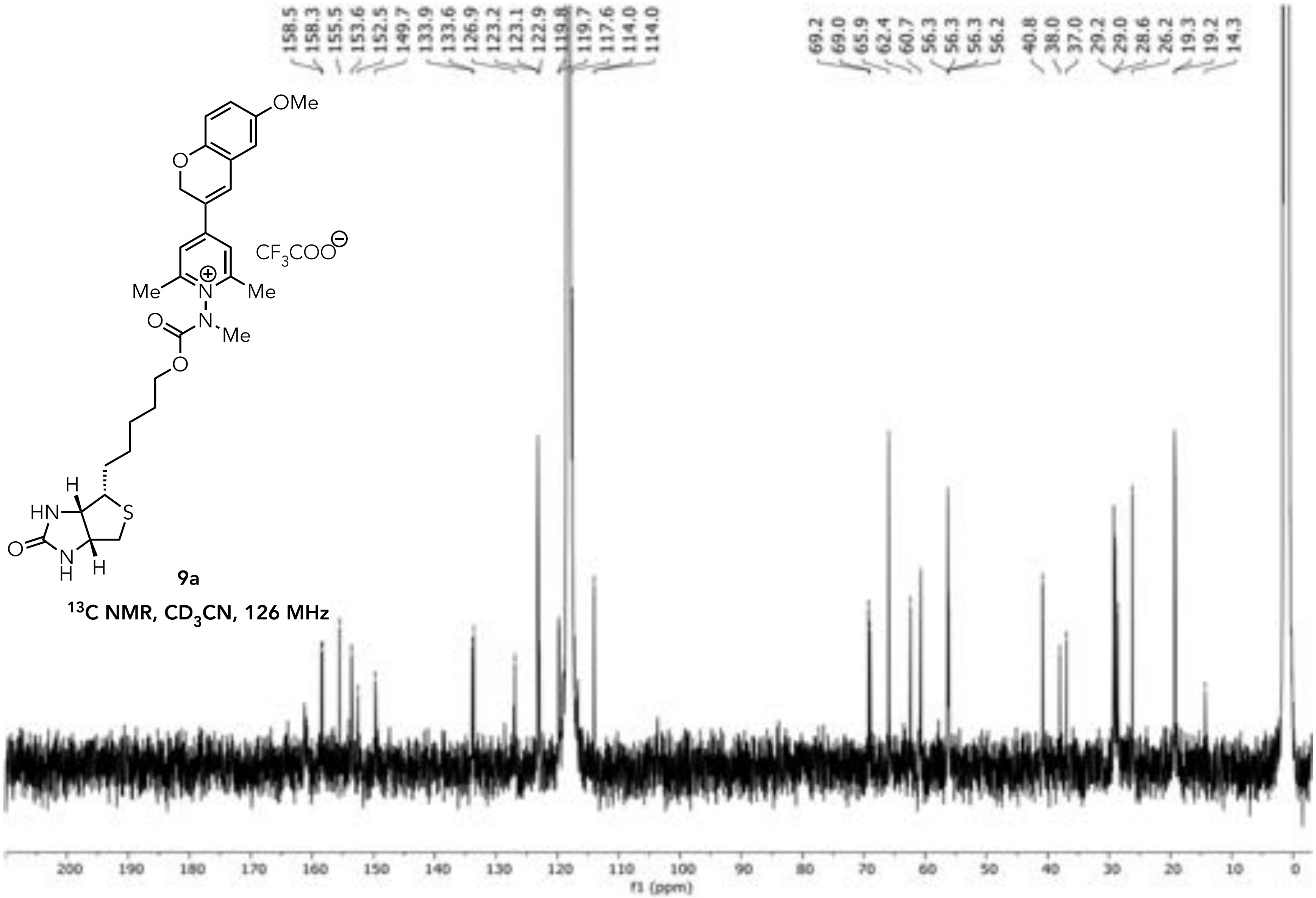


**9a**

**$^{13}C$ NMR, $CD_3CN$, 126 MHz**

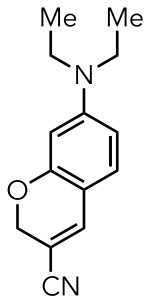


**S21**
**$^{1}H$ NMR, $CD_3CN$, 500 MHz**

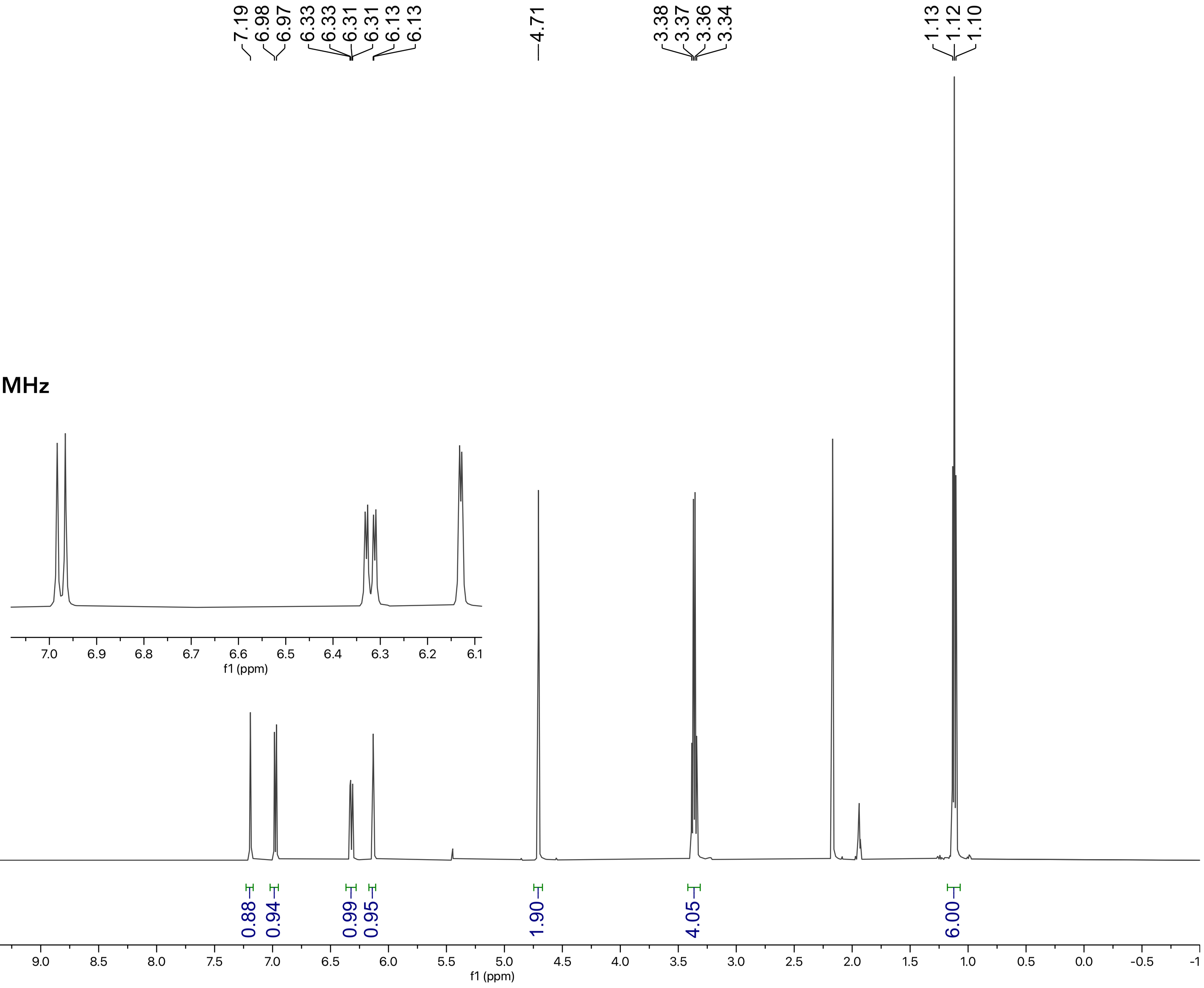

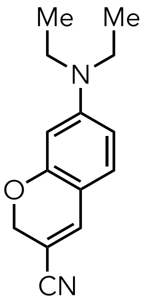

**S21**

**$^{13}C$ NMR, $CD_3CN$, 126 MHz**

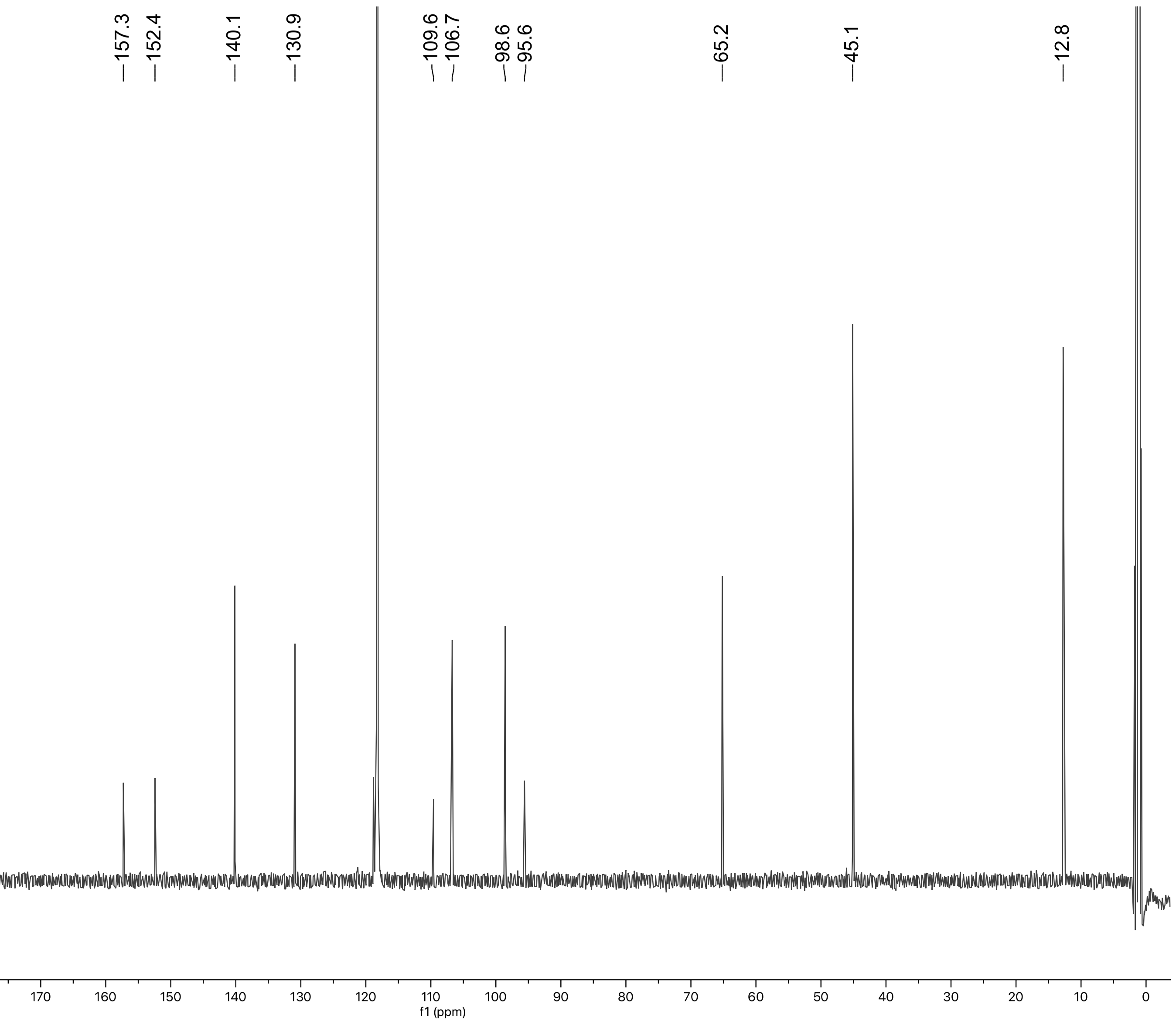

**S22**

**$^1$H NMR, $CD_3OD$, 500 MHz**

O= unknown impurity

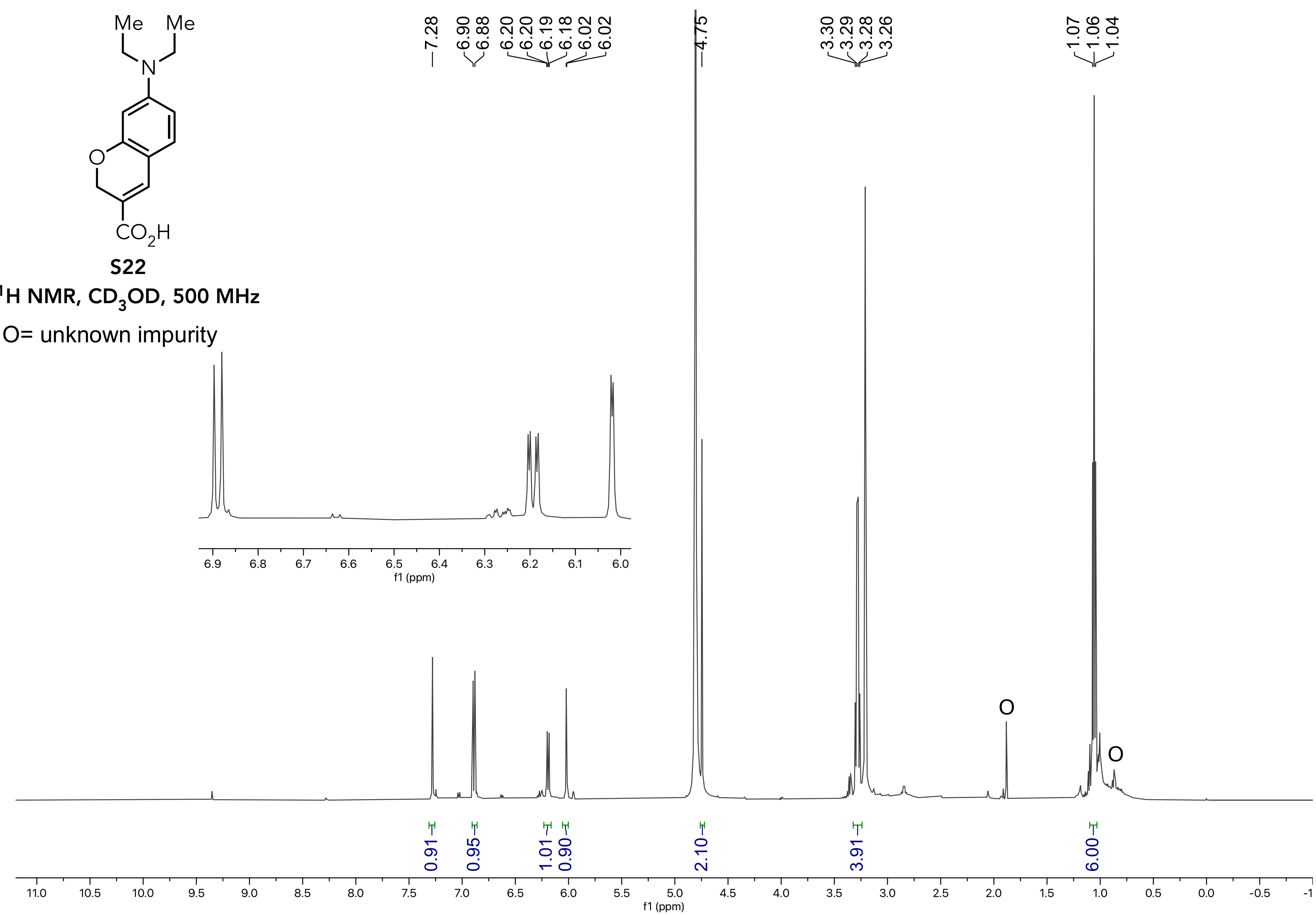

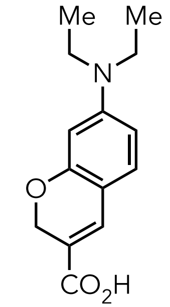


**S22**

**$^{13}C$ NMR, $CD_3OD$, 126 MHz**

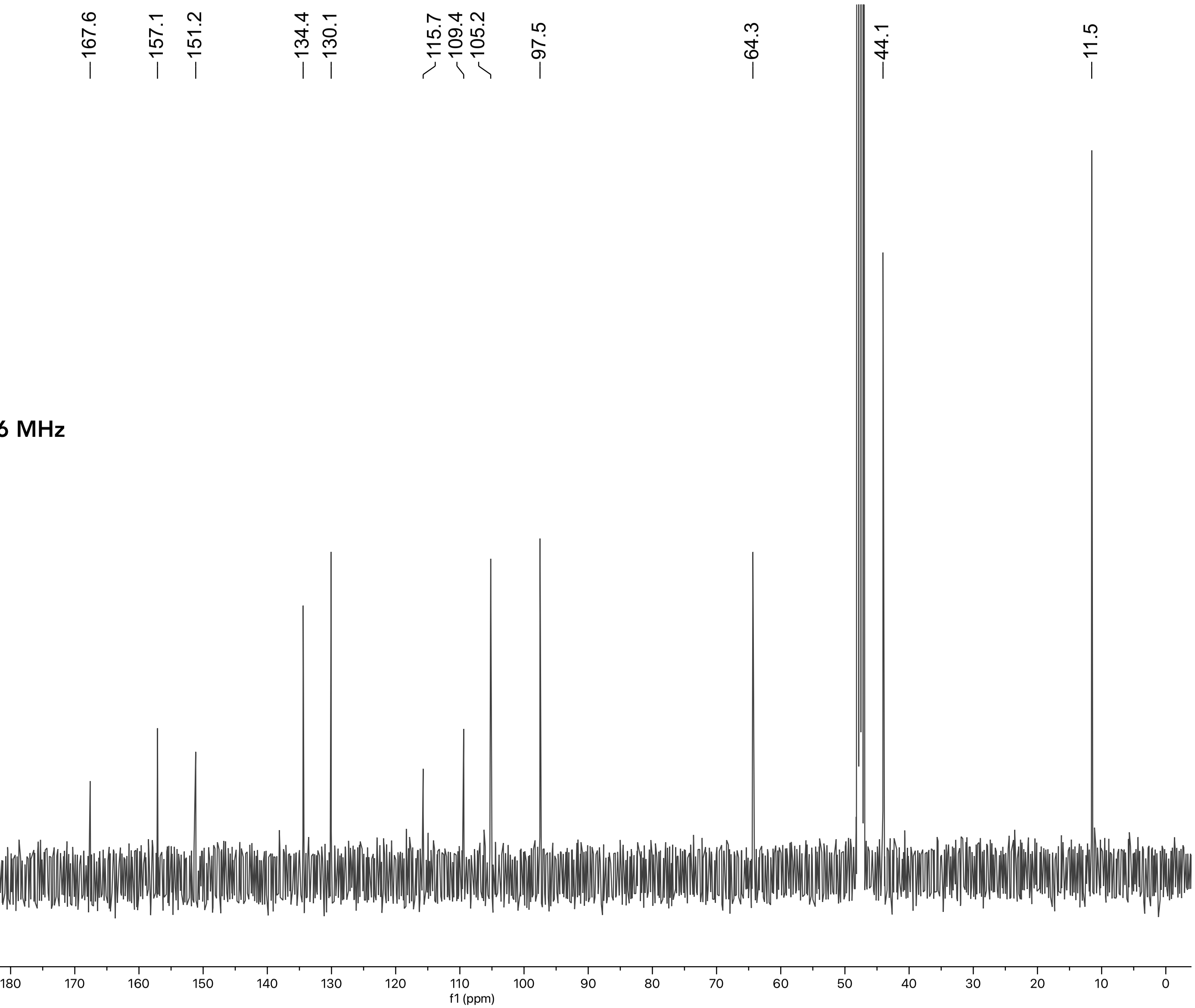

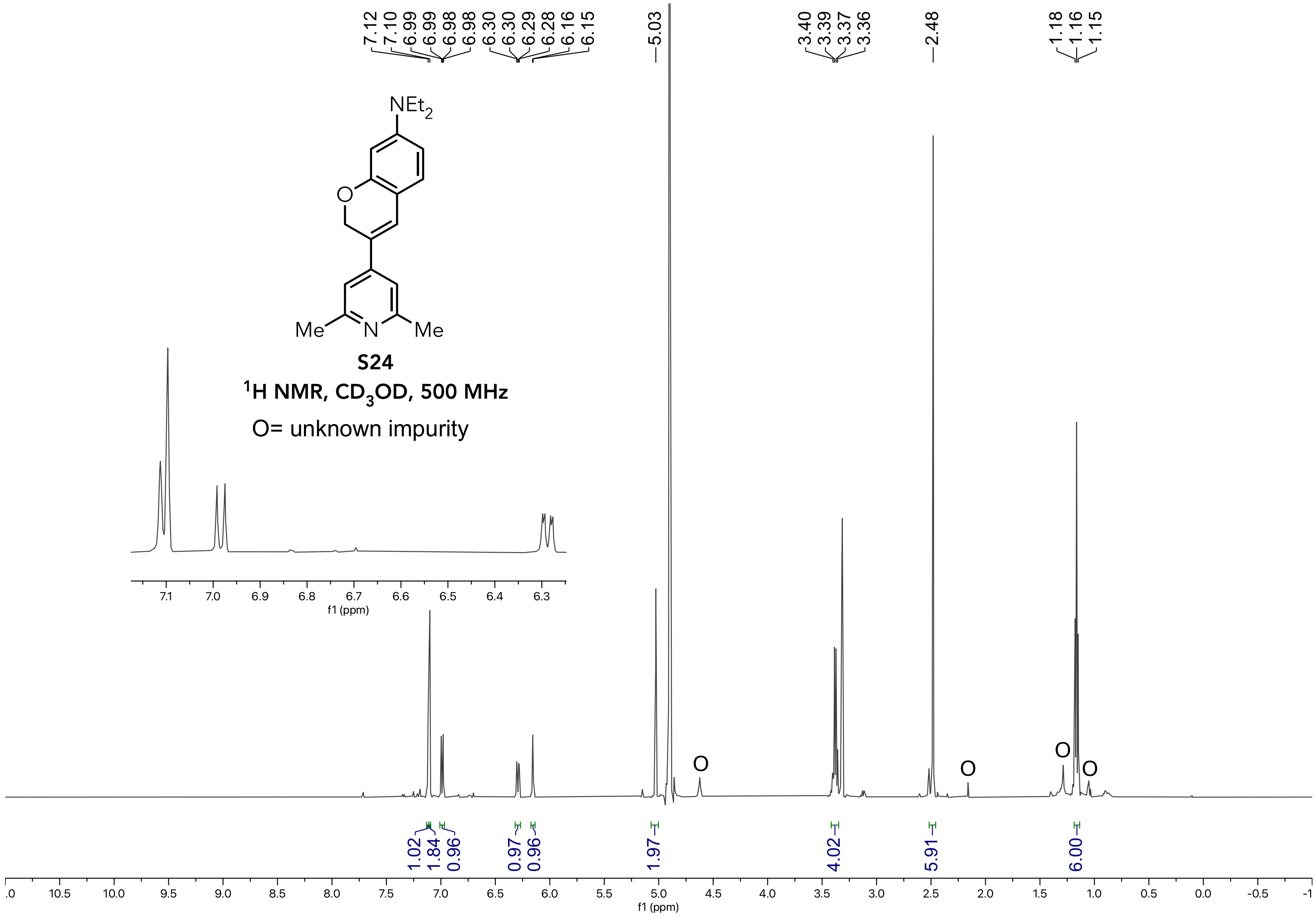
7.12
7.10
6.99
6.99
6.98
6.98
6.30
6.30
6.29
6.28
6.16
6.15
5.03
3.40
3.39
3.37
3.36
2.48
1.18
1.16
1.15
NEt2
O
Me
N
Me
S24
1H NMR, CD3OD, 500 MHz
O= unknown impurity
7.1
7.0
6.9
6.8
6.7
6.6
6.5
6.4
6.3
f1 (ppm)
O
O
O
O
1.02
1.84
0.96
0.97
0.96
1.97
4.02
5.91
6.00
.0
10.5
10.0
9.5
9.0
8.5
8.0
7.5
7.0
6.5
6.0
5.5
5.0
4.5
4.0
3.5
3.0
2.5
2.0
1.5
1.0
0.5
0.0
-0.5
-1
f1 (ppm)

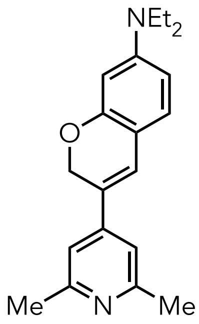


**S24**

**$^{13}$C NMR, $CD_3OD$, 126 MHz**

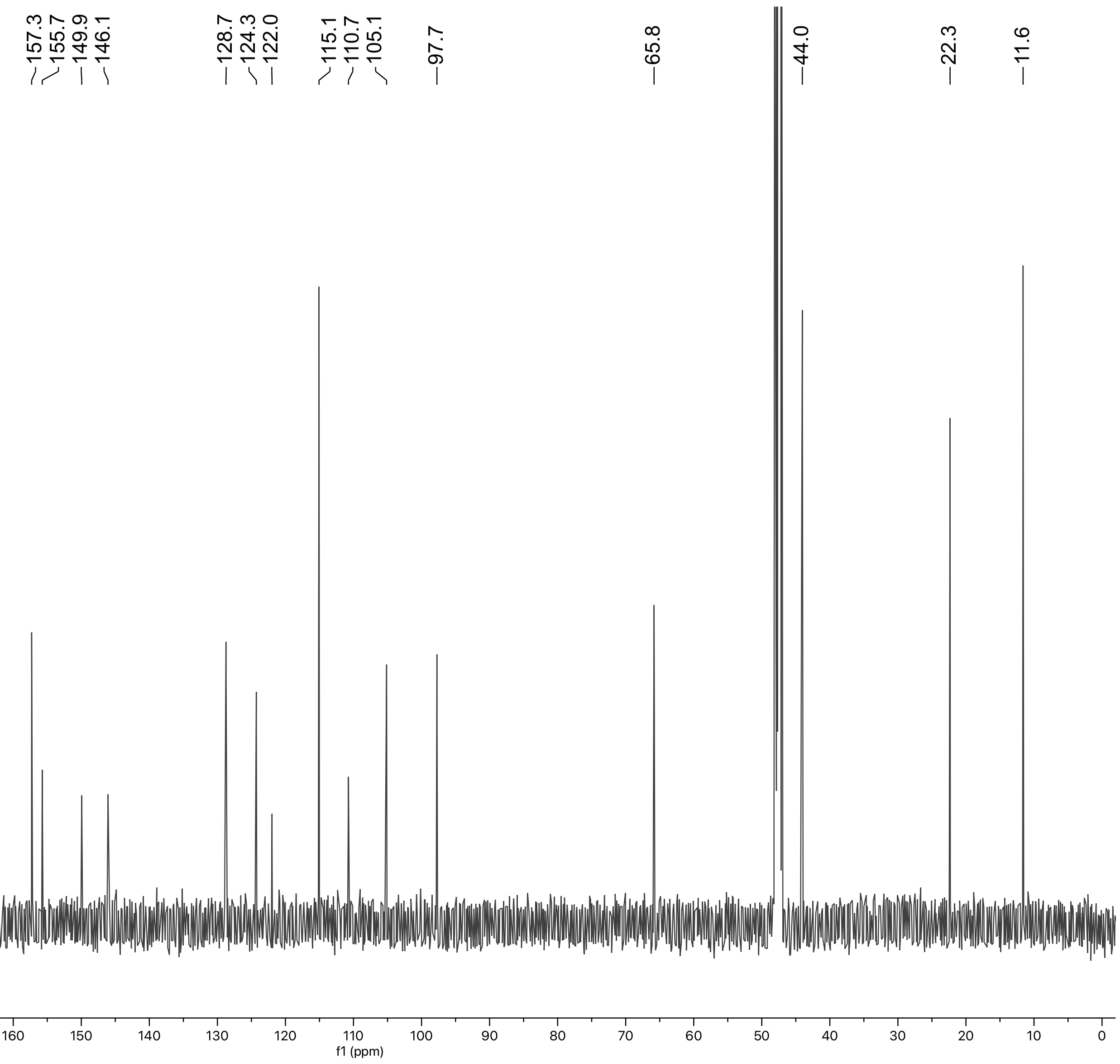

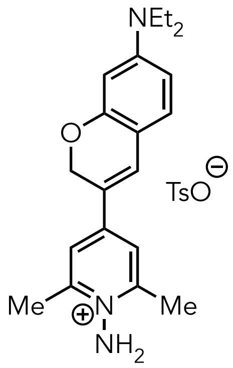


**S25**

**$^1$H NMR, $CD_3CN$, 500 MHz**

O= unknown impurity

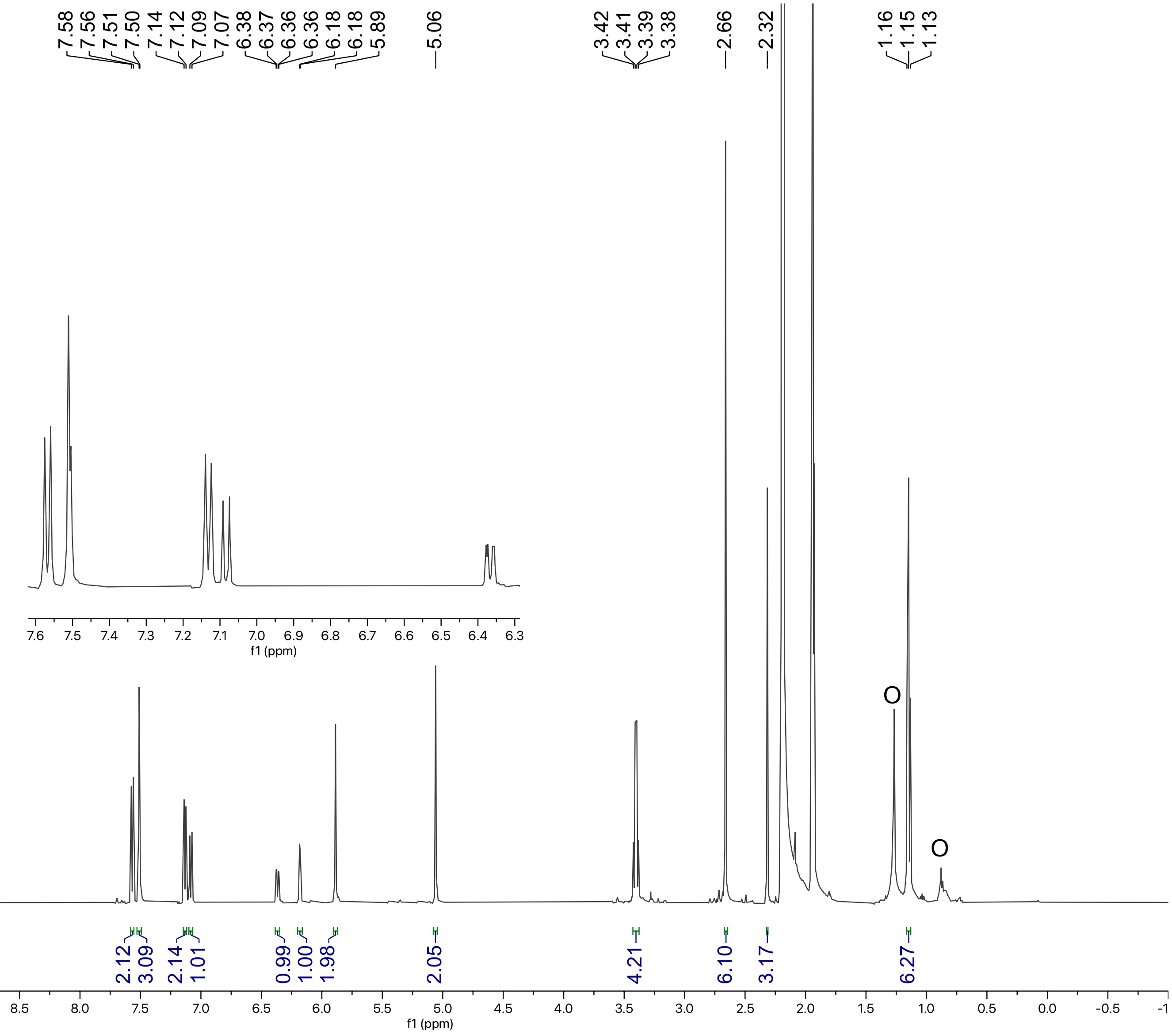

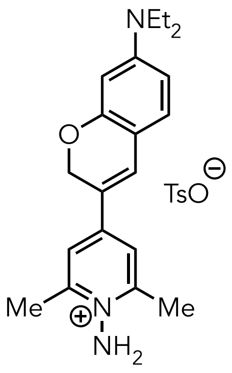

**S25**

**$^{13}$C NMR, $CD_3CN$, 126 MHz**

O= unknown impurity

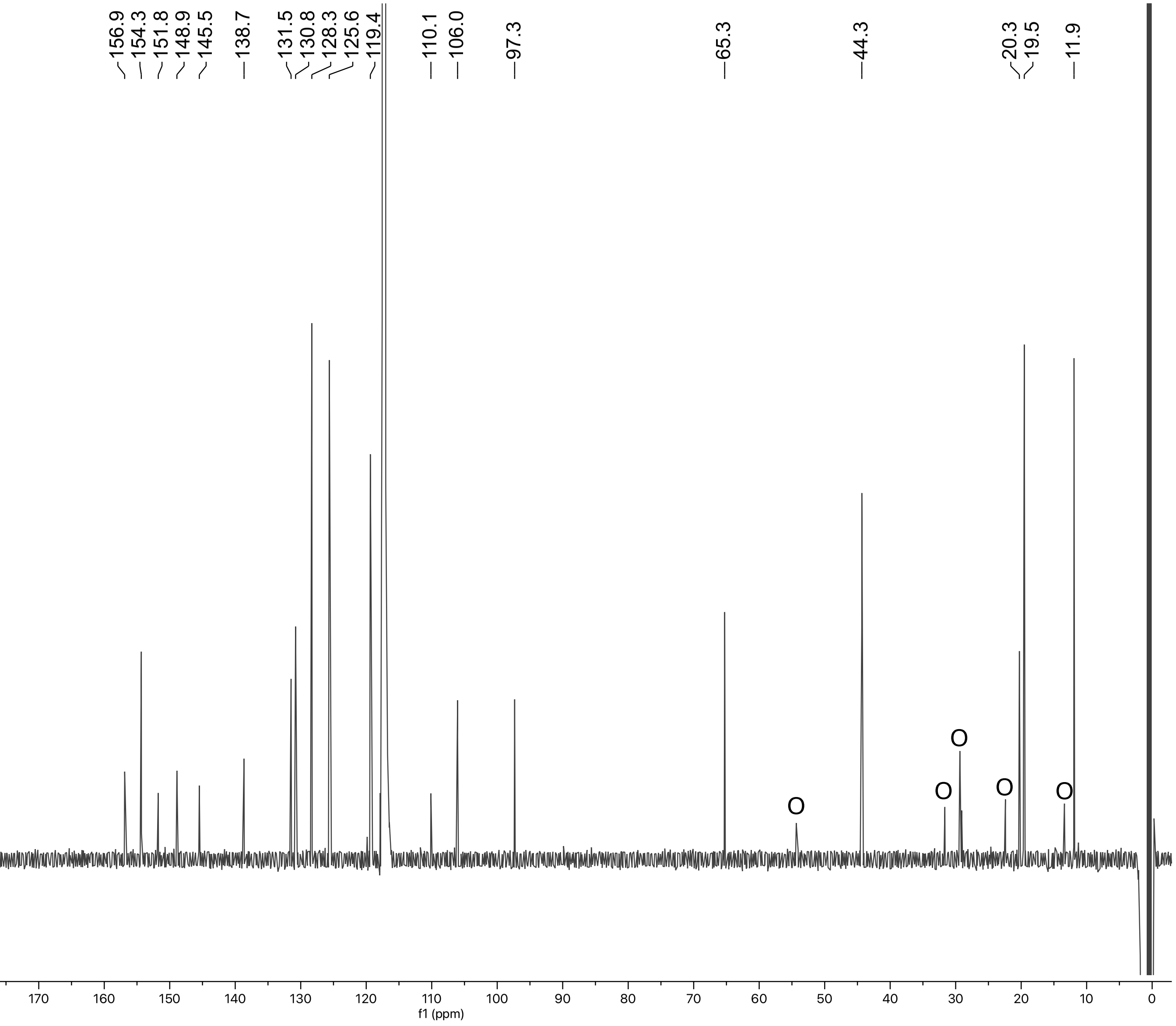

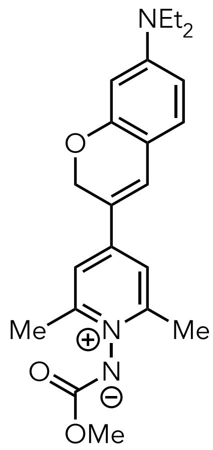


**S26**

**[1]H NMR, $CD_3CN$, 500 MHz**

O= unknown impurity

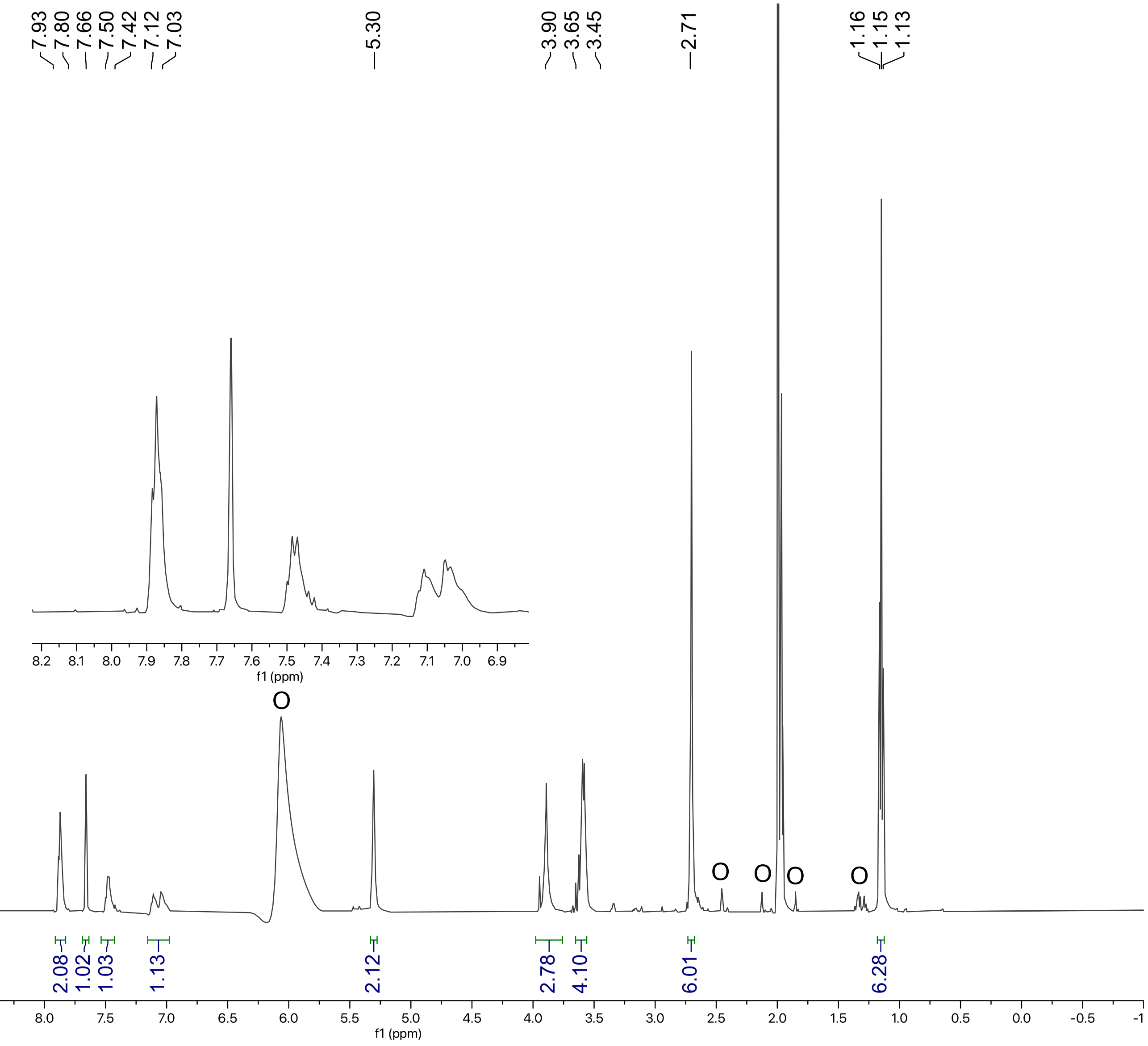

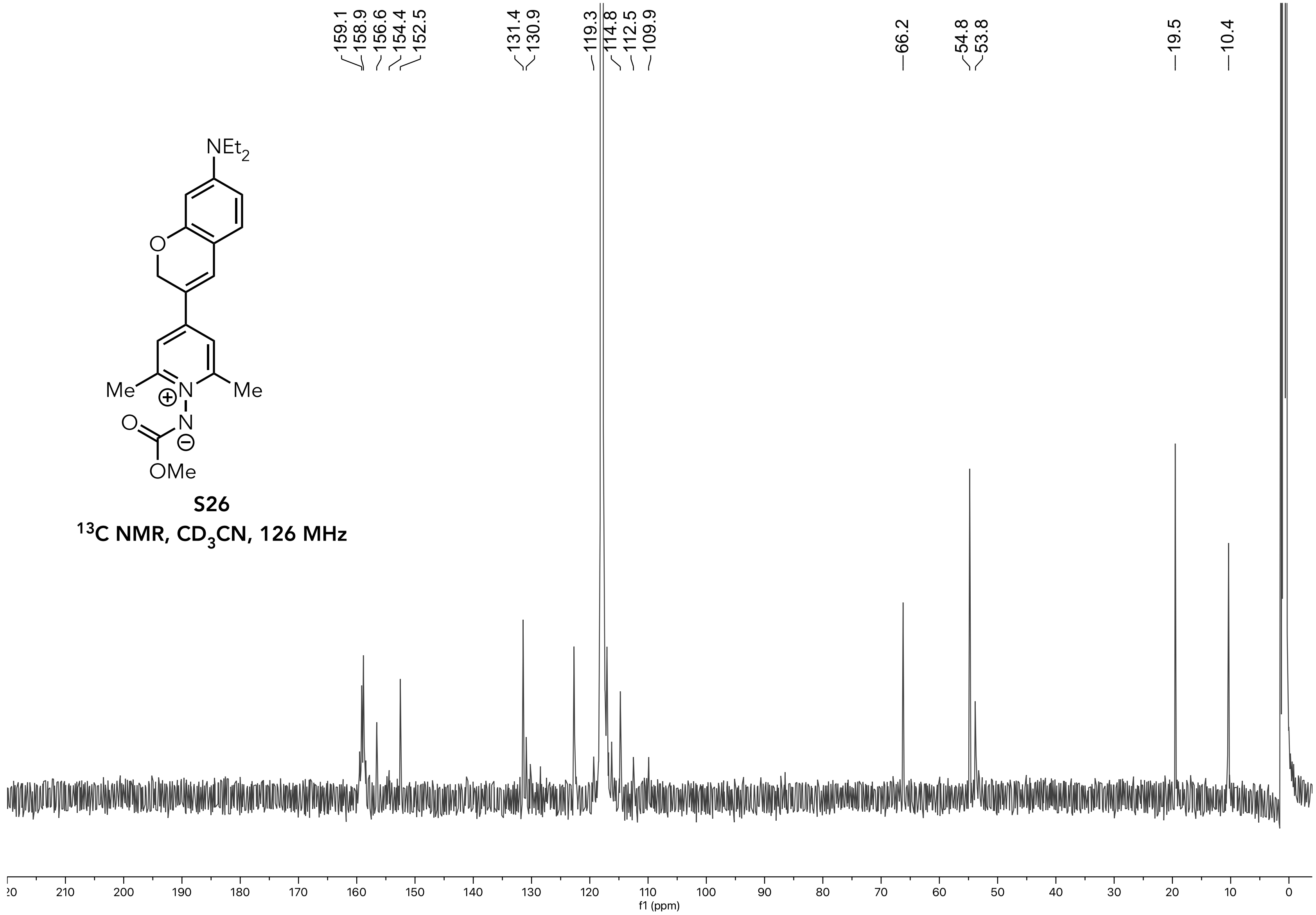


**S26**

$^{13}C$ NMR, $CD_3CN$, 126 MHz

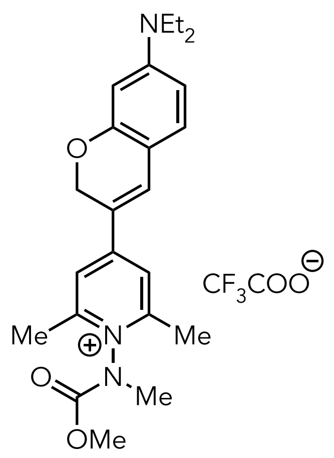


**10**

**[1]H NMR, $CD_3CN$, 500 MHz**

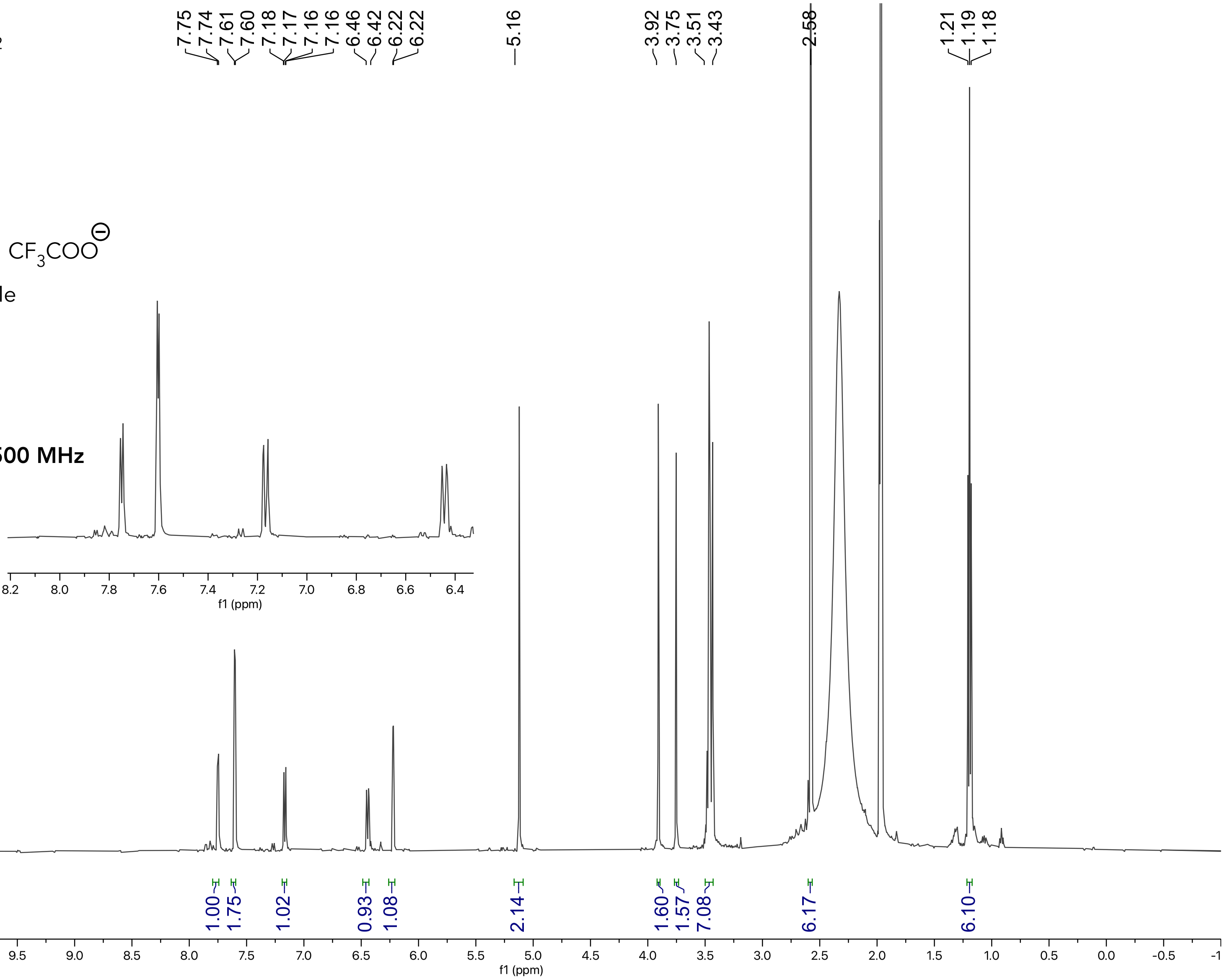

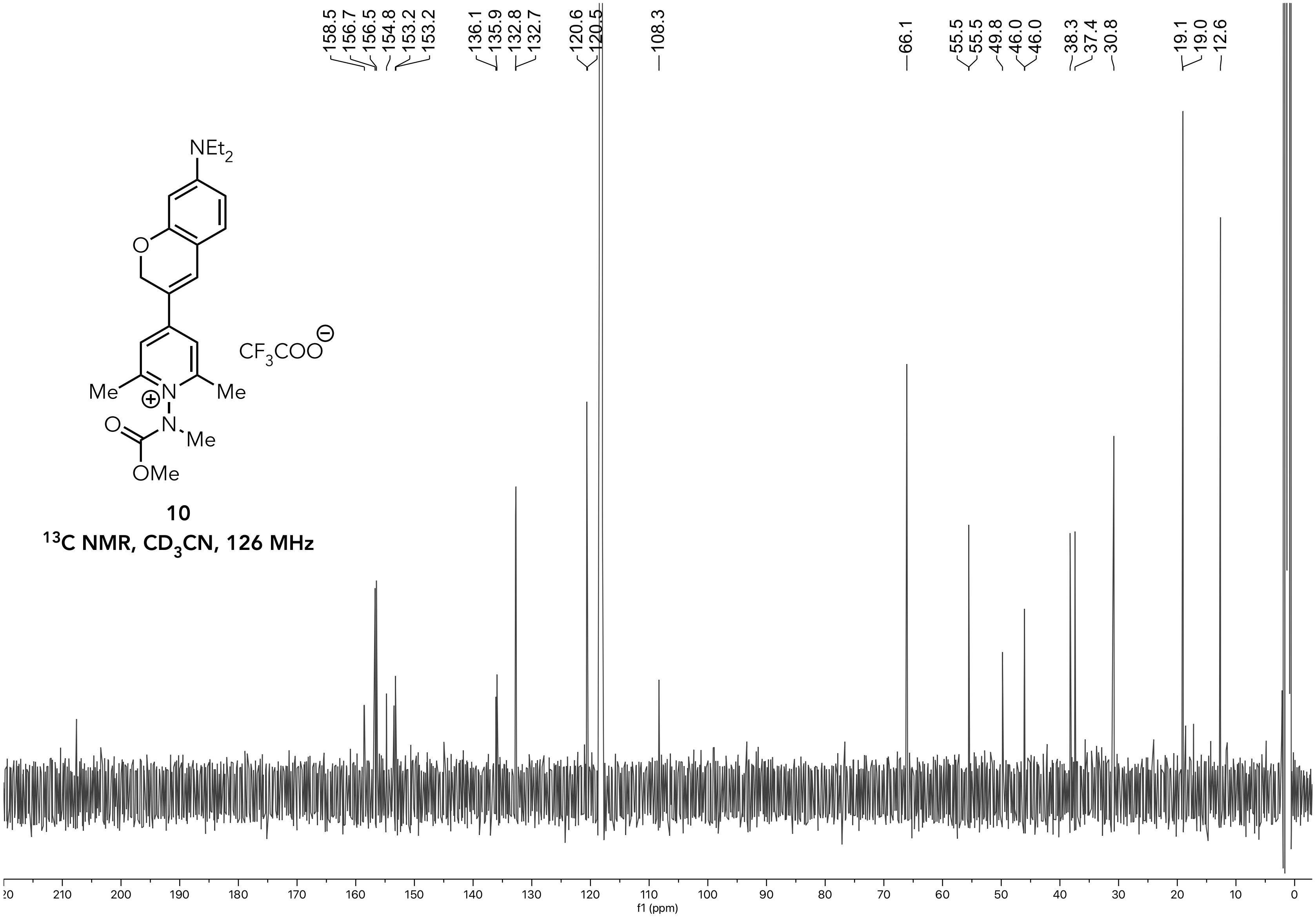


**10**

**$^{13}C$ NMR, $CD_3CN$, 126 MHz**

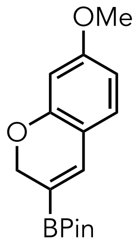


**S27**
$^1$H NMR, $CDCl_3$, 500 MHz
O= unknown impurity

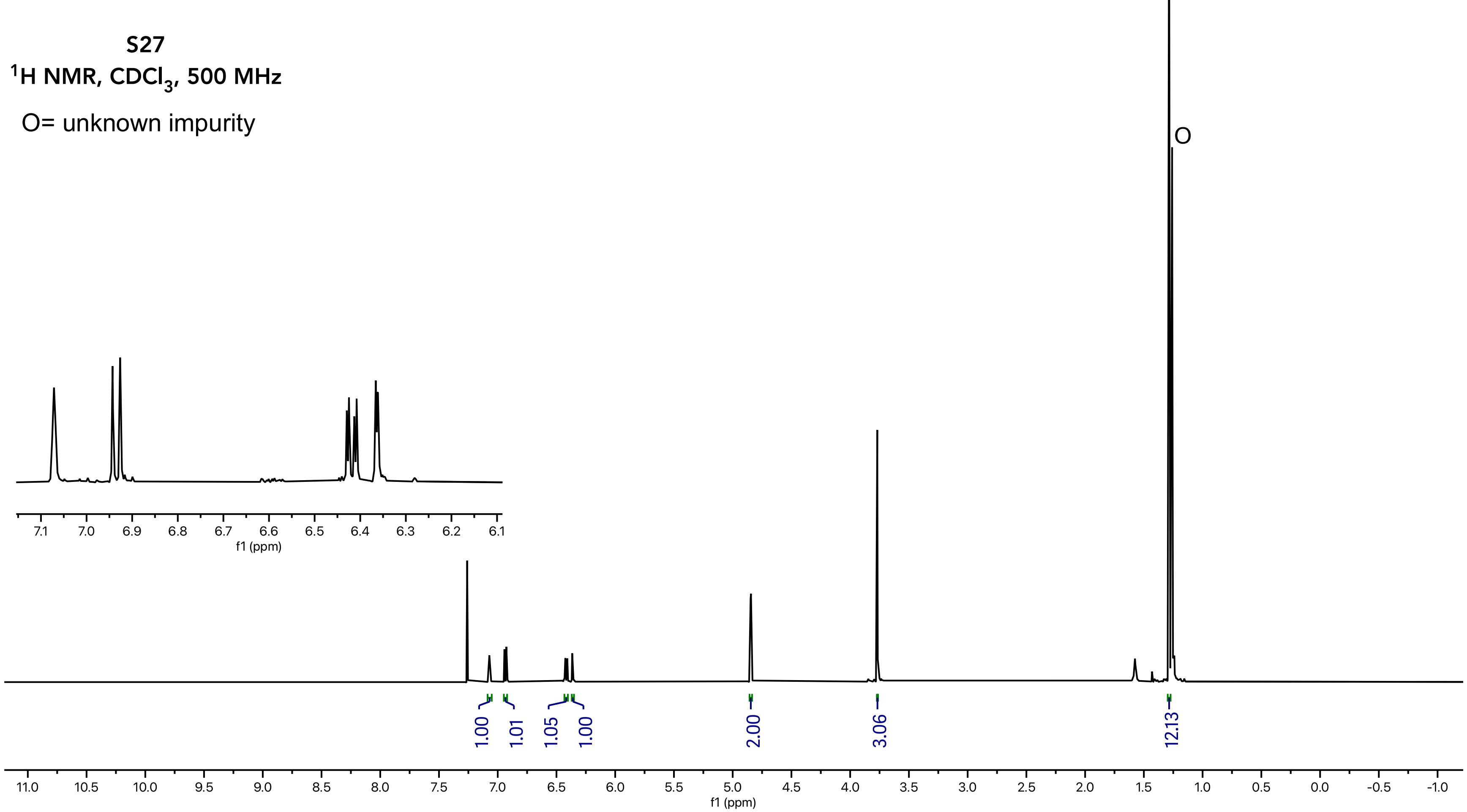

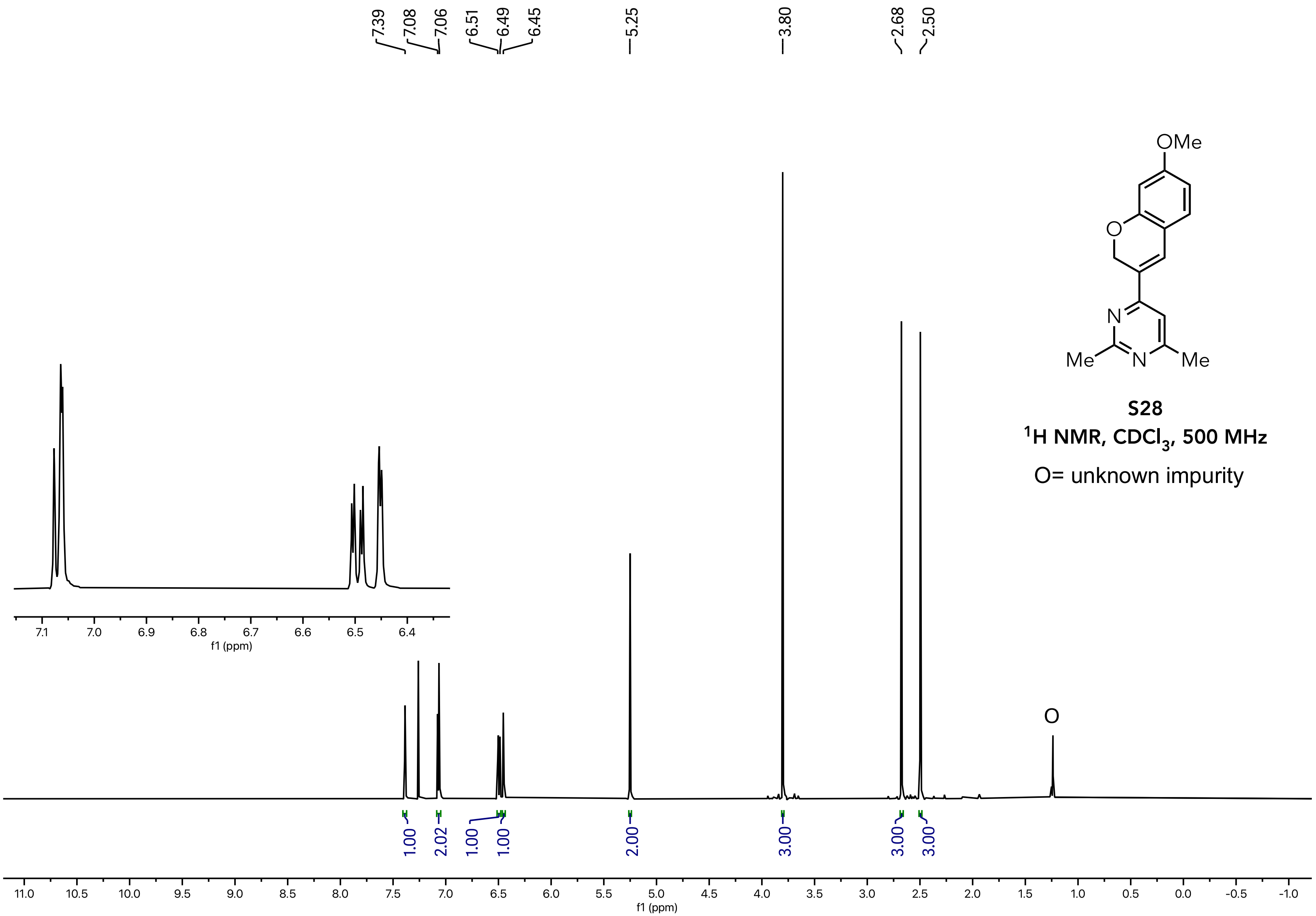


**S28**
**$^1$H NMR, $CDCl_3$, 500 MHz**
O= unknown impurity

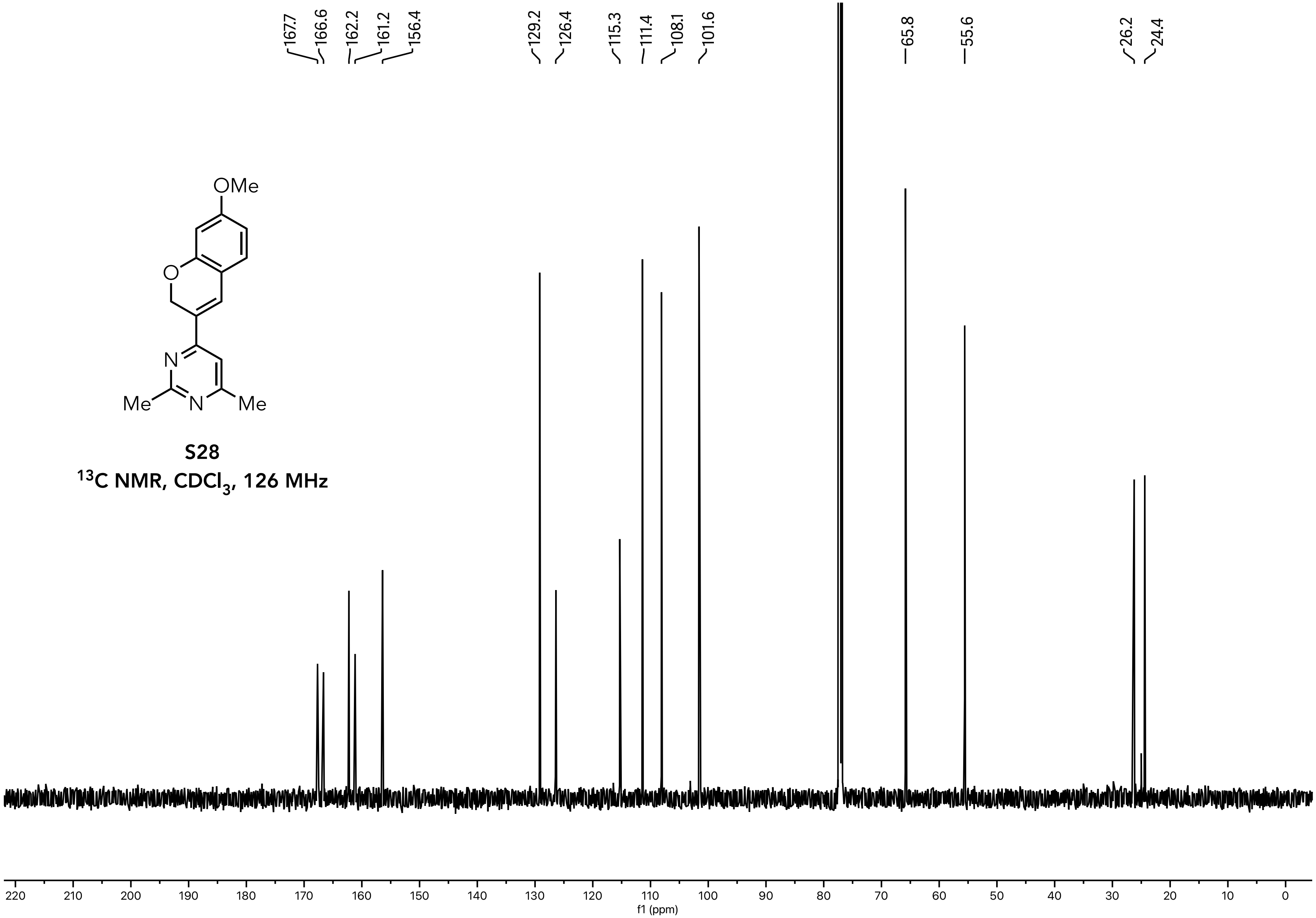


**S28**

**$^{13}C$ NMR, $CDCl_3$, 126 MHz**

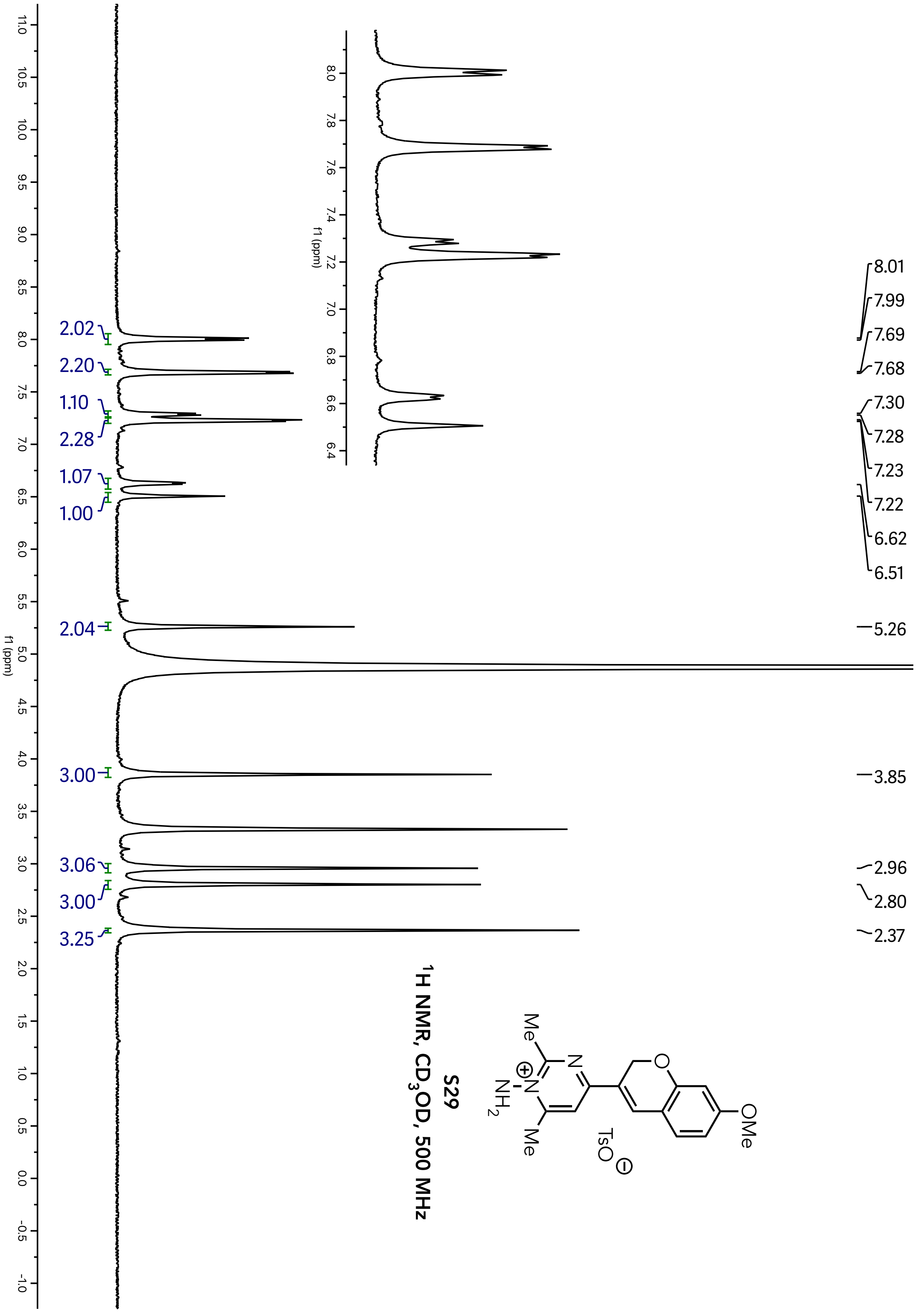

S29
$^1$H NMR, $CD_3OD$, 500 MHz
8.01
7.99
7.69
7.68
7.30
7.28
7.23
7.22
6.62
6.51
5.26
3.85
2.96
2.80
2.37
2.02
2.20
1.10
2.28
1.07
1.00
2.04
3.00
3.06
3.00
3.25
f1 (ppm)

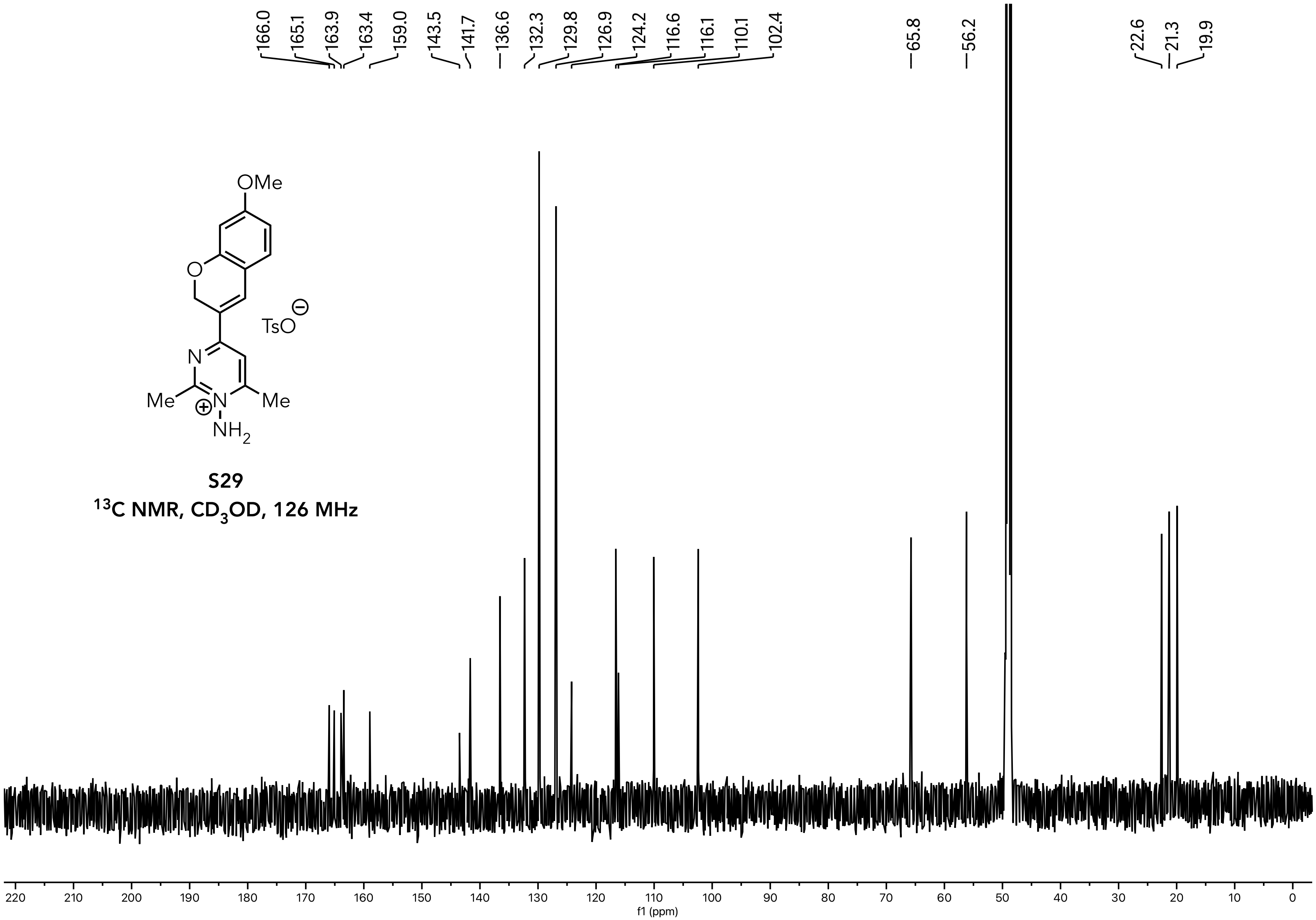


**S29**

**$^{13}$C NMR, $CD_3OD$, 126 MHz**

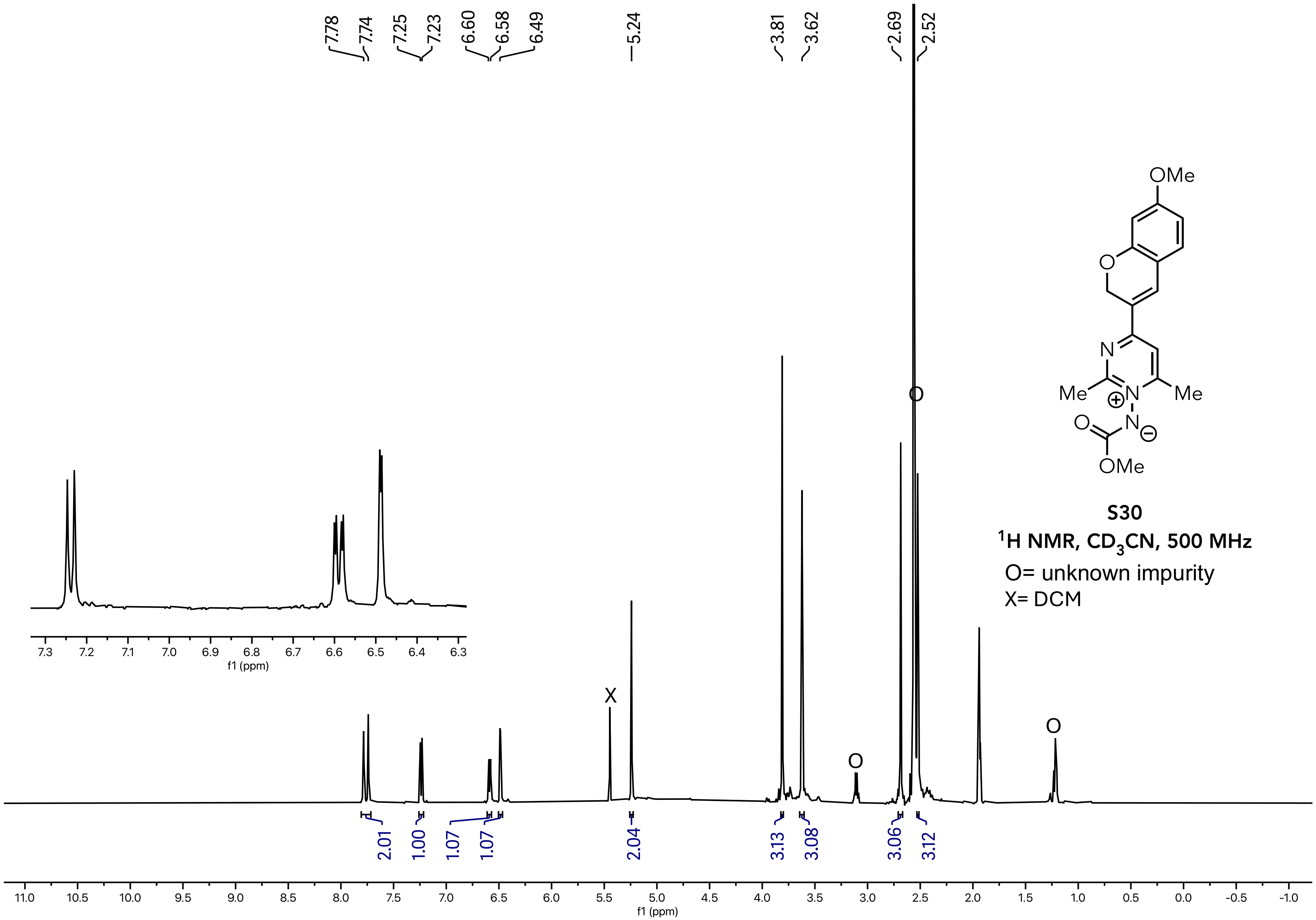


**S30**

**$^1$H NMR, $CD_3CN$, 500 MHz**

O= unknown impurity

X= DCM

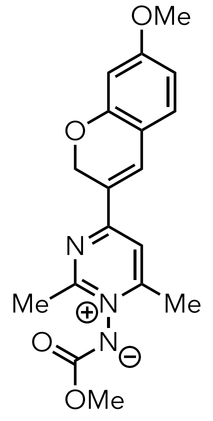


**S30**

**$^{13}C$ NMR, $CD_3CN$, 126 MHz**

O= unknown impurity

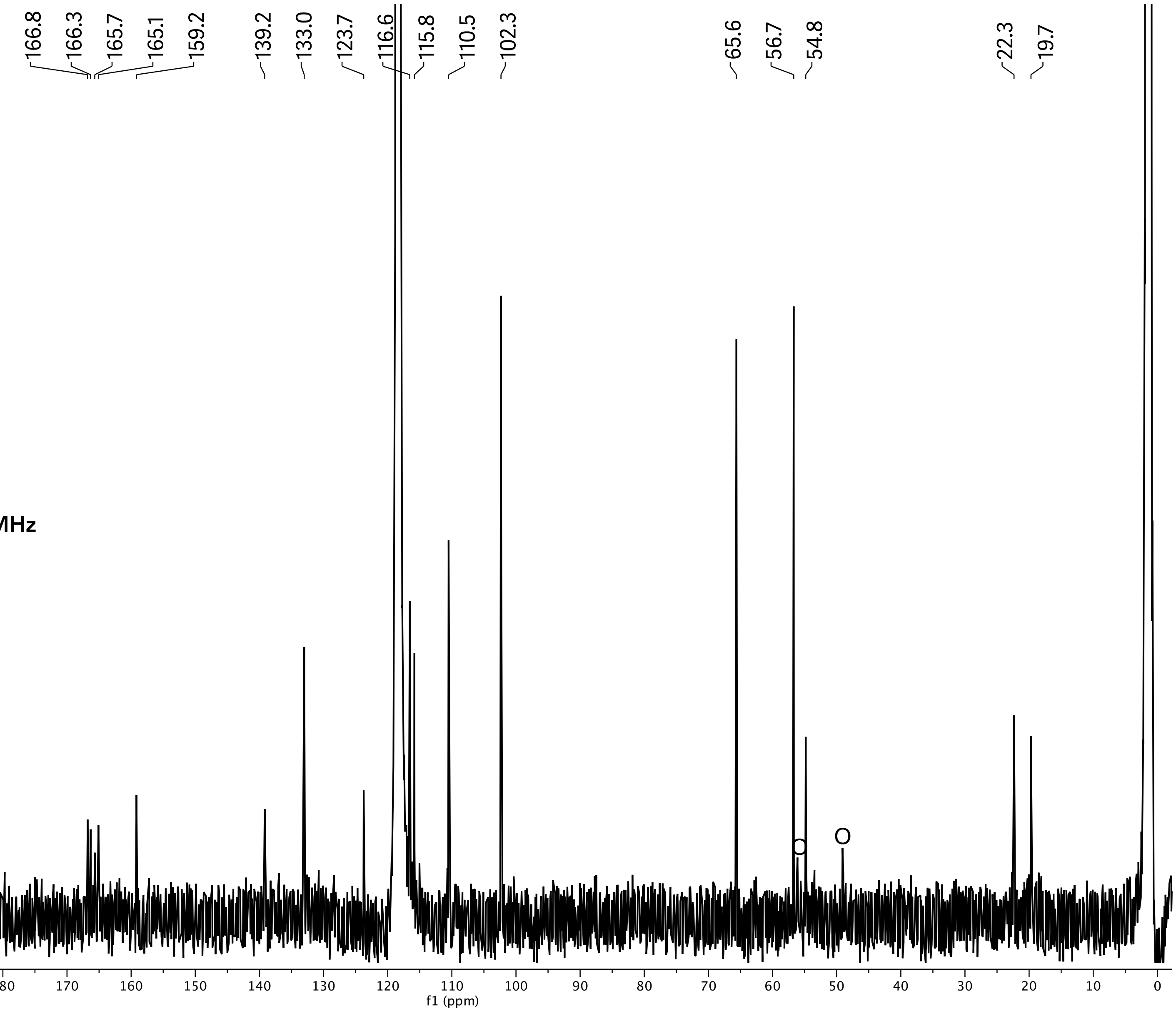

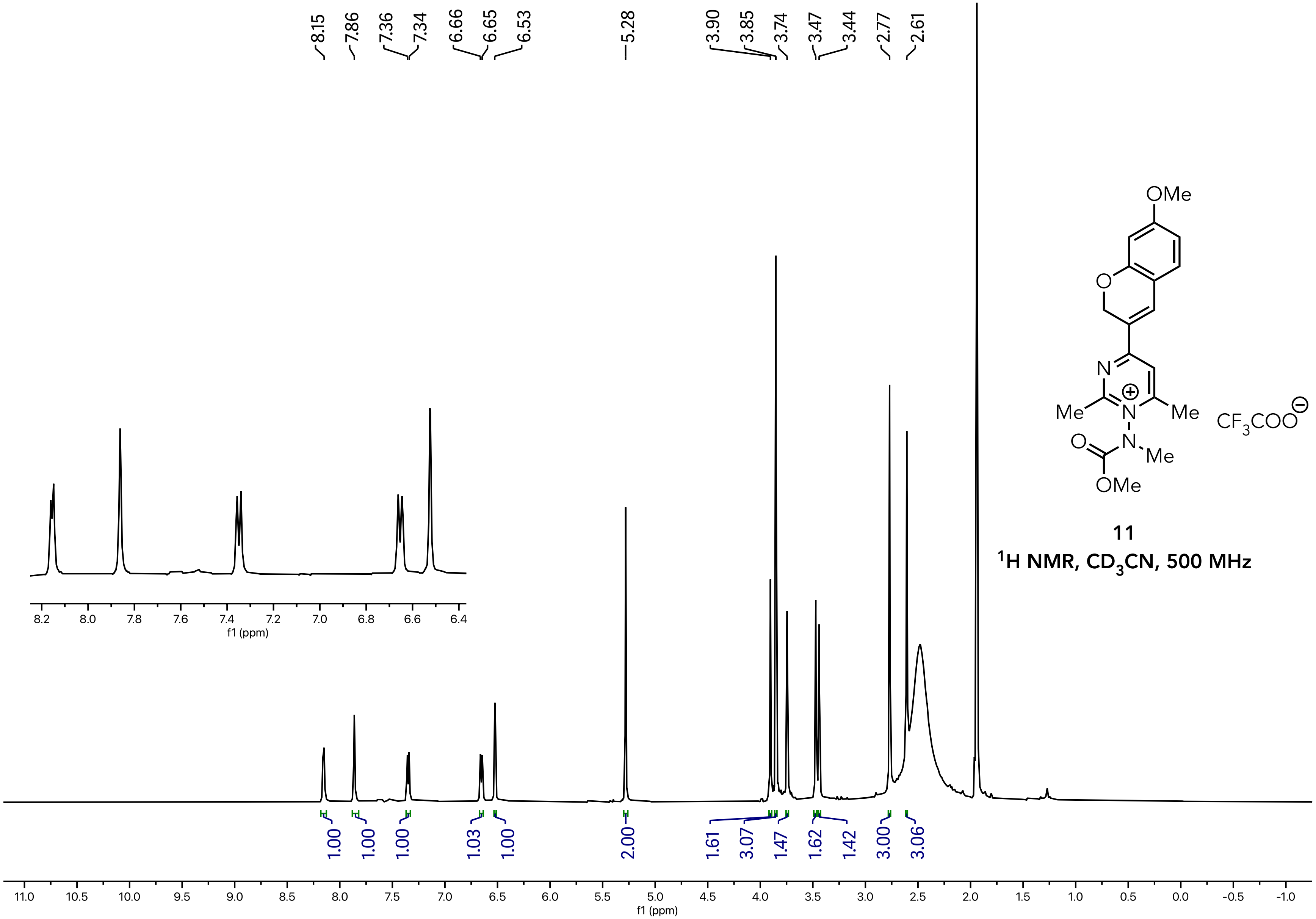


**11**

**$^1H$ NMR, $CD_3CN$, 500 MHz**

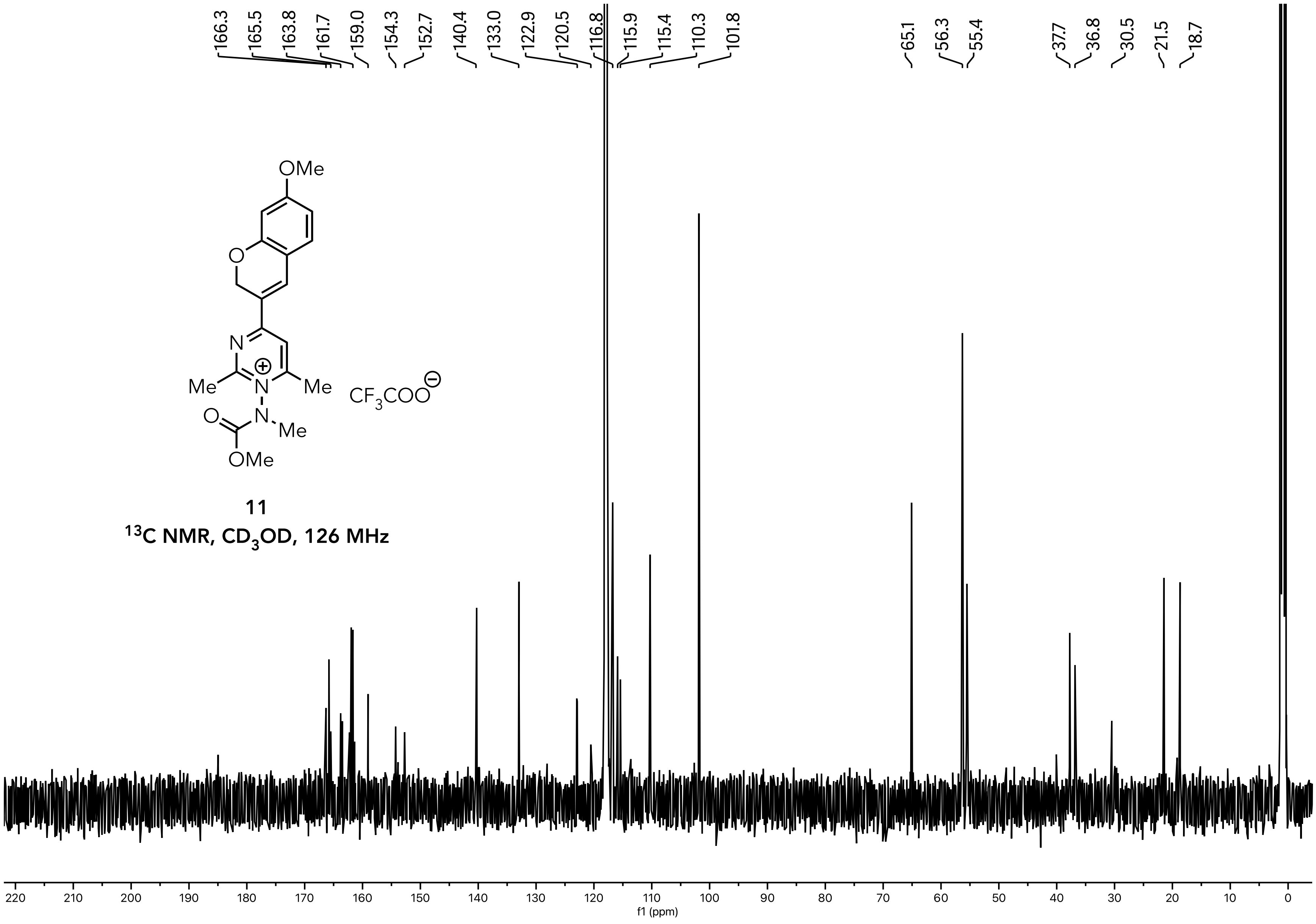


**11**

$^{13}C$ NMR, $CD_3OD$, 126 MHz

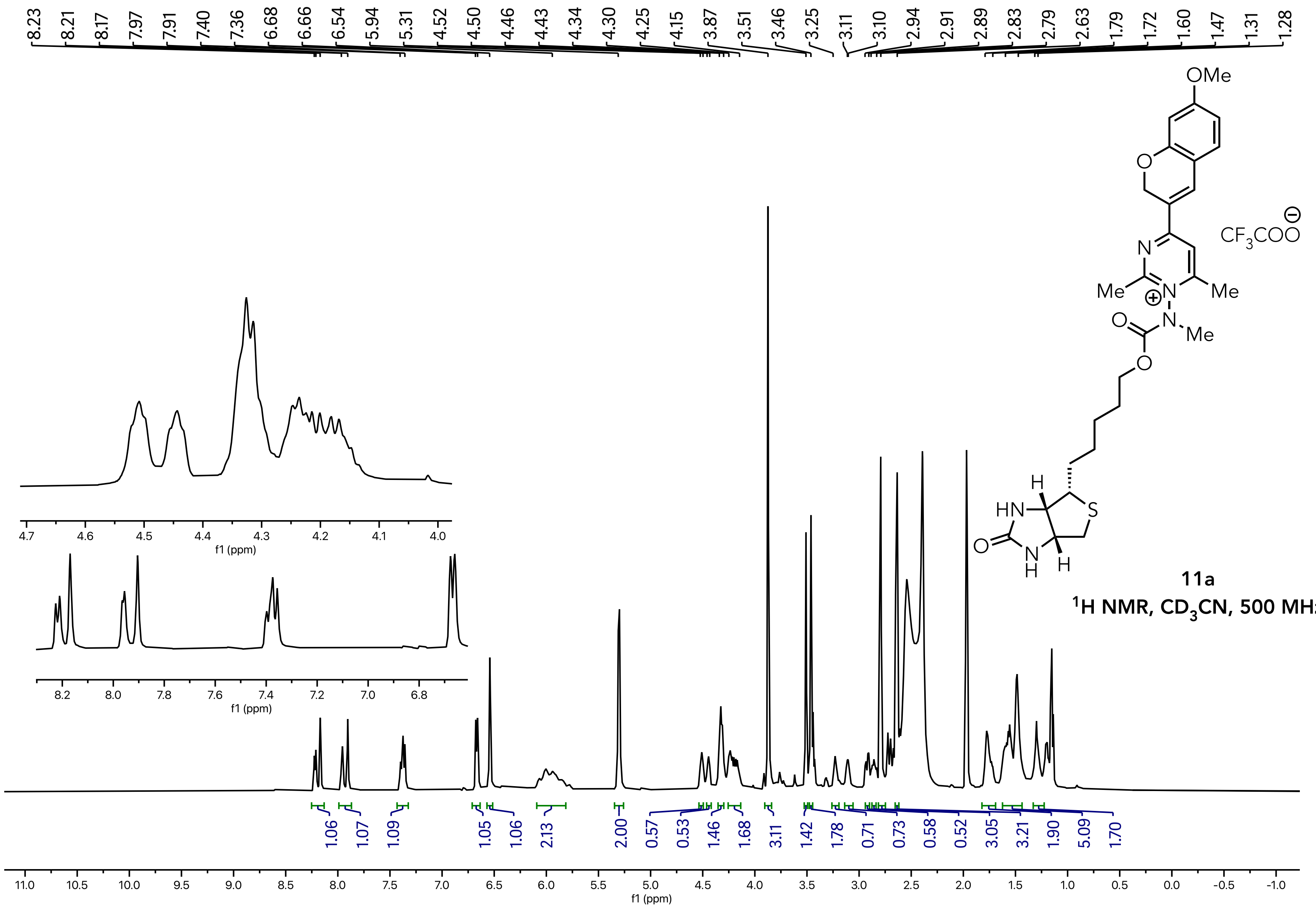


**11a**

**$^1$H NMR, $CD_3CN$, 500 MHz**

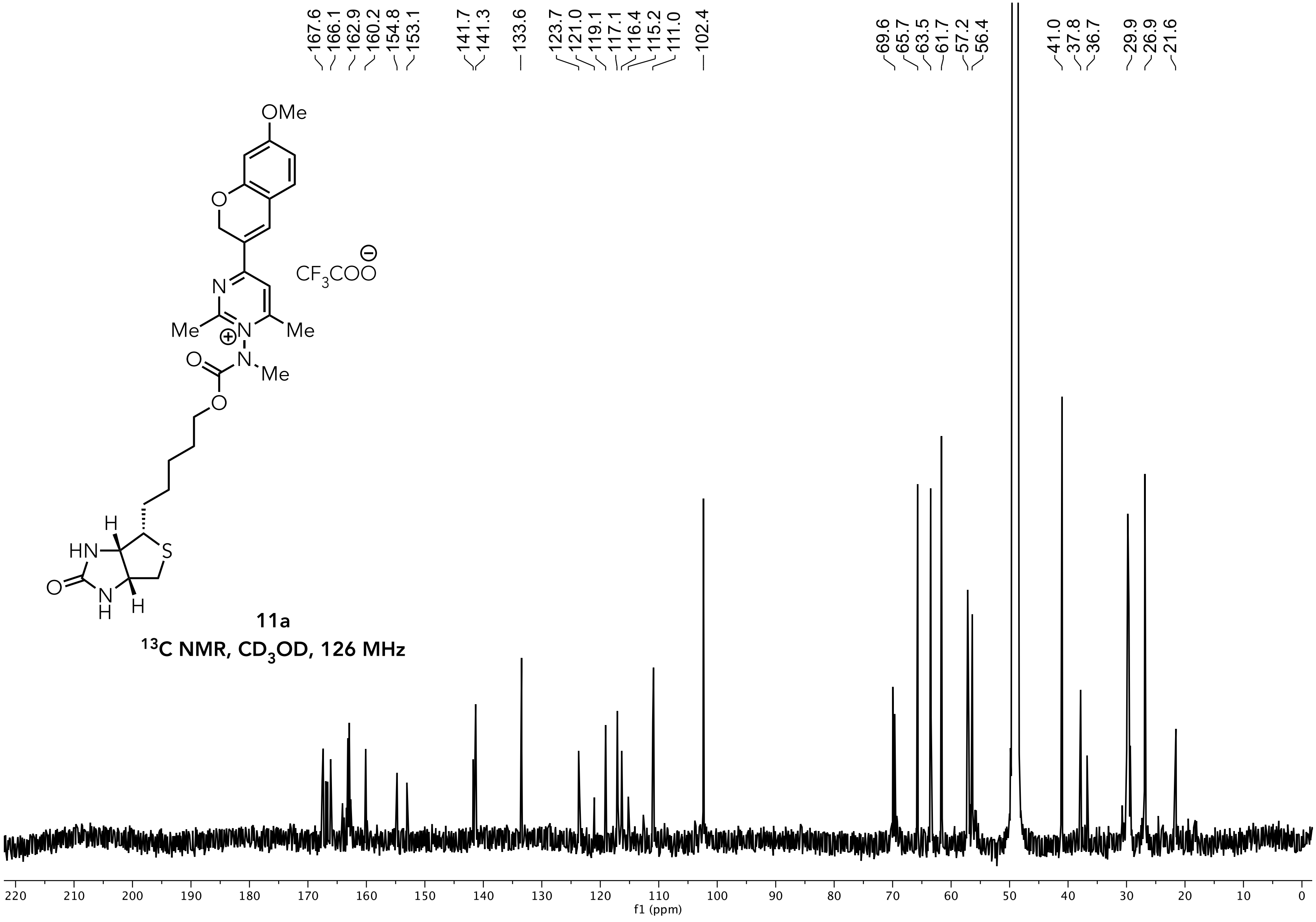


**11a**

**$^{13}C$ NMR, $CD_3OD$, 126 MHz**
