## Supplementary material for "Designer Aromatic Cations for Photo-Induced Protein Ligation, Imaging, and Intracellular Labelling at Extended Wavelengths": Computation Details

### Table of Contents

1. **Ground State Calculations of Probes 2-11**……………………………….………CSI2
2. **Estimation of the excited state oxidation potentials**…………………………CSI22
3. **Fluorophore Volume Calculations**…………………………..…………………CSI26
4. **Excited State Calculations** ..……………………………………………...…CSI26
5. **References**…………………………………………………………………...…CSI38

**Computational Details**

All calculations were performed using the Gaussian 16 computational chemistry software package[1]. Optimized structures were viewed and assessed in gaussview[2].

### 1. Ground State Calculations of Probes 2-11.

The ground state geometries of **2-11** were calculated using UMN12-SX/6-31G(d)/SMD=$H_2O$ level of theory. In each instance, frequency calculations were performed at the same level of theory and revealed no negative frequencies in any instance; confirming the calculated properties of **2-11** are from true energetic minima.

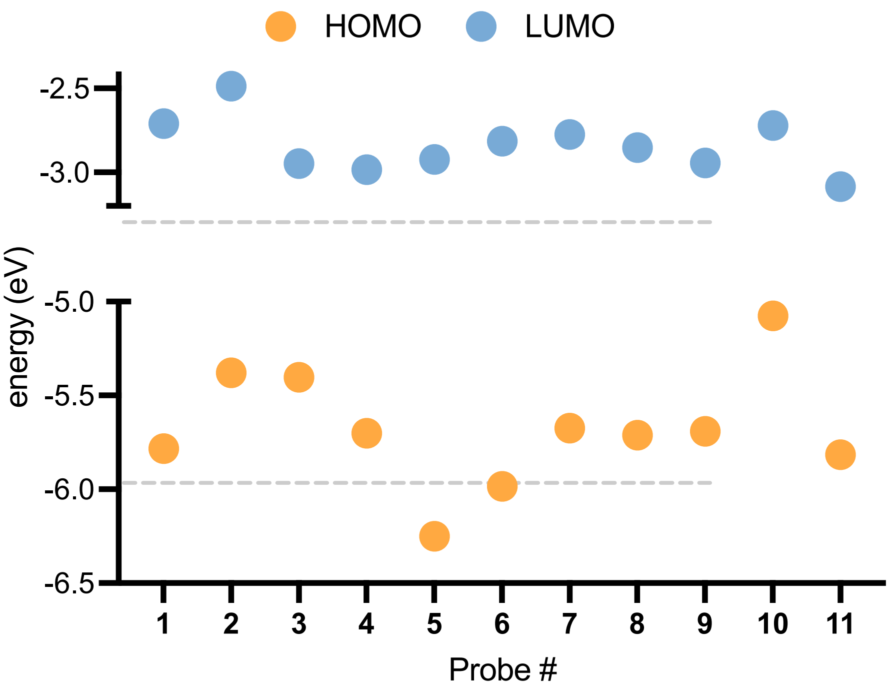

**Figure CSI1**. Calculated HOMO and LUMO energies derived from ground state calculations of probes **1-11**.

**Ground state geometry for probes 1-11**.
Geometry of **1**:

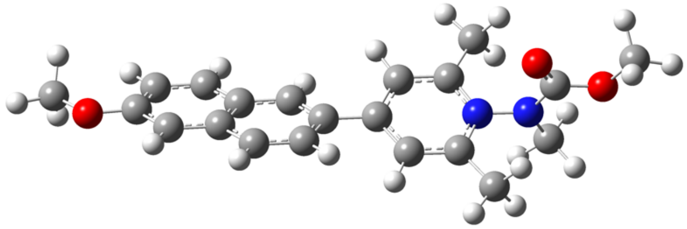

C,0,-3.3926867746,-1.4480171135,-1.743314606
C,0,-2.6602855277,-0.6989967967,-0.7897574333
C,0,-3.3793161361,0.0008849898,0.2293013872
C,0,-4.7842380224,-0.0664627871,0.2536982023
C,0,-5.467924137,-0.8051130767,-0.6938333662
H,0,-0.7263488588,-1.1932067711,-1.5927675991
C,0,-1.2512976942,-0.6299161018,-0.8186245745
C,0,-0.5477767064,0.0902404282,0.131930794
C,0,0.9195627847,0.1256978794,0.1072038823
C,0,1.634859346,-0.0373516001,-1.0902551399
C,0,3.0385359633,0.34163394,1.2689826126
C,0,3.0120841545,-0.01550646,-1.1121416838
H,0,1.1198235922,-0.1558237896,-2.0417519546
N,0,3.6698977964,0.1629506787,0.0717846132
C,0,3.8161304387,-0.1581880345,-2.3558269869
H,0,4.4088143242,0.7483174075,-2.5427843978
H,0,4.5089304532,-1.0076484671,-2.2878333056
H,0,3.1434403001,-0.3182841783,-3.2032807494
C,0,3.8694830613,0.5369338406,2.487778611
H,0,4.5466580253,-0.3128909546,2.6480334057
H,0,4.4808946078,1.4458976985,2.4023727947
H,0,3.2148310925,0.6363118636,3.3583498064
N,0,5.0597440237,0.1760404795,0.0541022838
C,0,5.6386237329,1.4027959567,-0.1199740354
O,0,5.0106303308,2.4372136306,-0.2562052293
C,0,5.7798968953,-1.0803164507,0.2271330279
H,0,5.0421095592,-1.8802279087,0.3432148889
H,0,6.4013105433,-1.2890525313,-0.652240553
H,0,6.4109499782,-1.039668355,1.1233845846
O,0,6.9597189085,1.2893272871,-0.1170847447

C,0,7.6756095065,2.51725358,-0.2892303791
H,0,8.7325861877,2.2419279222,-0.2622276774
H,0,7.4233187724,2.9708949094,-1.2543456778
H,0,7.4402317555,3.2100639541,0.5265036929
H,0,-5.3454313769,0.4629177905,1.0244175896
C,0,-1.2713659371,0.7814353808,1.1484763274
H,0,-0.7306656236,1.3770855535,1.8844824102
C,0,-4.7636609162,-1.5059092594,-1.7060016814
H,0,-2.8478487585,-1.98539418,-2.5215400856
H,0,-5.2990182129,-2.0883703253,-2.4532204922
C,0,-2.6377509088,0.7443693593,1.1887902188
H,0,-3.1796073868,1.2926275883,1.9610178523
C,0,1.6604639039,0.3222823239,1.2834110528
H,0,1.1682227201,0.4351751972,2.2474591095
O,0,-6.8202698378,-0.8127118206,-0.5894736471
C,0,-7.5618524891,-1.568636924,-1.5309671509
H,0,-8.6128870622,-1.4402371169,-1.254742337
H,0,-7.407388311,-1.1968213576,-2.5539498435
H,0,-7.3008560701,-2.6354169185,-1.4819216066

Zero-point correction= 0.409306 (Hartree/Particle)
Thermal correction to Energy= 0.433908
Thermal correction to Enthalpy= 0.434853
Thermal correction to Gibbs Free Energy= 0.353735
Sum of electronic and zero-point Energies= -1147.808516
Sum of electronic and thermal Energies= -1147.783914
Sum of electronic and thermal Enthalpies= -1147.782970
Sum of electronic and thermal Free Energies= -1147.864088

Geometry of **2**:

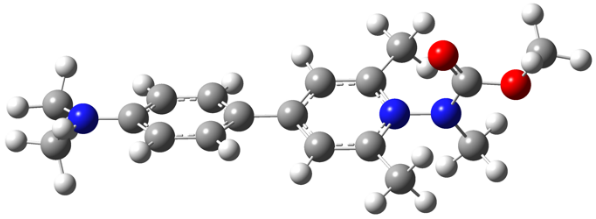

H,0,-0.6247198044,-1.3306552707,-1.2542116972
C,0,-1.1609556475,-0.710185974,-0.5352971292
C,0,-0.4475320463,0.1117310712,0.3598378156
C,0,1.0039176898,0.1925822647,0.3268161436
C,0,1.7396704938,-0.1662084276,-0.8217933991

C,0,3.1184174299,0.7043849978,1.4160833226
C,0,3.1120618772,-0.0963639426,-0.8533095294
H,0,1.2390708099,-0.4806751507,-1.7351060758
N,0,3.7648628028,0.3267045244,0.2716632838
C,0,3.9274255861,-0.4445534318,-2.0487346565
H,0,4.5025105676,0.4243790102,-2.3977332721
H,0,4.6378153054,-1.2515071719,-1.8233148732
H,0,3.2660265464,-0.7736415883,-2.8553865303
C,0,3.9407005679,1.1585032887,2.5703273967
H,0,4.6507947506,0.3816475524,2.8846243882
H,0,4.516602438,2.0573616355,2.3089615693
H,0,3.2840137281,1.397135191,3.4118618568
N,0,5.1517421676,0.390365096,0.246091965
C,0,5.6838138119,1.5783650962,-0.1697155449
O,0,5.0191670641,2.542637336,-0.5048461408
C,0,5.9223769955,-0.7772610924,0.6565575715
H,0,5.2181128285,-1.5618068378,0.9495559777
H,0,6.5413167552,-1.140511771,-0.1731660204
H,0,6.562584991,-0.5314538605,1.5126786911
O,0,7.0093449204,1.5181371745,-0.1519006396
C,0,7.6785410681,2.7113987472,-0.5728045081
H,0,8.7452165614,2.4872317492,-0.4963296894
H,0,7.4118696502,2.9501308553,-1.6086165323
H,0,7.4157878067,3.5467399401,0.086031555
C,0,-1.2059594506,0.8444429324,1.2945696948
H,0,-0.7098916793,1.5151361042,1.9970359569
C,0,1.7460688144,0.6340687513,1.4417733509
H,0,1.2546175896,0.9048798568,2.3737273527
C,0,-2.5818589393,0.7775857865,1.330608476
C,0,-3.2962443772,-0.04897707,0.4262256658
C,0,-2.535436781,-0.8039246839,-0.5025566672
H,0,-3.1138542285,1.3778059228,2.065342726
H,0,-3.0284241657,-1.478575372,-1.1988876075
N,0,-4.6561551903,-0.1105113704,0.4453802463
C,0,-5.3427423743,-1.0889832809,-0.3755594592
H,0,-6.4209774611,-0.9841431253,-0.2278687552
H,0,-5.1339909921,-0.9300598383,-1.4423424871
H,0,-5.0564298574,-2.1217374539,-0.1191583322
C,0,-5.3890209328,0.5280829525,1.5215821506
H,0,-5.2158403097,1.6129520875,1.5322733318
H,0,-6.4592309837,0.3634994501,1.3703020006

H,0,-5.1118658269,0.1261375598,2.5093468375

Zero-point correction= 0.403602 (Hartree/Particle)
Thermal correction to Energy= 0.427296
Thermal correction to Enthalpy= 0.428240
Thermal correction to Gibbs Free Energy= 0.349631
Sum of electronic and zero-point Energies= -1013.701205
Sum of electronic and thermal Energies= -1013.677511
Sum of electronic and thermal Enthalpies= -1013.676567
Sum of electronic and thermal Free Energies= -1013.755176

Geometry of **3**:

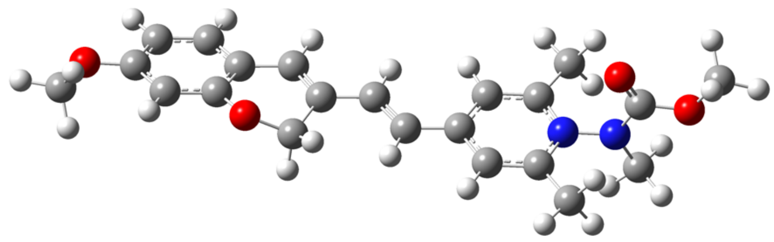

C,0,-4.6028950003,-1.1567194973,-2.2076875797
C,0,-3.4126871479,-0.6926943859,-1.6205700497
C,0,-2.7273310413,-1.5596803293,-0.7486390159
C,0,-3.2018573802,-2.8310698445,-0.4536870178
C,0,-4.3817399666,-3.2618972074,-1.0643902196
H,0,-3.5238298961,1.3843798081,-2.2864644272
C,0,-2.8992639966,0.6365981395,-1.7911128167
C,0,0.6043736828,4.0457493094,-1.371606607
C,0,-0.1347037054,5.0890893132,-1.9688839079
C,0,2.4666870375,5.5940254723,-1.1379732244
C,0,0.408981042,6.3371825875,-2.1505600225
H,0,-1.1595761913,4.9411012237,-2.302811382
N,0,1.6980781057,6.5524249011,-1.7395965097
C,0,-0.3292425743,7.4718936682,-2.7702076454
H,0,-0.4286985045,8.3071713326,-2.0629817759
H,0,0.190574066,7.8450823443,-3.6629560149
H,0,-1.3296475917,7.1372545118,-3.0596094074
C,0,3.8455039109,5.9571633935,-0.712489467
H,0,4.44526276,6.3097354962,-1.5623680701
H,0,3.8247888011,6.7561987387,0.0417984995
H,0,4.331669613,5.0789812538,-0.2778743105
N,0,2.2480104094,7.8131439261,-1.9305691399

C,0,2.0614328001,8.7044003511,-0.9114354439
O,0,1.4543012258,8.4503482422,0.1135493262
C,0,2.9662455113,8.0912494435,-3.1683829057
H,0,2.943482995,7.1862194667,-3.7829551706
H,0,2.4837486073,8.9101150015,-3.7159891977
H,0,4.0094408643,8.3556988242,-2.9558884835
O,0,2.6332608239,9.8648905499,-1.2063565639
C,0,2.503374942,10.8775723289,-0.203105087
H,0,3.0209806787,11.7516194569,-0.6056022906
H,0,1.4458120797,11.106374747,-0.0296745098
H,0,2.9762369802,10.5497943085,0.729598129
H,0,-2.6415222428,-3.4531401163,0.2404160524
C,0,-0.825196637,-0.1324747895,-0.7295974675
H,0,-0.1629628141,0.2396686367,0.0603962879
H,0,-0.1968533349,-0.6076255937,-1.5043842764
C,0,-5.0823397517,-2.4233475998,-1.948364586
O,0,-1.6136532709,-1.1393648219,-0.0924508124
H,0,-5.1496254376,-0.4925563281,-2.8791433881
C,0,1.9191987728,4.3456318024,-0.9607736371
H,0,2.5311497231,3.5785904521,-0.487701149
C,0,-1.1447486746,2.28833712,-1.5185904456
H,0,-1.8332126932,2.9821529864,-2.0103018508
C,0,0.0974083111,2.7135000458,-1.1614306635
H,0,0.796305031,2.0242818699,-0.6835065765
C,0,-1.6609299793,0.9671858993,-1.3342479455
O,0,-4.9237577563,-4.4834864608,-0.8631602325
C,0,-4.2593294126,-5.3687145906,0.0221261574
H,0,-4.2037095356,-4.9482508872,1.0362553108
H,0,-4.8605301534,-6.2825986335,0.0391460453
H,0,-3.2471349976,-5.6029192907,-0.3372086962
H,0,-5.9995066671,-2.7919939667,-2.405004657

Zero-point correction= 0.442128 (Hartree/Particle)
Thermal correction to Energy= 0.469167
Thermal correction to Enthalpy= 0.470111
Thermal correction to Gibbs Free Energy= 0.383347
Sum of electronic and zero-point Energies= -1262.188474
Sum of electronic and thermal Energies= -1262.161434
Sum of electronic and thermal Enthalpies= -1262.160490
Sum of electronic and thermal Free Energies= -1262.247255

Geometry of **4**:

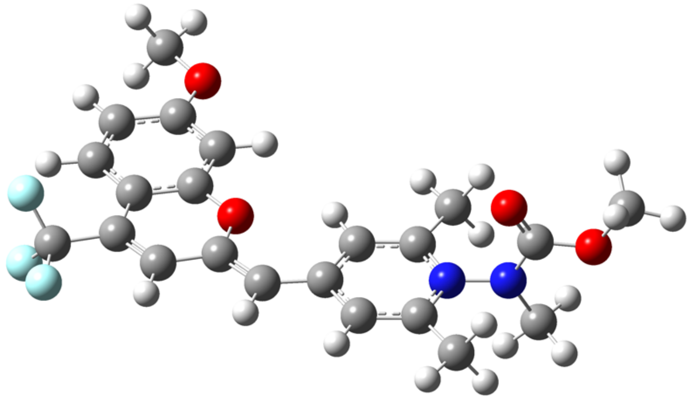

C,0,0.3688346562,3.9088059517,-0.7263800629
C,0,0.5866018606,4.252187995,-2.0819472516
C,0,1.8233989838,5.7654206318,-0.1030886327
C,0,1.393078106,5.3101923286,-2.4286011317
H,0,0.1249572416,3.6969676398,-2.8916993802
N,0,1.9967130454,6.0319512455,-1.435633561
C,0,1.6470562706,5.7111824441,-3.8399105943
H,0,1.2985234833,6.7365441415,-4.0259501388
H,0,2.7187377803,5.6715359737,-4.0779055936
H,0,1.112299997,5.0323079332,-4.5103508071
C,0,2.5167400431,6.634049643,0.8870297526
H,0,3.603817606,6.6288392682,0.7302987751
H,0,2.1688582561,7.6737562833,0.8100105863
H,0,2.3054322813,6.27017549,1.8968327299
N,0,2.8074178342,7.0993273501,-1.7969740813
C,0,2.1889223269,8.3156635845,-1.8720378602
O,0,0.9993405462,8.4886808841,-1.6746098255
C,0,4.2306496794,6.8670661035,-2.0092180488
H,0,4.8282284416,7.4059828898,-1.2631146383
H,0,4.4160661605,5.7931277886,-1.9114483456
H,0,4.5217886619,7.1905027806,-3.0158478429
O,0,3.0676079618,9.2585189753,-2.1885899008
C,0,2.5281145405,10.579440475,-2.3004276589
H,0,3.3764571494,11.2174939955,-2.5594929901
H,0,1.769371287,10.6138152079,-3.0905420467
H,0,2.0921488426,10.8931929527,-1.3452042475
C,0,1.0184669419,4.7105019564,0.2428089743
H,0,0.8838985125,4.4957722161,1.3021447749

C,0,-0.4442596456,2.8342634896,-0.2510517734
H,0,-0.5049970576,2.724214885,0.8310489758
C,0,-1.1662694189,1.9090032425,-0.9569769573
C,0,-1.9339049137,0.8893288334,-0.3045529947
C,0,-1.864384378,1.0512584545,-3.0656244898
C,0,-2.6387476717,-0.0088827654,-1.0285198174
H,0,-1.9211899076,0.8736464058,0.7833901873
C,0,-2.6349987533,0.0369345785,-2.4687764865
O,0,-1.1619398659,1.9456455505,-2.3096207894
C,0,-3.3248976694,-0.8346543829,-3.3246528175
H,0,-3.9354795146,-1.6363647092,-2.9110544678
C,0,-2.468447703,0.3196927443,-5.2583927604
C,0,-3.2505004178,-0.704059422,-4.7002628453
H,0,-3.7991718563,-1.3988802173,-5.3316602075
C,0,-1.7708036648,1.2045390175,-4.4336822412
H,0,-1.1643608005,2.0001783672,-4.8629938037
C,0,-3.4380079811,-1.0747999301,-0.3274663886
F,0,-3.0368186186,-2.3015287631,-0.6898634761
F,0,-3.3348599656,-0.9982170367,0.9991740004
F,0,-4.740207919,-0.9889045654,-0.6334607784
O,0,-2.3289964362,0.5277278929,-6.5831707034
C,0,-3.0130740917,-0.3424492032,-7.4710141873
H,0,-4.1005998612,-0.2807700733,-7.3253930252
H,0,-2.7594964769,0.0010607789,-8.4781111995
H,0,-2.6784404088,-1.3813039116,-7.3426625757

Zero-point correction= 0.418095 (Hartree/Particle)
Thermal correction to Energy= 0.447393
Thermal correction to Enthalpy= 0.448337
Thermal correction to Gibbs Free Energy= 0.355894
Sum of electronic and zero-point Energies= -1559.807061
Sum of electronic and thermal Energies= -1559.777763
Sum of electronic and thermal Enthalpies= -1559.776819
Sum of electronic and thermal Free Energies= -1559.869262

Geometry of **5**:

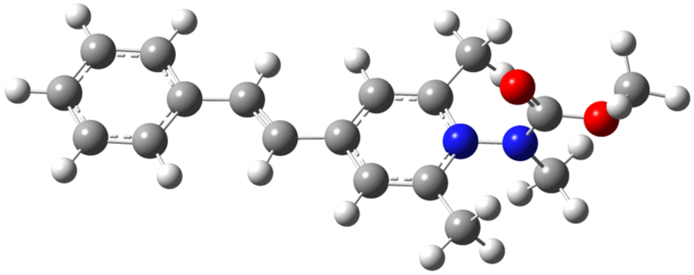

H,0,-3.2328007473,1.3410883485,-2.6787725299
C,0,-2.7213606596,0.5455369507,-2.1334135271
C,0,0.7863151102,3.9311580645,-1.3923957631
C,0,0.127845827,4.9440611541,-2.1153566197
C,0,2.5912502708,5.4999965984,-0.9597579658
C,0,0.6846915451,6.1928519001,-2.2606239132
H,0,-0.8409720266,4.7746641906,-2.5808257319
N,0,1.9048235218,6.4295904021,-1.6873185034
C,0,0.0303663617,7.3018768427,-3.0068650886
H,0,-0.1745809169,8.1534811702,-2.3433429002
H,0,0.6659694722,7.6585746826,-3.828525623
H,0,-0.916867393,6.9469737156,-3.4231735259
C,0,3.8967588389,5.8887887491,-0.3618097129
H,0,4.6032207571,6.2272618383,-1.1315101992
H,0,3.7663863435,6.7072844722,0.3600232651
H,0,4.3258965038,5.027541902,0.1583040767
N,0,2.4675079178,7.6908573144,-1.8393762911
C,0,2.1380505901,8.605264827,-0.8780641045
O,0,1.4004246447,8.3685772965,0.0616756877
C,0,3.3437682837,7.9426644981,-2.9773904414
H,0,3.3961045324,7.0262617359,-3.5730331651
H,0,2.9401670712,8.7520805427,-3.5978614016
H,0,4.3516245331,8.2084684181,-2.635511702
O,0,2.7352551191,9.7636351398,-1.1244079128
C,0,2.4590074629,10.8027895775,-0.179380728
H,0,3.0230912288,11.6702711632,-0.5302624442
H,0,1.386129263,11.0255116573,-0.163946084
H,0,2.797038624,10.5061091871,0.8199399768
C,0,-0.8150181377,-0.2197445198,-0.8585791178
H,0,0.161369951,-0.0458967458,-0.405167475
C,0,2.0299284236,4.2506853336,-0.8188308064
H,0,2.5761324711,3.5037731878,-0.244137619

C,0,-0.9079708924,2.1475490698,-1.7178772082
H,0,-1.5247519703,2.8257247368,-2.3147969192
C,0,0.2600881735,2.5937873249,-1.2073069641
H,0,0.8882928922,1.9328561917,-0.6068452424
C,0,-1.4673594746,0.8121505976,-1.5580567757
C,0,-2.6532463852,-1.718634628,-1.316022804
H,0,-3.110970704,-2.7036621184,-1.2195065594
C,0,-3.3103347979,-0.7072810537,-2.0131864776
H,0,-4.28436922,-0.8957912602,-2.4652973904
C,0,-1.4048325275,-1.4695735651,-0.7409983392
H,0,-0.8887669211,-2.2611628103,-0.1970167493

Zero-point correction= 0.363036 (Hartree/Particle)
Thermal correction to Energy= 0.384618
Thermal correction to Enthalpy= 0.385562
Thermal correction to Gibbs Free Energy= 0.310409
Sum of electronic and zero-point Energies= -957.226341
Sum of electronic and thermal Energies= -957.204759
Sum of electronic and thermal Enthalpies= -957.203815
Sum of electronic and thermal Free Energies= -957.278968

Geometry of **6**:

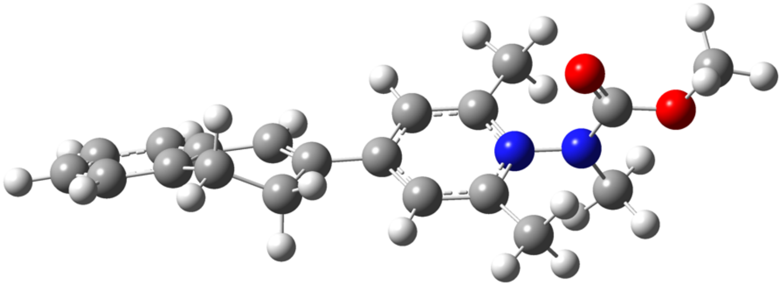

C,0,-3.3193041271,-1.5982707634,-1.5965416602
C,0,-2.6851029413,-0.6212919738,-0.8164175355
C,0,-3.4514265956,0.2179691465,0.019161094
C,0,-4.8311029978,0.0496471864,0.064784277
C,0,-5.4538817441,-0.9343202551,-0.704337433
H,0,-0.7031564471,-1.019657665,-1.6173997522
C,0,-1.2413125889,-0.4667973896,-0.8438210054
C,0,-0.5717166189,0.242760481,0.0976954941
C,0,0.8875710649,0.2793177565,0.1096048884
C,0,1.6533105284,0.0420312079,-1.0476564701

C,0,2.9634645829,0.5676965218,1.3377683714
C,0,3.0281603808,0.0559118777,-1.0145582255
H,0,1.1781350978,-0.1306056948,-2.0113833586
N,0,3.6414575367,0.3027347355,0.1831001981
C,0,3.8828385545,-0.167325517,-2.2123058523
H,0,4.4988425418,0.718139562,-2.4228056953
H,0,4.5577750853,-1.0215196099,-2.0675949107
H,0,3.2452659855,-0.3653848067,-3.0787750051
C,0,3.7471703601,0.8506536638,2.5703453701
H,0,4.4252246504,0.0215683751,2.812660991
H,0,4.3533057804,1.7590121344,2.4463132004
H,0,3.0603652874,1.001424836,3.408041535
N,0,5.0305354066,0.3012935162,0.2207696052
C,0,5.6311562435,1.511825013,0.0138852009
O,0,5.0231884085,2.5430541827,-0.2105239551
C,0,5.7272737121,-0.9482991003,0.5046207332
H,0,4.9772886558,-1.7385911028,0.6067391251
H,0,6.4053292001,-1.2011813262,-0.3193558286
H,0,6.2967586555,-0.8683586375,1.4388463945
O,0,6.9493504786,1.3887067989,0.0933540672
C,0,7.6856404036,2.5998302809,-0.1084261604
H,0,8.7369380332,2.319276979,-0.0089299155
H,0,7.4901621186,2.9999166968,-1.1097421945
H,0,7.4138679899,3.3394158455,0.6531628887
H,0,-5.4286346733,0.6981024183,0.708519268
C,0,-1.3452561725,0.8898632439,1.2236141695
H,0,-0.8113593781,1.7711710871,1.6026821436
H,0,-1.4167316481,0.1799858488,2.0664019155
C,0,-4.6984569684,-1.7574052918,-1.5389560335
H,0,-2.7147826161,-2.2385201383,-2.2421017333
H,0,-5.18638722,-2.5228071397,-2.1425627441
H,0,-6.5364898356,-1.0564500389,-0.6531729901
C,0,-2.7381865259,1.3115222589,0.7664247664
H,0,-2.6389340644,2.181083844,0.0937939557
H,0,-3.3365668757,1.6449317704,1.6241926893
C,0,1.5861841484,0.5601044955,1.2975522598
H,0,1.0589402975,0.7583488065,2.2287680171

Zero-point correction= 0.399319 (Hartree/Particle)
Thermal correction to Energy= 0.421877
Thermal correction to Enthalpy= 0.422821

Thermal correction to Gibbs Free Energy= 0.346918
Sum of electronic and zero-point Energies= -1034.561566
Sum of electronic and thermal Energies= -1034.539008
Sum of electronic and thermal Enthalpies= -1034.538064
Sum of electronic and thermal Free Energies= -1034.613967

Geometry of **7**:

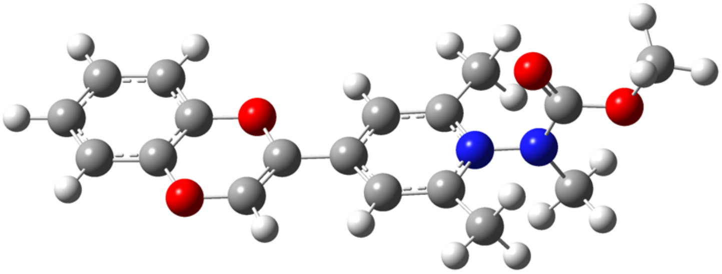

C,0,-3.3193041271,-1.5982707634,-1.5965416602
C,0,-2.6851029413,-0.6212919738,-0.8164175355
C,0,-3.4514265956,0.2179691465,0.019161094
C,0,-4.8311029978,0.0496471864,0.064784277
C,0,-5.4538817441,-0.9343202551,-0.704337433
H,0,-0.7031564471,-1.019657665,-1.6173997522
C,0,-1.2413125889,-0.4667973896,-0.8438210054
C,0,-0.5717166189,0.242760481,0.0976954941
C,0,0.8875710649,0.2793177565,0.1096048884
C,0,1.6533105284,0.0420312079,-1.0476564701
C,0,2.9634645829,0.5676965218,1.3377683714
C,0,3.0281603808,0.0559118777,-1.0145582255
H,0,1.1781350978,-0.1306056948,-2.0113833586
N,0,3.6414575367,0.3027347355,0.1831001981
C,0,3.8828385545,-0.167325517,-2.2123058523
H,0,4.4988425418,0.718139562,-2.4228056953
H,0,4.5577750853,-1.0215196099,-2.0675949107
H,0,3.2452659855,-0.3653848067,-3.0787750051
C,0,3.7471703601,0.8506536638,2.5703453701
H,0,4.4252246504,0.0215683751,2.812660991
H,0,4.3533057804,1.7590121344,2.4463132004
H,0,3.0603652874,1.001424836,3.408041535
N,0,5.0305354066,0.3012935162,0.2207696052
C,0,5.6311562435,1.511825013,0.0138852009
O,0,5.0231884085,2.5430541827,-0.2105239551

C,0,5.7272737121,-0.9482991003,0.5046207332
H,0,4.9772886558,-1.7385911028,0.6067391251
H,0,6.4053292001,-1.2011813262,-0.3193558286
H,0,6.2967586555,-0.8683586375,1.4388463945
O,0,6.9493504786,1.3887067989,0.0933540672
C,0,7.6856404036,2.5998302809,-0.1084261604
H,0,8.7369380332,2.319276979,-0.0089299155
H,0,7.4901621186,2.9999166968,-1.1097421945
H,0,7.4138679899,3.3394158455,0.6531628887
H,0,-5.4286346733,0.6981024183,0.708519268
C,0,-1.3452561725,0.8898632439,1.2236141695
H,0,-0.8113593781,1.7711710871,1.6026821436
H,0,-1.4167316481,0.1799858488,2.0664019155
C,0,-4.6984569684,-1.7574052918,-1.5389560335
H,0,-2.7147826161,-2.2385201383,-2.2421017333
H,0,-5.18638722,-2.5228071397,-2.1425627441
H,0,-6.5364898356,-1.0564500389,-0.6531729901
C,0,-2.7381865259,1.3115222589,0.7664247664
H,0,-2.6389340644,2.181083844,0.0937939557
H,0,-3.3365668757,1.6449317704,1.6241926893
C,0,1.5861841484,0.5601044955,1.2975522598
H,0,1.0589402975,0.7583488065,2.2287680171

Zero-point correction= 0.350146 (Hartree/Particle)
Thermal correction to Energy= 0.372288
Thermal correction to Enthalpy= 0.373232
Thermal correction to Gibbs Free Energy= 0.297691
Sum of electronic and zero-point Energies= -1106.372876
Sum of electronic and thermal Energies= -1106.350734
Sum of electronic and thermal Enthalpies= -1106.349790
Sum of electronic and thermal Free Energies= -1106.425331

Geometry of **8**:

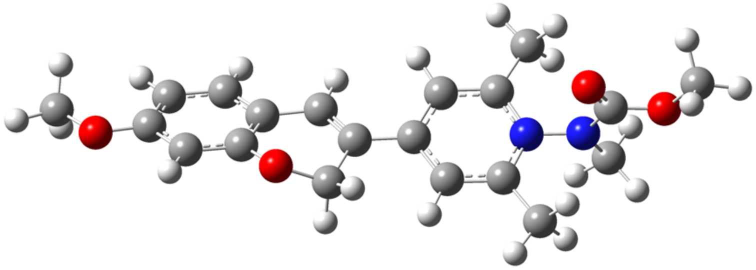

C,0,-3.3561566797,-1.4995494496,-1.7518079109
C,0,-2.6176941592,-0.675102199,-0.8942766228
C,0,-3.3256683375,0.1473191798,0.0071036702
C,0,-4.7056517322,0.1572253274,0.0408040114
C,0,-5.4155599352,-0.6856241996,-0.8241184505
H,0,-0.6623415315,-1.0554980066,-1.7544793917
H,0,-2.8199980691,-2.1343053291,-2.4595151185
C,0,-1.1852916483,-0.5675351318,-0.9297210606
C,0,-0.51947357,0.0965690708,0.0484366771
C,0,0.927666231,0.1862638892,0.0934383896
C,0,1.7320407661,-0.0789498561,-1.0337354582
C,0,2.9645402895,0.5959893245,1.3557903628
C,0,3.1028855459,-0.0184259798,-0.9682031377
H,0,1.2900837632,-0.3154852482,-1.9993647714
N,0,3.6800368957,0.3033190798,0.2312112274
C,0,3.9946180291,-0.2693571269,-2.1331881925
H,0,4.5945287949,0.6218502146,-2.3649351258
H,0,4.6847788113,-1.1005428466,-1.9354770889
H,0,3.3854523077,-0.5185357326,-3.0069276981
C,0,3.7072136437,0.9639283088,2.5910538782
H,0,4.4076113061,0.1719475878,2.8878641934
H,0,4.2848754812,1.8860302807,2.4363284707
H,0,2.995846884,1.129670122,3.4050842939
N,0,5.0660716374,0.3583681624,0.3024526056
C,0,5.627439456,1.5674668805,-0.0005112708
O,0,4.9870937006,2.5562419356,-0.3099129064
C,0,5.803486695,-0.8455957762,0.6674540695
H,0,5.0773865753,-1.6393966874,0.867166951
H,0,6.461794104,-1.1571704714,-0.152933702
H,0,6.3996072759,-0.667916972,1.5707060012
O,0,6.9484768611,1.4961303283,0.0954290823
C,0,7.6449665794,2.7135708931,-0.190960184

H,0,8.7039883351,2.4787345431,-0.0599230283
H,0,7.4486927047,3.0292163311,-1.2218525965
H,0,7.3383630221,3.4991711221,0.5088250518
H,0,-5.2407726227,0.8080915869,0.7302896366
C,0,-1.3355449931,0.6354572693,1.2005173121
H,0,-0.8986224095,1.5501840243,1.6180733859
H,0,-1.4065716269,-0.1116553658,2.0124429477
C,0,-4.7417532232,-1.5253628123,-1.7229916116
H,0,-5.2853580566,-2.1816528594,-2.3981367728
O,0,-2.6520473761,1.0184148946,0.8059192396
O,0,-6.7606537413,-0.6210963278,-0.7145992746
C,0,-7.5376405117,-1.458054159,-1.5550788946
H,0,-7.3177950502,-2.5190698442,-1.3708323209
H,0,-8.5811700194,-1.2523104223,-1.2989143745
H,0,-7.3679011307,-1.2230180894,-2.6152785246
C,0,1.5902494526,0.5359998873,1.2860808766
H,0,1.0407913355,0.7517833992,2.2001649344

| | |
|---|---|
| Zero-point correction= | 0.408459 (Hartree/Particle) |
| Thermal correction to Energy= | 0.433376 |
| Thermal correction to Enthalpy= | 0.434320 |
| Thermal correction to Gibbs Free Energy= | 0.352790 |
| Sum of electronic and zero-point Energies= | -1184.877884 |
| Sum of electronic and thermal Energies= | -1184.852967 |
| Sum of electronic and thermal Enthalpies= | -1184.852023 |
| Sum of electronic and thermal Free Energies= | -1184.933553 |

Geometry of **9**:

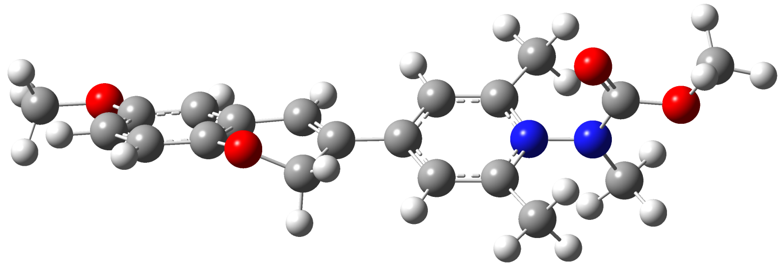

C,0,-3.4157256711,-1.4775616688,-1.648615792
C,0,-2.6795128965,-0.6169777117,-0.8326257155

C,0,-3.3589085153,0.2301147583,0.0608695787
C,0,-4.7416699033,0.2250942449,0.1186203849
C,0,-5.4714799901,-0.6407707502,-0.6966611366
H,0,-0.730107587,-1.0214622586,-1.7191644176
C,0,-1.2384200513,-0.5188120946,-0.8943874949
C,0,-0.5608419896,0.1482735221,0.0679171937
C,0,0.893824794,0.2189937456,0.0983384371
C,0,1.6774752296,-0.0453311354,-1.040166862
C,0,2.9464616064,0.5873571469,1.3422017118
C,0,3.0512608981,-0.0048536705,-0.989411468
H,0,1.219639231,-0.2668108489,-2.0020484241
N,0,3.6434753032,0.2969152629,0.2064482939
C,0,3.9256759757,-0.257535072,-2.1667103494
H,0,4.5289304172,0.6304897704,-2.4019366168
H,0,4.6121194712,-1.0945589042,-1.981522878
H,0,3.3029896134,-0.4976864932,-3.0333387093
C,0,3.7075107446,0.9319422183,2.5726448843
H,0,4.3997687654,0.1271413632,2.8534644743
H,0,4.2960682918,1.8476249473,2.4207388034
H,0,3.0073280471,1.0990862718,3.3959912108
N,0,5.0315112684,0.3282635616,0.2629742293
C,0,5.6089791519,1.5343250197,-0.0230550941
O,0,4.9809506488,2.5364773595,-0.3134510744
C,0,5.7514104837,-0.8904541375,0.6144331441
H,0,5.0144107711,-1.6793494721,0.7929212004
H,0,6.4126513297,-1.1945100576,-0.2062060712
H,0,6.3418585986,-0.7359736701,1.5257174891
O,0,6.9288662035,1.44183901,0.0646388475
C,0,7.6423175468,2.6522612456,-0.2105355681
H,0,8.6983367126,2.3994287083,-0.0893038832
H,0,7.4443685465,2.9839733416,-1.2360399061
H,0,7.3521241591,3.4332159284,0.5013102112
H,0,-5.2528421849,0.901490853,0.8030860773
C,0,-1.3577737537,0.7236997493,1.2169487004
H,0,-0.8916244623,1.6313487013,1.6181607071
H,0,-1.4396651858,-0.0124321779,2.0384777268
C,0,-4.8069953919,-1.4991095857,-1.5780472417
O,0,-2.6596444751,1.1329932085,0.8143064059
H,0,-2.9043191018,-2.1388548675,-2.3492716576
H,0,-6.5577813432,-0.6351347444,-0.6284710242
O,0,-5.432917686,-2.3813107693,-2.4089633462

C,0,-6.8466235399,-2.4169427337,-2.3715264182
H,0,-7.2796350123,-1.4459142037,-2.6541114318
H,0,-7.1494957747,-3.1743625153,-3.101201891
H,0,-7.2138757382,-2.7047191556,-1.3752758008
C,0,1.5693351448,0.547240596,1.286761287
H,0,1.0313383696,0.7618899037,2.2080884238

Zero-point correction= 0.408500 (Hartree/Particle)
Thermal correction to Energy= 0.433434
Thermal correction to Enthalpy= 0.434378
Thermal correction to Gibbs Free Energy= 0.352505
Sum of electronic and zero-point Energies= -1184.873131
Sum of electronic and thermal Energies= -1184.848197
Sum of electronic and thermal Enthalpies= -1184.847253
Sum of electronic and thermal Free Energies= -1184.929126

Geometry of **10**:

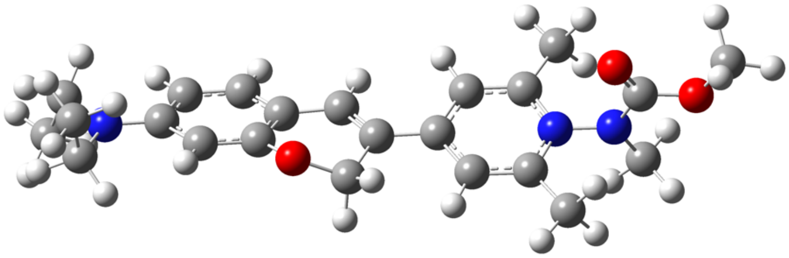

C,0,-3.1727936854,-2.3219402671,-1.0956405852
C,0,-2.4742202507,-1.2008821808,-0.607033092
C,0,-3.2499668079,-0.090302873,-0.2048245951
C,0,-4.6244560641,-0.0810980557,-0.2877233224
C,0,-5.3181999036,-1.2204845405,-0.7708070007
H,0,-0.4801680356,-1.8749758825,-1.0762321265
C,0,-1.0563573617,-1.0864798427,-0.5874003539
C,0,-0.4462457057,-0.035293923,0.0377902658
C,0,0.9825776507,0.1160704171,0.1134280703
C,0,1.8721680308,-0.6408012079,-0.6861493545
C,0,2.933899119,1.1795210368,1.1136910799
C,0,3.2319290132,-0.4995419349,-0.5864263029
H,0,1.5023204585,-1.3447199525,-1.4283354852
N,0,3.7302448215,0.3989751258,0.3227925332

C,0,4.2020002294,-1.259120934,-1.422484969
H,0,4.8005800167,-0.5792311232,-2.0447872784
H,0,4.8935487621,-1.8436193353,-0.8006358823
H,0,3.6560856652,-1.943134399,-2.0786229066
C,0,3.5890642877,2.1393994343,2.043162141
H,0,4.2680776966,1.625070545,2.736480937
H,0,4.1758297162,2.8832125648,1.4860954706
H,0,2.8222250037,2.6620471348,2.62224731
N,0,5.1070058549,0.5418693956,0.4248446892
C,0,5.6700798727,1.4763617814,-0.397529664
O,0,5.0394403811,2.1630000619,-1.181801024
C,0,5.8378577614,-0.2917227041,1.3719325895
H,0,5.1154214375,-0.9376638168,1.8798054125
H,0,6.5722019286,-0.9157844448,0.8477696685
H,0,6.3494832334,0.3308561105,2.116244127
O,0,6.9837619983,1.5195271138,-0.2130271976
C,0,7.6803411505,2.4682336025,-1.0272910848
H,0,8.7323142946,2.3726538042,-0.7476871102
H,0,7.5444336009,2.2298336919,-2.0883038799
H,0,7.320978992,3.4826030986,-0.8202266237
H,0,-5.1446959322,0.8279038672,0.0035717189
C,0,-1.3462087613,0.9152347889,0.7924521165
H,0,-0.9441006233,1.934527411,0.8145901847
H,0,-1.4834057702,0.5796013073,1.8371976934
C,0,-4.5447675998,-2.3503148051,-1.170976561
O,0,-2.6299592238,1.0578664946,0.1842323004
H,0,-2.5997084425,-3.1946937097,-1.4149818689
C,0,1.5710525109,1.0364221883,1.0110427341
H,0,0.9640300763,1.6532182647,1.6702639298
H,0,-5.0331988922,-3.2528373747,-1.5289157967
N,0,-6.6735694843,-1.2337554683,-0.8442217429
C,0,-7.4889915604,-0.145901751,-0.3183192676
H,0,-8.4514310281,-0.5838007548,-0.0179813101
H,0,-7.0343464886,0.2436094933,0.6021197357
C,0,-7.7203778991,0.9694619375,-1.3264471384
H,0,-6.7739145462,1.4346529306,-1.6332805197
H,0,-8.2170809813,0.5825137596,-2.2266518803
H,0,-8.3603966305,1.7478880122,-0.8913789034
C,0,-7.4089936821,-2.337193965,-1.4509201735
H,0,-8.3428708394,-1.9166446871,-1.8500571053
H,0,-6.8555366233,-2.717752374,-2.3193266294

C,0,-7.7229699154,-3.4561406378,-0.4697873994
H,0,-6.8044974379,-3.9018328453,-0.0646373887
H,0,-8.3181430295,-3.0785940021,0.3729184369
H,0,-8.2986973088,-4.2474980015,-0.9669118908

Zero-point correction= 0.506628 (Hartree/Particle)
Thermal correction to Energy= 0.535896
Thermal correction to Enthalpy= 0.536840
Thermal correction to Gibbs Free Energy= 0.445043
Sum of electronic and zero-point Energies= -1282.748272
Sum of electronic and thermal Energies= -1282.719005
Sum of electronic and thermal Enthalpies= -1282.718060
Sum of electronic and thermal Free Energies= -1282.809857

Geometry of **11**:

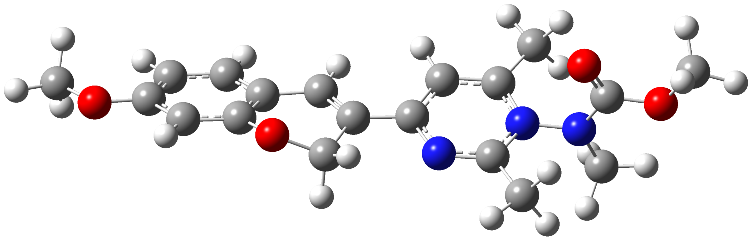

C,0,-3.3504161435,-1.4616645906,-1.7772530741
C,0,-2.5765995231,-0.686323104,-0.9009926161
C,0,-3.2477220249,0.0995929206,0.0639036577
C,0,-4.6258053513,0.1210332294,0.1363896014
C,0,-5.3692148662,-0.6725609051,-0.7468657159
H,0,-0.6606209157,-1.0308518439,-1.8521366429
H,0,-2.8406292593,-2.065034651,-2.5303823411
C,0,-1.1524110188,-0.5824767877,-0.9862516125
C,0,-0.4496263386,0.0457708193,-0.0038045892
C,0,0.9846859644,0.1483068205,-0.0125473752
C,0,1.7794823102,-0.1469106161,-1.1377514618
C,0,2.8670710355,0.6343980921,1.2344753991
C,0,3.1436936107,-0.0737890542,-1.0387372567
H,0,1.3441449784,-0.408601023,-2.0989926121
N,0,3.664595773,0.2783194621,0.1825861023
C,0,4.0736405182,-0.3130683522,-2.1713928421
H,0,4.6516873336,0.5964873991,-2.3877934879
H,0,4.7837675015,-1.1198810853,-1.9497737237

H,0,3.4963379146,-0.584035711,-3.059981804
C,0,3.5071144067,1.127425721,2.4820360212
H,0,4.1886359999,0.3817875125,2.909938261
H,0,4.0853431763,2.0394029815,2.2794302077
H,0,2.7214387293,1.3542540178,3.2063895718
N,0,5.0449295582,0.3295329453,0.3390875326
C,0,5.6208374699,1.5276173016,-0.0014160105
O,0,4.9754911926,2.4849288198,-0.3888048967
C,0,5.6997571481,-0.8946331348,0.7751329282
H,0,5.4970979395,-1.7096030961,0.068842507
H,0,6.7755463983,-0.7138472658,0.8158001239
H,0,5.3442916416,-1.1832317198,1.7721868909
O,0,6.9350147752,1.4950190081,0.1536699838
C,0,7.614349126,2.712237723,-0.1752218229
H,0,8.6715442926,2.5120076206,0.0144666903
H,0,7.4556547848,2.960933635,-1.2305332022
H,0,7.2588179011,3.5287101253,0.463176257
H,0,-5.1345726649,0.7431349381,0.8707764936
C,0,-1.2118433293,0.5258920681,1.2065361917
H,0,-0.7510779906,1.4132124954,1.6509838
H,0,-1.2511897409,-0.2627112597,1.979100176
C,0,-4.7319913428,-1.4764272222,-1.7068554746
H,0,-5.3044633441,-2.0932460442,-2.3951175733
O,0,-2.5458739094,0.9224210509,0.8834818325
O,0,-6.7065812205,-0.6004109243,-0.5956000391
C,0,-7.5211906204,-1.3889038439,-1.4487481953
H,0,-7.3083202287,-2.4592907326,-1.3202186736
H,0,-8.5527923766,-1.1830407102,-1.1487163154
H,0,-7.3820306766,-1.1055300917,-2.5013397989
N,0,1.563202146,0.5640322724,1.1451034783

Zero-point correction= 0.396416 (Hartree/Particle)
Thermal correction to Energy= 0.420286
Thermal correction to Enthalpy= 0.421230
Thermal correction to Gibbs Free Energy= 0.342349
Sum of electronic and zero-point Energies= -1200.926619
Sum of electronic and thermal Energies= -1200.902748
Sum of electronic and thermal Enthalpies= -1200.901804
Sum of electronic and thermal Free Energies= -1200.980685

### 2. Estimation of the excited state oxidation potentials

The pyridinium and pyrimidinium salt structures used in this study undergo rapid and irreversible degradation upon single electron reduction, which makes experimental $E_{1/2}$ reduction half potential measurements impractible with available resources. Thus, estimation of excited state redox potentials were estimated as described previously by Nicewicz[3]. Ground state geometries of each probe studied were calculated using UMN12SX/6-31G(d)/SMD=$H_2O$. This is a modification of the original Nicewicz approach, which used B3LYP. We opted for this theory as we assessed it to provide electronic energies that were more representative of experimental data. Frequency calculations at the same level of theory revealed no negative frequencies of either structure.

Estimation of photo-oxidation potential:

E*([**pyridinium**$^+$]*/**pyridinium•**)=$E_{red}$(**pyridinium**$^+$/**pyridinium•**)+$E_{0,0}$

Where $E_{red}$ is the measured $E_{1/2}$ vs. SCE of the analyte, and $E_{0,0}$ is the intersect point of the normalized absorption/emission spectra.

Since the structures in this study undergo rapid and irreversible fragmentation upon SET, an experimental $E_{1/2}$ was not obtainable. We thus opted to use the method of Nicewicz to estimate the photooxidation potentials. Ground state geometries of each probe studied were calculated using UMN12SX/6-31G(d)/SMD=$H_2O$. This is a modification of the original Nicewicz approach, which used B3LYP. We opted for this theory as we assessed it to provide electronic energies that were more representative of experimental data. Frequency calculations at the same level of theory revealed no negative frequencies of either structure.

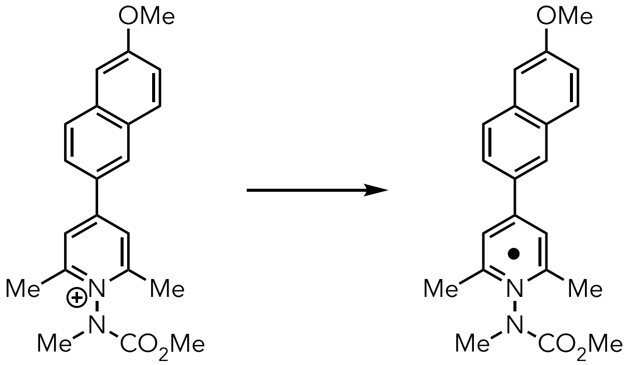

**1** **1•**

| | **1** | **1•** |
|---|---|---|
| G (kcal/mol) | -720506.68 | -720572.85 |
| $\Delta G_{1/2}$ (kcal/mol) | -66.169 | |
| $E_{1/2}$ vs. sce (V) | -1.553 | |
| $E_{0,0}$ (449 nm) | 2.7613 eV | |
| **[1+]*/1•** | +1.209 eV | |

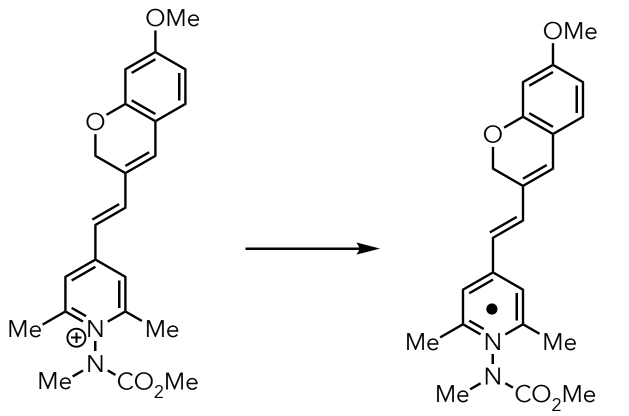

**3** **3•**

| | **3** | **3•** |
|---|---|---|
| G (kcal/mol) | -792300.4 | -792371.791 |
| $\Delta G_{1/2}$ (kcal/mol) | -71.386 | |
| $E_{1/2}$ vs. sce (V) | -1.326 | |
| $E_{0,0}$ (537 nm) | 2.3088 eV | |
| [**3+**]*/**3•** | +0.982 eV | |

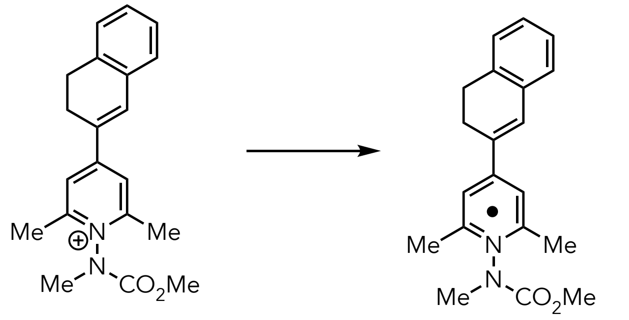

**6** **6•**

| | **6** | **6•** |
|---|---|---|
| G (kcal/mol) | -649437.96 | -649506.86 |
| $\Delta G_{1/2}$ (kcal/mol) | -68.900 | |
| $E_{1/2}$ vs. sce (V) | -1.434 | |
| $E_{0,0}$ (430nm) | 2.8834 eV | |
| [**6+**]*/**6•** | +1.449 eV | |

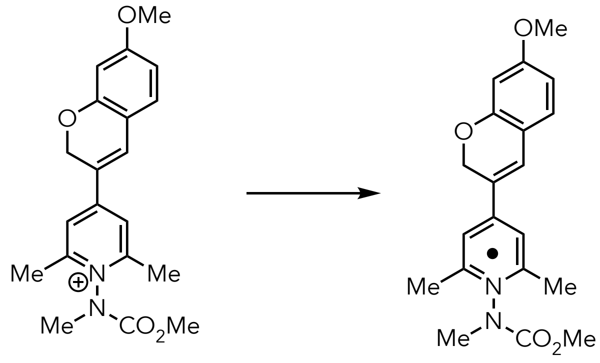

**8** **8•**

| | **8** | **8•** |
|---|---|---|
| G (kcal/mol) | -743767.18 | -743836.47 |
| $\Delta G_{1/2}$ (kcal/mol) | -69.285 | |
| $E_{1/2}$ vs. sce (V) | -1.418 | |
| $E_{0,0}$ (486nm) | 2.5511 eV | |
| **[8+]***/**8•** | +1.134 eV | |

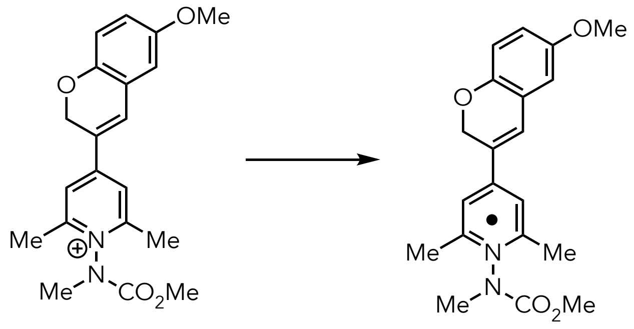

**9** **9•**

| | **9** | **9•** |
|---|---|---|
| G (kcal/mol) | -743764.22 | -743835.9034 |
| $\Delta G_{1/2}$ (kcal/mol) | -71.680 | |
| $E_{1/2}$ vs. sce (V) | -1.314 | |
| $E_{0,0}$ (506nm) | 2.4503 eV | |
| **[9+]***/**9•** | +1.137 eV | |

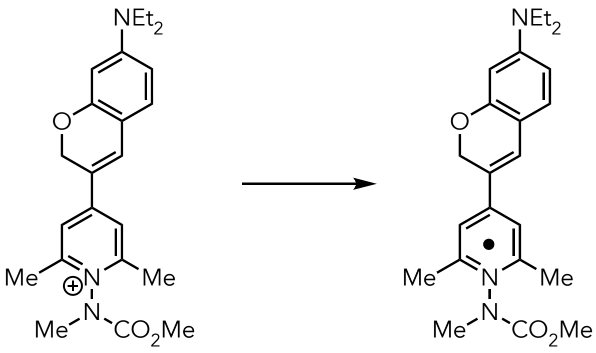

**10** **10•**

| | **10** | **10•** |
|---|---|---|
| G (kcal/mol) | -805242.45 | -805308.37 |
| $\Delta G_{1/2}$ (kcal/mol) | -65.920 | |
| $E_{1/2}$ vs. sce (V) | -1.564 | |
| $E_{0,0}$ (591nm) | 2.0979 eV | |
| **[10+]*/10•** | +0.5339 eV | |

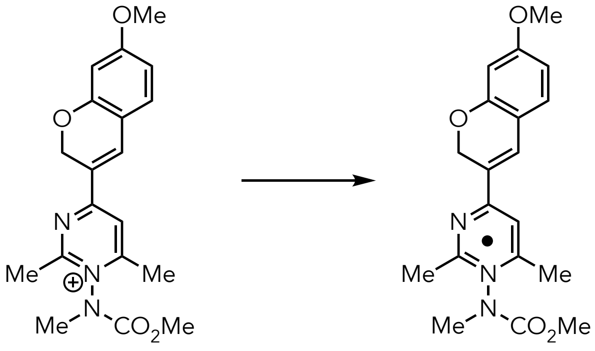

**11** **11•**

| | **11** | **11•** |
|---|---|---|
| G (kcal/mol) | -753830.2 | -753905.13 |
| $\Delta G_{1/2}$ (kcal/mol) | -74.925 | |
| $E_{1/2}$ vs. sce (V) | -1.173 | |
| $E_{0,0}$ (529 nm) | 2.3437 eV | |
| **[11+]*/11•** | +1.171 eV | |

### 3. Fluorophore Volume Calculations

Fluorophore volume calculations were performed on the chromophore component of probes **1**, **6**, **8**, **9**, **10**, and **11** by performing ground state geometry minimization calculations on their *aldehyde analogs* shown in Figure CS2. Calculations were performed at the B3LYP/6-31G(d)/SMD=$H_2O$ level of theory level of theory and volumes were estimated using the "Volume" command in Gaussview. Frequency calculations were performed on all structures to ensure that each calculated structure represents a true energetic minima.

**Core chromophore structures**

| 1 | 6 | 8 | 9 | 10 | 11 |
|---|---|---|---|---|---|

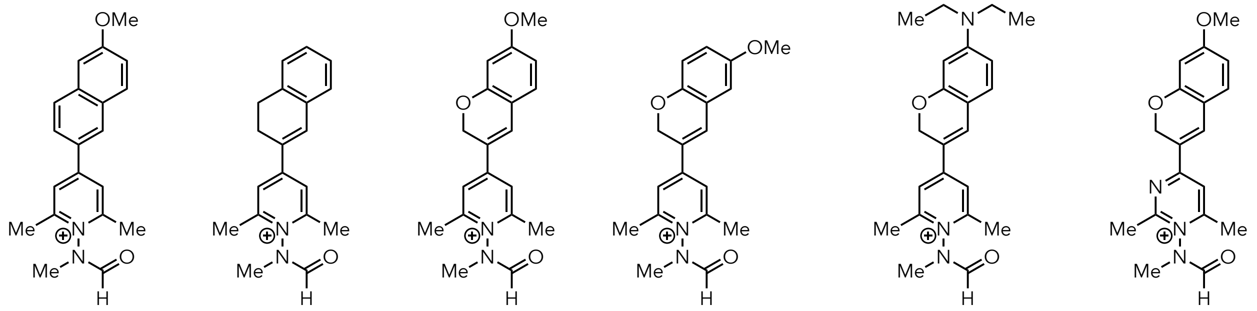

<u>Volumes (Å$^3$)</u>

| 1 | 6 | 8 | 9 | 10 | 11 |
|---|---|---|---|---|---|
| 395 | 396 | 390 | 452 | 433 | 352 |

**Figure CS2.** Calculated volumes of the aromatic cation core chromophores of **1**, **6**, **8**, **9**, **10**, **11**.

### 4. Excited State Calculations

Excited state calculations were performed on **1-11** using optimized ground state geometries that were calculated using B3LYP/6-31G(d)/SMD=$H_2O$ level of theory. Single point time dependent DFT (TDDFT) calculations were performed for S1 with 6 electronic transitions at two different levels of theory: CAMB3LYP/6-311+G(d,p)/SMD=$H_2O$, and ωB97xd/6-311+G(d,p)/SMD=$H_2O$.

#### Pyridinium Probe 2

Excitation energies and oscillator strengths:

```
Excited state symmetry could not be determined.
Excited State   1:      Singlet-?Sym    2.8523 eV  434.68 nm  f=1.4983  <S**2>=0.000
     81 -> 85       -0.10506
     84 -> 85        0.69157
This state for optimization and/or second-order correction.
Total Energy, E(TD-HF/TD-DFT) =  -1014.35215640
Copying the excited state density for this state as the 1-particle RhoCI density.
```

```
Excited state symmetry could not be determined.
Excited State   2:      Singlet-?Sym    4.3560 eV  284.63 nm  f=0.0549  <S**2>=0.000
     83 -> 85        -0.25358
     84 -> 86        -0.57097
     84 -> 89         0.29670

Excited state symmetry could not be determined.
Excited State   3:      Singlet-?Sym    4.7481 eV  261.12 nm  f=0.1325  <S**2>=0.000
     82 -> 85         0.62296
     83 -> 85         0.17042
     84 -> 89         0.23798

Excited state symmetry could not be determined.
Excited State   4:      Singlet-?Sym    4.8411 eV  256.11 nm  f=0.0345  <S**2>=0.000
     82 -> 85        -0.11726
     83 -> 85         0.59887
     84 -> 86        -0.31179

Excited state symmetry could not be determined.
Excited State   5:      Singlet-?Sym    5.2431 eV  236.47 nm  f=0.4068  <S**2>=0.000
     81 -> 86        -0.12139
     82 -> 85        -0.27167
     83 -> 85         0.17296
     84 -> 86         0.22463
     84 -> 89         0.56468

Excited state symmetry could not be determined.
Excited State   6:      Singlet-?Sym    5.2823 eV  234.72 nm  f=0.0551  <S**2>=0.000
     84 -> 87         0.43514
     84 -> 88         0.47560
     84 -> 96         0.13600
     84 -> 98         0.12244
     84 ->100         0.12438
     84 ->101        -0.11663
```

**Pyridinium Probe 3**

Excitation energies and oscillator strengths:

Excited state symmetry could not be determined.
Excited State 1: Singlet-?Sym 2.0189 eV 614.12 nm f=1.8097 <S**2>=0.000
101 ->102 -0.69581
This state for optimization and/or second-order correction.
Total Energy, E(TD-HF/TD-DFT) = -1262.97466005
Copying the excited state density for this state as the 1-particle RhoCI density.

Excited state symmetry could not be determined.
Excited State 2: Singlet-?Sym 3.8050 eV 325.84 nm f=0.1695 <S**2>=0.000
100 ->102 -0.60883
100 ->103 -0.13522
101 ->103 0.28583

Excited state symmetry could not be determined.
Excited State 3: Singlet-?Sym 4.2061 eV 294.77 nm f=0.0217 <S**2>=0.000
99 ->102 -0.37841
100 ->102 0.24625
100 ->103 0.13185
101 ->103 0.47966
101 ->105 0.13659

Excited state symmetry could not be determined.
Excited State 4: Singlet-?Sym 4.5031 eV 275.33 nm f=0.0972 <S**2>=0.000
98 ->102 -0.52654
98 ->103 0.17381
99 ->102 -0.20758
101 ->103 -0.15891
101 ->104 0.30375

Excited state symmetry could not be determined.
Excited State 5: Singlet-?Sym 4.5450 eV 272.79 nm f=0.0791 <S**2>=0.000
98 ->102 -0.20445
99 ->102 0.48166
99 ->103 0.10635
100 ->102 0.12032
101 ->103 0.37138
101 ->104 0.15519

Excited state symmetry could not be determined.
Excited State 6: Singlet-?Sym 4.7806 eV 259.35 nm f=0.0830 <S**2>=0.000
98 ->102 0.34810
99 ->104 0.11528
101 ->104 0.56627

**Pyridinium Probe 4**

Excitation energies and oscillator strengths:

Excited state symmetry could not be determined.
Excited State 1: Singlet-?Sym 2.2228 eV 557.79 nm f=1.2647 <S**2>=0.000
113 ->114 -0.69997
This state for optimization and/or second-order correction.
Total Energy, E(TD-HF/TD-DFT) = -1560.66337001
Copying the excited state density for this state as the 1-particle RhoCI density.

Excited state symmetry could not be determined.
Excited State 2: Singlet-?Sym 3.8092 eV 325.49 nm f=0.1844 <S**2>=0.000
112 ->114 0.65524
113 ->115 0.21964

Excited state symmetry could not be determined.
Excited State 3: Singlet-?Sym 4.1411 eV 299.40 nm f=0.4242 <S**2>=0.000
110 ->114 -0.12842
112 ->114 0.19341
112 ->115 0.12001
113 ->115 -0.64330

Excited state symmetry could not be determined.
Excited State 4: Singlet-?Sym 4.4489 eV 278.69 nm f=0.2758 <S**2>=0.000
110 ->114 -0.10880
110 ->115 0.15556
111 ->114 -0.59499
113 ->116 0.26958

Excited state symmetry could not be determined.
Excited State 5: Singlet-?Sym 4.5284 eV 273.79 nm f=0.0843 <S**2>=0.000
110 ->114 -0.59525
111 ->115 0.16025

112 ->114 -0.11788
113 ->115 0.10625
113 ->117 -0.22555

Excited state symmetry could not be determined.
Excited State 6: Singlet-?Sym 4.8499 eV 255.64 nm f=0.0967 <S**2>=0.000
111 ->114 -0.28616
113 ->116 -0.62002

**Pyridinium Probe 5**

Excitation energies and oscillator strengths:

Excited state symmetry could not be determined.
Excited State 1: Singlet-?Sym 2.7081 eV 457.83 nm f=1.4544 <S**2>=0.000
79 -> 80 0.69942
This state for optimization and/or second-order correction.
Total Energy, E(TD-HF/TD-DFT) = -957.802081936
Copying the excited state density for this state as the 1-particle RhoCI density.

Excited state symmetry could not be determined.
Excited State 2: Singlet-?Sym 4.2760 eV 289.96 nm f=0.0764 <S**2>=0.000
78 -> 80 0.65447
78 -> 81 0.15236
78 -> 82 0.11610
79 -> 83 -0.15191

Excited state symmetry could not be determined.
Excited State 3: Singlet-?Sym 4.4302 eV 279.86 nm f=0.2168 <S**2>=0.000
77 -> 80 0.66585
77 -> 81 -0.13195
79 -> 82 -0.10382

Excited state symmetry could not be determined.
Excited State 4: Singlet-?Sym 5.0416 eV 245.92 nm f=0.0538 <S**2>=0.000
75 -> 80 -0.34760
76 -> 80 -0.15090
79 -> 81 -0.52924
79 -> 82 -0.19692

Excited state symmetry could not be determined.

Excited State   5:      Singlet-?Sym    5.0607 eV  244.99 nm  f=0.0066  <S**2>=0.000
76 -> 80        0.64839
76 -> 81       -0.16426
79 -> 81       -0.10886

Excited state symmetry could not be determined.
Excited State   6:      Singlet-?Sym    5.2129 eV  237.84 nm  f=0.1203  <S**2>=0.000
75 -> 80       -0.31154
75 -> 81       -0.10236
77 -> 80       -0.13141
79 -> 81        0.36578
79 -> 82       -0.46650

**Pyridinium Probe 6**

Excitation energies and oscillator strengths:

Excited state symmetry could not be determined.
Excited State   1:      Singlet-?Sym    2.4880 eV  498.33 nm  f=1.1862  <S**2>=0.000
86 -> 87        0.70002
This state for optimization and/or second-order correction.
Total Energy, E(TD-HF/TD-DFT) =  -1035.20080681
Copying the excited state density for this state as the 1-particle RhoCI density.

Excited state symmetry could not be determined.
Excited State   2:      Singlet-?Sym    4.0673 eV  304.84 nm  f=0.1125  <S**2>=0.000
85 -> 87       -0.66277
85 -> 88       -0.10084
85 -> 89       -0.14784
86 -> 90        0.12207

Excited state symmetry could not be determined.
Excited State   3:      Singlet-?Sym    4.4280 eV  280.00 nm  f=0.2235  <S**2>=0.000
84 -> 87        0.66193
84 -> 89       -0.12554
86 -> 88       -0.13112

Excited state symmetry could not be determined.
Excited State   4:      Singlet-?Sym    4.9295 eV  251.51 nm  f=0.0108  <S**2>=0.000
82 -> 87       -0.26555
86 -> 88        0.52599

86 -> 89 0.32321
86 -> 90 0.10141

Excited state symmetry could not be determined.
Excited State 5: Singlet-?Sym 5.0462 eV 245.70 nm f=0.1740 <S**2>=0.000
83 -> 87 0.19938
84 -> 87 -0.16016
86 -> 88 -0.35809
86 -> 89 0.51161
86 -> 90 0.11565

Excited state symmetry could not be determined.
Excited State 6: Singlet-?Sym 5.0740 eV 244.35 nm f=0.0326 <S**2>=0.000
80 -> 87 -0.10325
83 -> 87 -0.62602
83 -> 89 0.15081
86 -> 89 0.20259

**Pyridinium Probe 7**

Excitation energies and oscillator strengths:

Excited state symmetry could not be determined.
Excited State 1: Singlet-?Sym 2.4949 eV 496.95 nm f=0.3978 <S**2>=0.000
86 -> 87 0.68914
86 -> 89 -0.10757
This state for optimization and/or second-order correction.
Total Energy, E(TD-HF/TD-DFT) = -1106.99302266
Copying the excited state density for this state as the 1-particle RhoCI density.

Excited state symmetry could not be determined.
Excited State 2: Singlet-?Sym 4.4835 eV 276.54 nm f=0.4328 <S**2>=0.000
83 -> 87 0.40687
84 -> 87 0.32865
85 -> 87 0.15233
86 -> 88 0.21073
86 -> 89 -0.33281
86 -> 90 0.13676

Excited state symmetry could not be determined.

```
Excited State   3:      Singlet-?Sym    4.5100 eV  274.91 nm  f=0.3239  <S**2>=0.000
     83 -> 87        -0.37964
     84 -> 87         0.18898
     85 -> 87         0.33697
     85 -> 89        -0.11892
     86 -> 88        -0.20891
     86 -> 89        -0.31075
     86 -> 90        -0.14658

Excited state symmetry could not be determined.
Excited State   4:      Singlet-?Sym    4.5899 eV  270.12 nm  f=0.3042  <S**2>=0.000
     83 -> 87        -0.34169
     84 -> 87         0.12364
     84 -> 90         0.11132
     85 -> 87        -0.12568
     85 -> 89         0.13050
     86 -> 88         0.15965
     86 -> 90         0.52674

Excited state symmetry could not be determined.
Excited State   5:      Singlet-?Sym    4.8022 eV  258.18 nm  f=0.1097  <S**2>=0.000
     84 -> 87        -0.32394
     85 -> 87        -0.31287
     86 -> 87        -0.11934
     86 -> 88         0.10427
     86 -> 89        -0.48610

Excited state symmetry could not be determined.
Excited State   6:      Singlet-?Sym    4.8840 eV  253.86 nm  f=0.1452  <S**2>=0.000
     83 -> 87        -0.14069
     84 -> 87        -0.21937
     84 -> 88        -0.12572
     85 -> 87         0.33164
     86 -> 88         0.52623
```

**Pyridinium Probe 8**

Excitation energies and oscillator strengths:

Excited state symmetry could not be determined.
Excited State 1: Singlet-?Sym 2.3029 eV 538.39 nm f=1.1961 <S**2>=0.000
94 -> 95 -0.69699
This state for optimization and/or second-order correction.
Total Energy, E(TD-HF/TD-DFT) = -1185.59681392
Copying the excited state density for this state as the 1-particle RhoCI density.

Excited state symmetry could not be determined.
Excited State 2: Singlet-?Sym 3.9164 eV 316.58 nm f=0.2680 <S**2>=0.000
93 -> 95 0.66189
93 -> 97 0.16009

Excited state symmetry could not be determined.
Excited State 3: Singlet-?Sym 4.5171 eV 274.48 nm f=0.1533 <S**2>=0.000
91 -> 95 -0.35697
92 -> 95 0.51917
92 -> 97 -0.11336
94 -> 96 -0.23581

Excited state symmetry could not be determined.
Excited State 4: Singlet-?Sym 4.7393 eV 261.61 nm f=0.1945 <S**2>=0.000
91 -> 95 0.15479
92 -> 95 0.29407
93 -> 95 0.14177
94 -> 96 0.51076
94 -> 97 0.27639

Excited state symmetry could not be determined.
Excited State 5: Singlet-?Sym 4.8729 eV 254.44 nm f=0.3075 <S**2>=0.000
91 -> 95 -0.21956
94 -> 96 0.18926
94 -> 97 -0.38681
94 -> 99 0.47109

Excited state symmetry could not be determined.
Excited State 6: Singlet-?Sym 4.9945 eV 248.24 nm f=0.0057 <S**2>=0.000
91 -> 95 0.20332

93 -> 95 0.10741
94 -> 96 -0.30651
94 -> 97 0.33063
94 -> 99 0.44422

**Pyridinium Probe 9**

Excitation energies and oscillator strengths:

Excited state symmetry could not be determined.
Excited State 1: Singlet-?Sym 2.2021 eV 563.02 nm f=0.7165 <S**2>=0.000
93 -> 95 0.10978
94 -> 95 -0.68643
This state for optimization and/or second-order correction.
Total Energy, E(TD-HF/TD-DFT) = -1185.59368690
Copying the excited state density for this state as the 1-particle RhoCI density.

Excited state symmetry could not be determined.
Excited State 2: Singlet-?Sym 3.4567 eV 358.68 nm f=0.5314 <S**2>=0.000
93 -> 95 -0.67362
94 -> 97 -0.10472

Excited state symmetry could not be determined.
Excited State 3: Singlet-?Sym 4.4360 eV 279.50 nm f=0.2291 <S**2>=0.000
92 -> 95 -0.65956
92 -> 97 0.12381
94 -> 96 -0.11768

Excited state symmetry could not be determined.
Excited State 4: Singlet-?Sym 4.6665 eV 265.69 nm f=0.1026 <S**2>=0.000
93 -> 95 0.13479
94 -> 96 -0.45685
94 -> 97 -0.41589
94 -> 99 -0.25651

Excited state symmetry could not be determined.
Excited State 5: Singlet-?Sym 4.9086 eV 252.58 nm f=0.4093 <S**2>=0.000
89 -> 95 -0.12365
93 -> 97 -0.16302
94 -> 97 0.37253
94 -> 99 -0.50998

94 ->103 0.12068

Excited state symmetry could not be determined.
Excited State 6: Singlet-?Sym 5.0019 eV 247.87 nm f=0.0649 <S**2>=0.000
92 -> 95 0.16158
93 -> 96 0.24615
94 -> 96 -0.43794
94 -> 97 0.34928
94 -> 99 0.23967

**Pyridinium Probe 10**

Excitation energies and oscillator strengths:

Excited state symmetry could not be determined.
Excited State 1: Singlet-?Sym 2.2083 eV 561.45 nm f=1.6097 <S**2>=0.000
106 ->107 0.68894
This state for optimization and/or second-order correction.
Total Energy, E(TD-HF/TD-DFT) = -1283.53871592
Copying the excited state density for this state as the 1-particle RhoCI density.

Excited state symmetry could not be determined.
Excited State 2: Singlet-?Sym 3.9032 eV 317.65 nm f=0.1719 <S**2>=0.000
104 ->107 -0.17038
105 ->107 0.65286
105 ->109 -0.16039

Excited state symmetry could not be determined.
Excited State 3: Singlet-?Sym 4.5003 eV 275.50 nm f=0.0150 <S**2>=0.000
104 ->107 0.55904
105 ->107 0.17361
106 ->108 -0.14084
106 ->109 0.29126
106 ->110 0.15694

Excited state symmetry could not be determined.
Excited State 4: Singlet-?Sym 4.6080 eV 269.06 nm f=0.0380 <S**2>=0.000
103 ->107 0.41310
103 ->109 0.10019
104 ->108 -0.14910

106 ->108 -0.51737

Excited state symmetry could not be determined.
Excited State 5: Singlet-?Sym 4.7881 eV 258.94 nm f=0.1008 <S**2>=0.000
106 ->109 -0.44646
106 ->110 0.50856

Excited state symmetry could not be determined.
Excited State 6: Singlet-?Sym 4.9100 eV 252.52 nm f=0.2045 <S**2>=0.000
103 ->107 0.53232
103 ->109 0.11081
106 ->108 0.40493
106 ->109 0.10450
106 ->110 0.10918

**Pyridinium Probe 11**

Excited state symmetry could not be determined.
Excited State 1: Singlet-?Sym 2.2980 eV 539.54 nm f=1.1956 <S**2>=0.000
94 -> 95 0.69682
This state for optimization and/or second-order correction.
Total Energy, E(TD-HF/TD-DFT) = -1201.60743148
Copying the excited state density for this state as the 1-particle RhoCI density.

Excited state symmetry could not be determined.
Excited State 2: Singlet-?Sym 3.8047 eV 325.87 nm f=0.2985 <S**2>=0.000
93 -> 95 0.67263
93 -> 97 0.15546

Excited state symmetry could not be determined.
Excited State 3: Singlet-?Sym 4.3397 eV 285.70 nm f=0.0000 <S**2>=0.000
88 -> 95 0.19766
89 -> 95 -0.28345
91 -> 95 0.57684
91 -> 97 -0.14813

Excited state symmetry could not be determined.
Excited State 4: Singlet-?Sym 4.6338 eV 267.56 nm f=0.0442 <S**2>=0.000
90 -> 95 -0.13996
92 -> 96 0.15570

94 -> 96 0.65980

Excited state symmetry could not be determined.
Excited State 5: Singlet-?Sym 4.7908 eV 258.80 nm f=0.0539 <S**2>=0.000
90 -> 95 -0.14617
92 -> 95 0.55152
93 -> 95 -0.11472
94 -> 96 -0.12620
94 -> 97 -0.35687

Excited state symmetry could not be determined.
Excited State 6: Singlet-?Sym 5.0927 eV 243.46 nm f=0.0857 <S**2>=0.000
90 -> 95 -0.12112
92 -> 95 0.32531
94 -> 97 0.57828
94 -> 98 -0.13484

### 5. References

(1) Gaussian 16, Revision C.01, Frisch, M. J.; Trucks, G. W.; Schlegel, H. B.; Scuseria, G. E.; Robb, M. A.; Cheeseman, J. R.; Scalmani, G.; Barone, V.; Petersson, G. A.; Nakatsuji, H.; Li, X.; Caricato, M.; Marenich, A. V.; Bloino, J.; Janesko, B. G.; Gomperts, R.; Mennucci, B.; Hratchian, H. P.; Ortiz, J. V.; Izmaylov, A. F.; Sonnenberg, J. L.; Williams-Young, D.; Ding, F.; Lipparini, F.; Egidi, F.; Goings, J.; Peng, B.; Petrone, A.; Henderson, T.; Ranasinghe, D.; Zakrzewski, V. G.; Gao, J.; Rega, N.; Zheng, G.; Liang, W.; Hada, M.; Ehara, M.; Toyota, K.; Fukuda, R.; Hasegawa, J.; Ishida, M.; Nakajima, T.; Honda, Y.; Kitao, O.; Nakai, H.; Vreven, T.; Throssell, K.; Montgomery, J. A., Jr.; Peralta, J. E.; Ogliaro, F.; Bearpark, M. J.; Heyd, J. J.; Brothers, E. N.; Kudin, K. N.; Staroverov, V. N.; Keith, T. A.; Kobayashi, R.; Normand, J.; Raghavachari, K.; Rendell, A. P.; Burant, J. C.; Iyengar, S. S.; Tomasi, J.; Cossi, M.; Millam, J. M.; Klene, M.; Adamo, C.; Cammi, R.; Ochterski, J. W.; Martin, R. L.; Morokuma, K.; Farkas, O.; Foresman, J. B.; Fox, D. J. Gaussian, Inc., Wallingford CT, 2016.

(2) GaussView, Version 6, Dennington, Roy; Keith, Todd A.; Millam, John M. Semichem Inc., Shawnee Mission, KS, 2016.

(3) Roth, H.G.; Romero, N.A.; Nicewicz, D.A. Experimental and Calculated Electrochemical Potentials of Common Organic Molecules for Applications to Single-Electron Redox Chemistry. *Synlett*. **2016**, 27, 714-723.
